# Population structure in Arctic marine forests is shaped by diverse recolonisation pathways and far northern glacial refugia

**DOI:** 10.1101/2020.03.19.999466

**Authors:** Trevor T. Bringloe, Heroen Verbruggen, Gary W. Saunders

**Affiliations:** Centre for Environmental and Molecular Algal Research (CEMAR), Biology Department, University of New Brunswick, P.O. Box 4400, Fredericton, New Brunswick, Canada, E3B 5A3; School of BioSciences, University of Melbourne, Parkville Campus, Victoria, Australia, 3010

**Keywords:** Pleistocene glaciation, Species Distribution Models, Genetics

## Abstract

The Arctic is experiencing a rapid shift towards warmer regimes, calling for a need to understand levels of biodiversity and ecosystem responses to climate cycles. This study examines marine refugial locations during the Last Glacial Maximum in order to link recolonization pathways to patterns of genetic diversity in Arctic marine forests. We present genetic data for 109 species of seaweed to infer community-level patterns, and hindcast species distributions during the Last Glacial Maximum to further pinpoint likely refugial locations. Sequence data revealed contiguous populations extending from the Bering Sea to the Northwest Atlantic, with high levels of genetic diversity in the East Canadian Arctic. One fifth of the species sampled appeared restricted to Arctic waters. Hindcasted species distributions highlighted refugia in the Bering Sea, Northwest Atlantic, South Greenland, and Europe. We hypothesize that Arctic coastal systems were recolonized from many geographically disparate refugia leading to enriched diversity levels in the East Canadian Arctic, with important contributions stemming from northerly refugia likely centered along Southern Greenland. Moreover, we hypothesize these northerly refugia likely played a key role in promoting polar endemic diversity, as reflected by abundant unique population haplotypes and endemic species in the East Arctic.

**Significance Statement:** Our work challenges the existing paradigm that marine Arctic ecosystems are depauperate extensions of southerly (temperate) communities established in the wake of recent glaciation, fundamentally changing how these systems should be viewed and interpreted. We forward novel hypotheses regarding the recent history of Arctic marine systems, particularly with regards to endemism being an integral feature of Arctic biomes, and present a firm framework for future evolutionary research in this system typically viewed as “ecologically immature.”

## Introduction

The Arctic is characterized by a turbulent climatic history and the prospect of further change. Repeated glaciations over the past 2.6 Ma had a lasting impact on biological communities, forcing populations to repeatedly contract and expand with the formation and retreat of ice-sheets (1). Today, warming in the Arctic is significantly exceeding the Northern Hemisphere average, and boreal and temperate regimes are expected to shift northwards as a result (2). A prescient need exists to understand the responses of Arctic marine communities to climate change, a need that will inherently depend on understanding levels of biodiversity, the recent history of Arctic ecosystems, and ultimately the potential for adaptation.

Marine forests are a model system for providing such insight on Arctic marine communities. Marine forests are structurally complex seascapes created by seaweeds, are ubiquitous worldwide, and provide valuable ecosystem and economic services in the forms of habitat, nursery grounds, primary productivity, and harvesting resources (3). Marine forests may also sequester large amounts of carbon (4), giving them a potent role in ocean-based climate change mitigation (5). Arctic marine forests are broadly distributed, with circumpolar species distributions extending from the Pacific through to the Atlantic with large gaps along the Siberian Arctic coastline owing to unsuitable soft substrate (6) (see Fig. S1 for exemplar species). Marine forests in the Arctic can also grow to incredible depths in some locations, particularly off the coasts of Greenland and Svalbard where macroalgae have been recorded as deep as 60 m (7). Annual ice cover is often a limiting factor in the Arctic, preventing a lush intertidal community from flourishing, but nonetheless allowing annuals and hardier flora to take advantage of the short growing season (6). Some species are particularly resilient to these conditions. The kelp *Laminaria solidungula*, for instance, completes nearly all its growth under ice, using the summer months when photosynthetic rate is high to focus exclusively on carbon capture and storage (8).

The presence of species that are finely tuned to marine Arctic conditions raises an interesting paradox, in that Arctic marine forests were historically viewed as entirely derived from southerly European refugia following the Last Glacial Maximum (LGM; 9, 10). Under this view, Arctic marine forests were regarded as an extension of cold-tolerant temperate species, precluding the notion of Arctic endemism. While the notion of survival in refugia south of ice sheets persists, alternative hypotheses regarding the recent origins of Arctic marine forests have been brought forward, emphasizing contributions from the Northern Pacific (11, 12, 13). A few genetic studies suggest there may have been multiple recolonization pathways for marine Arctic communities, and have also revealed cryptic diversity in the Arctic, reviving the notion of Arctic endemism (11, 14, 15). Being foundational to coastal ecosystems, such novel insight derived from Arctic marine forests likely apply across a wide range of taxa, and have the potential to dramatically shift how we view and interpret biodiversity and recent origins in Arctic marine communities, along with their potential to adapt to ongoing climatic changes.

The notion of Arctic marine forests as a depauperate extension of temperate communities is an antiquated view. Genetic signature at odds with the hypothesis of recolonization predominantly out of European refugia and the presence of cryptic diversity in the Arctic challenge researchers to revise our understanding of Arctic marine forests and how they persist through cycles of glaciation. Our objective was to clarify the recent origin of Arctic marine forests, emphasizing a community-level approach by summarizing genetic data results across many species, and using species distribution models to identify likely refugial locations during the LGM. We evaluated several hypotheses, employing two lines of evidence in the forms of genetic surveys (DNA barcoding) and hindcasting species distributions using ecological niche modelling. If Arctic marine populations were recently recolonized from southerly refugia, particularly from Europe, then we should observe a decline in genetic diversity along the recolonization pathway (east to west and south to north). In addition, recolonized areas should be genetically closer to the source basin as compared to conspecifics in alternate basins owing to vicariance during glaciation (i.e. Atlantic vs Pacific). Finally, if refugia were only available in Europe, then hindcasting species distributions should reflect this, that is areas of likely species occurrences lacking persistent ice cover during the LGM should be restricted to Europe. If these hypotheses are rejected, our approach will otherwise yield insight on alternate locations of persistence and the subsequent dynamics of recolonization following glaciation.

## Results

### Analysis of sequence data

In order to infer refugial locations and recolonization pathways in Arctic marine forest assemblages, we surveyed and sequenced DNA barcode markers in specimens from several areas. For analytical purposes, we pooled specimens according to the following geographic distinctions: the Northeast Pacific (British Columbia, Canada, and Washington state, USA); Nome, Bering Sea, Alaska (proxy for Pacific migrants into the Arctic); Beaufort, Arctic Ocean, Alaska (proxy for the West North American Arctic); East Canadian Arctic (Cambridge Bay, Nunavut, through to Nain, Labrador); Northwest Atlantic (Makkovik, Labrador, and southwards, including the Canadian Atlantic provinces and New England States, USA); and the Northeast Atlantic (Europe). In total, 4631 specimen records were amalgamated, 2018 of which were collected during this study (2014-2018), representing 42 species of red algae, 49 brown algae, and 18 green algae, for a total of 109 marine macroalgal species with Arctic populations. Of these, 21 species (19%) were sampled exclusively within the Arctic basin, predominantly from the East Arctic region (Table S1; Fig. S2). The genetic results averaged across 31 species-marker combinations revealed Arctic populations that were typically distinct from conspecifics in the Northeast Atlantic and Northeast Pacific. A Principal Coordinates Analysis (PCoA) of genetic distances grouped populations extending from the Bering Sea through to the Northwest Atlantic, within which the East Arctic fell in the middle (Table 1, Fig. 1). Northeast Pacific and Northwest Atlantic populations featured greater genetic diversity, in contrast to comparable populations in Europe, while genetic diversity was depleted in Nome populations and was lowest in the Beaufort (Table 2). The East Arctic, on the other hand, featured levels of genetic diversity comparable to the Northwest Atlantic and Northeast Pacific (Table 2). Interestingly, private haplotypes were detected in all the regions, with the lowest occurrences in the Beaufort. Tajima’s D was negative in southerly populations (Northeast Pacific, Northwest Atlantic, and Northeast Atlantic), and was 0 on average for northern populations (Nome, Beaufort, East Arctic). Of the variables analysed, significant differences were detected across the regions in the number of polymorphic sites, the number of haplotypes, the number of private haplotypes, and Tajima’s D, generally corresponding to large differences in values between the Beaufort and the Northwest Atlantic. Post hoc tests, however, were unable to detect differences between specific regions (Table S2).

**Table 1.** Genetic distances (Φ_ST_) between pairwise populations in 26 species of marine macroalgae with Arctic populations. Genetic distances were evaluated based on four genetic markers (COI-5P, *tufA*, ITS, *ycf35*). Note, the pairwise value between Beaufort and the Northeast Pacific was “normalized” by taking the average of genetic distances between Nome and the NE Pacific, and the East Arctic and NE Pacific. Sample sizes for pairwise distances are in the top right corner of the table. SE for genetic distances are indicated in brackets.

|  | **Northeast Pacific** | **Nome, Alaska** | **Beaufort, Alaska** | **East Arctic** | **Northwest Atlantic** | **Northeast Atlantic** |
| --- | --- | --- | --- | --- | --- | --- |
| **Northeast Pacific** | - | 6 | - | 8 | 8 | 4 |
| **Nome, Alaska** | 0.438(0.158) | - | 11 | 20 | 17 | 5 |
| **Beaufort, Alaska** | 0.471(0.141) | 0.251(0.089) | - | 15 | 12 | 5 |
| **East Arctic** | 0.503(0.124) | 0.300(0.059) | 0.338(0.090) | - | 27 | 11 |
| **Northwest Atlantic** | 0.751(0.101) | 0.518(0.085) | 0.675(0.089) | 0.347(0.063) | - | 11 |
| **Northeast Atlantic** | 0.822(0.141) | 0.741(0.149) | 0.867(0.086) | 0.539(0.109) | 0.437(0.131) | - |

**Table 2.** Summary statistics for populations of Arctic marine macroalgae with Arctic populations. *n* = sample size (number of species), bp = number of basepairs, N_poly_ = number of polymorphic nucleotide sites, Na = number of haplotypes, Ne = number of effective alleles, N_PH_ = number of private haplotypes, *h* = haplotype diversity, θ_π_ = nucleotide diversity, *D* = Tajima’s test for neutralilty, *p*= Kruskal-Wallis tests for independent distributions of values (Dunn’s post hoc tests with Bonferroni correstion did not detect disgnificant pairwise differences); NE=Northeast, NW=Northwest. Note sample sizes from NE Pacific to NE Atlantic in N_PH_ are 10, 20, 16, 31, 29, 13, and from NE Pacific to Overall in *D*, 6, 15, 5, 23, 19, 5, 31. Values for 1 SE are presented below values for each variable.

| Location | NE Pacific | Nome | Beaufort | East Arctic | NW Atlantic | NE Atlantic | Overall | $p$ |
| --- | --- | --- | --- | --- | --- | --- | --- | --- |
| $n$ | 8 | 20 | 15 | 31 | 27 | 11 | 31 | |
| $bp$ | 649 | 639 | 625 | 631 | 632 | 593 | 631 | |
| $N_{poly}$ | 5.375 | 1.600 | 1.000 | 4.387 | 4.444 | 1.727 | 9.516 | **0.021** |
|  | 2.187 | 0.461 | 0.390 | 0.931 | 0.894 | 0.740 | 1.433 |  |
| $N_a$ | 4.000 | 2.300 | 1.933 | 2.935 | 4.148 | 2.273 | 8.097 | **0.041** |
|  | 1.102 | 0.242 | 0.330 | 0.321 | 0.574 | 0.702 | 0.995 |  |
| $N_e$ | 1.782 | 1.542 | 1.312 | 1.753 | 1.821 | 1.474 | 2.834 | 0.157 |
|  | 0.334 | 0.115 | 0.136 | 0.141 | 0.192 | 0.248 | 0.265 |  |
| $N_{PH}$ | 2.500 | 1.000 | 0.625 | 1.323 | 2.655 | 1.538 | - | **0.035** |
|  | 0.806 | 0.218 | 0.202 | 0.287 | 0.526 | 0.656 |  |  |
| $h$ | 0.32325 | 0.29515 | 0.14760 | 0.32958 | 0.32674 | 0.20782 | 0.53132 | 0.143 |
|  | 0.09617 | 0.04919 | 0.05954 | 0.04680 | 0.05140 | 0.07419 | 0.04820 |  |
| $\theta_\pi$ | 0.001544 | 0.000744 | 0.000399 | 0.001873 | 0.001383 | 0.000852 | 0.002605 | 0.093 |
|  | 0.000794 | 0.000268 | 0.000181 | 0.000407 | 0.000569 | 0.000345 | 0.000395 |  |
| $D$ | -1.19122 | -0.02702 | 0.01276 | -0.00114 | -1.06532 | -0.60136 | -0.25835 | **0.012** |
|  | 0.35562 | 0.29170 | 0.44359 | 0.28512 | 0.21389 | 0.53250 | 0.21314 |  |

**Fig. 1.**
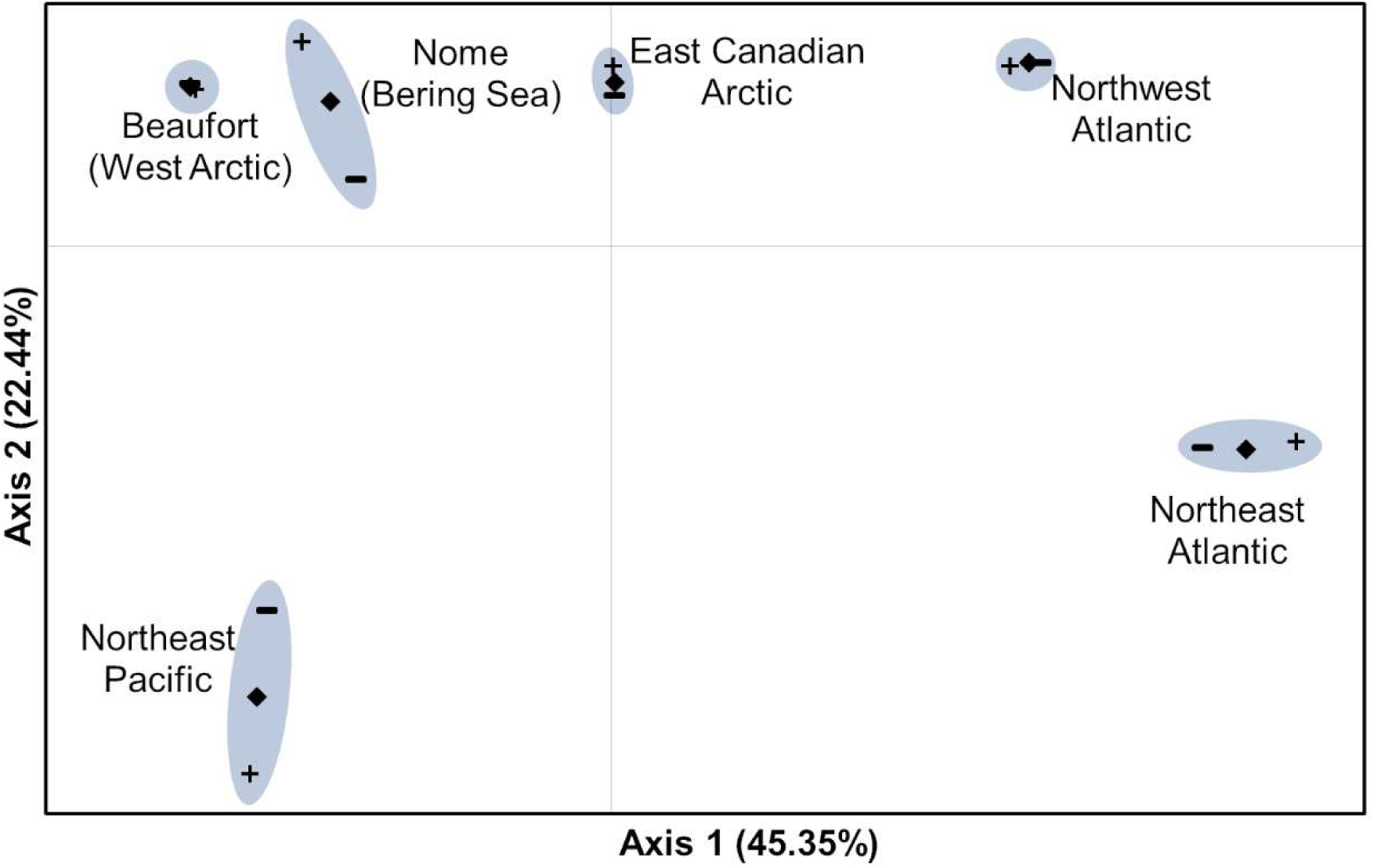
Principal Coordinates Analysis results for genetic distances between the sampled regions. Genetic distances are inferred from 31 species-marker combinations. Positive and negative symbols represent the distributions of coordinates with ± SE.

The presence of private haplotypes in Arctic populations was further highlighted in the haplotype distributions of some species, particularly in the East Arctic. Noteworthy examples included *Alaria esculenta* (Fig. S44 and S45), *Devaleraea ramentacea* (Figs. S13 and S14), *Rhodomela* sp. 1virgata (Fig. S36), and *Pylaiella washingtoniensis* (Fig. S74). Some species showed signs of admixture between North Pacific and North Atlantic populations, particularly in Churchill (Hudson Bay) and Northern Labrador (Northwest Atlantic); these species included *Coccotylus truncatus* (Fig. S12), *Eudesme borealis* (Fig. S58), *Saccharina latissima* (Fig. S76), *Scagelia pylaisaei* (Fig. S39), and possibly *Chaetopteris plumosa* (Fig. S49) and *Phycodrys fimbriata* (Fig. S26). As well, the genetic makeup of several species populations in the Beaufort was markedly different from conspecifics in Nome (Tables 1, 2; Fig. 1), including *Coccotylus truncatus* (Fig. S11 and S12), *Odonthalia dentata* (Fig. S23), *Phycodrys fimbriata* (Fig. S26), and *Rhodomela sibirica* (Figs. S34 and S35).

### Hindcasting species distributions during the Last Glacial Maximum

In order to further pinpoint marine refugial locations, we used species distribution models to hindcast the availability of suitable habitat during the LGM. Hindcasting results differed between Arctic endemic marine forest species (as proxied by the kelp *Laminaria solidungula*) and cold-temperate species ranging into the Arctic (as proxied by the red alga *Odonthalia dentata*). Cold-temperate species featured a present-day circumpolar distribution with large areas of probable occurrence in the North Pacific and North Atlantic. Projections during the LGM similarly indicated areas of probable occurrence throughout the Pacific and Atlantic basins, though highly reduced in total area compared to the present-day projection and particularly limited in South Greenland and the Northwest Atlantic (Fig. 2). Areas of probable occurrence for Arctic endemics for both time projections were more northernly as compared to cold-temperate species, with clear areas of probable occurrence projected in Southern Greenland during the LGM.

**Fig. 2.**
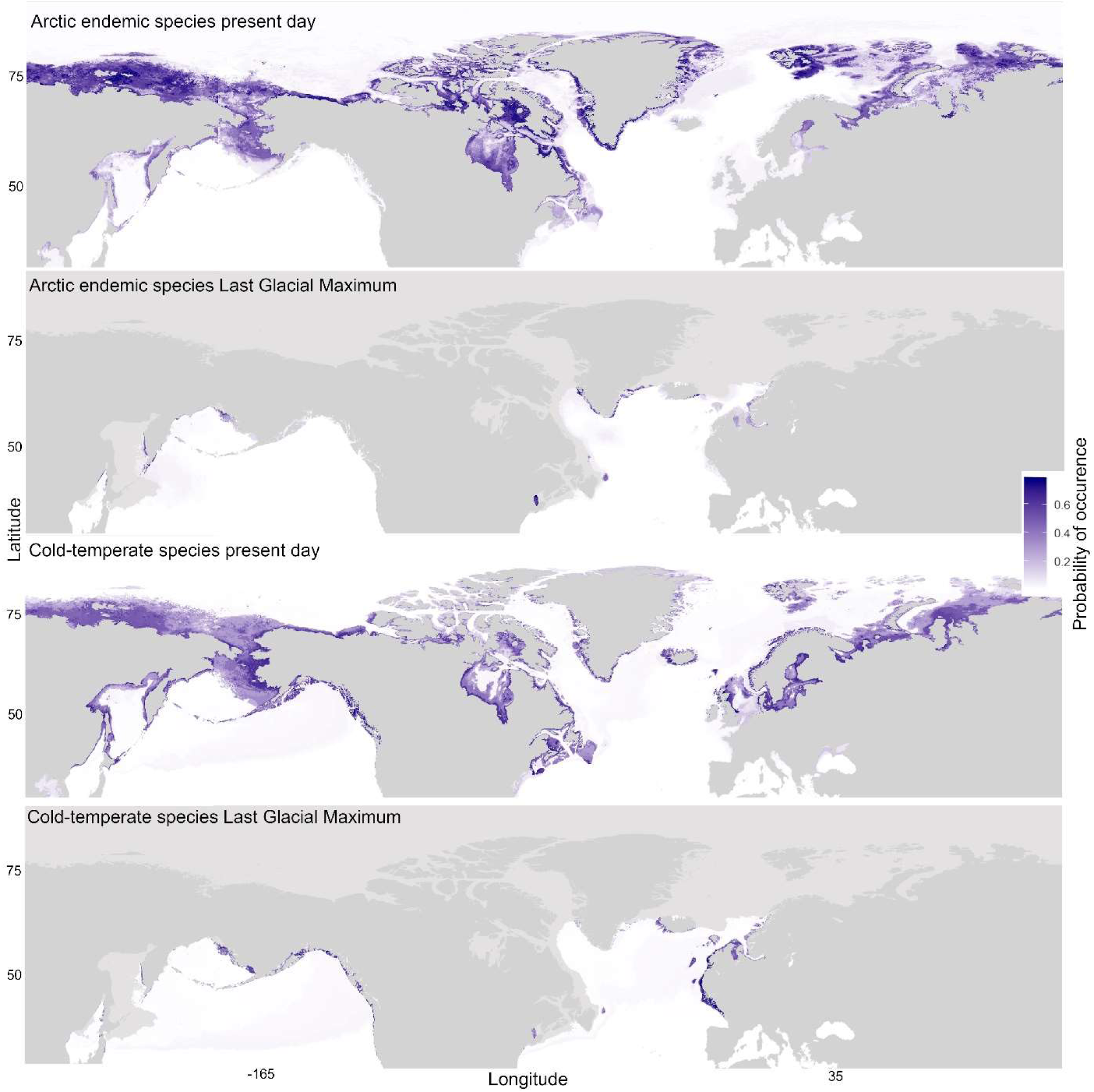
Present day and Last Glacial Maximum distributions of Arctic marine forest species. Endemic species are proxied by the kelp *Laminaria solidungula*, and cold-temperate species ranging into the Arctic are proxied by the red alga *Odonthalia dentata*. The heatmap represents relative probability of occurrence amongst global marine locations as proxied by mean annual sea surface temperatures and temperatures of the coldest ice-free month. Light grey areas represent persistent ice cover during the LGM.

## Discussion

### Recolonization of the Arctic via multiple refugial nuclei

Our understanding of biodiversity patterns in Arctic marine forests has lagged compared to marine animals. The reported similarity of Arctic marine floral assemblages to Atlantic communities was an observation historically at odds with the Pacific origins inferred in Arctic marine fauna, particularly in the Western Arctic (10). Besides noting the “paradox of marine benthic fauna and flora,” Dunton (10) also predicted the importance of molecular data in resolving such disjunct perspectives. Today, with over 4500 sequence records for more than 100 species, the increased fidelity to resolve historical events in populations of Arctic marine forests can shed light on not just recent origins, but also the potential for evolution and northern adaptation in Arctic marine communities.

The results presented here reject the hypothesis that Arctic marine forests are entirely or predominantly derived from European conspecifics (9, 10). This was first evident in the genetic distances, which indicated a series of populations extending from the Northwest Atlantic through to the Northern Bering Sea distinct from the Northeast Atlantic and Northeast Pacific (Fig. 1, Table 1). Furthermore, a decline in diversity from the Atlantic into the Arctic was not evident, which ought to occur in the wake of recolonization (Table 2). In fact, levels of diversity were typically highest in the Northwest Atlantic and maintained in the East Arctic, and Tajima’s D test suggested at a general trend towards population expansion in the Northwest Atlantic, consistent with the presence of refugia (Table 2, Table S3). Hindcasting also showcased areas of probable occurrence during the LGM in the Bering Sea, Southern Greenland, Northwest Atlantic (though highly limited), in addition to Europe (Fig. 2). Finally, haplotype patterns in several species presented here strongly imply the presence of multiple refugia in the Northwest Atlantic (e.g. *Ceramium virgatum*; Fig. S6).

Clearly the European recolonization hypothesis, stemming from a legacy of morphological identifications and the persistent perception of inhospitable glacial conditions in the Northwest Atlantic (9, 10), can be put to rest. For one, refugia in the Northwest Atlantic have been increasingly recognized, downplaying the importance of postglacial recolonization from European refugia (2, 16, 17). Trans-Atlantic populations of marine flora also typically exhibit genetic divergence, corresponding to isolation events during the Pleistocene, further supporting the notion that an abundant marine flora was available to recolonize the Artic out of the Northwest Atlantic (18). Assis *et al.* (2) also used ecological niche modelling to hindcast kelp distributions during the LGM, which suggested several species likely survived glaciation in the Northwest Atlantic, a result further supported in the hindcasting results presented here (Fig. 2). Other modelling work indicated Arctic and sub-Arctic Atlantic marine forests have evolutionary origins in the North Pacific owing to Pleistocene climate and geography, suggesting recurring recolonization occurred out of the Pacific (though not necessarily clarifying origins since the LGM; 13).

Rather, the emerging story of Arctic marine recolonization is more nuanced than anticipated, even in more recent studies. Pacific origins to Arctic flora since the LGM have been speculated in the past (6, 9), interpreted in more recent reviews of floristic surveys (19), and increasingly inferred by molecular data (11; 12). Molecular work has been slow to recognize the significance of unique Arctic lineages that are difficult to reconcile with the emphasis on recolonization hypotheses stemming from refugia south of continental ice-sheets. For instance, Neiva *et al.* (12) report “temperate” and “cold” Northwest Atlantic phylogroups of *Saccharina latissima*, the later of which was largely restricted to the Canadian Arctic and Western Greenland, but invoke backcrossing or genotyping errors as possible explanations for the closely related genetic groups. The hesitation to consider northern refugia as a possible explanation (e.g. Southern Greenland) likely stems from several sources, including a lack of comprehensive geographic coverage in genetic sampling, limited insight from single species, and confusion regarding the extent of seasonal ice cover during the LGM. Here, the position of East Arctic genetic distances between populations sampled in the Bering Sea and Northwest Atlantic (Fig. 1) and sustained levels of genetic diversity in East Arctic populations (Table 2) can be attributed to a combination of admixture between Pacific and Atlantic populations and vicariance, the latter of which invoke the presence of high northern refugia. Hindcasting identified the most likely area of occurrence for these refugia along the coastlines of Southern Greenland where ice cover was seasonal during the LGM (Fig. 2; 20), particularly in species specifically outfitted to thrive in Arctic conditions (e.g. *Laminaria solidungula*). It is worth noting hindcasting results are consistent with the work of Assis *et al.* (2), though these authors also inferred high Arctic refugia in areas we identified as having persistent ice cover (namely Baffin Island and Northern Labrador). In sum, the origin of Arctic marine forests, as revealed by our community level analysis, forwards the West Arctic as likely derived from Bering Sea refugia, while the East Canadian Arctic exhibits a “melting pot” character, with input from the Pacific, the Atlantic (predominantly the Northwest), and Arctic refugia.

### High northern refugia drive Arctic endemism

The conclusion that refugia were abundant and occurred far north offers an explanation for marine Arctic endemism, which is generally assumed to be rare or impossible given the perception of intolerable ice conditions during the LGM. Lee (9) suggested less than 7% of the Arctic flora was confined to the Arctic and perceived low levels of adaptation, characterizing the flora as “ecologically immature.” More recent estimates bump up the level of Arctic endemism to 13.5% (21). Molecular taxonomic work reflects the trend towards recognizing Arctic affinities, for example, in the recently described species *Ahnfeltia borealis, Chorda borealis*, and *Eudesme borealis* (22). Here, 21 species (19%) of the sampled flora appeared to be confined to the Arctic, not to mention the cryptic lineages reported by Laughinghouse *et al.* (14), Küpper *et al.* (15), and species with distinct Arctic populations reported here (e.g. *Alaria esculenta*; Fig. S45 and S46). Altogether, these results suggest that a large portion of the Arctic flora has persisted far north through cycles of glaciation, raising the possibility that Arctic marine forests do not simply tolerate but rather are adapted to polar conditions.

The implications of these findings extend beyond marine forests. Patterns in accompanying hard-bottom marine fauna, some of which thrive in the habitat provided by macroalgae, are readily resolved by invoking northern refugia. A phylogeographic review of Arctic marine fauna highlighted substantial geographical subdivision in the COI-5P complex of the polychaete *Harmothoe imbricata*, with lineages that appeared to be restricted to the East Canadian Arctic (23). The presence of cryptic Arctic lineages was further revealed in polychaetes (24), molluscs (25) and amphipods (26). In concert with the results for Arctic forests, these studies demonstrate the resilience of Arctic marine communities to cycles of glaciation and the potential for adaptation to extreme-cold environments. Given the relative scarcity of genetic surveys in the Arctic, these results also suggest levels of Arctic marine endemism remain underestimated, a disparity that is likely further exacerbated at the population level.

On a final note, it is possible northern refugia are not confined to the Southern coast of Greenland. Though the Western Arctic was locked in multi-year sea ice, portions of the Siberian coastline as far east as the Laptev Sea appear to have remained seasonally ice-free at least as early as 16 ka (27). This was due to warm Atlantic water entering the Arctic through the Fram Strait, which may have provided Arctic refuge for marine forests. The role of katabatic winds and the formation of polynyas, recurrent areas of ice-free water, also may have played a role maintaining a seasonally ice-free shoreline (27). The modern day Arctic features numerous polynyas, particularly along the margins of the Arctic basin, enhancing early spring productivity and creating biodiversity hotspots (28). Lee (9) even describes seaweed communities in a polynya near Brock island (78°N), including *Laminaria solidungula, Desmarestia viridis*, and *Turnerella pennyi*. Hypotheses regarding the role of polynyas maintaining Arctic refugial locations during the LGM may need to be invoked to explain the population structure of the Beaufort, which is oddly more genetically distant from Atlantic conspecifics than Nome populations (which are geographically further away, Fig. 1). We do caution in interpreting this pattern, however, given the limited sampling of the West Arctic/Bering Sea, lack of sampling from Russia, and “founder-takes-all” effects, which can result in sharp demarcations in macroalgal population structure, even at small spatial scales (29). Even so, if history is an indication, we need to temper our assumptions regarding the inability for marine forests to flourish in areas with “harsh” ice conditions.

### Conclusions

Over the decades, our understanding of Arctic marine forests has gradually evolved from a one-vector ecosystem stemming from European refugia to many melding pathways, and from a biologically depauperate expanse into a complex genetic landscape. Ultimately, our goal was to provide a checkpoint in our understanding of Arctic marine diversity, particularly by sampling Arctic locations where DNA barcode surveys are rare or absent (i.e. Bering Sea, East Arctic, Northern Europe) and by summarizing genetic patterns across macroalgal species with sequence data from the Arctic. We emphasize the significance of summarizing insight across species, with over 100 examined through this work spanning three phyla and two kingdoms of life, and extensively so in 26 (Table S1; Fig. S1). Single-species studies have struggled to reconcile post-glacial dynamics, particularly where findings do not align with prevailing views (e.g. 12, 14). As well, a population genetic approach to resolving these patterns in Arctic marine communities oftentimes falls short of conclusive owing the inherent difficulty in sampling any individual location and subsequent limited geographical coverage for analysis. Ecological niche modelling can close these gaps in our knowledge by highlighting likely locations of persistence during glaciation and aiding in the interpretation of genetic patterns.

We forward the view that the marine Arctic environment was recolonized from numerous and globally distributed source populations, including unrecognized far northern refugia that additionally contribute to endemism in polar waters. This work raises the possibility of investigating complex evolutionary processes born out of the Arctic environment, particularly the interplay between incipient speciation and secondary contact, and adaptation to the marine Arctic environment. Ecosystem management will need to acknowledge diversity endemic to the Arctic and the potential impacts of climate change on these populations. Future work should emphasize sampling in Russia where DNA barcodes are virtually absent, and Southern Greenland as a putative refugial location. A consensus must also be reached regarding the extent of persistent sea ice cover during the LGM, and what role, if any, continental ice-sheets had in extirpating marine populations. As well, given the range of challenges and limited opportunity to sample the marine Arctic environment, future collectors should consider sampling protocols amendable to downstream genomic analyses, whatever the primary purpose for Arctic fieldwork. The DNA barcode surveys presented here are a checkpoint towards the genomic era of Arctic investigations, a realm that will certainly reveal novel and exciting insight into levels of biodiversity in the Arctic, and potential for further adaptation and evolution in a changing climate.

## Materials and Methods

### Genetic data and analyses

Marine macroalgae were sampled in several key locations across the Arctic over the course of five years, including Japan; Kamchatka (Russia); Haida Gwaii, British Columbia, Canada; Nome, Alaska (Northern Bering Sea); the Beaufort Sea, Northern Alaska; Cambridge Bay (Nunavut, Canada); Hudson Bay, Manitoba, Canada; Baffin Island, through Northern Labrador to Makkovik, Canada; and Bergen, Norway. Specimen records, including pictures, collection information, sequence data, and GenBank accessions can be accessed via the Barcode of Life Data System (BOLD; doi: dx.doi.org/10.5883/DS-TAMMA) and FigShare: (https://doi.org/10.6084/m9.figshare.11301929.v1). Marine macroalgae were generally collected from the intertidal and via scuba, but occasionally via dredge. A portion of each specimen (approx. 1 cm^2^) was preserved in silica gel for DNA extraction, while several representatives of putative species were preserved as pressed vouchers. Specimens were brought back to the University of New Brunswick (where specimens are stored) for DNA extraction. Several genes were amplified, including the 5’ end of the cytochrome *c* oxidase subunit I gene (COI-5P) in red and brown algae, *tufA* in green algae, and partial reads of the ribulose-1, 5-biphosphate carboxylase large subunit (*rbc*L-3P; Table S4) in red and brown algae. Secondary markers were acquired in select species, including the full length nuclear internal transcribed spacer region (ITS) and plastid *ycf35*, in order to further clarify or support COI-5P patterns (Table S4). Successful PCR products were sent to Genome Quebec for forward and reverse sequencing. All genetic data were edited in Geneious v.8.0 (30), and any relevant previously published DNA barcodes were added to the dataset.

Populations were pooled for analysis according to broad geographic regions: Northeast Pacific (British Columbia, Canada, and Washington state, USA); Nome, Alaska (proxy for Pacific migrants into the Arctic); Beaufort, Alaska (proxy for the West North American Arctic); East Canadian Arctic with the southern distribution delimited using the 10°C air temperature isotherm for July (Cambridge Bay, Nunavut, through to Nain, Labrador); Northwest Atlantic (Makkovik, Labrador, and southwards, including the Canadian Atlantic provinces and New England States, USA); Northeast Atlantic (Europe; Fig. 2). The Northwest Pacific was excluded given the paucity of data. Species generally corresponded to Barcode Index Numbers (BINs), a binning system provided through BOLD which utilizes a fluid threshold to proxy species units based on levels of intra- and inter-specific genetic variation (31). Species were included in genetic analyses provided they featured in at least one Arctic region for which ≥10 individuals were sampled. Out of 109 species sampled with Artic populations, we analysed 26 (Fig. S1), five of which had sequence data from multiple markers, for a total of 31 species-marker combinations. All 109 species were considered in additional analyses evaluating total diversity levels in the Arctic, which were used to quantify the number of Arctic endemic species. These analyses are presented as supplementary material, and include accumulation curves of all genotypes sampled (32; Fig. S2), a table interpreting haplotype patterns in all 109 species sampled (Table S1), and haplotype maps and networks (Figs. S3-S95).

Various populations statistics were calculated for each region, including measures of genetic diversity, genetic differentiation, and Tajima’s D test for neutrality. Sequences were truncated to the shortest length sequence within each species prior to all genetic analyses. GenAlEx 6.51 (33) was used to run an Analysis of Molecular Variance (AMOVA) and derive values of PhiST (Φ_ST_), an analogue to Fst that incorporates nucleotide diversity in distance calculations (i.e. haplotypes are not assumed to be equidistant from each other). Calculations for Φ_ST_ followed that of Miermans (34). Pairwise tests for significant Φ_ST_ values were conducted using 9999 permutations of the dataset. Null values for Φ_ST_ occurred wherein two populations were monotypic for the same haplotype and were changed to 0 (no genetic differentiation). Species specific analyses are presented in Table S5. Φ_ST_ values were averaged for each region across all species, and a Principal Coordinates Analysis (PCoA) was conducted on the pairwise distance matrix using the covariance-standardized method in GenAlEx. Given notably few measurements were available between the Northeast Pacific and the Beaufort (only two species with 2-7 records from the Beaufort, and no genetic differentiation), we “normalized” this pairwise distance by taking the average of genetic distances between the Northeast Pacific and Nome, and the Northeast Pacific and the East Arctic. The unaltered PCoA figure is presented in Figure S96. We also conducted the same analysis with low sample size populations removed (<10 individuals; Fig. S97). Alternate PCoAs revealed the same pattern, except the Beaufort grouped closer to the Northeast Pacific when not “normalized.”

Frequency based parameters were also calculated for each region within each species, including the number of haplotypes (Na), the number of effective alleles (Ne), the number of private haplotypes within populations (N_PH_), and haplotype diversity (*h*). Ambiguous sites were removed for these calculations. DnaSP v6 (35) was also used to calculate the number of polymorphic sites, nucleotide diversity (θ_π_), and Tajima’s D statistic with accompanying *p*-values. These results were again averaged across all species-marker combinations, and differences in measures between populations were assessed using Kruskal-Wallis H tests (36). Dunn’s post hoc tests with Bonferroni corrections were performed for variables yielding significant Kruskal-Wallis H tests. The analyses for individual species are presented in Table S3, and the post hoc tests are presented in Table S2.

### Hindcasting species distributions

The distribution of marine forests during the Last Glacial Maximum was inferred using ecological niche modelling. Modern and paleo marine environmental data layers were downloaded from MARSPEC (37, 38), in particular, bathymetry, mean annual sea surface temperatures (SST) and SST of the coldest ice-free month. Seaweed distributions are highly tuned to marine isotherms, and ecological niche modelling consistently indicate SST is the most important variable in modelling macroalgal species distributions (2). Paleo environmental data layers represent average values as derived from six coupled ocean atmosphere general circulation climatic models, including salinity-adjusted CCSM3. Occurrence records were gathered for *Odonthalia dentata* and *Laminaria solidungula*. These species were selected to proxy distributions in boreal to temperate species with Arctic populations (i.e. the classical view of Arctic marine forests; *O. dentata*), and putative Arctic endemic species (*L. solidungula*). These species were also selected given they are reliably distinguished morphologically, and as such, historical records would not be conflated by cryptic species. Occurrence records were derived from Lüning (6). Distribution maps were scanned and georeferenced, and GPS locations for occurrences were subsequently derived. For continuous distributions, GPS coordinates were haphazardly recorded at approximately 250 km intervals. Historical occurrence records were then pooled with locations from DNA barcode records (BOLD), and the Macroalgal Herbarium Portal (https://macroalgae.org/portal/index.php). To correct for sampling bias during training of the ecological niche models, occurrence records in close proximity to each other were randomly removed using the R package spThin (39), with a thinning parameter of 100 kms. The resulting datasets kept 111/260 occurrence records for *Odonthalia dentata*, and 87/164 for *Laminaria solidungula* (FigShare: https://doi.org/10.6084/m9.figshare.11301929.v1). The thinned occurrence records and environmental layers were then trained using Maxent (40) and projected onto conditions during the LGM. Models were built using threshold features in order to better reflect lethal temperature limits in macroalgae, and model performance was assessed using cross validation. A regularization multiplier of 1 was used, as was a default prevalence of 0.5 (the probability of presence at average presence locations). Clamping was used to restrict variables outside the training range. Multivariate Environmental Similarity Surfaces (MESS) were also used to evaluate the distribution of environmental values outside the training data range projected during the LGM, which functioned to indirectly map persistent ice cover and restrict inferences of refugial locations to seasonally ice-free waters. Output asc files were converted to figures in R using ggmap and ggplot2 packages (39).

## Acknowledgments

We thank the many people who facilitated the collection of specimens: Dr. Meghann Bruce, Dr. Kyatt Dixon, Kirby Morrill, Dr. Amanda Savoie; Dr. Kenneth Dunton at the University of Texas for facilitating our sampling in the Beaufort (Northern Alaska); Dr. Yotsukura Norishige at Hokkaido University for providing collections from Hokkaido; Dr. Kjersti Sjøtun at the University of Bergen for facilitating our sampling in Norway; Don Stiles and James Horner for facilitating the collection of samples in Nome, Alaska; Laura Borden for providing collections from Cambridge Bay (Nunavut); Drs. Selivanova and Zhigadlova for providing specimens from Kamchatka (Russia). We thank those who helped generate COI-5P data, particularly Tanya Moore, as well as Alex Geoffroy and Line Le Gall for providing trace files. Drs. Jorge Assis and Ester Serrão provided critical feedback regarding analyses. Dr. Jason Addison also provided critical input leading to the inception of this manuscript. This project was funded by the Northern Scientific Training Program, the Natural Sciences & Engineering Research Council of Canada through an NSERC Post-Graduate Scholarship to T.T. Bringloe and a Discovery Grant to G.W. Saunders (170151-2013), the New Brunswick Innovation Foundation, and the University of Melbourne McKenzie Postdoctoral Fellowship program.

