## Supplemental material for "Population structure in Arctic marine forests is shaped by diverse recolonisation pathways and far northern glacial refugia"

**This PDF file includes:**

Fig. S1. Marine Arctic forest species used for genetic analyses.
Fig. S2. Accumulation curves for genotypes (haplotypes) in all specimens sampled.
Figs. S3-S95. Haplotype distribution and networks in each species of macroalgae sampled.
Fig. S96. PCoA analysis without "normalizing" Beaufort, Northeast Pacific relationship
Fig. S97. PCoA analysis with low sample populations (<10 individuals) removed from averages.
Table S1. General haplotype patterns and inferred origins in Arctic species of marine macroalgae.
Table S2. Kruskal-Wallis test results with Dunn's post hoc tests with Bonferroni corrections for groups wherein the null hypothesis was rejected.
Table S3. Summary statistics for populations of Arctic marine macroalgae with Arctic populations.
Table S4. Primers used for amplification of various genes in marine macroalgae.
Table S5. Pairwise $\Phi_{ST}$ values for species of macroalgae with Arctic populations.

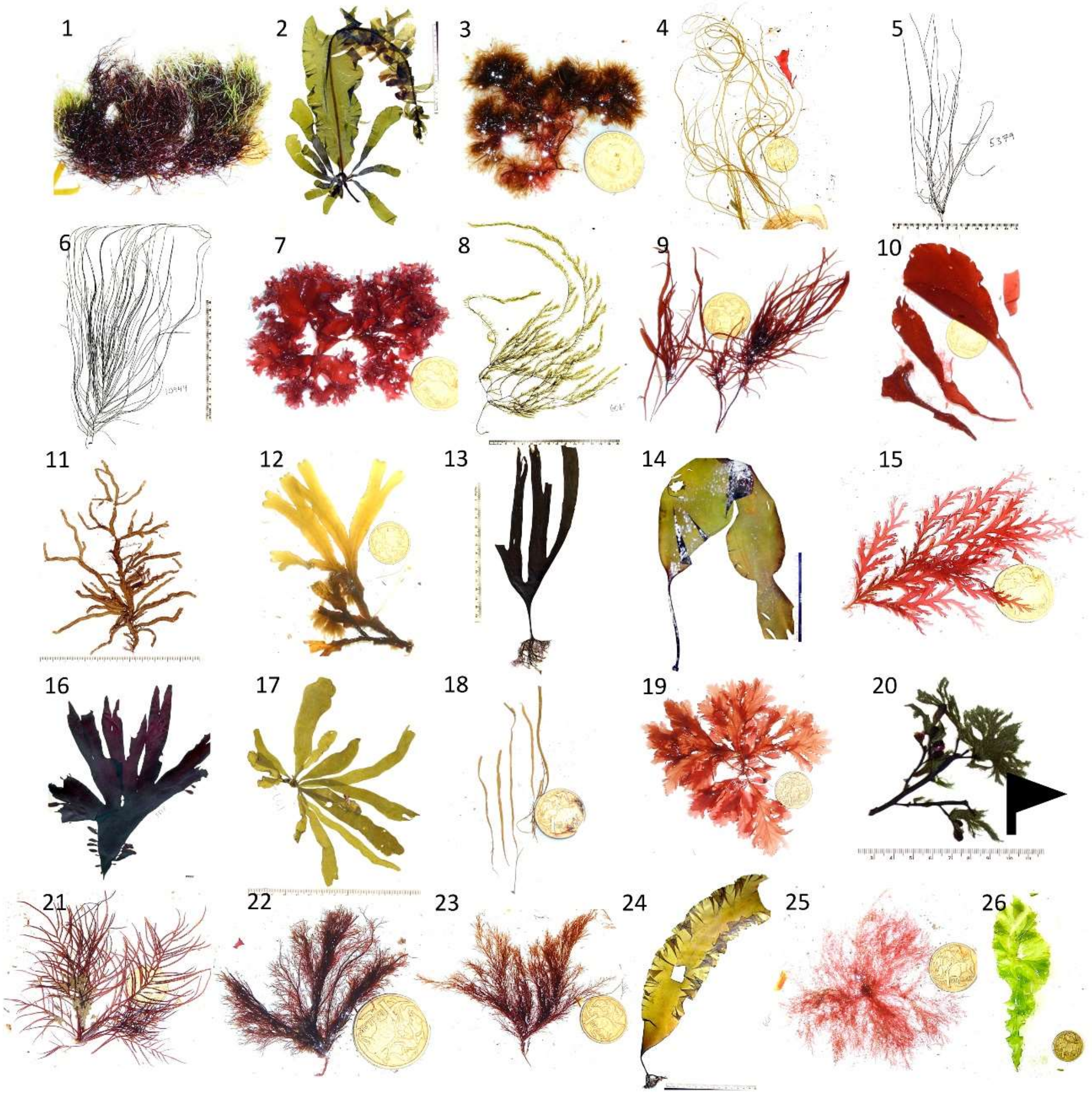

**Figure S1.** Marine Arctic forest species used for genetic analyses. For scale there is an Australian one dollar coin (~2.5 cm) or a centimeter ruler. 1. *Ahnfeltia borealis* (GWS041938); 2. *Alaria esculenta* (GWS005372); 3. *Chaetopteris plumosa* (GWS040974); 4. *Chorda borealis* (GWS040219); 5. *Chordaria chordaeformis* (GWS005379); 6. *Chordaria flagelliformis* (GWS010944); 7. *Coccotylus truncatus* (GWS039645); 8. *Desmarestia* sp. 1aculeata (GWS006068); 9. *Devaleraea ramentacea* (GWS034080). 10. *Dilsea socialis* (GWS041422); 11. *Eudesme borealis* (GWS007814). 12. *Fucus distichus* (GWS041940). 13. *Hedophyllum nigripes* (GWS002500). 14. *Laminaria solidungula* (GWS005422). 15. *Odonthalia dentata* (GWS041454). 16. *Palmaria palmata* (GWS003518). 17. *Petalonia fascia* (GWS003601). 18. *Petalonia filiformis* (GWS040249). 19. *Phycodrys fimbriata* (GWS030218). 20. *Pylaiella washingtoniensis* (epiphyte on *Ascophyllum nodosum*; GWS003682). 21. *Rhodomela sibirica* (GWS042349). 22. *Rhodomela virgata* (GWS042344). 23. *Rhodomela* sp. 1virgata (GWS039418). 24. *Saccharina latissima* (GWS006005). 25. *Scagelia pylaisaei* (GWS039369). 26. *Ulva fenestrata* (GWS036952).

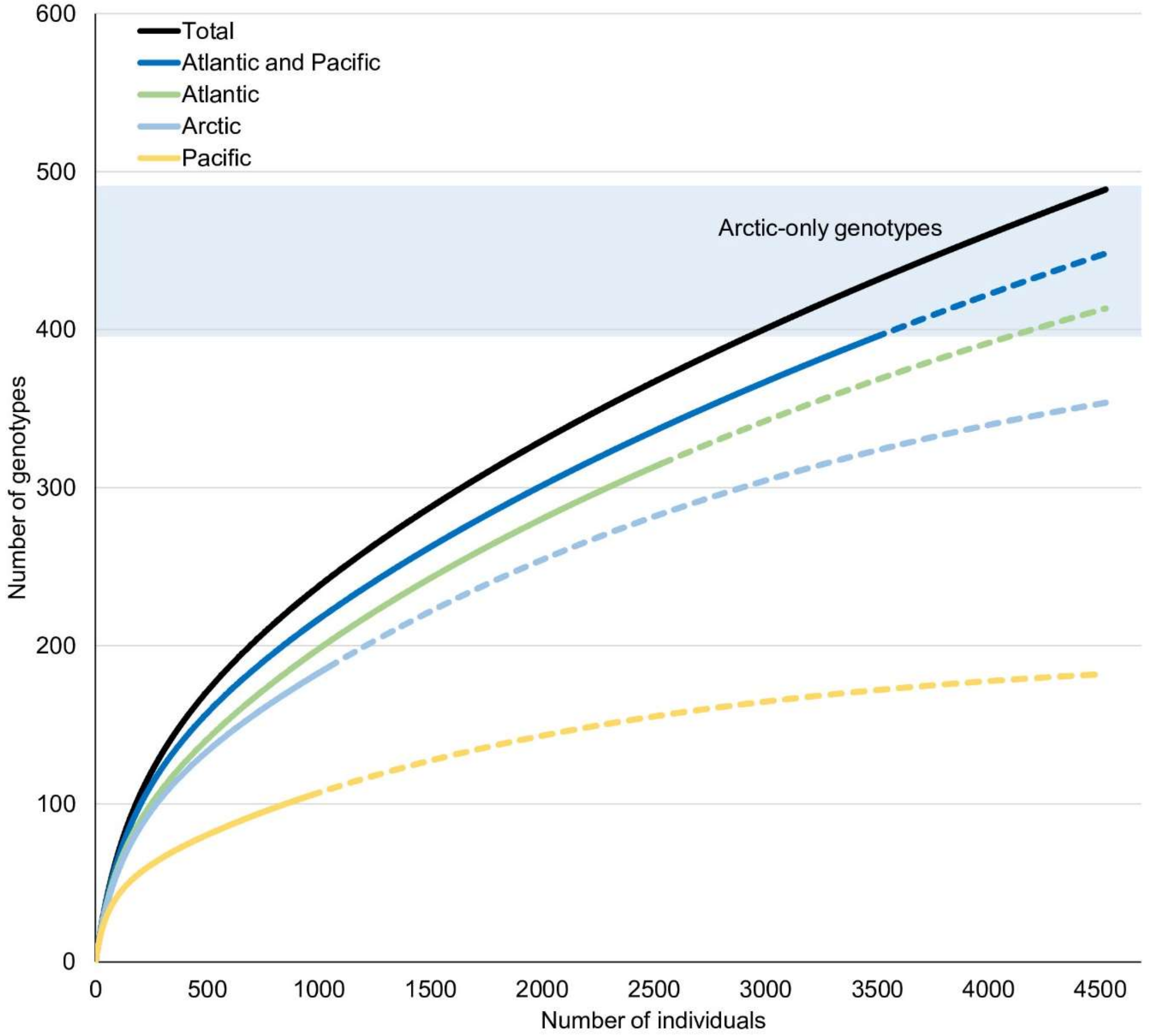

**Figure S2.** Accumulation curves for genotypes (haplotypes) in all specimens sampled. The Pacific curve includes all sequences from the Northwest, the Bering Sea (Nome, Alaska), and Northeast Pacific; the Arctic curve includes all sequences from the Beaufort (Alaska) and East Canadian Arctic; the Atlantic curve includes all sequences from the Northwest and Northeast Atlantic.

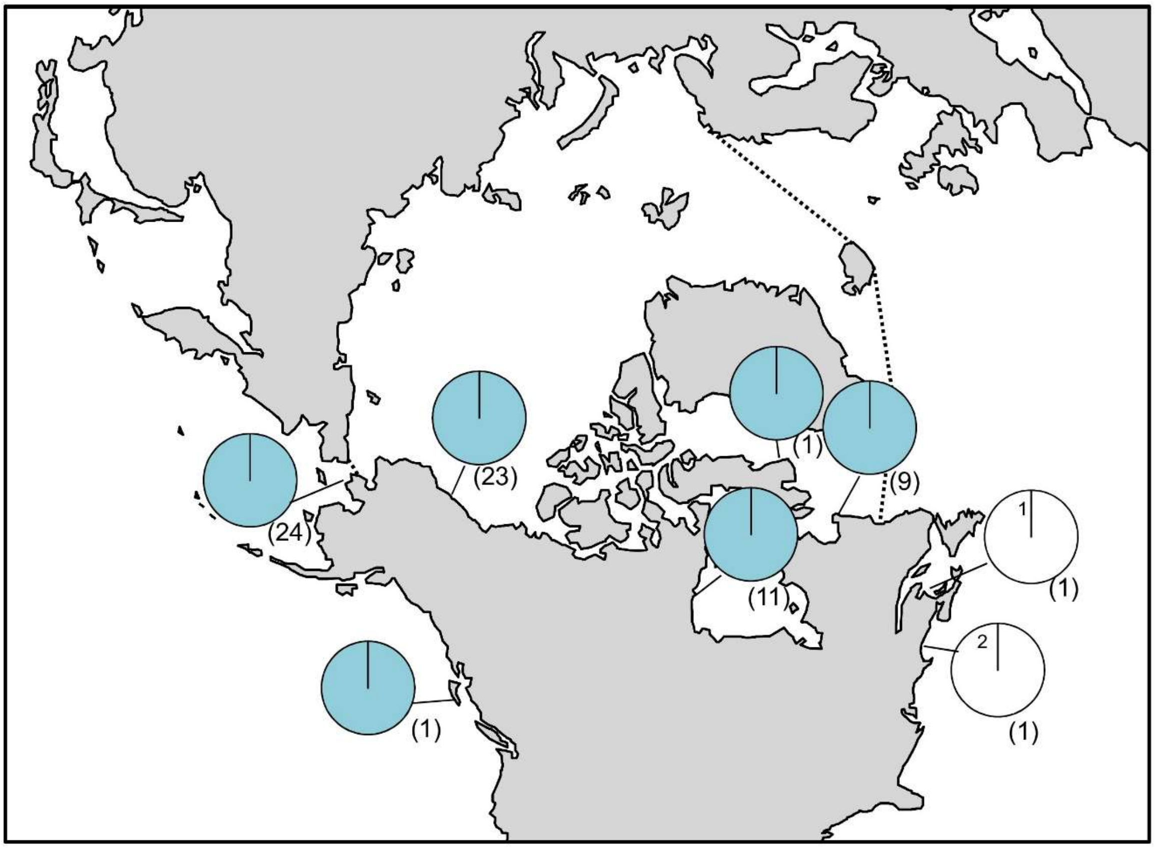

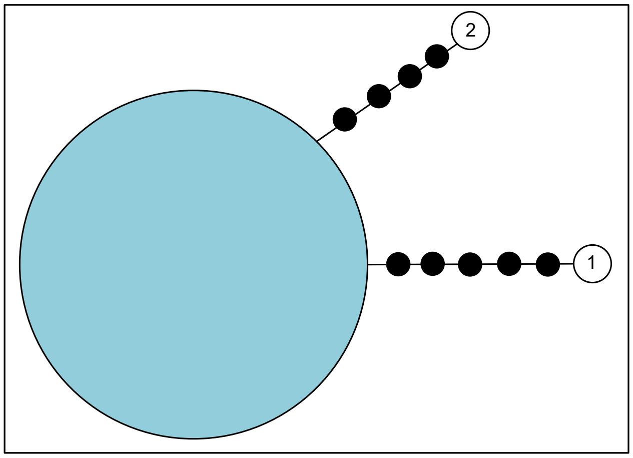

**Figure S3.** Haplotype map and network based on COI-5P data. In the map, numbers in parentheses refer to sample sizes from given locales, whereas numbers adjacent to white portions of pie charts refer to a haplotype sampled only once in the accompanying network. The dashed line indicates delineation of the Arctic Ocean. In the haplotype network, numbered haplotypes in white reference back to the map. Black circles indicate hypothesized (e.g. unsampled) haplotypes between clades. Circle size is proportional to the sampling frequency of a given haplotype.

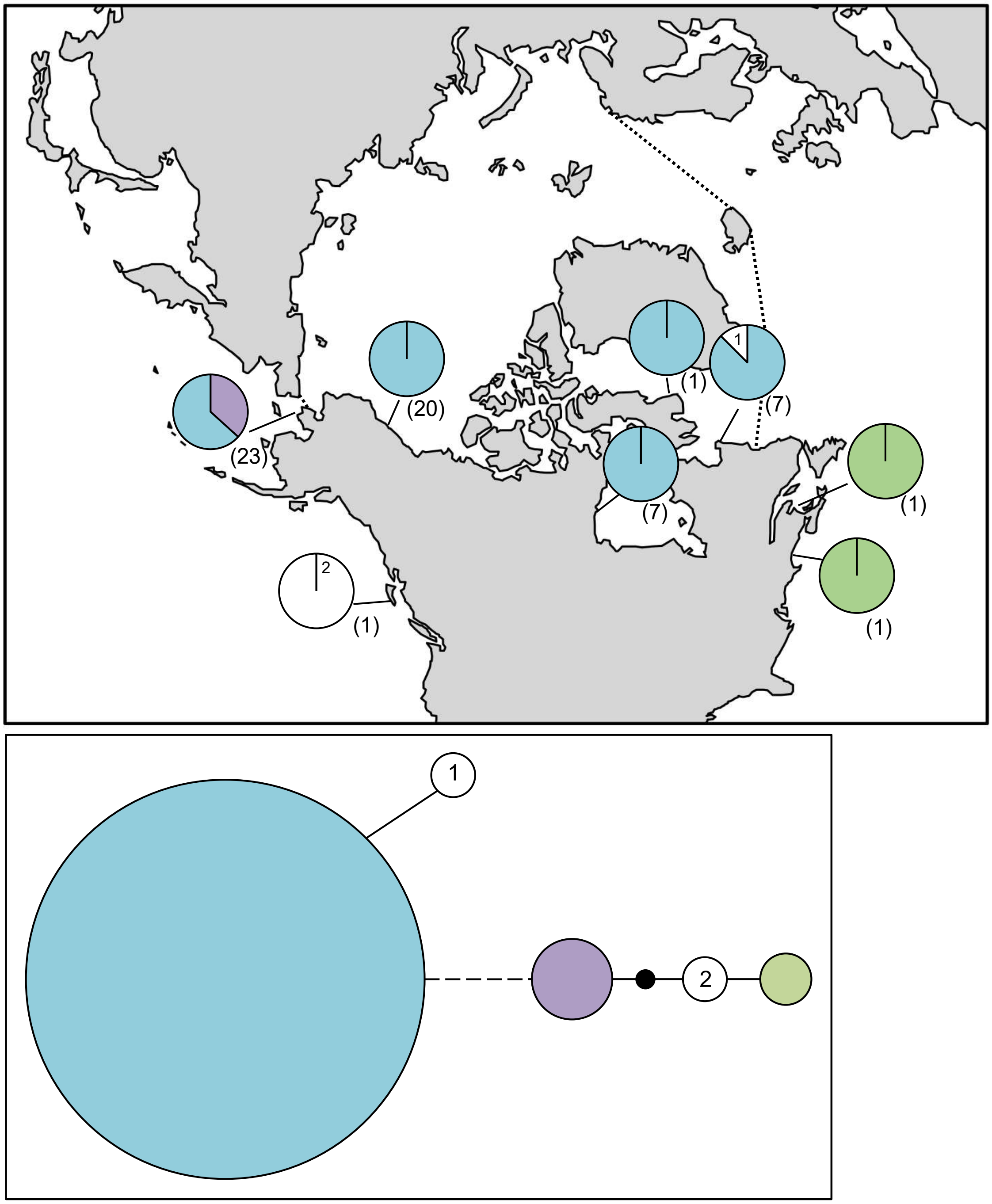

**Figure S4.** *Ahnfeltia borealis* haplotype map and network based on ycf35 data. In the map, numbers in parentheses refer to sample sizes from given locales, whereas numbers adjacent to white portions of pie charts refer to a haplotype sampled only once in the accompanying network. In the map, the dashed line indicates delineation of the Arctic Ocean. In the haplotype network, numbered haplotypes in white reference back to the map, the black circle represents a hypothesized (e.g. unsampled) haplotype between clades, and the dashed line refers to a two base pair insertion. Circle size is proportional to the sampling frequency of a given haplotype.

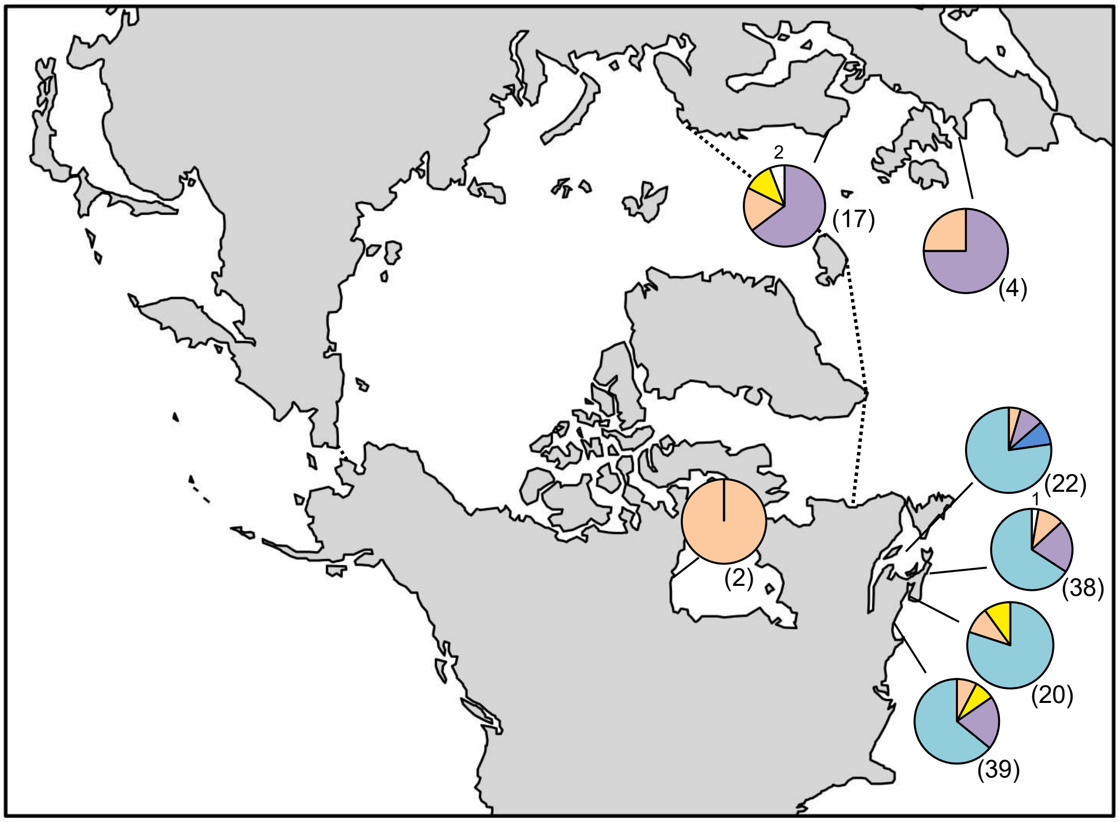

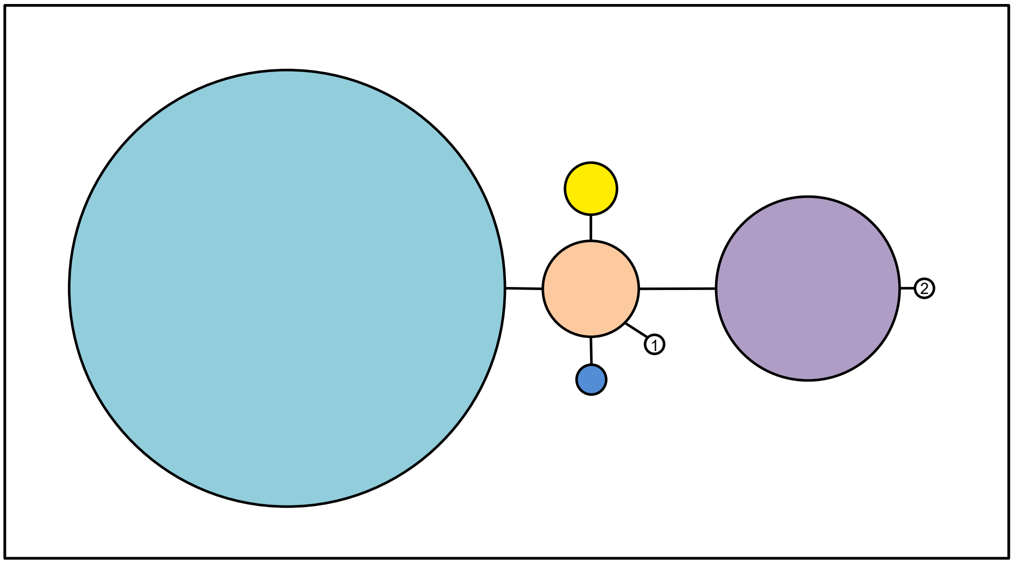

**Figure S5.** *Ahnfeltia plicata* haplotype map and network based on COI-5P data. In the map, numbers in parentheses refer to sample sizes from given locales, whereas numbers adjacent to white portions of pie charts refer to a haplotype sampled only once in the accompanying network. The dashed line indicates delineation of the Arctic Ocean. In the haplotype network, numbered haplotypes in white reference back to the map. Circle size is proportional to the sampling frequency of a given haplotype.

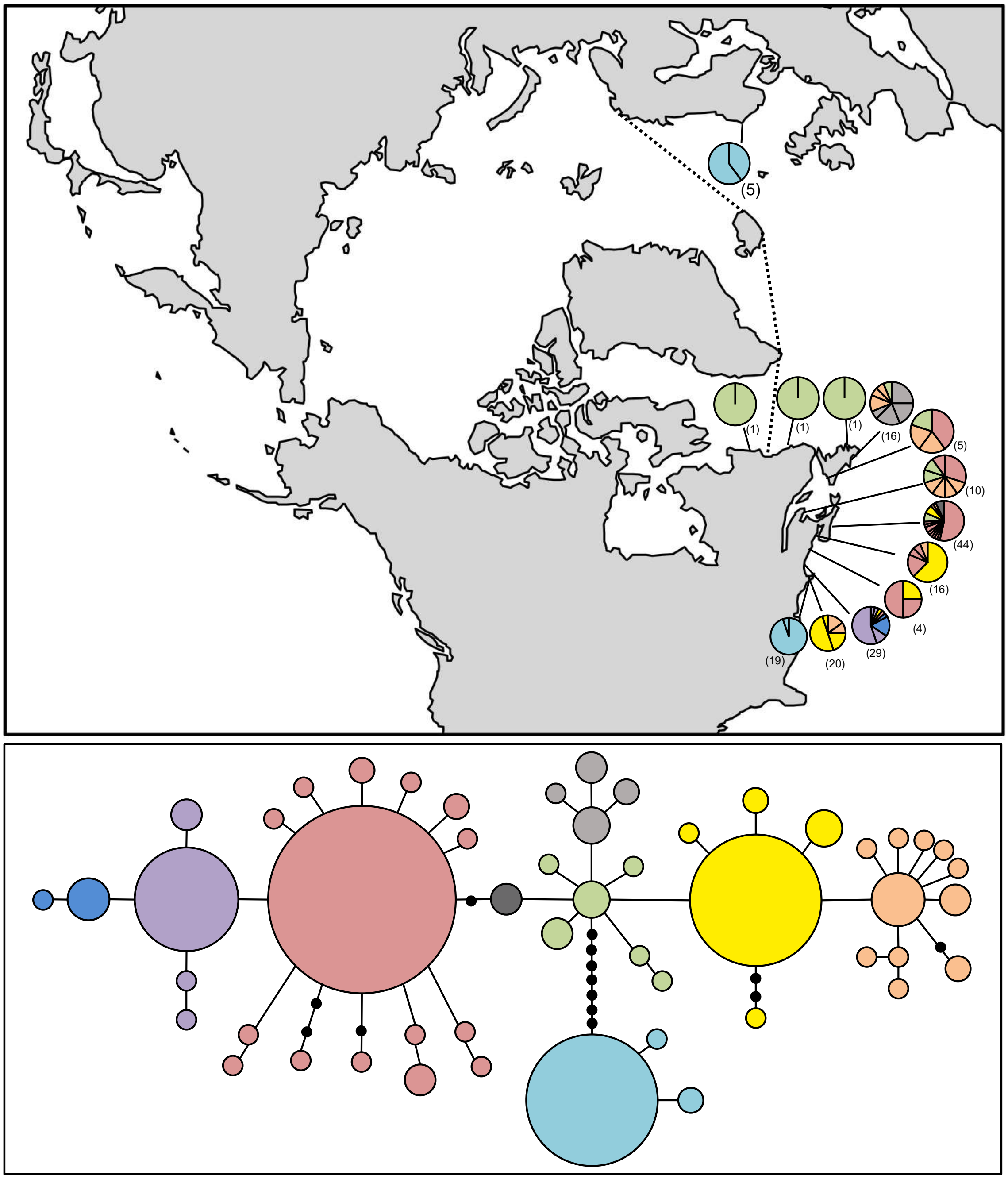

**Figure S6.** *Ceramium virgatum* haplotype map and network based on COI-5P data. In the map, numbers in parentheses refer to sample sizes from given locales. The dashed line indicates delineation of the Arctic Ocean. Circle size is proportional to the sampling frequency of a given haplotype. Black circles indicate hypothesized (e.g. unsampled) haplotypes between clades.

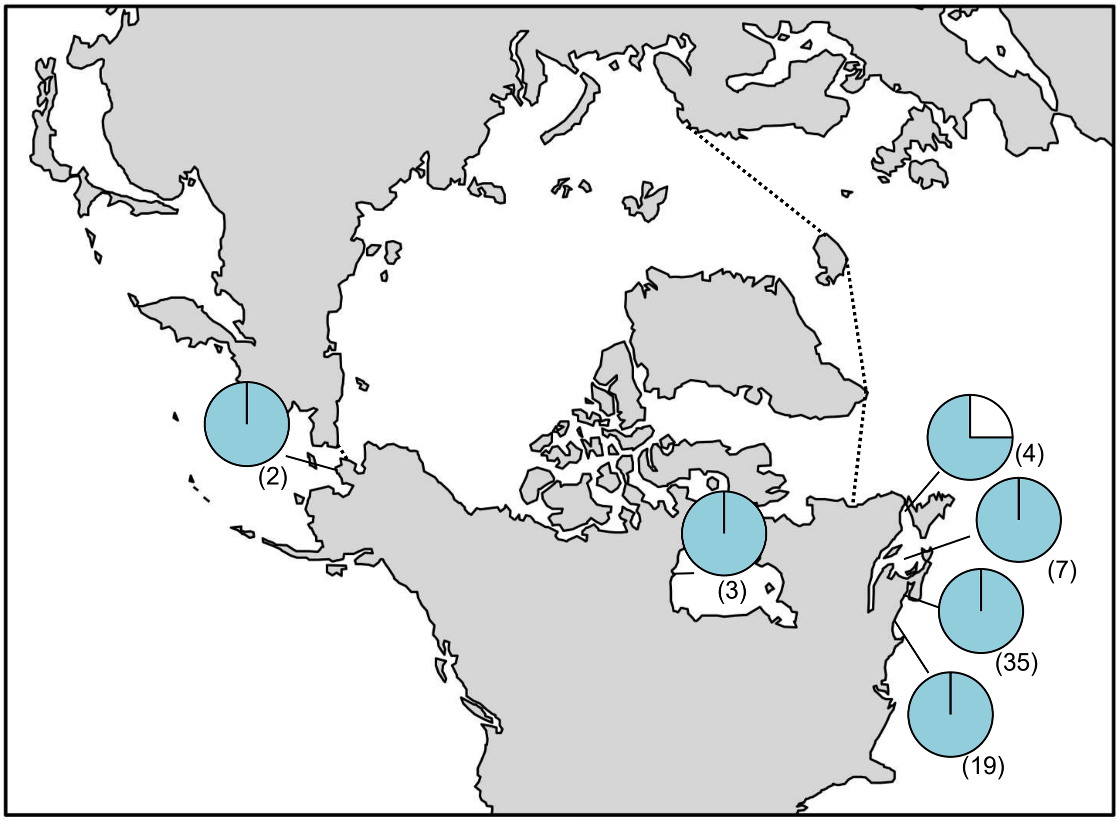

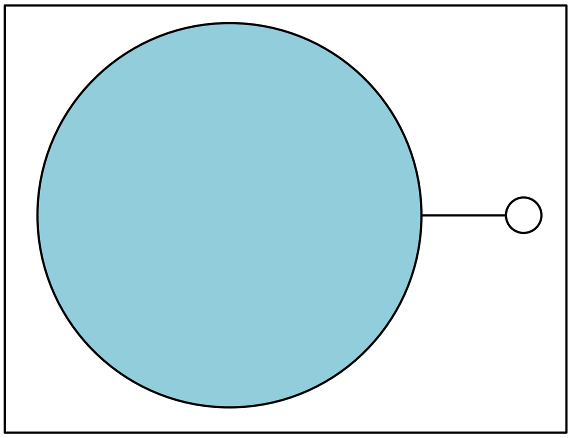

**Figure S7.** *Clathromorphum* sp. 9GWS haplotype map and network based on COI-5P data. In the map, numbers in parentheses refer to sample sizes from given locales. The dashed line indicates delineation of the Arctic Ocean. In the haplotype network, circle size is proportional to the sampling frequency of a given haplotype.

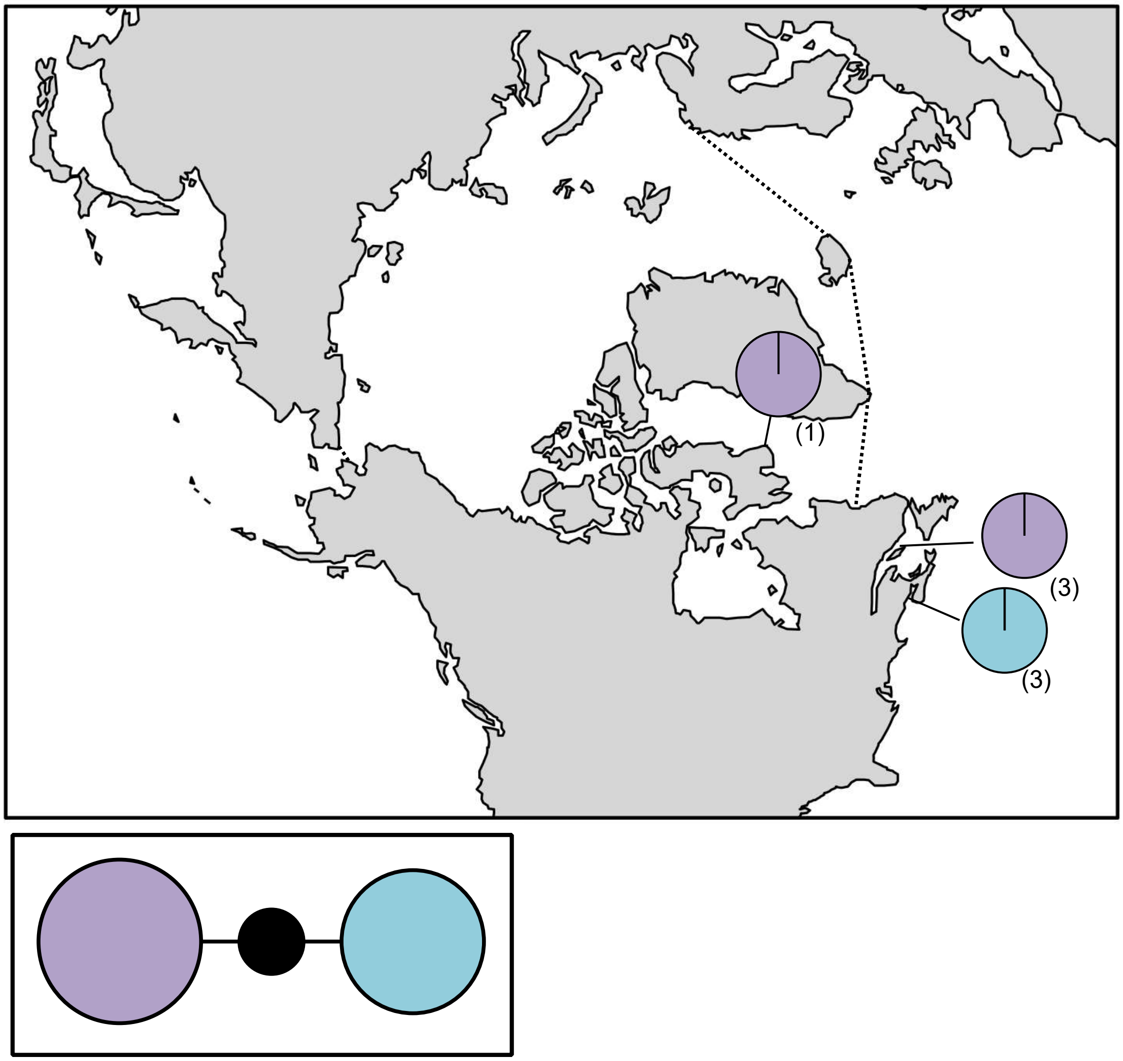

**Figure S8.** *Clathromorphum circumscriptum* haplotype map and network based on COI-5P data. In the map, numbers in parentheses refer to sample sizes from given locales. The dashed line indicates delineation of the Arctic Ocean. In the haplotype network, the black circle indicates a hypothesized (e.g. unsampled) haplotype between clades. Circle size is proportional to the sampling frequency of a given haplotype.

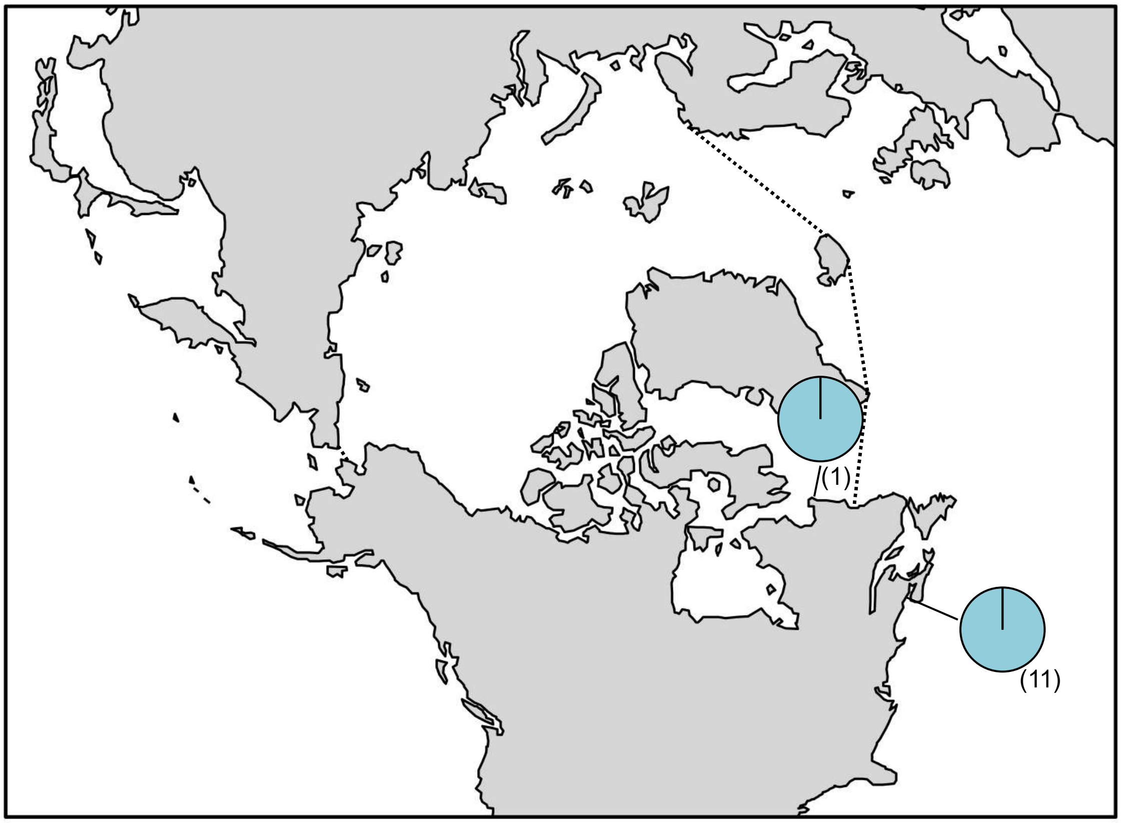

**Figure S9.** *Clathromorphum compactum* haplotype map and network based on COI-5P data. In the map, numbers in parentheses refer to sample sizes from given locales. The dashed line indicates delineation of the Arctic Ocean.

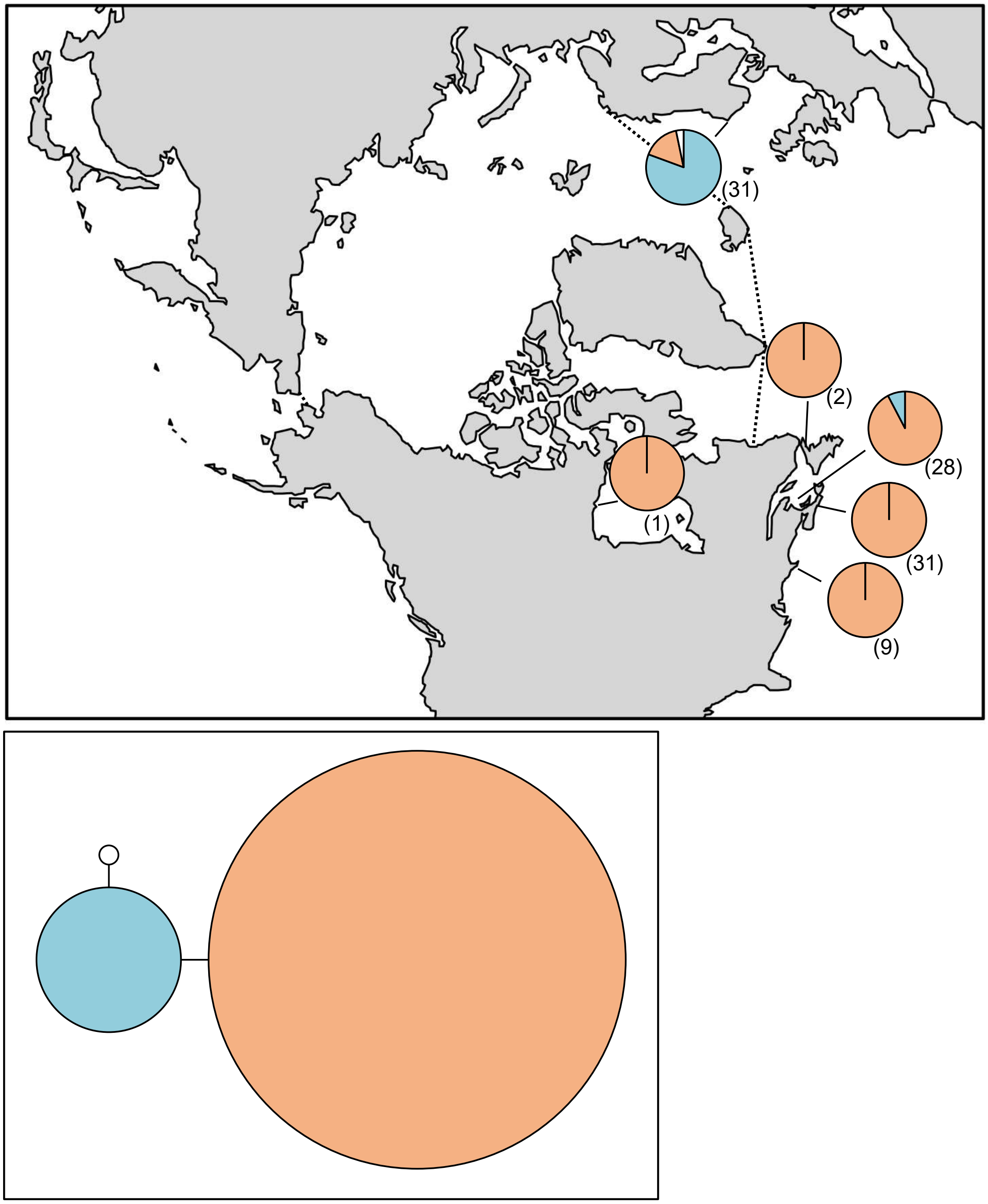

**Figure S10.** *Coccotylus brodiei* haplotype map and network based on COI-5P data. In the map, numbers in parentheses refer to sample sizes from given locales. The dashed line indicates delineation of the Arctic Ocean. In the haplotype network, circle size is proportional to the sampling frequency of a given haplotype.

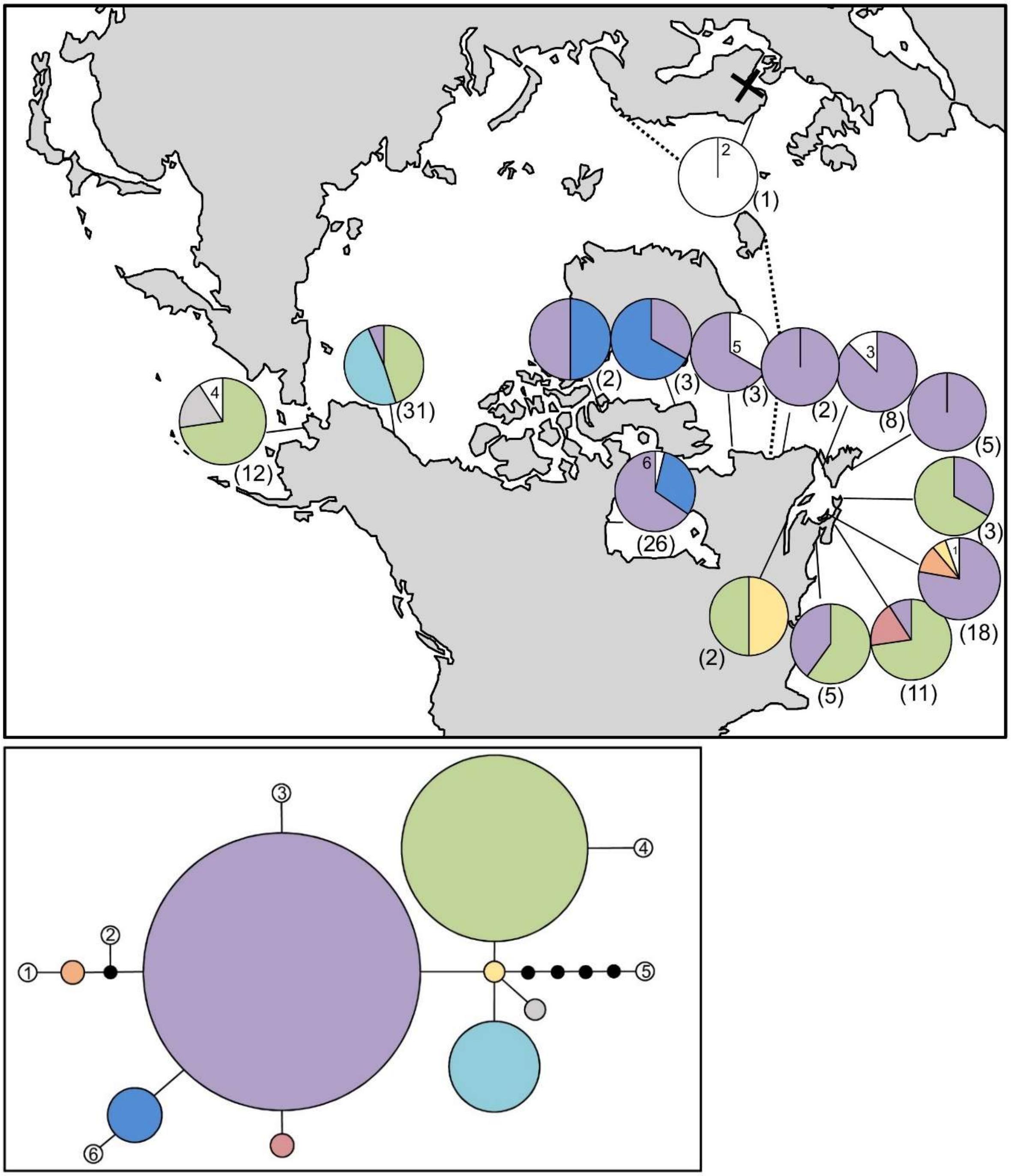

**Figure S11.** *Coccotylus truncatus* haplotype map and network based on COI-5P data. In the map, numbers in parentheses refer to sample sizes from given locales, whereas numbers adjacent to white portions of pie charts refer to a haplotype sampled only once in the accompanying network. The dashed line indicates delineation of the Arctic Ocean, while the black X indicates the location of genetically verified *Coccotylus truncatus* based on *rbc*L. In the haplotype network, numbered haplotypes in white reference back to the map. Black circles indicate hypothesized (e.g. unsampled) haplotypes between clades. Circle size is proportional to the sampling frequency of a given haplotype.

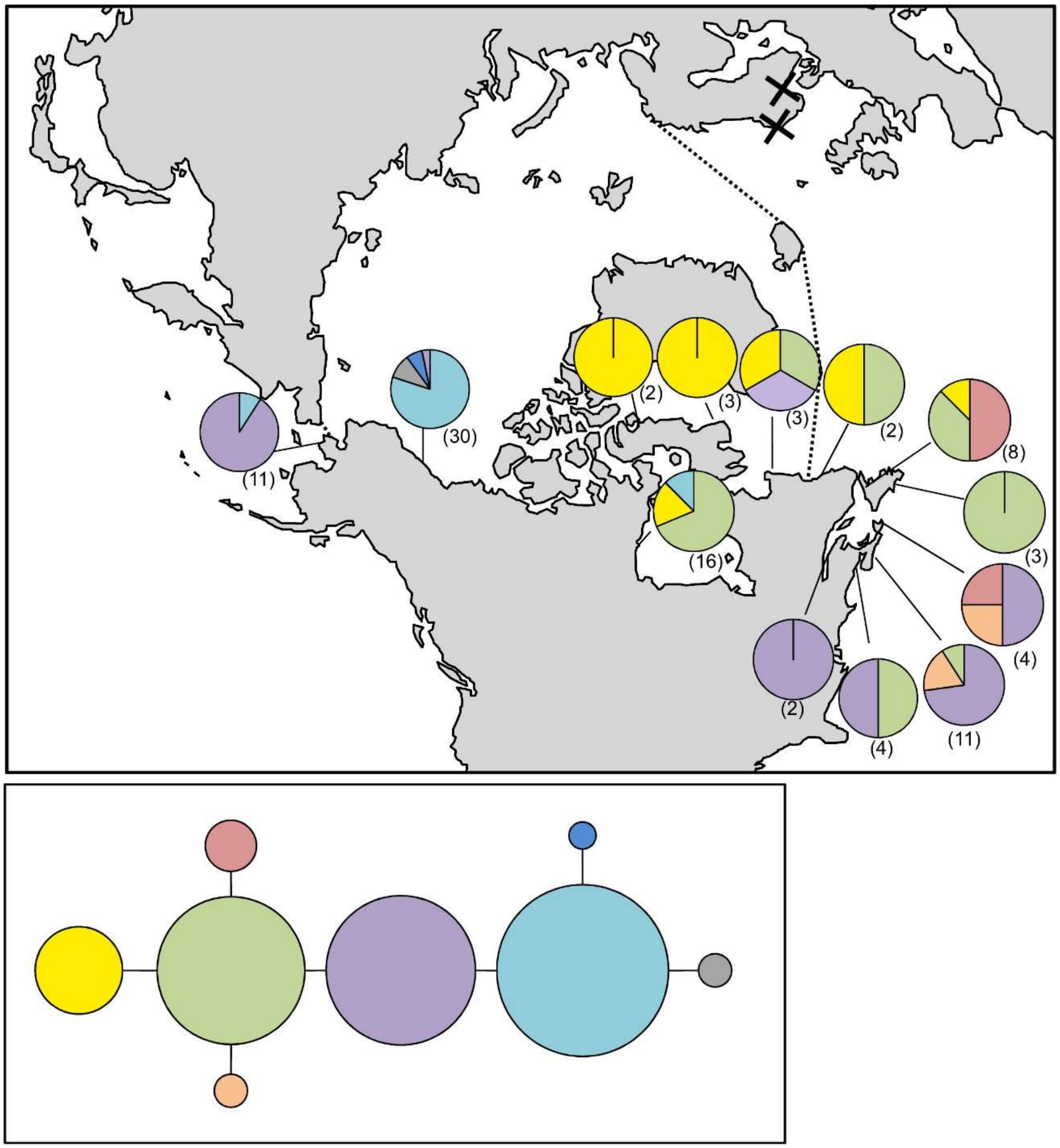

**Figure S12.** *Coccotylus truncatus* haplotype map and network based on ITS data. In the map, numbers in parentheses refer to sample sizes from given locales. The dashed line indicates delineation of the Arctic Ocean, while the black X indicates the location of genetically verified *Coccotylus truncatus* based on COI-5P and *rbc*L. In the haplotype network, circle size is proportional to the sampling frequency of a given haplotype.

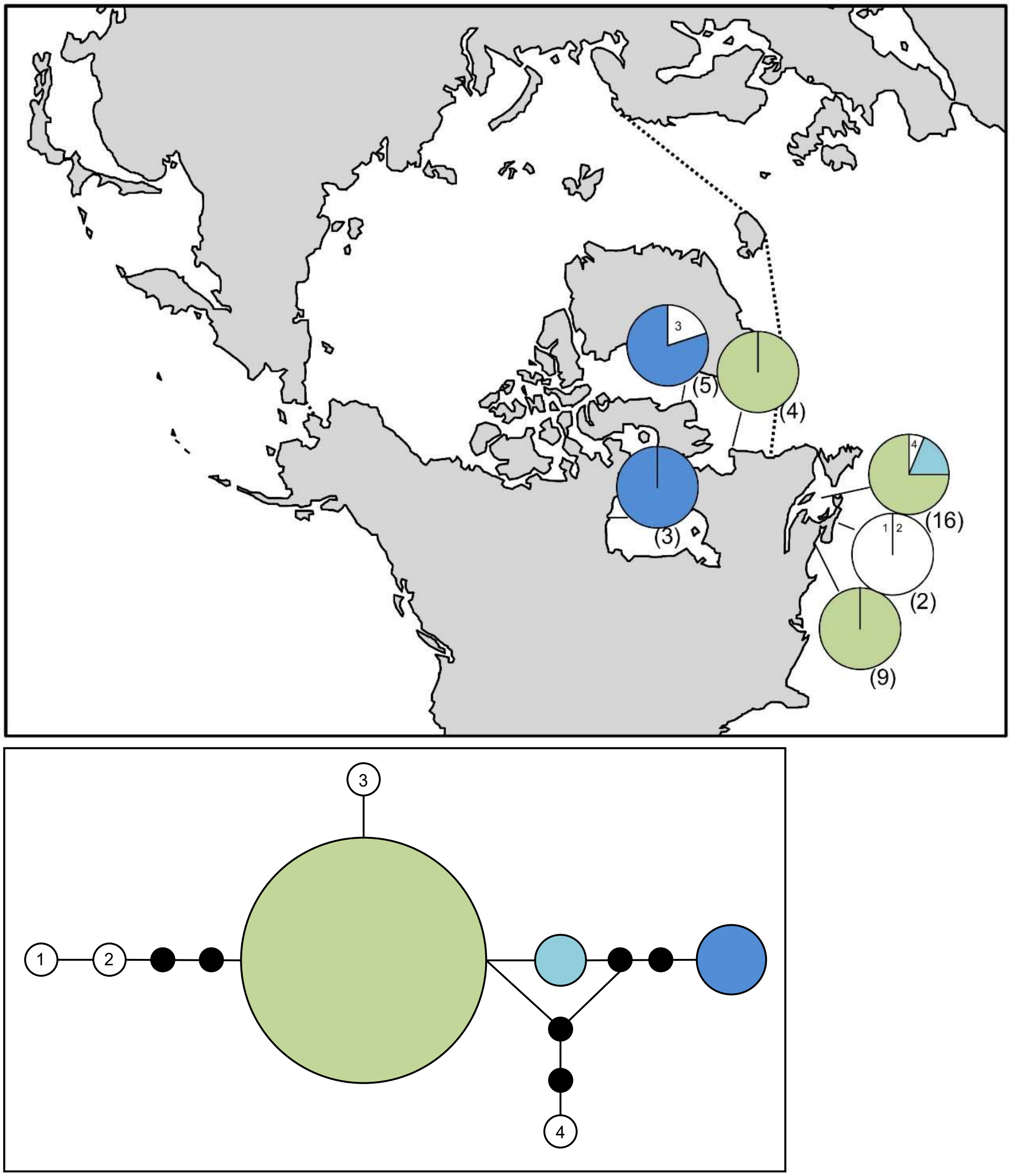

**Figure S13.** *Devaleraea ramentacea* haplotype map and network based on COI-5P data. In the map, numbers in parentheses refer to sample sizes from given locales, whereas numbers adjacent to white portions of pie charts refer to a haplotype sampled only once in the accompanying network. The dashed line indicates delineation of the Arctic Ocean. In the haplotype network, numbered haplotypes in white reference back to the map. Black circles indicate hypothesized (e.g. unsampled) haplotypes between clades. Circle size is proportional to the sampling frequency of a given haplotype.

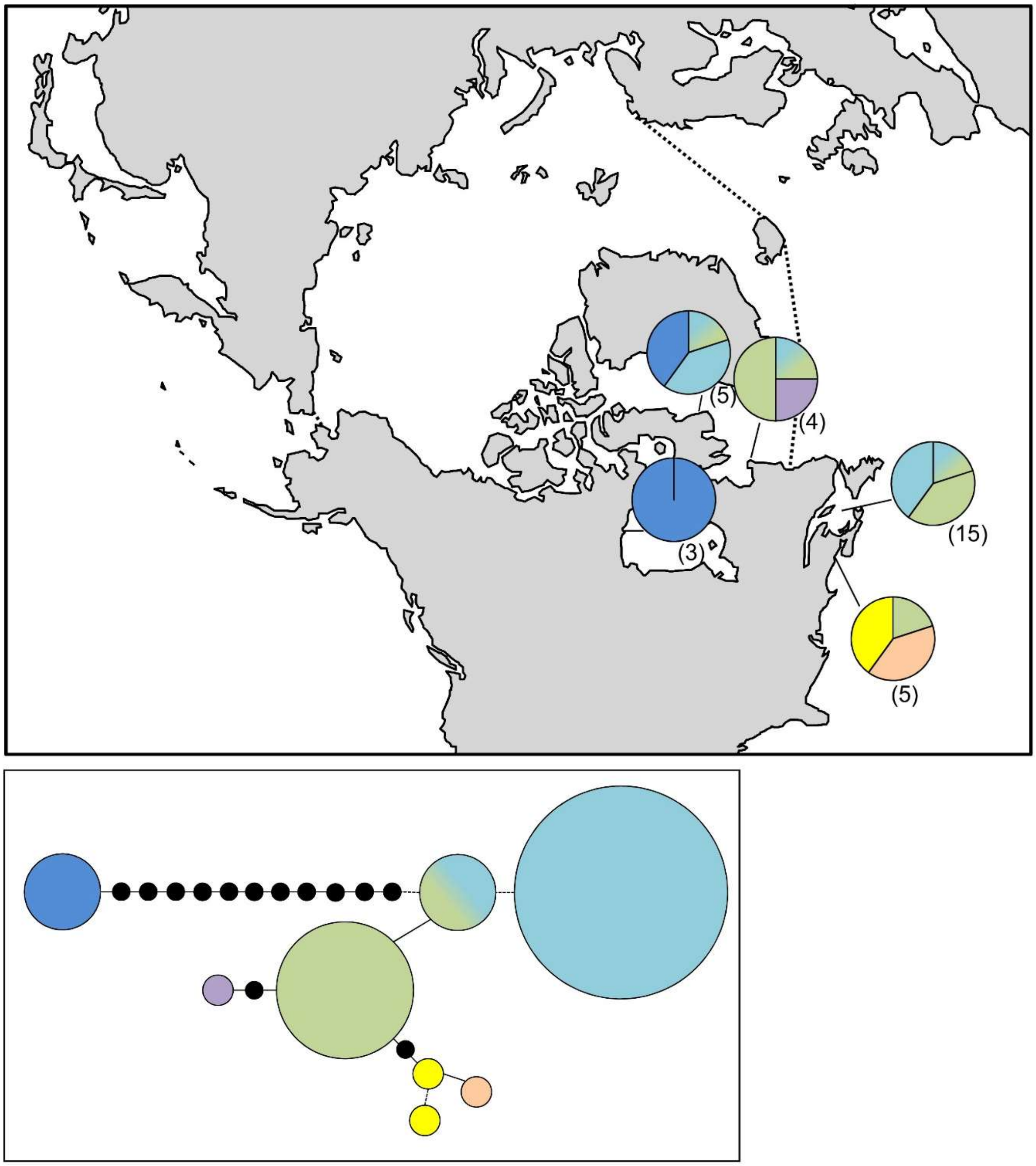

**Figure S14.** *Devaleraea ramentacea* haplotype map and network based on ITS data. In the map, numbers in parentheses refer to sample sizes from given locales. The dashed line indicates delineation of the Arctic Ocean. Black circles indicate hypothesized (e.g. unsampled) haplotypes and dashed lines represent insertions/deletions between clades. Circle size is proportional to the sampling frequency of a given haplotype.

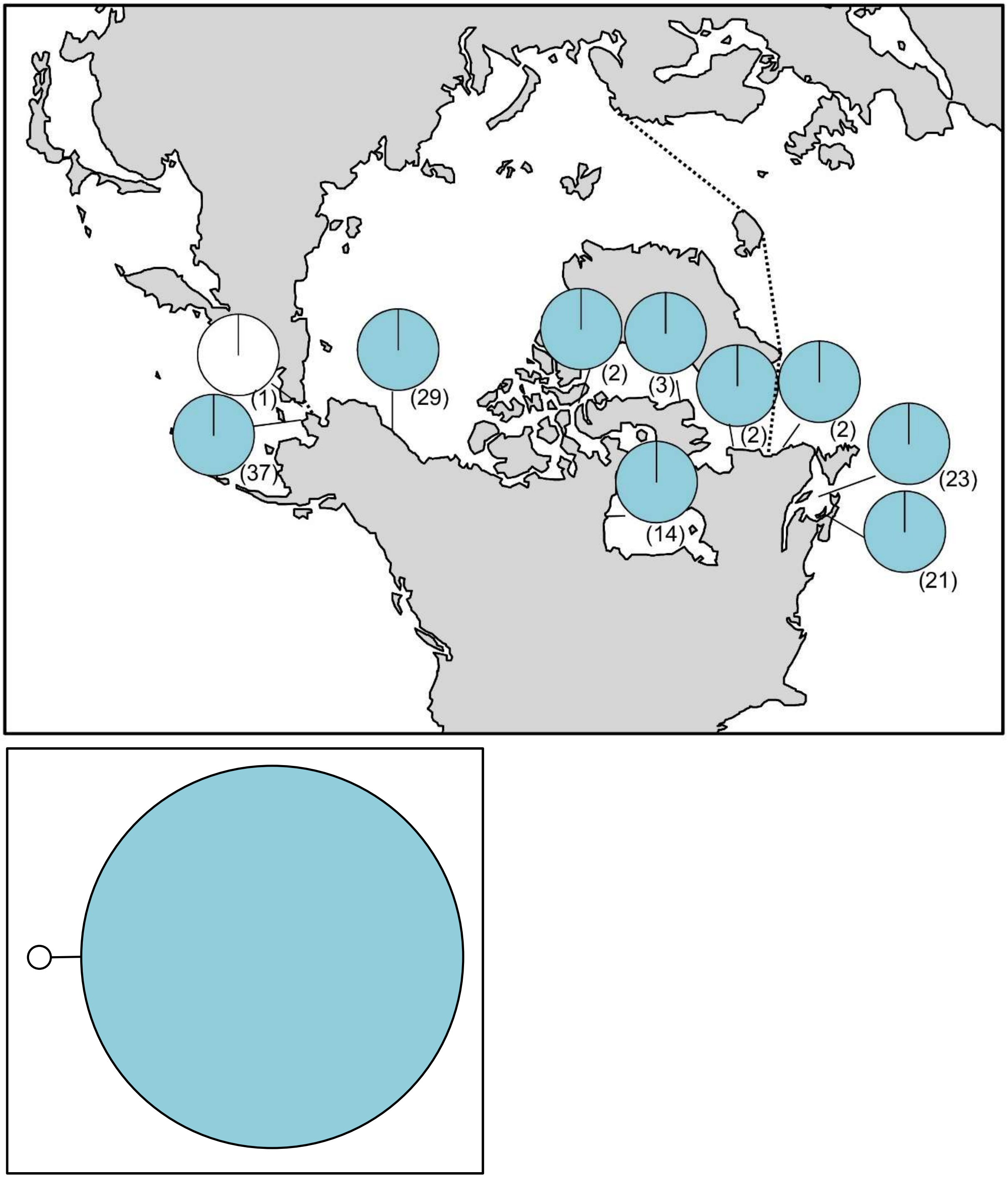

**Figure S15.** *Dilsea socialis* haplotype map and network based on COI-5P data. In the map, numbers in parentheses refer to sample sizes from given locales. The dashed line indicates delineation of the Arctic Ocean. In the haplotype network, circle size is proportional to the sampling frequency of a given haplotype.

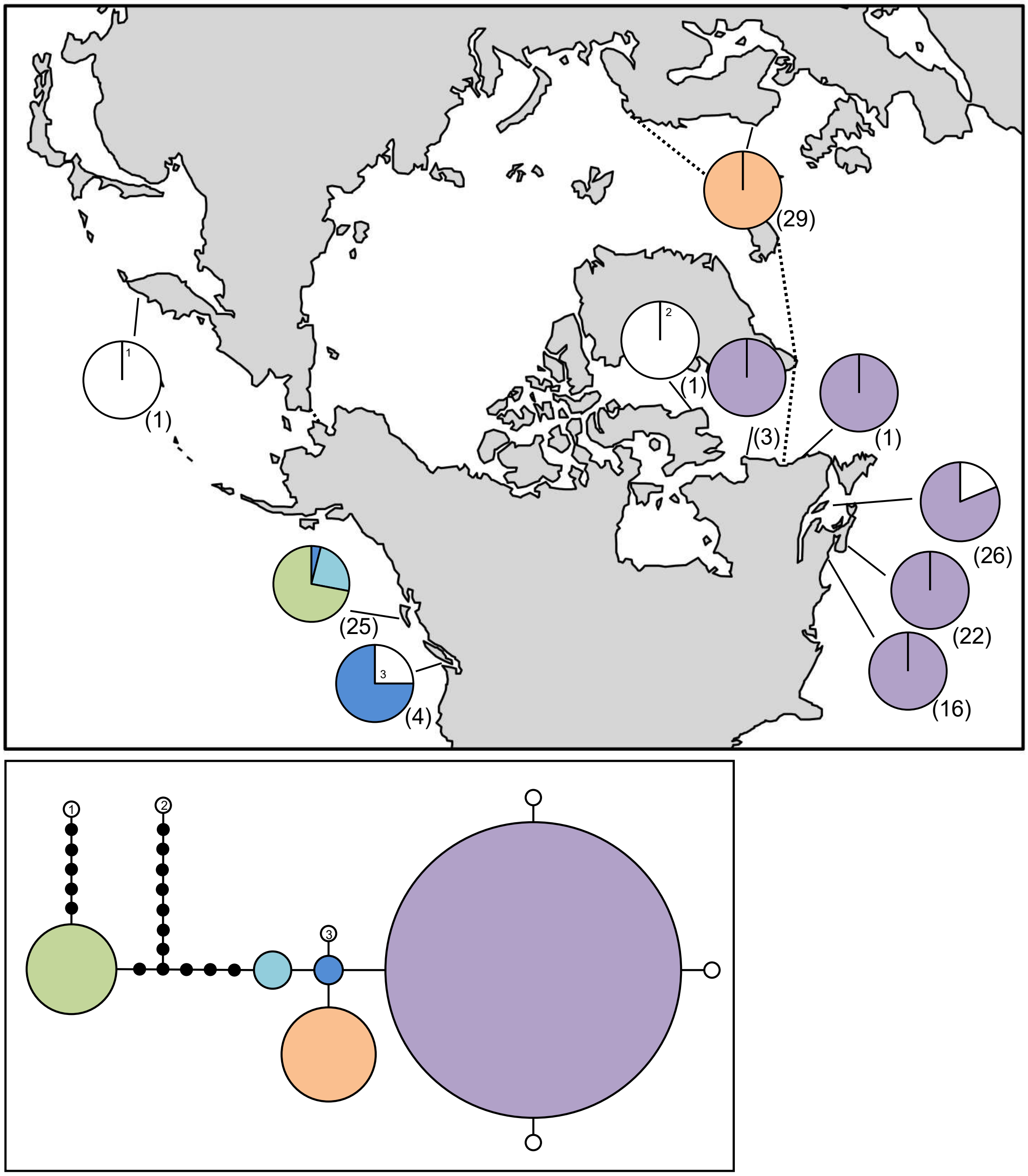

**Figure S16.** *Euthora cristata* haplotype map and network based on COI-5P data. In the map, numbers in parentheses refer to sample sizes from given locales. The dashed line indicates delineation of the Arctic Ocean. In the haplotype network, black circles indicate hypothesized (e.g. unsampled) haplotypes between clades. Circle size is proportional to the sampling frequency of a given haplotype. In the haplotype network, numbered haplotypes in white reference back to the map.

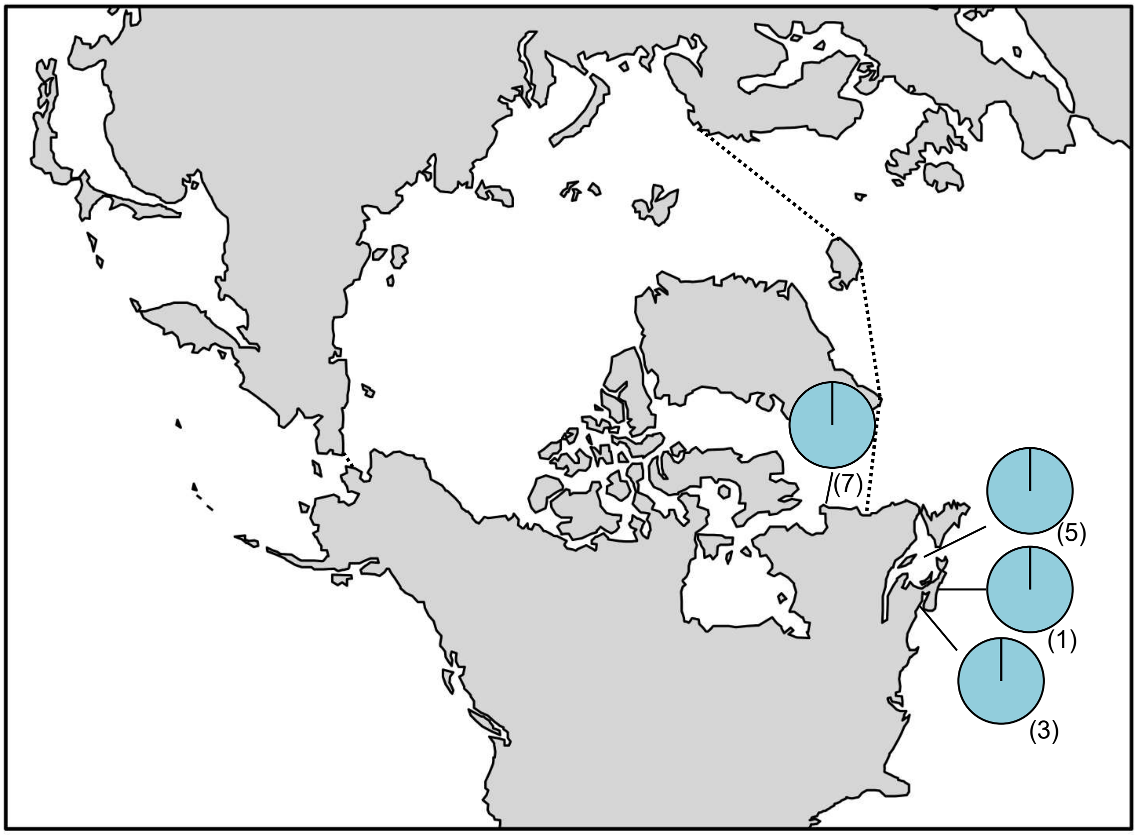

**Figure S17.** *Fimbrifolium dichotomum* haplotype map based on COI-5P data. In the map, numbers in parentheses refer to sample sizes from given locales. The dashed line indicates delineation of the Arctic Ocean.

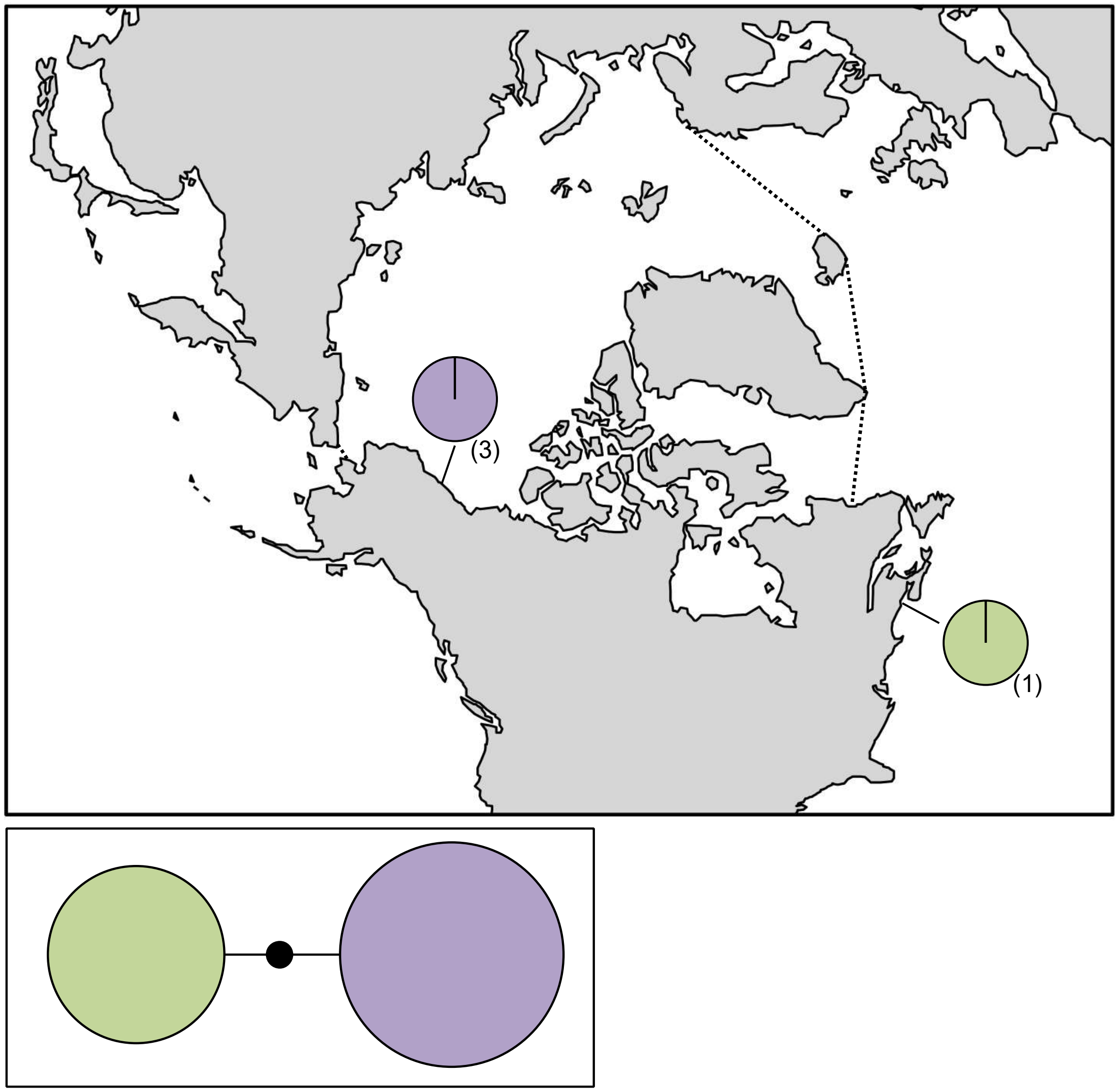

**Figure S18.** *Leptophytum foecundum* haplotype map and network based on COI-5P data. In the map, numbers in parentheses refer to sample sizes from given locales. The dashed line indicates delineation of the Arctic Ocean. In the haplotype network, the black circle represents a hypothesized (e.g. unsampled) haplotype between clades. Circle size is proportional to the sampling frequency of a given haplotype.

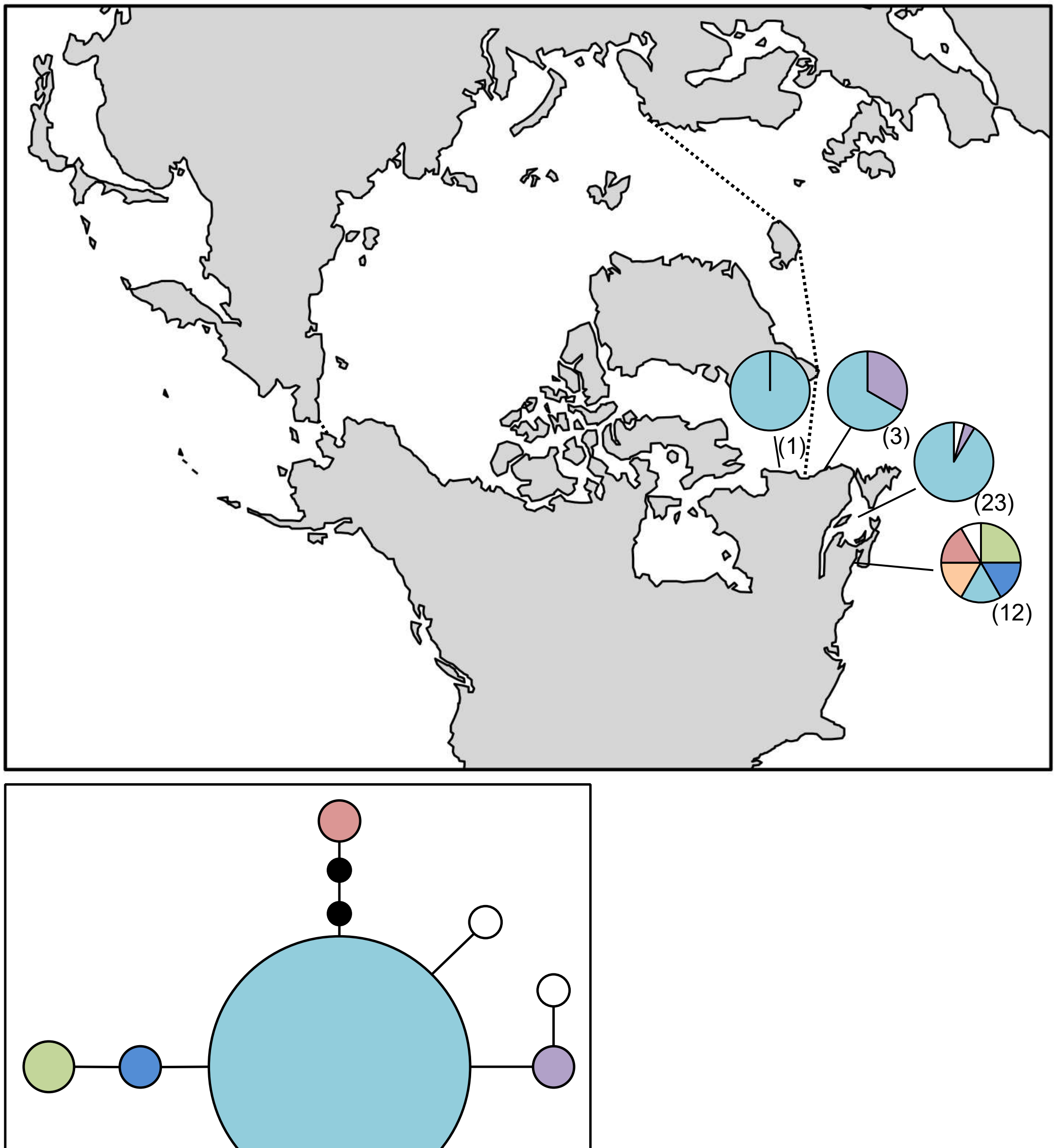

**Figure S19.** *Leptosiphonia flexicaulis* haplotype map based on COI-5P data. In the map, numbers in parentheses refer to sample sizes from given locales. The dashed line indicates delineation of the Arctic Ocean. In the haplotype network, black circles indicate hypothesized (e.g. unsampled) haplotypes between clades. Circle size is proportional to the sampling frequency of a given haplotype.

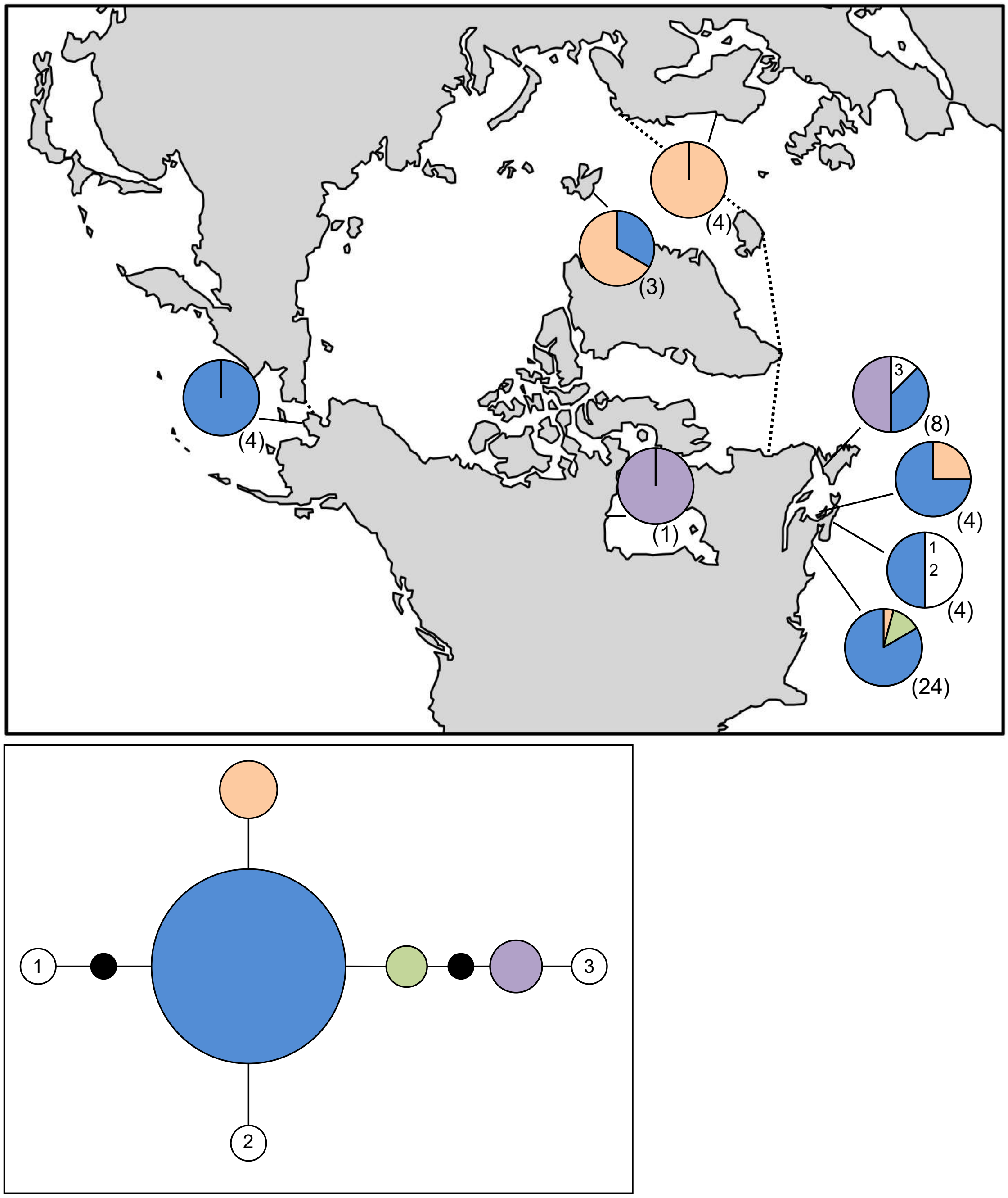

**Figure S20.** *Lithothamnion glaciale* haplotype map and network based on COI-5P data. In the map, numbers in parentheses refer to sample sizes from given locales, whereas numbers adjacent to white portions of pie charts refer to a haplotype sampled only once in the accompanying network. The dashed line indicates delineation of the Arctic Ocean. In the haplotype network, numbered haplotypes in white reference back to the map. Black circles indicate hypothesized (e.g. unsampled) haplotypes between clades. Circle size is proportional to the sampling frequency of a given haplotype.

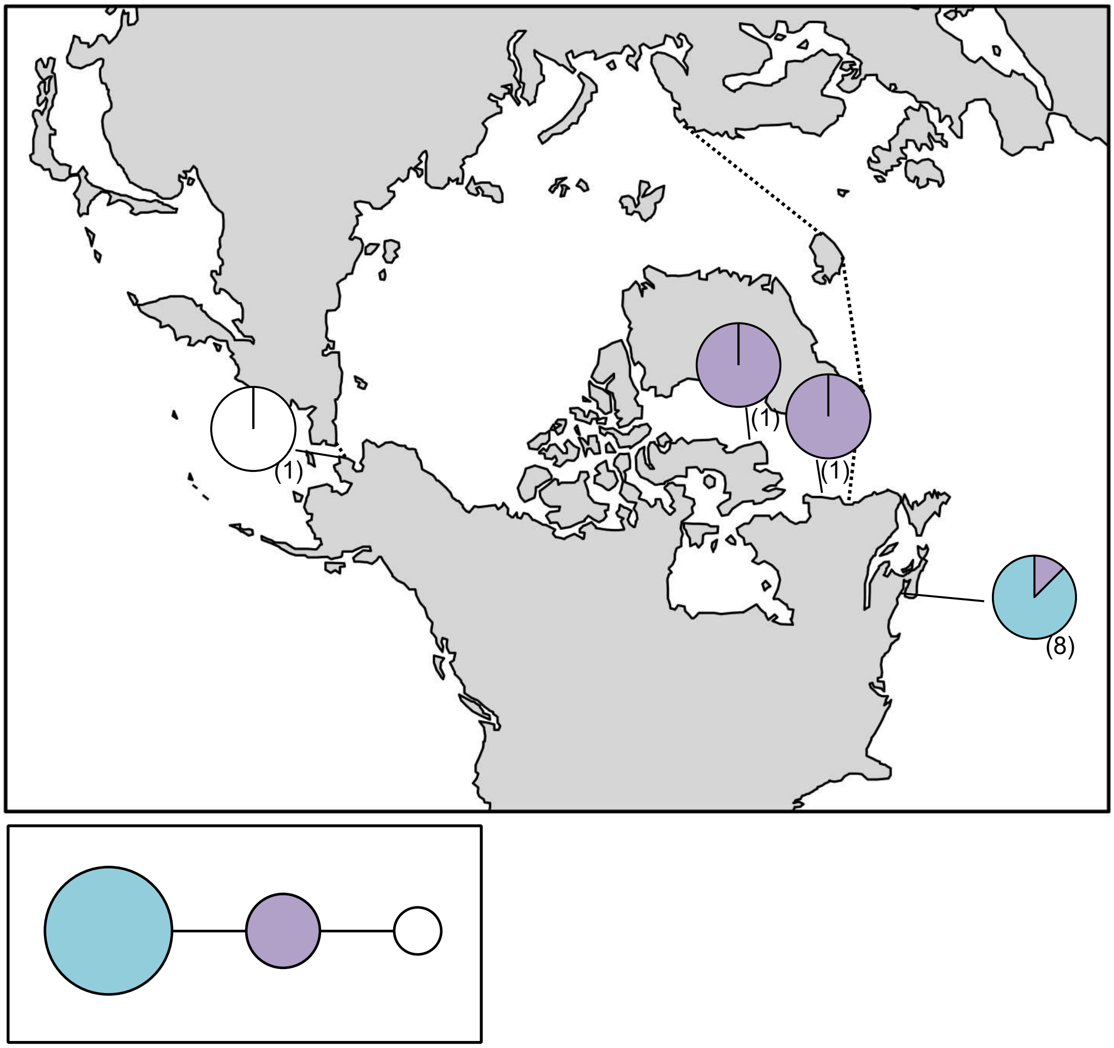

**Figure S21.** *Lithothamnion lemoineae* haplotype map based on COI-5P data. In the map, numbers in parentheses refer to sample sizes from given locales. The dashed line indicates delineation of the Arctic Ocean. In the haplotype network, circle size is proportional to the sampling frequency of a given haplotype.

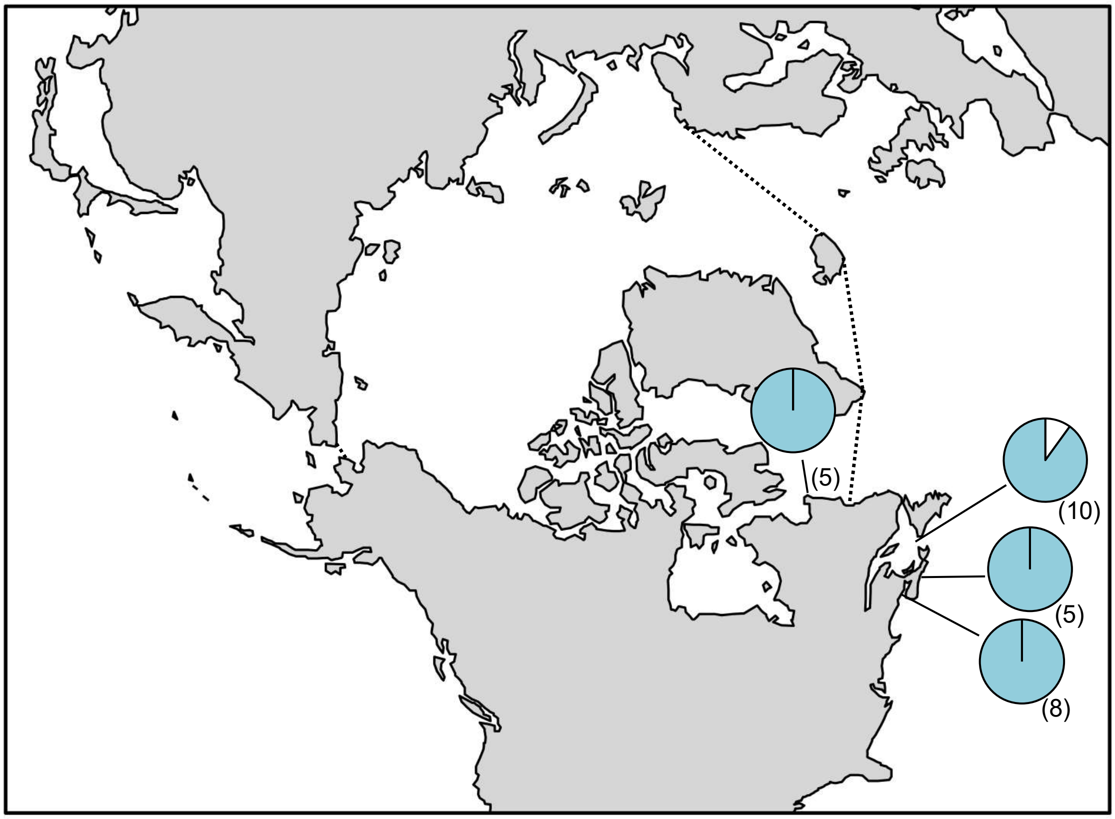

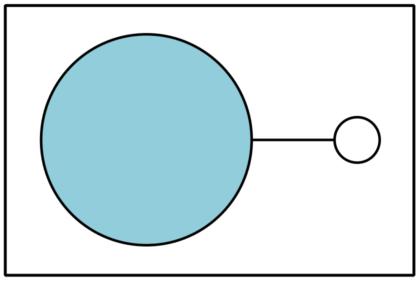

**Figure S22.** *Membranoptera fabriciana* haplotype map based on COI-5P data. In the map, numbers in parentheses refer to sample sizes from given locales. The dashed line indicates delineation of the Arctic Ocean. In the haplotype network, circle size is proportional to the sampling frequency of a given haplotype.

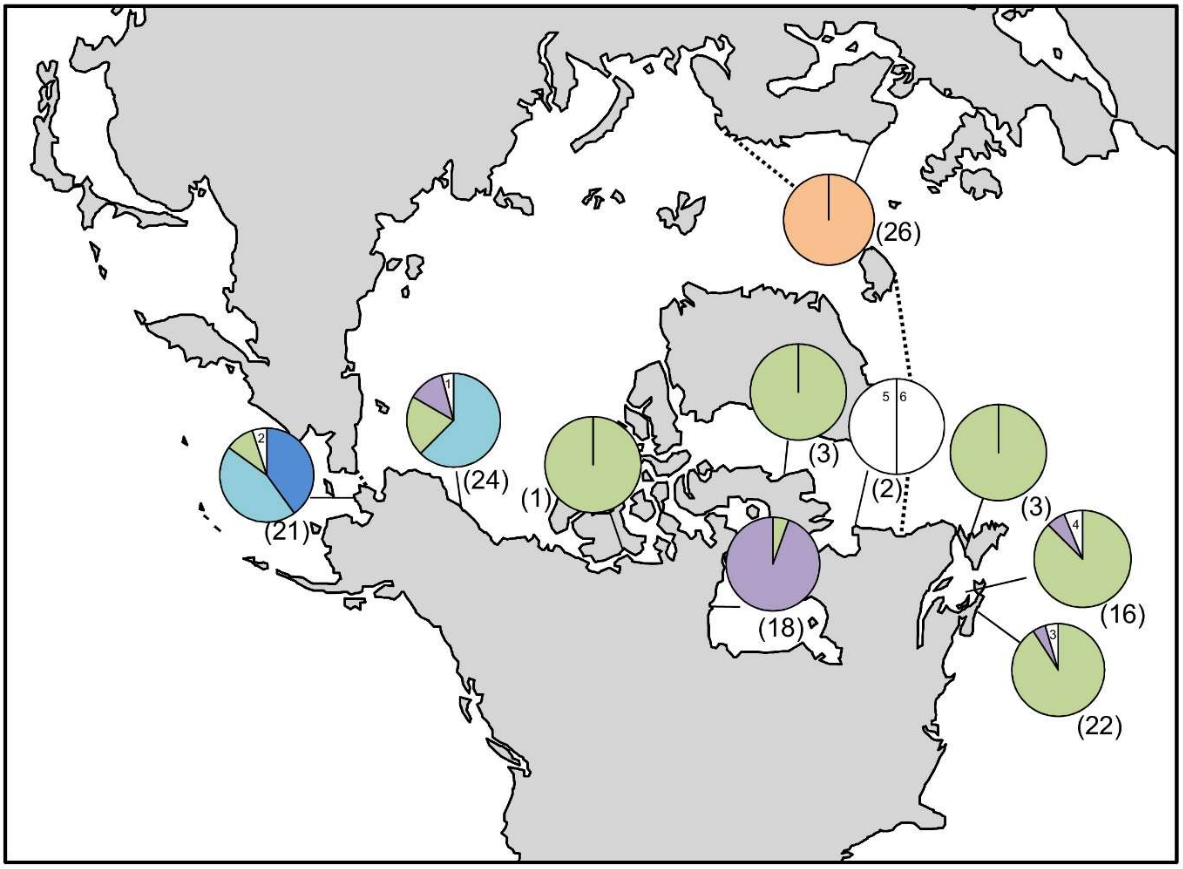

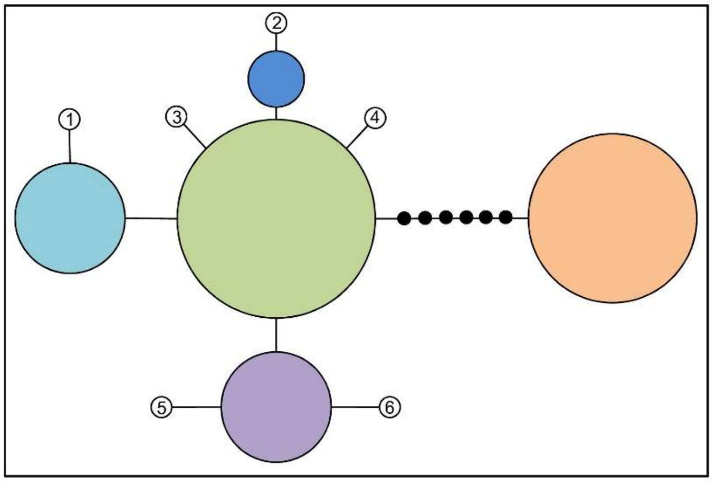

**Figure S23.** *Odonthalia dentata* haplotype map and network based on COI-5P data. In the map, numbers in parentheses refer to sample sizes from given locales, whereas numbers adjacent to white portions of pie charts refer to a haplotype sampled only once in the accompanying network. The dashed line indicates delineation of the Arctic Ocean. In the haplotype network, numbered haplotypes in white reference back to the map. Black circles indicate hypothesized (e.g. unsampled) haplotypes between clades. Circle size is proportional to the sampling frequency of a given haplotype.

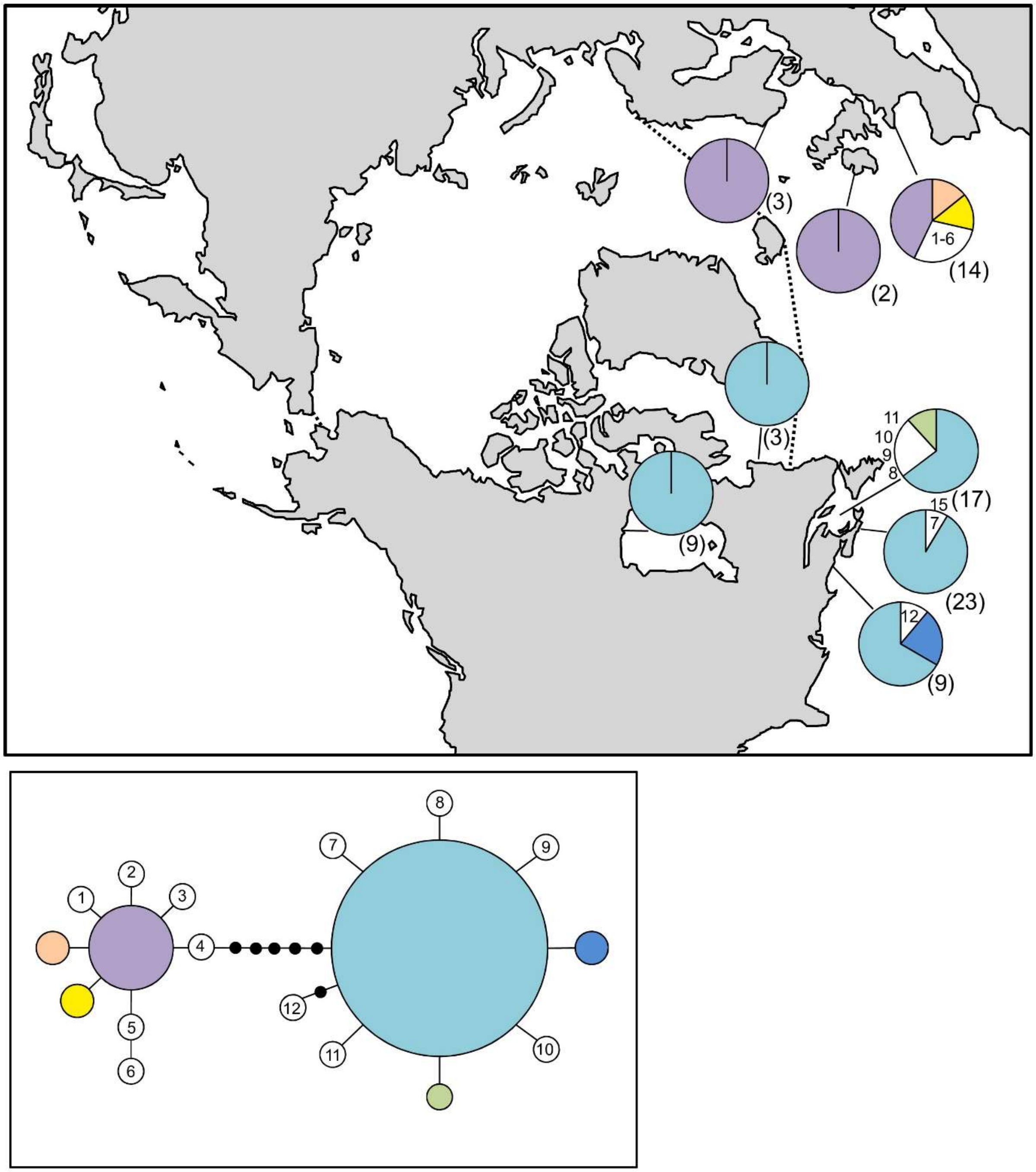

**Figure S24.** *Palmaria palmata* haplotype map and network based on COI-5P data. In the map, numbers in parentheses refer to sample sizes from given locales, whereas numbers adjacent to white portions of pie charts refer to a haplotype sampled only once in the accompanying network. The dashed line indicates delineation of the Arctic Ocean. In the haplotype network, numbered haplotypes in white reference back to the map. Black circles indicate hypothesized (e.g. unsampled) haplotypes between clades. Circle size is proportional to the sampling frequency of a given haplotype.

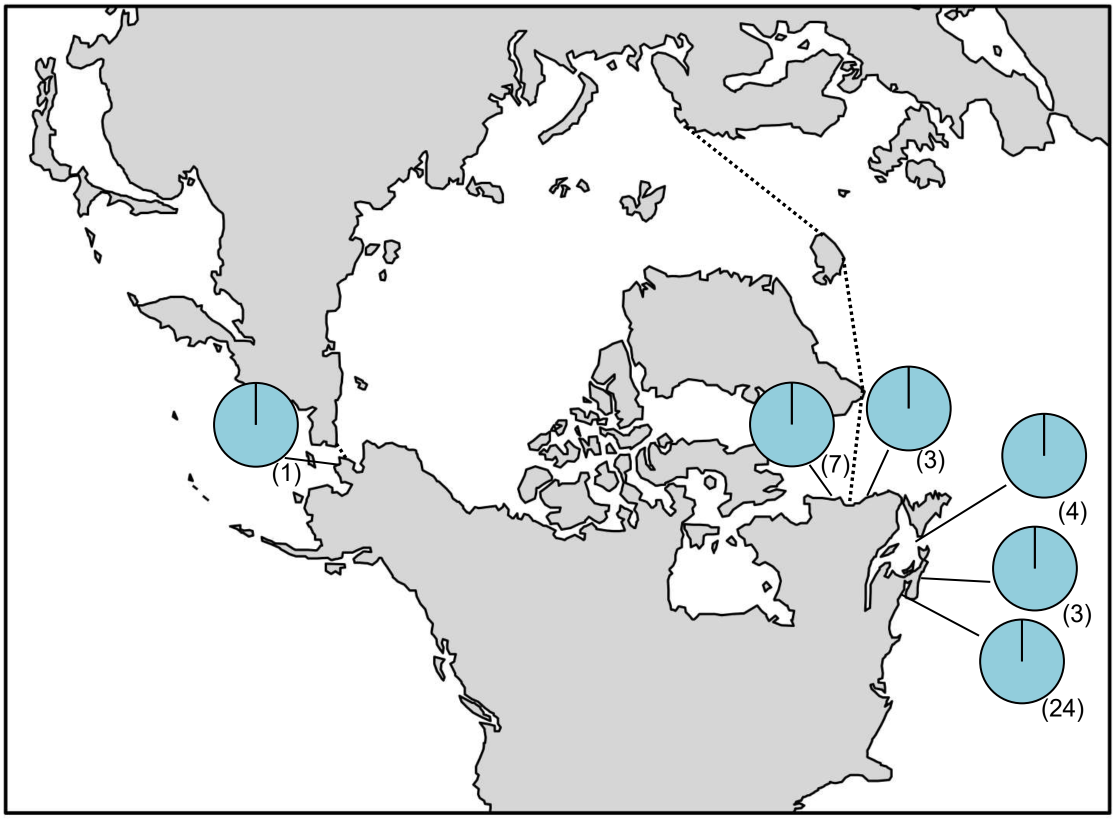

**Figure E25.** *Peyssonnelia rosenvingei* haplotype map based on COI-5P data. In the map, numbers in parentheses refer to sample sizes from given locales. The dashed line indicates delineation of the Arctic Ocean.

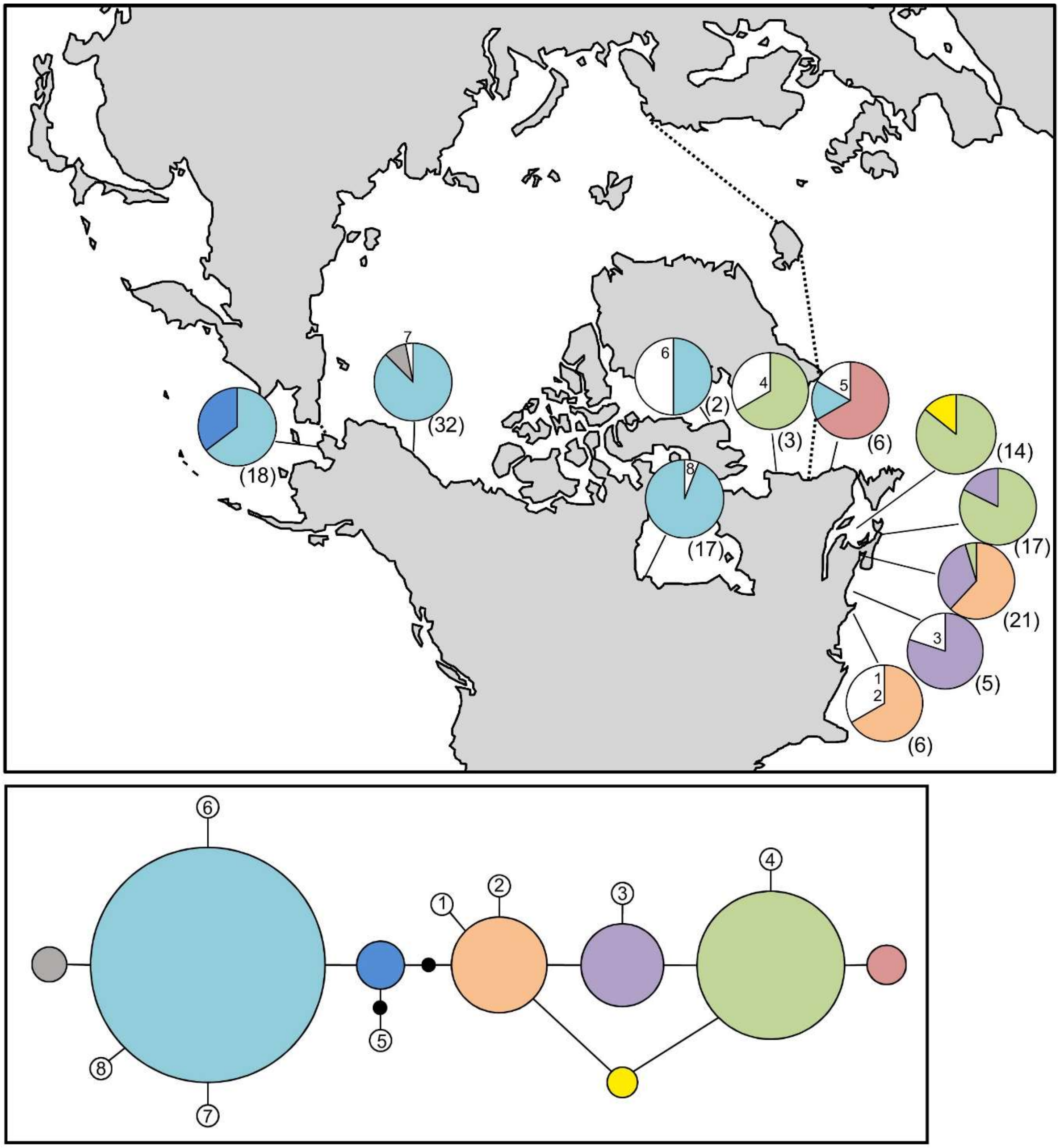

**Figure S26.** *Phycodrys fimbriata* haplotype map and network based on COI-5P data. In the map, numbers in parentheses refer to sample sizes from given locales, whereas numbers adjacent to white portions of pie charts refer to a haplotype sampled only once in the accompanying network. The dashed line indicates delineation of the Arctic Ocean. In the haplotype network, numbered haplotypes in white reference back to the map. Black circles indicate hypothesized (e.g. unsampled) haplotypes between clades. Circle size is proportional to the sampling frequency of a given haplotype.

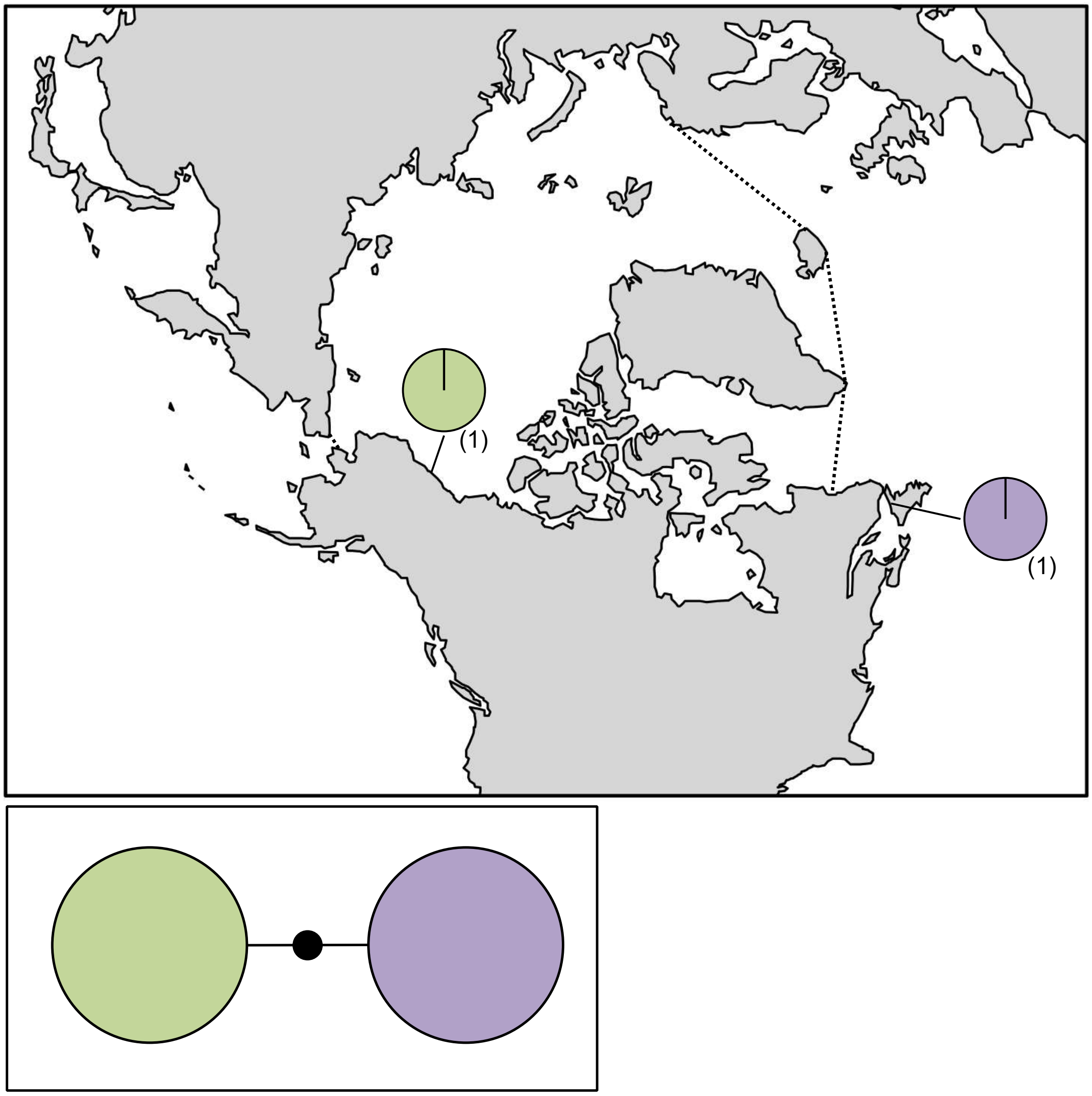

**Figure S27.** *Phymatolithon tenue* haplotype map and network based on COI-5P data. In the map, numbers in parentheses refer to sample sizes from given locales. The dashed line indicates delineation of the Arctic Ocean. In the haplotype network, the black circle represents a hypothesized (e.g. unsampled) haplotypes between clades. Circle size is proportional to the sampling frequency of a given haplotype.

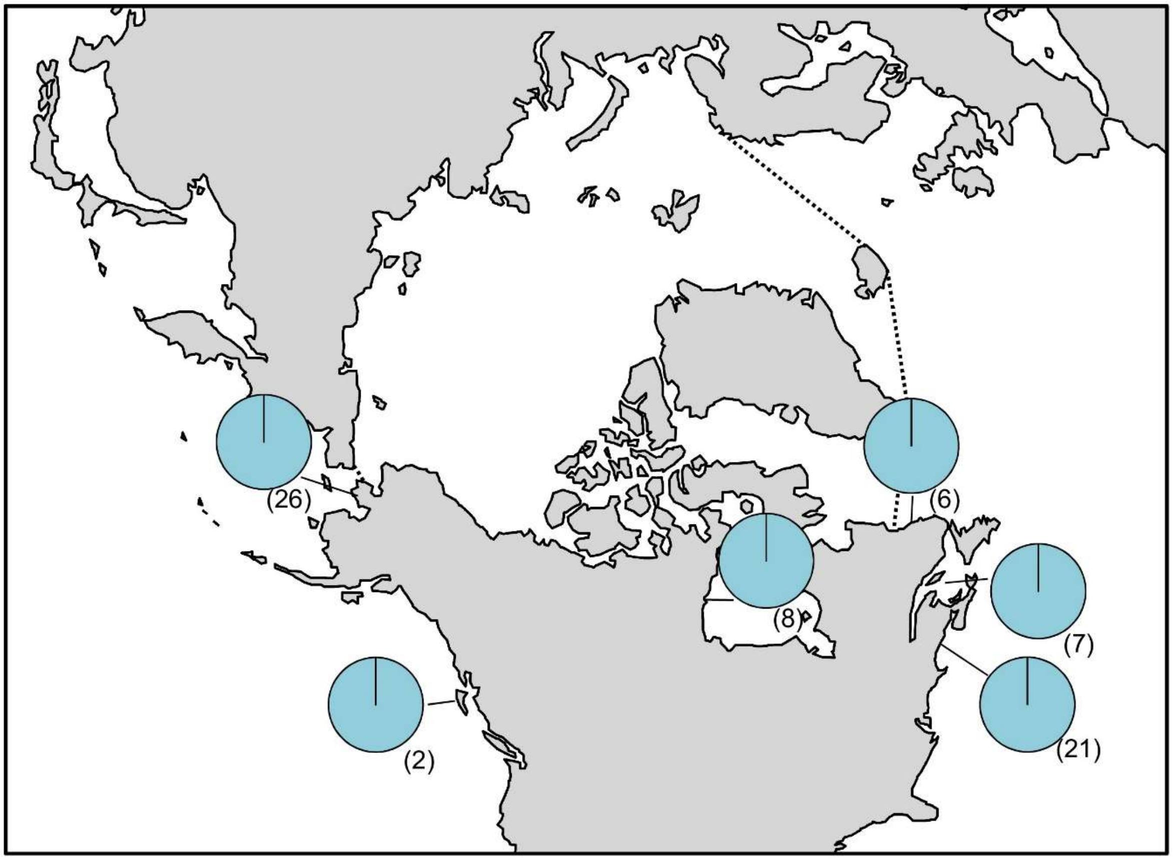

**Figure S28.** *Polysiphonia* sp. 1stricta haplotype map based on COI-5P data. In the map, numbers in parentheses refer to sample sizes from given locales. The dashed line indicates delineation of the Arctic Ocean.

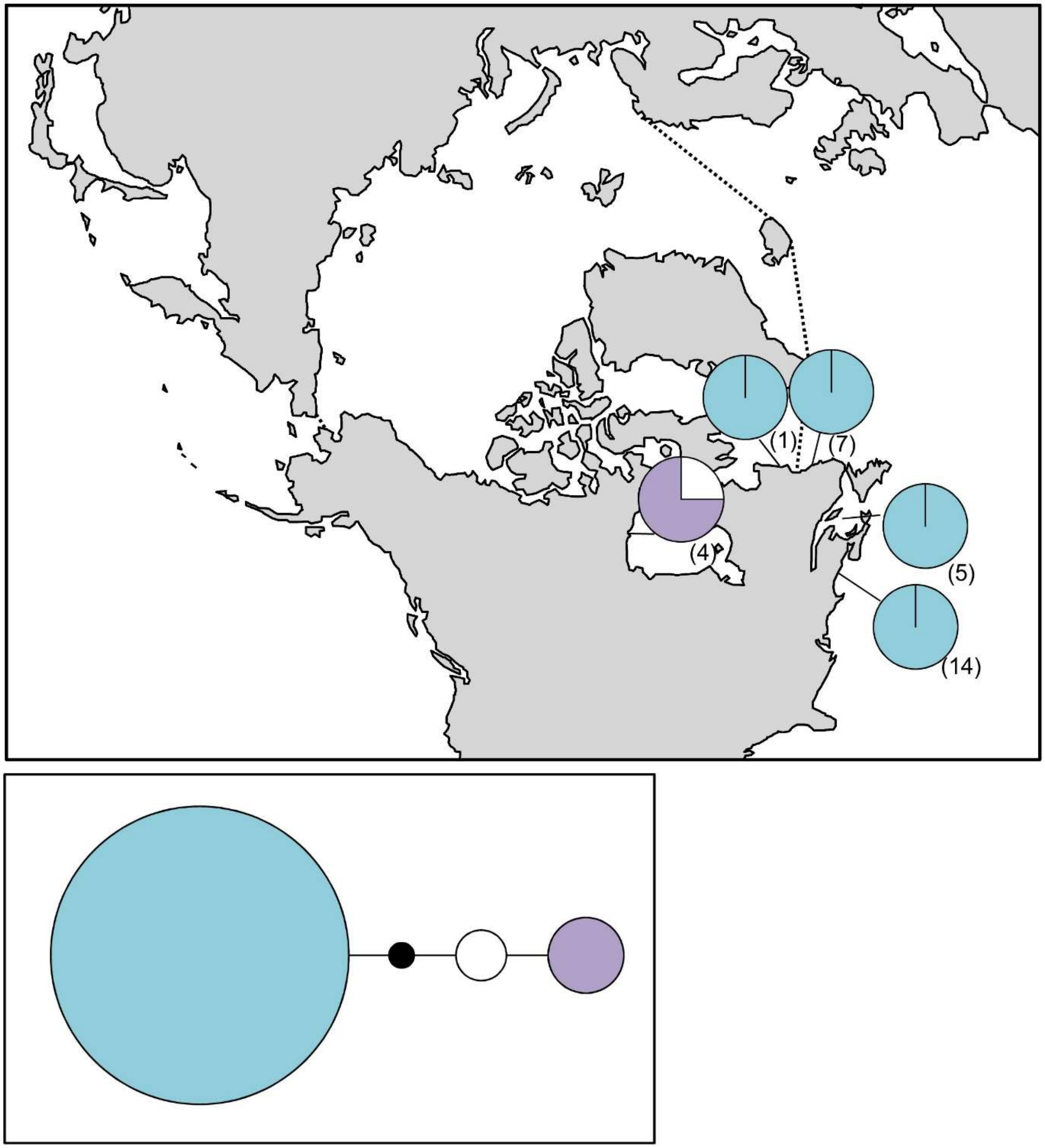

**Figure S29.** *Polysiphonia* sp. 3stricta map and network based on COI-5P data. In the map, numbers in parentheses refer to sample sizes from given locales. The dashed line indicates delineation of the Arctic Ocean. In the haplotype network, the black circle represents a hypothesized (e.g. unsampled) haplotypes between clades. Circle size is proportional to the sampling frequency of a given haplotype.

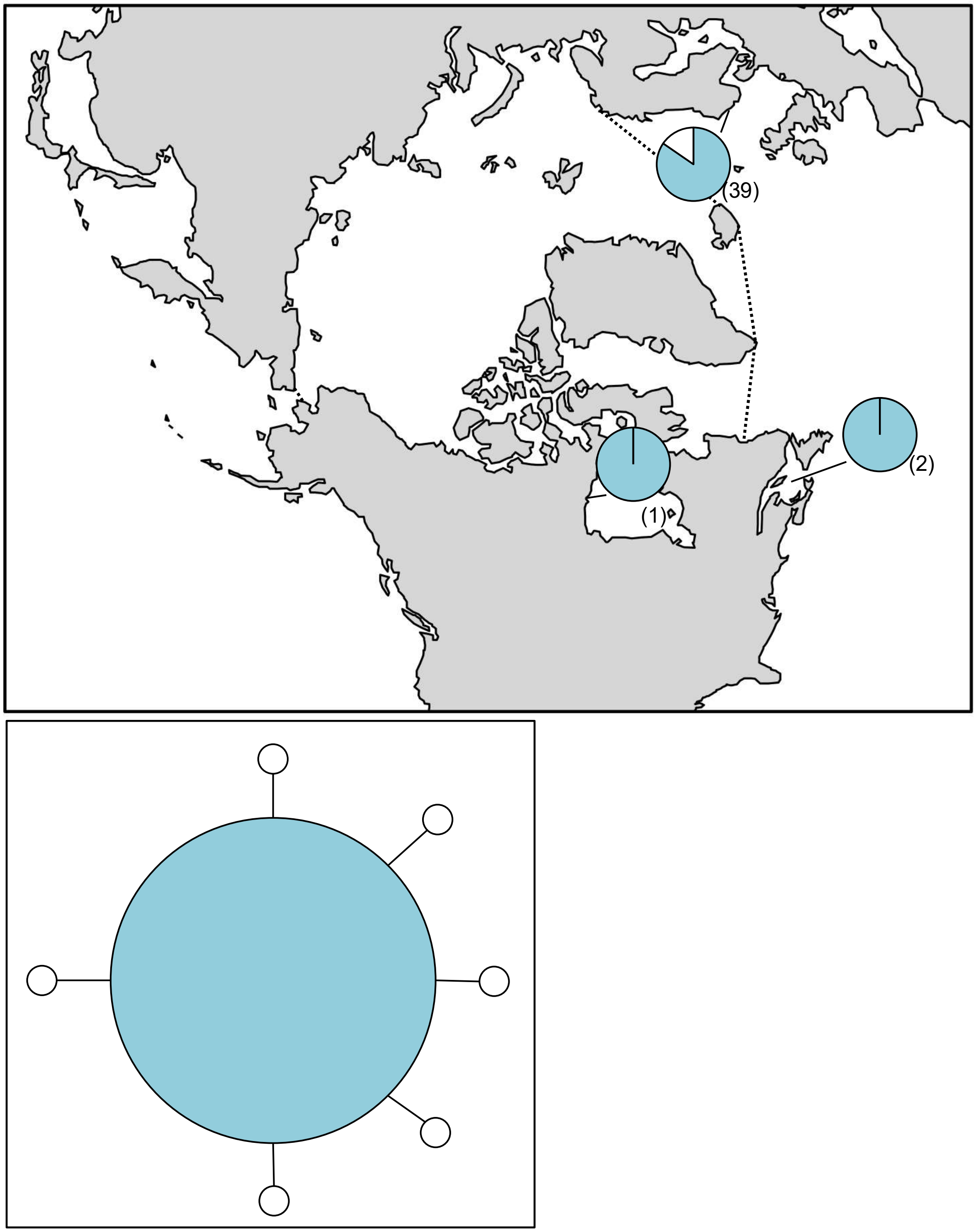

**Figure S30.** *Ptilota gunneri* haplotype map and network based on COI-5P data. In the map, numbers in parentheses refer to sample sizes from given locales. The dashed line indicates delineation of the Arctic Ocean. In the haplotype network, circle size is proportional to the sampling frequency of a given haplotype.

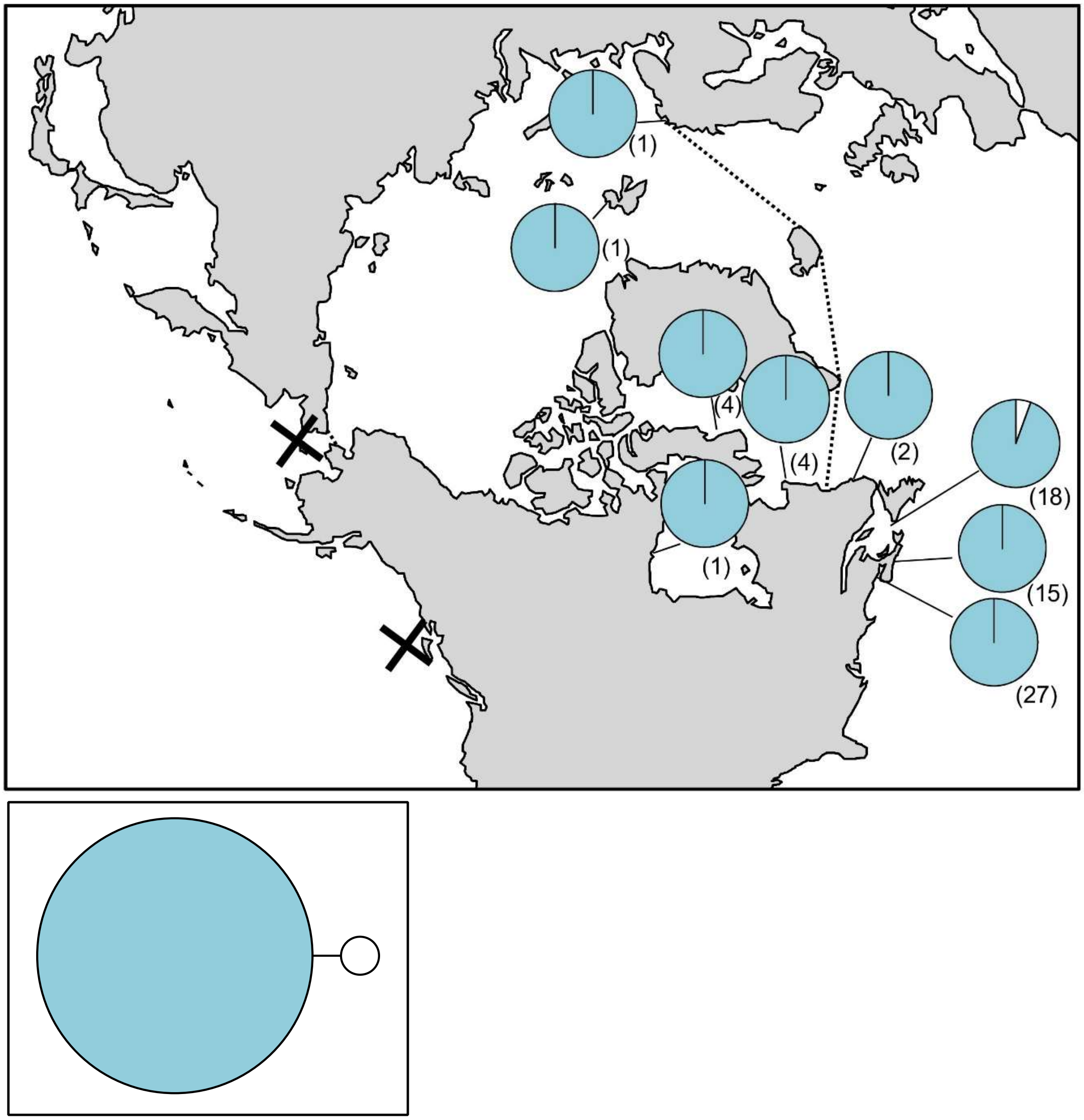

**Figure S31.** *Ptilota serrata* haplotype map and network based on COI-5P data. In the map, numbers in parentheses refer to sample sizes from given locales. The dashed line indicates delineation of the Arctic Ocean, while black X's indicate the locations of genetically verified *Ptilota serrata* based on *rbc*L. In the haplotype network, circle size is proportional to the sampling frequency of a given haplotype.

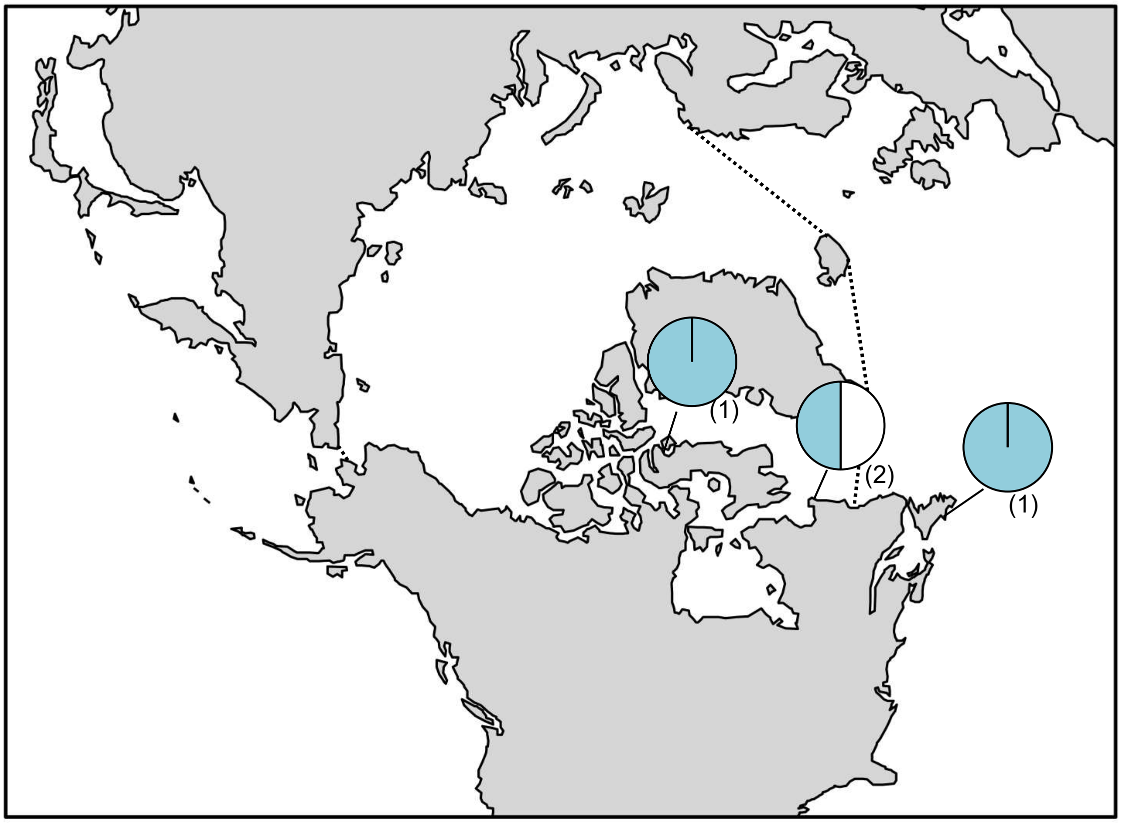

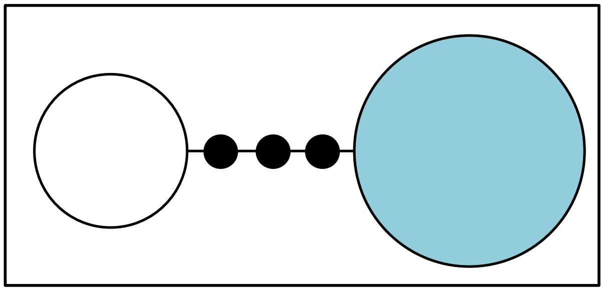

**Figure S32.** *Rhodochorton purpureum* haplotype map and network based on *rbc*L-3P data. In the map, numbers in parentheses refer to sample sizes from given locales. The dashed line indicates delineation of the Arctic Ocean. In the haplotype network. the black circles indicate hypothesized (e.g. unsampled) haplotypes between clades. Circle size is proportional to the sampling frequency of a given haplotype.

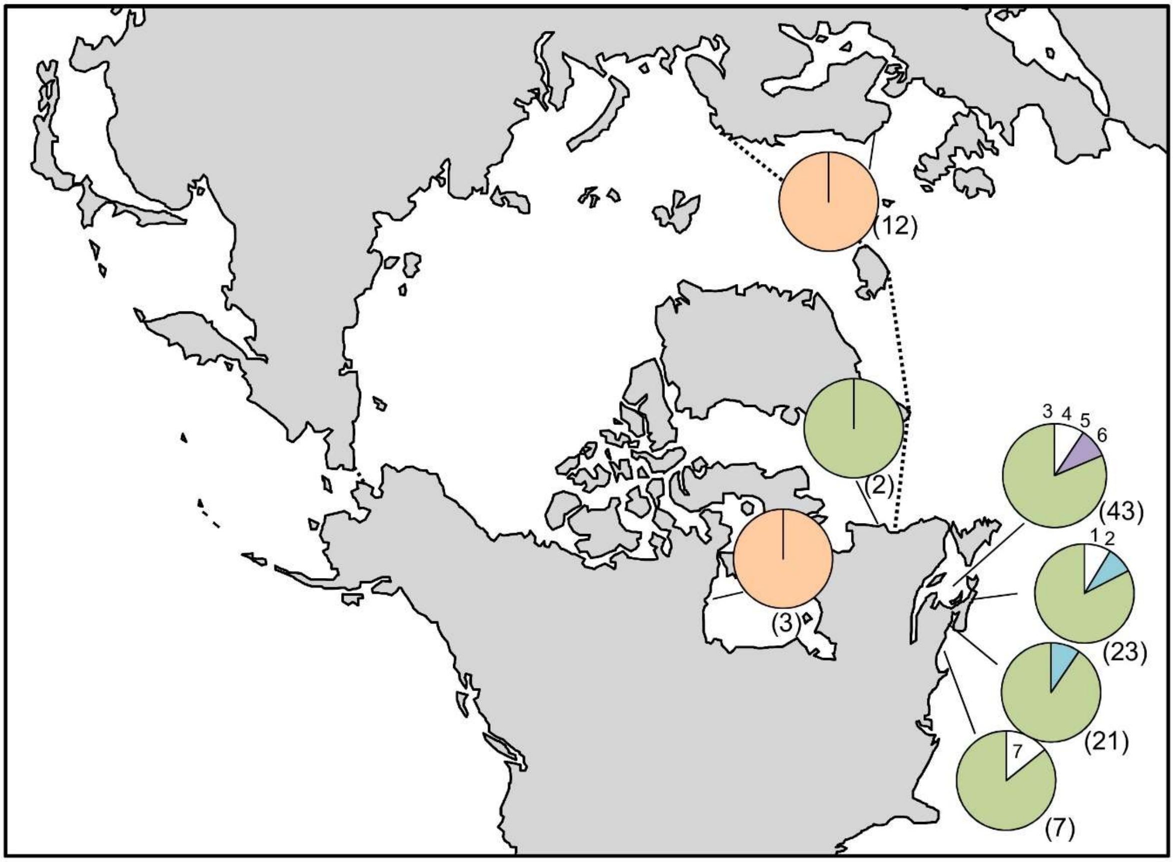

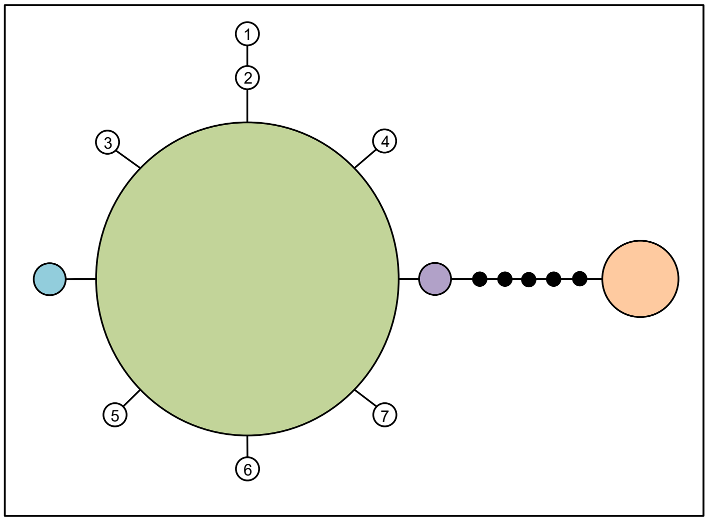

**Figure S33.** *Rhodomela lycopodioides* haplotype map and network based on COI-5P data. In the map, numbers in parentheses refer to sample sizes from given locales, whereas numbers adjacent to white portions of pie charts refer to a haplotype sampled only once in the accompanying network. The dashed line indicates delineation of the Arctic Ocean. In the haplotype network, numbered haplotypes in white reference back to the map. Black circles indicate hypothesized (e.g. unsampled) haplotypes between clades. Circle size is proportional to the sampling frequency of a given haplotype.

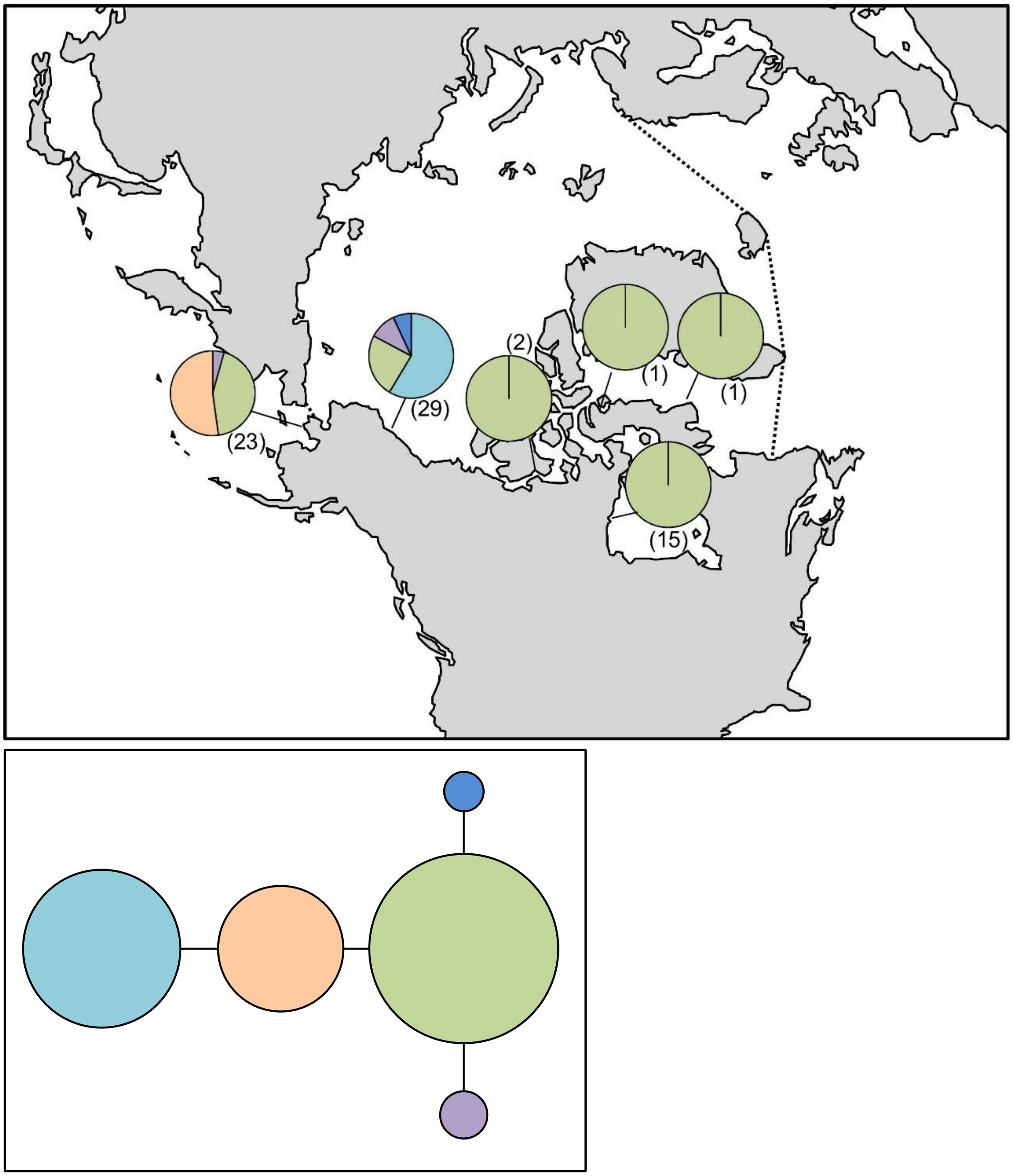

**Figure S34.** *Rhodomela sibirica* haplotype map and network based on COI-5P data. In the map, numbers in parentheses refer to sample sizes from given locales. The dashed line indicates delineation of the Arctic Ocean. In the haplotype network, circle size is proportional to the sampling frequency of a given haplotype.

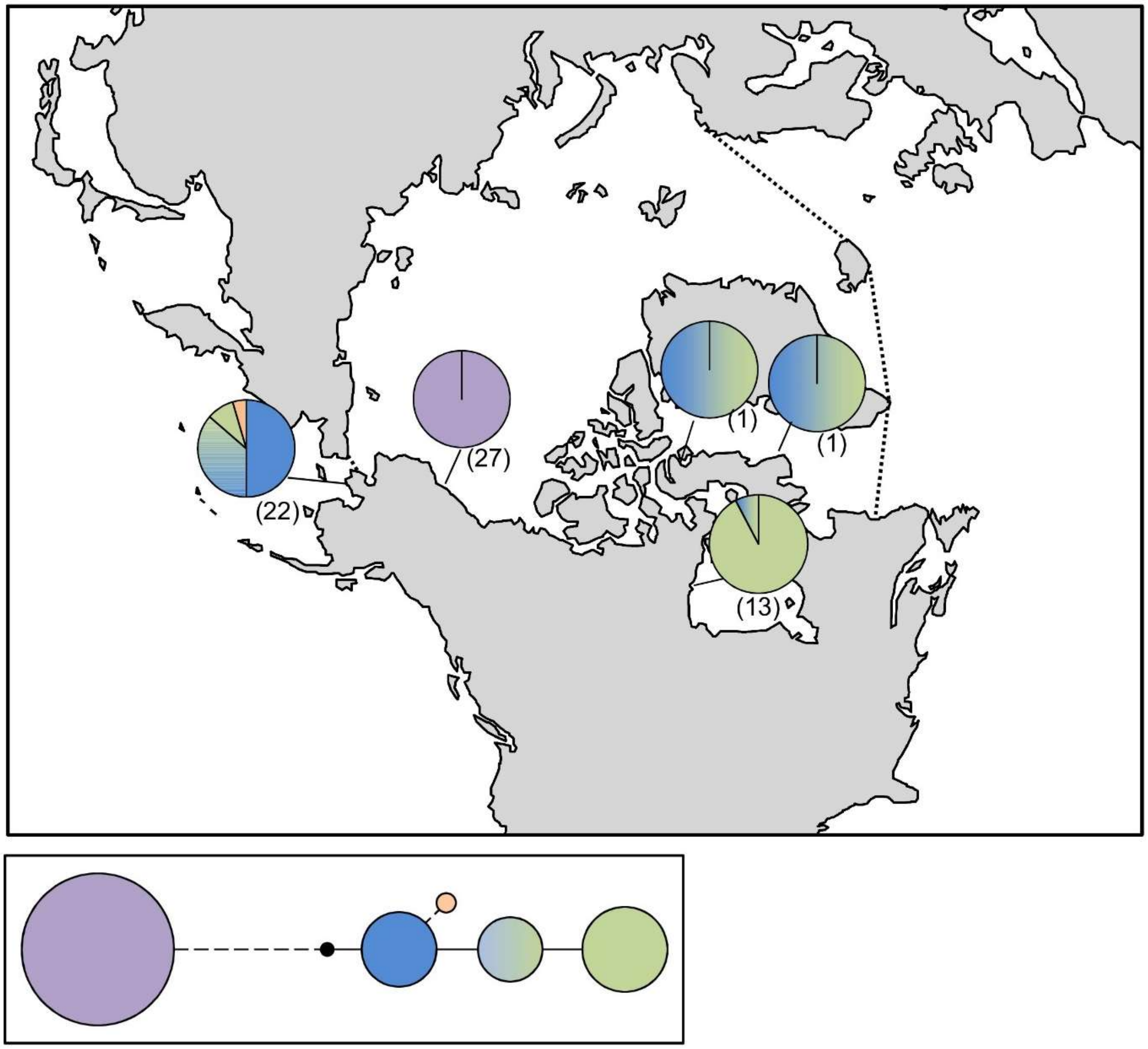

**Figure S35.** *Rhodomela sibirica* haplotype map and network based on ITS data. In the map, numbers in parentheses refer to sample sizes from given locales, the blue-green circle and pie chart segments indicate additivity in the ITS sequences, and the dashed line indicates delineation of the Arctic Ocean. In the haplotype network, the black circles indicate hypothesized (e.g. unsampled) haplotypes between clades, while the dashed line represents a 16 base-pair insert, and circle size is proportional to the sampling frequency of a given haplotype.

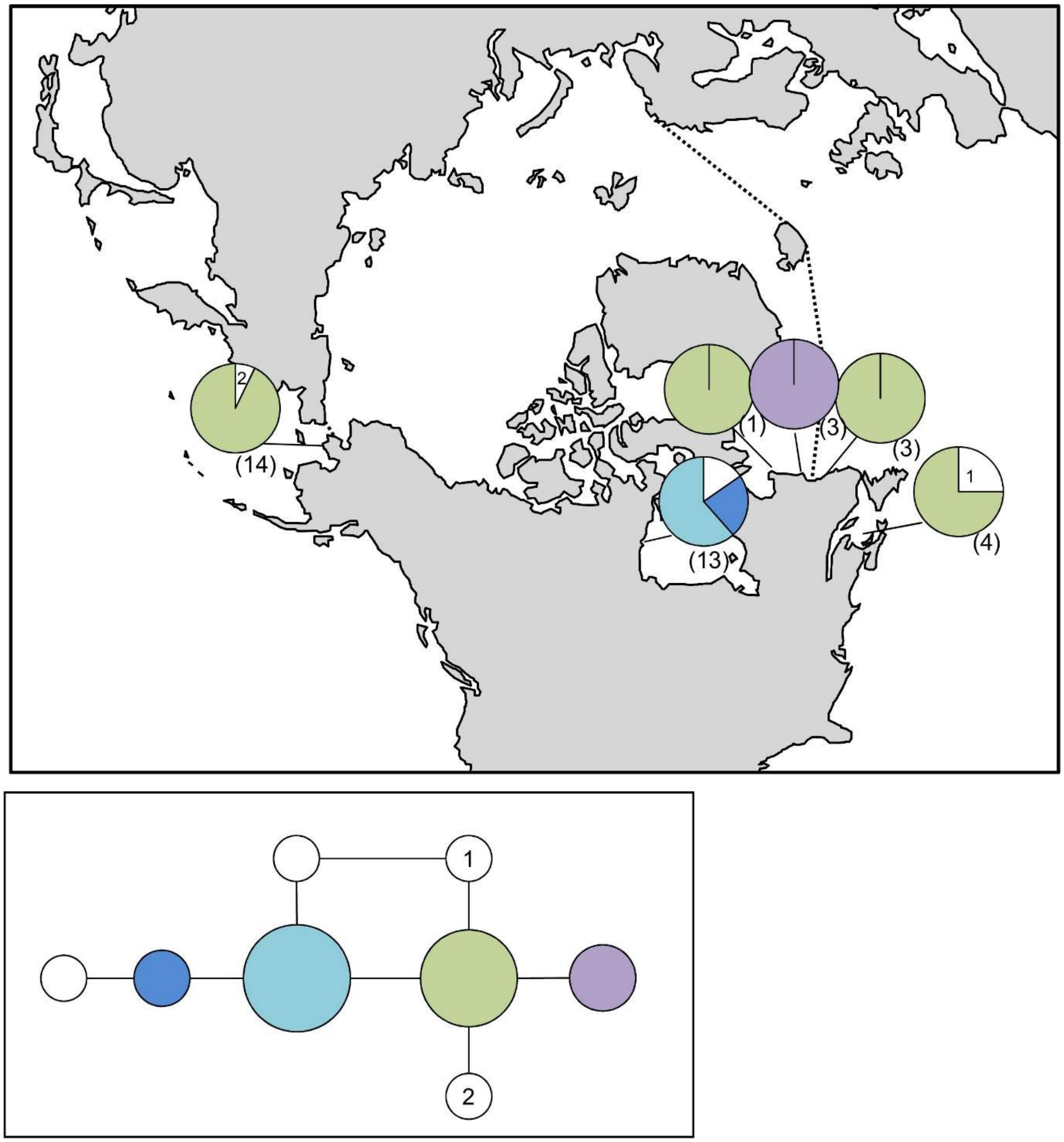

**Figure S36.** *Rhodomela* sp. 1virgata haplotype map and network based on COI-5P data. In the map, numbers in parentheses refer to sample sizes from given locales, whereas numbers adjacent to white portions of pie charts refer to a haplotype sampled only once in the accompanying network. The dashed line indicates delineation of the Arctic Ocean. In the haplotype network, numbered haplotypes in white reference back to the map. Circle size is proportional to the sampling frequency of a given haplotype.

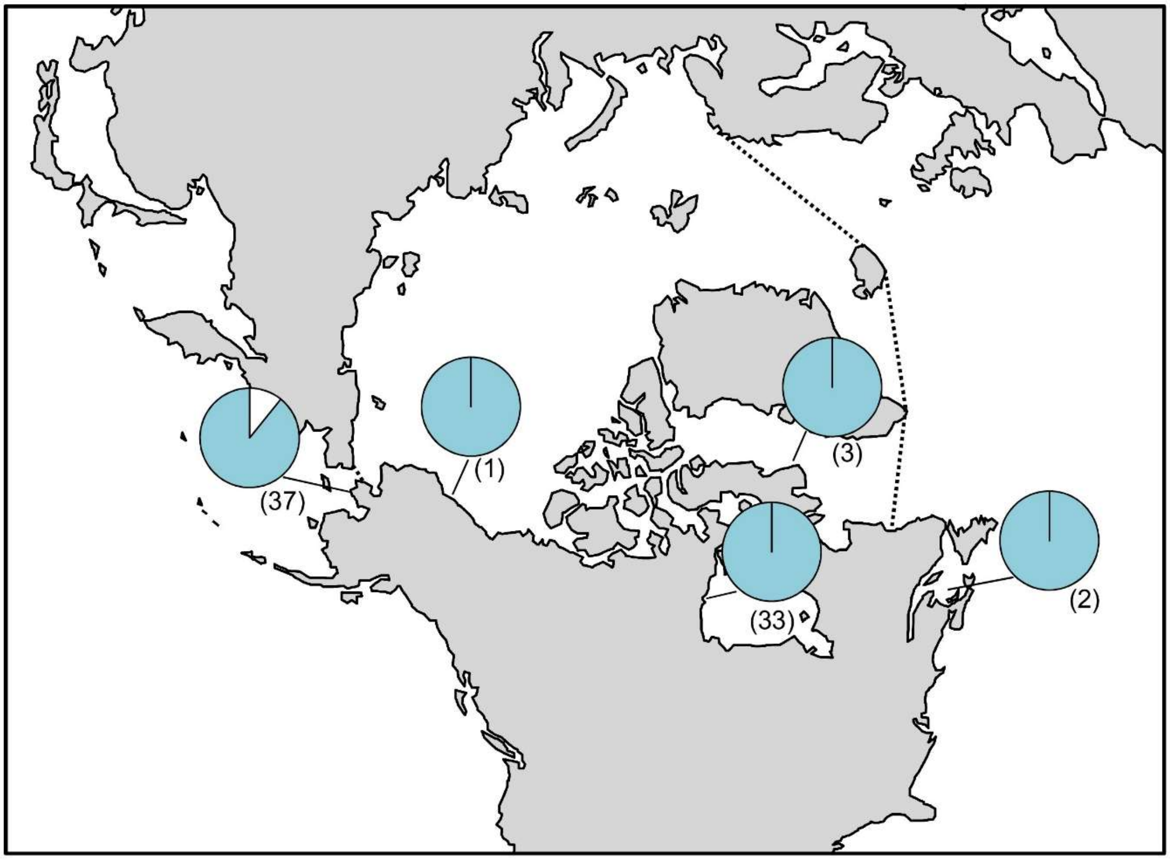

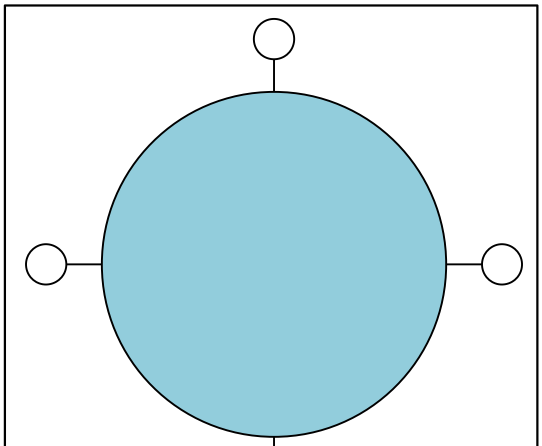

**Figure S37.** *Rhodomela virgata* haplotype map and network based on COI-5P data. In the map, numbers in parentheses refer to sample sizes from given locales. The dashed line indicates delineation of the Arctic Ocean. In the haplotype network, circle size is proportional to the sampling frequency of a given haplotype.

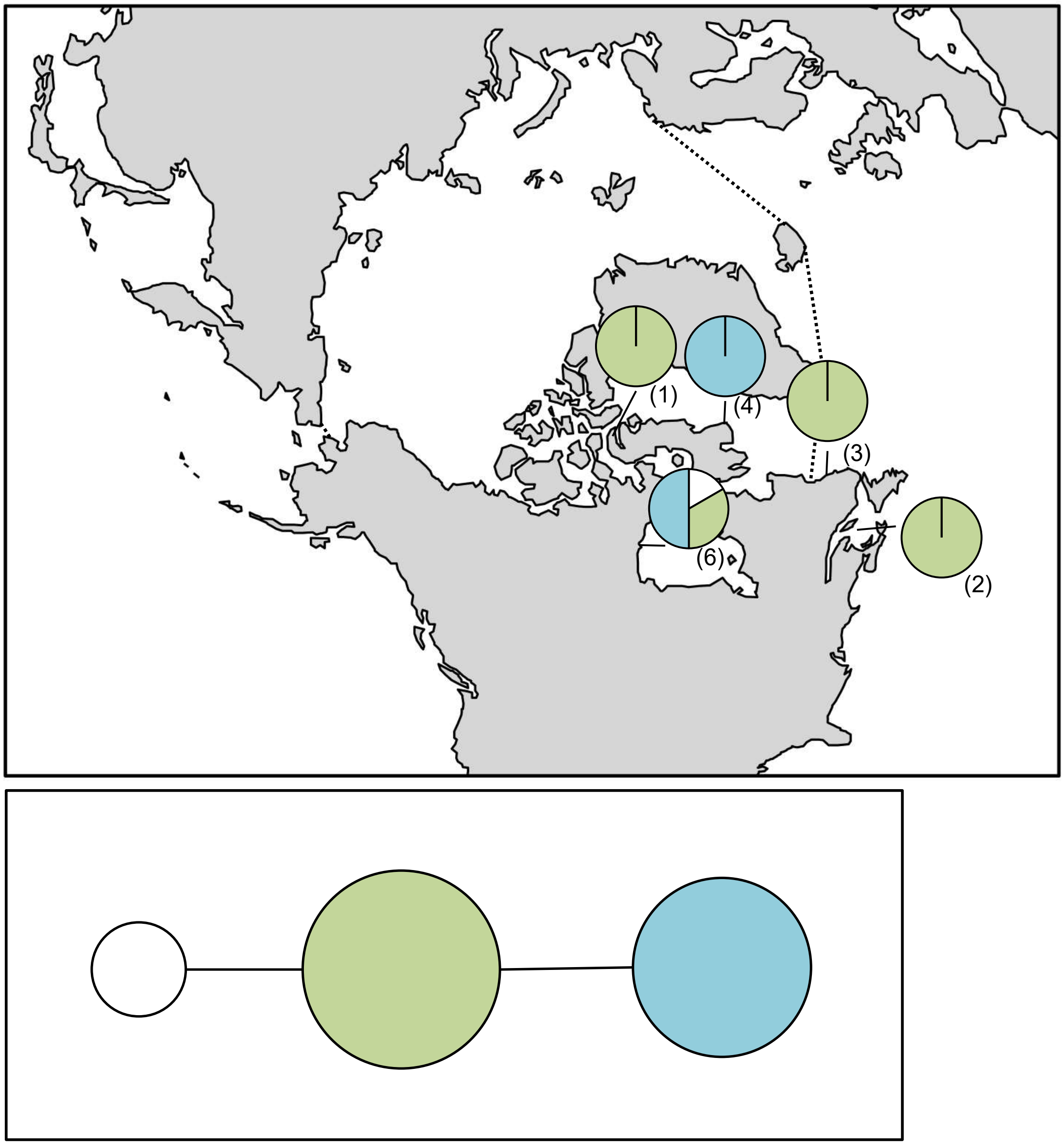

**Figure S38.** *Savoiea arctica* haplotype map and network based on COI-5P data. In the map, numbers in parentheses refer to sample sizes from given locales. The dashed line indicates delineation of the Arctic Ocean. In the haplotype network, circle size is proportional to the sampling frequency of a given haplotype.

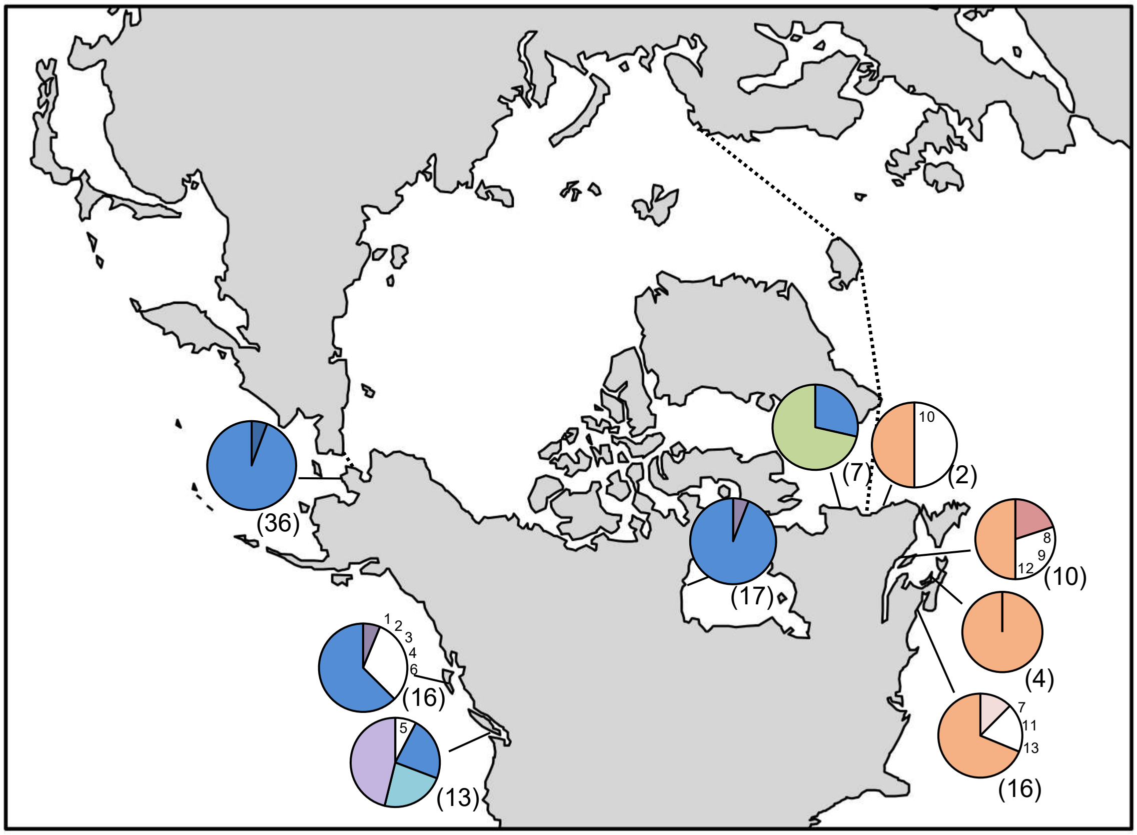

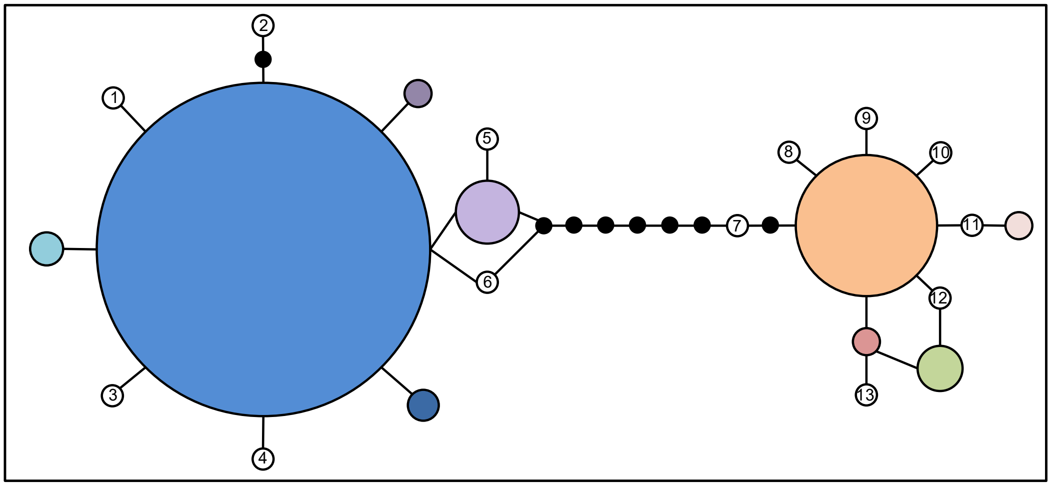

**Figure S39.** *Scagelia pylaisaei* haplotype map and network based on COI-5P data. In the map, numbers in parentheses refer to sample sizes from given locales, whereas numbers adjacent to white portions of pie charts refer to a haplotype sampled only once in the accompanying network. The dashed line indicates delineation of the Arctic Ocean. In the haplotype network, numbered haplotypes in white reference back to the map. Black circles indicate hypothesized (e.g. unsampled) haplotypes between clades. Circle size is proportional to the sampling frequency of a given haplotype.

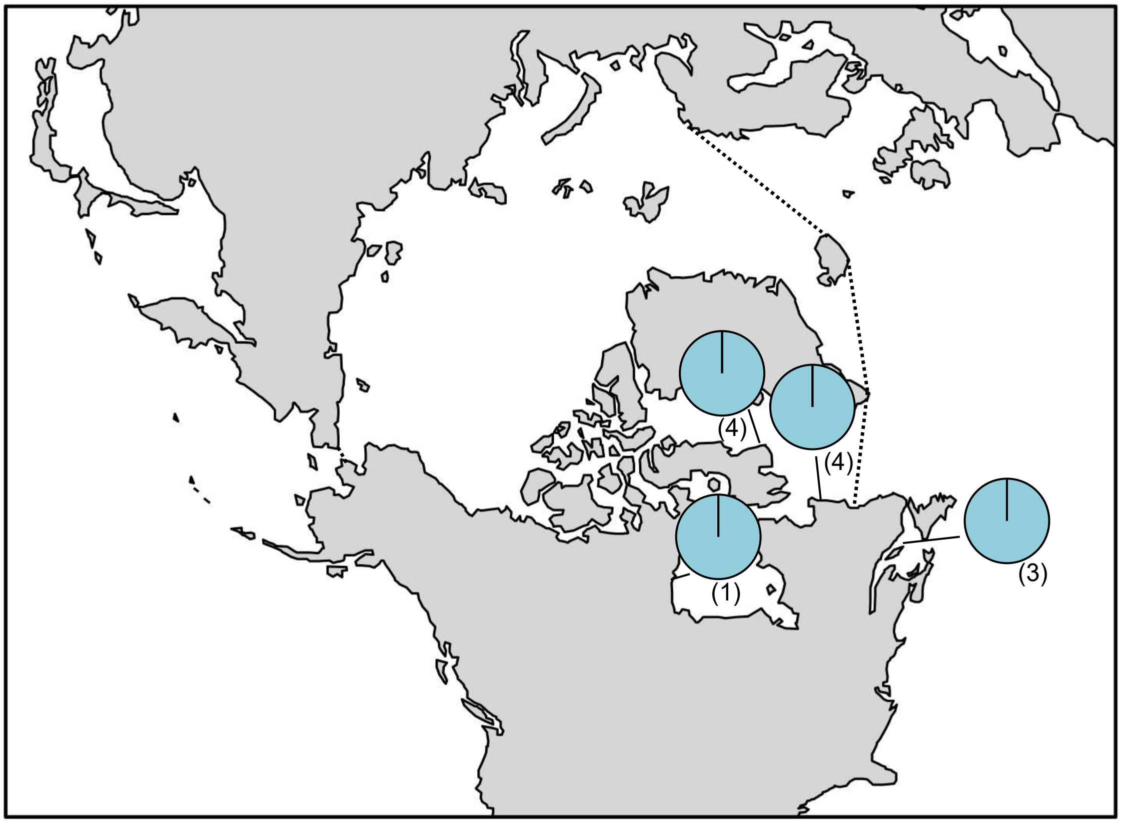

**Figure S40.** *Turnerella pennyi* haplotype map based on *rbc*L-3P data. In the map, numbers in parentheses refer to sample sizes from given locales. The dashed line indicates delineation of the Arctic Ocean.

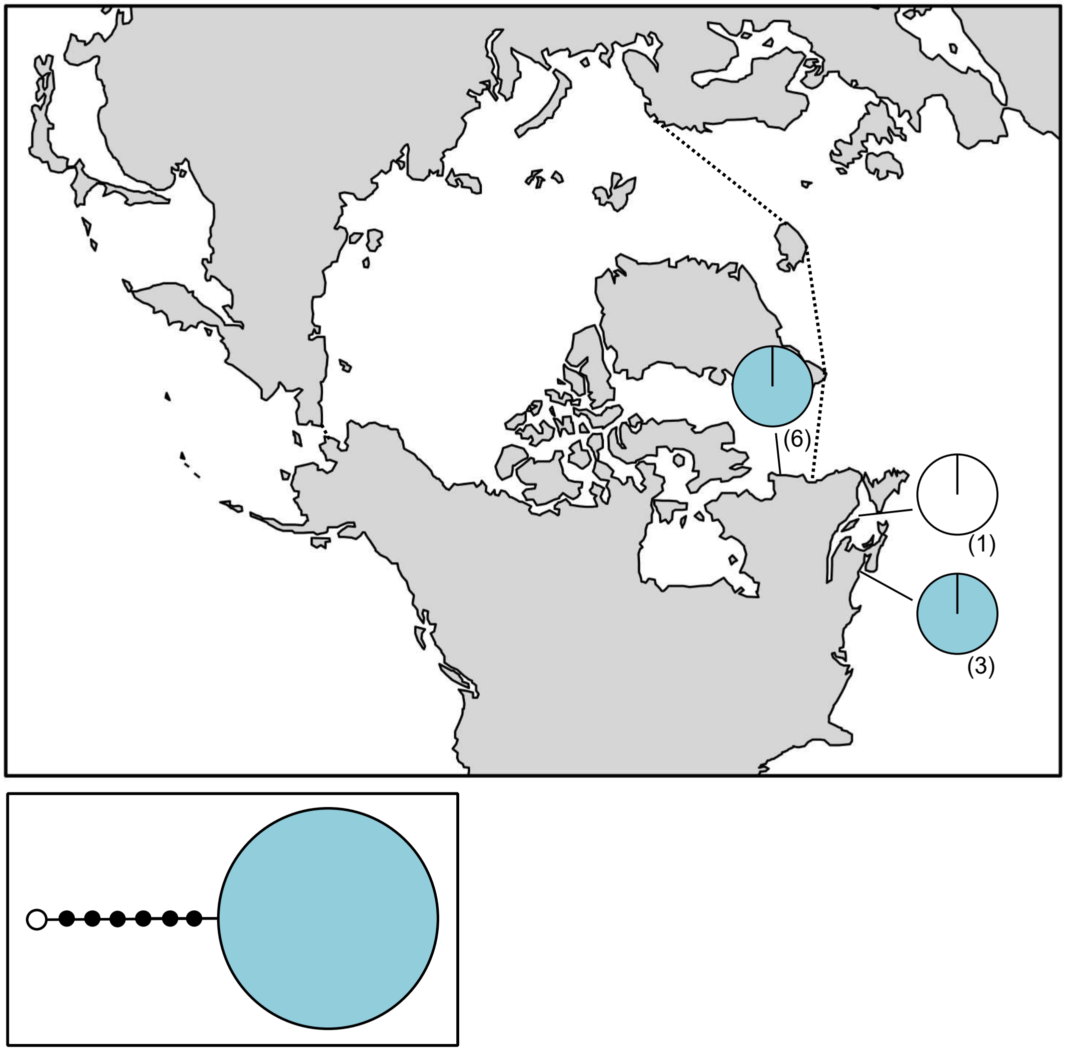

**Figure S41.** *Waernia mirabilis* haplotype map based on COI-5P data. In the map, numbers in parentheses refer to sample sizes from given locales. The dashed line indicates delineation of the Arctic Ocean. In the haplotype network, black circles indicate hypothesized (e.g. unsampled) haplotypes between clades. Circle size is proportional to the sampling frequency of a given haplotype.

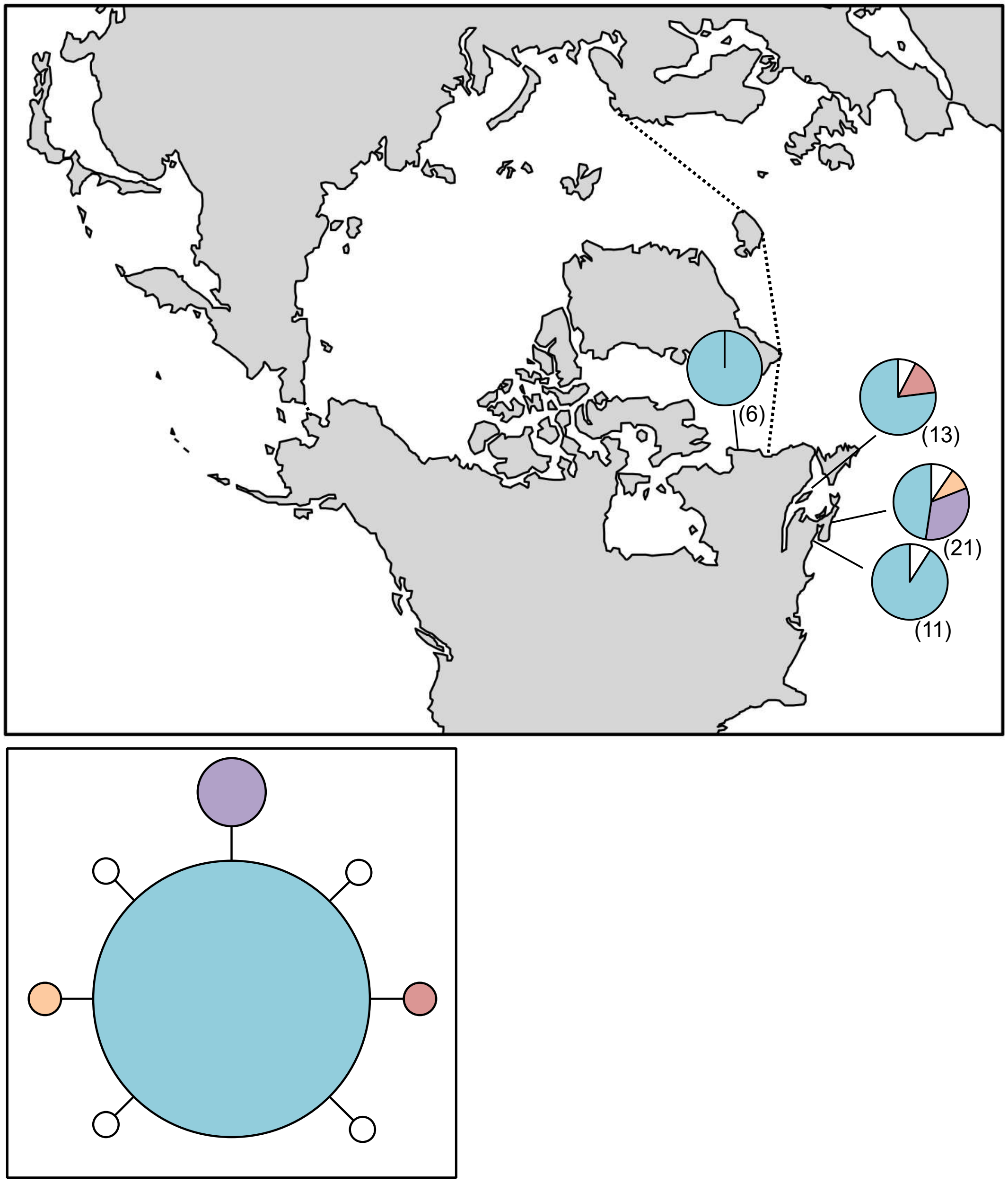

**Figure S42.** *Wildemania miniata* haplotype map based on COI-5P data. In the map, numbers in parentheses refer to sample sizes from given locales. The dashed line indicates delineation of the Arctic Ocean. In the haplotype network, circle size is proportional to the sampling frequency of a given haplotype.

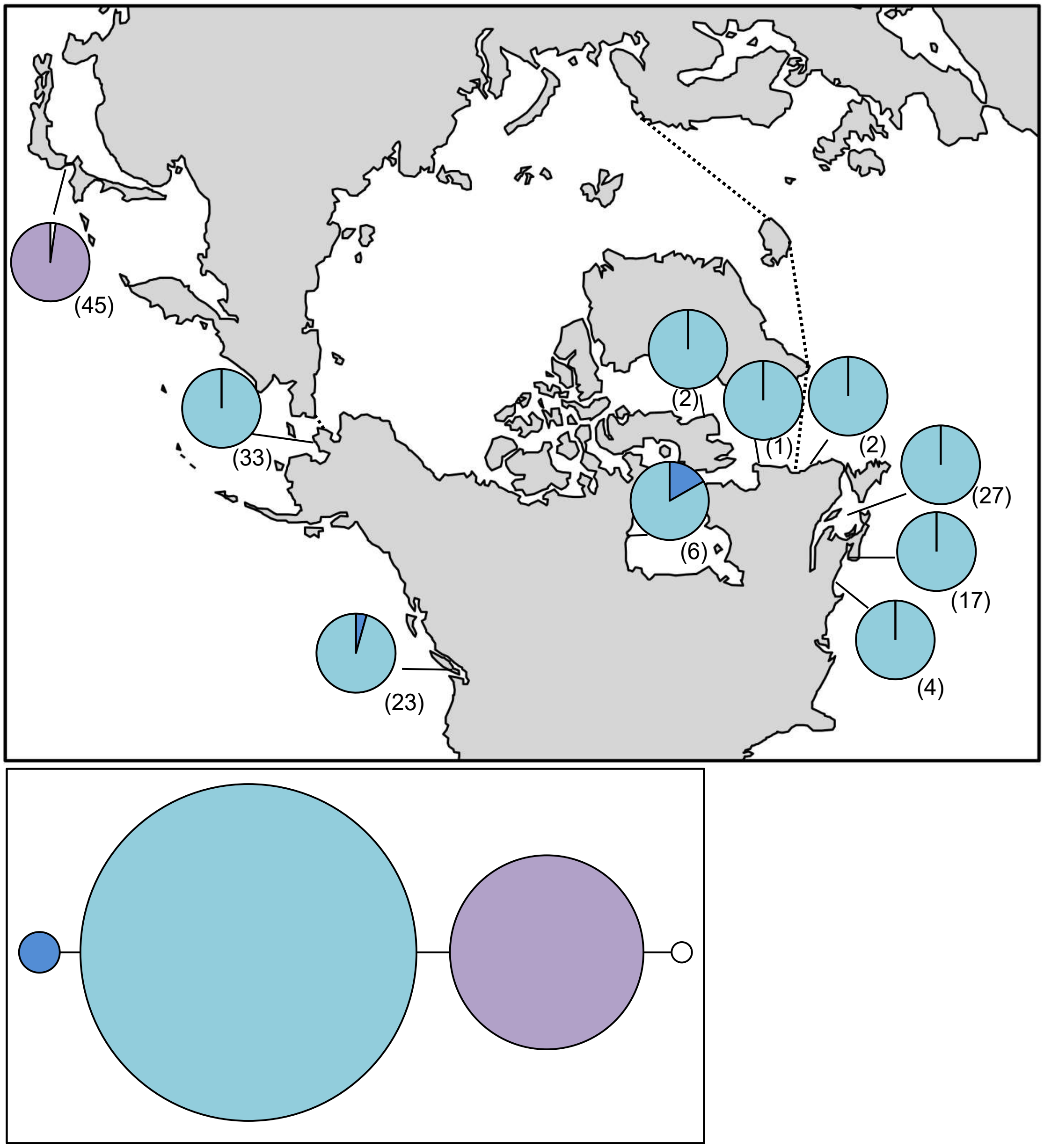

**Figure S43.** *Agarum clathratum* haplotype map and network based on COI-5P data. In the map, numbers in parentheses refer to sample sizes from given locales. The dashed line indicates delineation of the Arctic Ocean. In the haplotype network, circle size is proportional to the sampling frequency of a given haplotype.

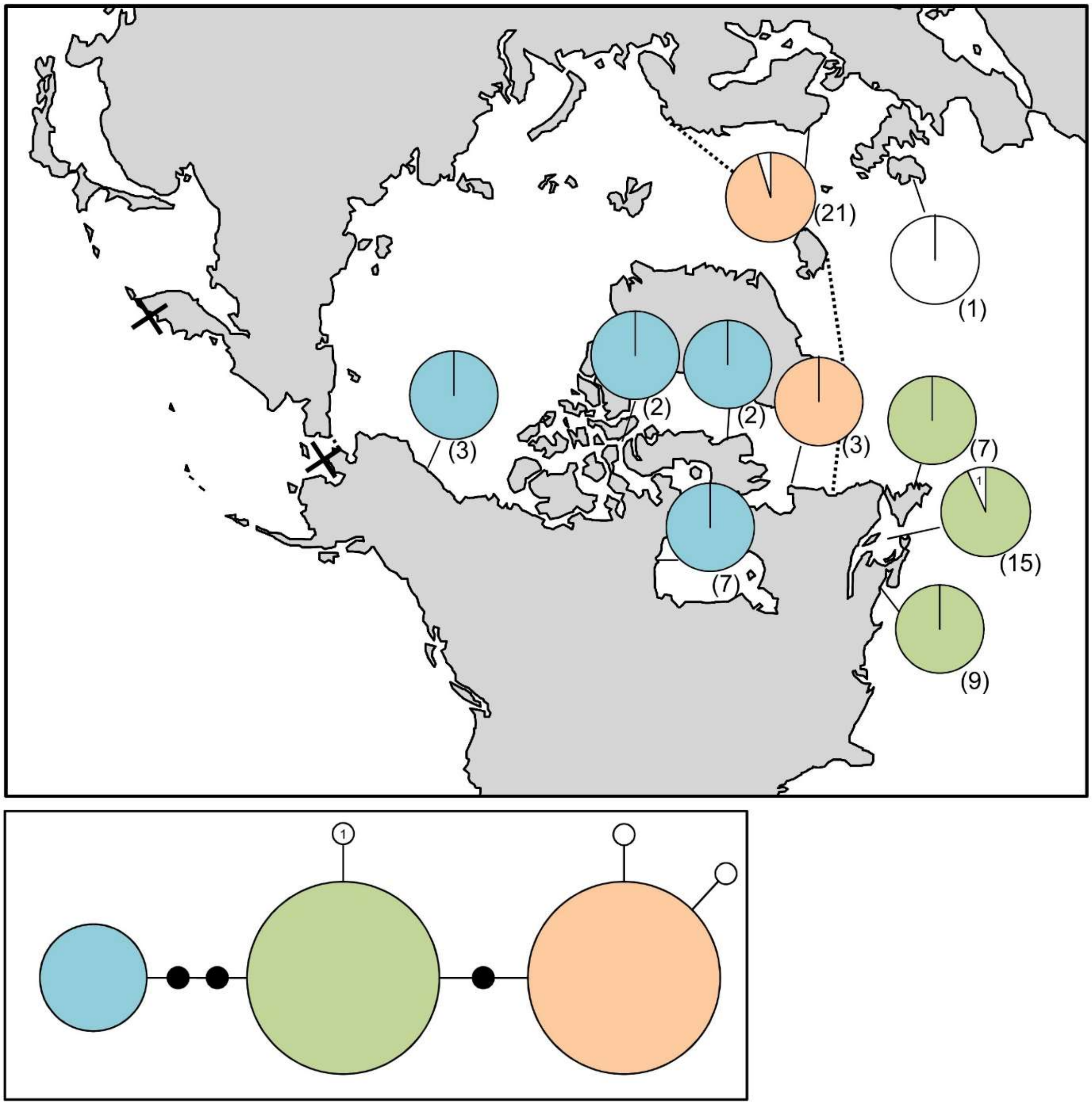

**Figure S44.** haplotype map and network based on COI-5P data. In the map, numbers in parentheses refer to sample sizes from given locales, whereas numbers adjacent to white portions of pie charts refer to a haplotype sampled only once in the accompanying network. The dashed line indicates delineation of the Arctic Ocean. In the haplotype network, numbered haplotypes in white reference back to the map. Black circles indicate hypothesized (e.g. unsampled) haplotypes between clades. Circle size is proportional to the sampling frequency of a given haplotype. The locations marked X indicate genetically confirmed locations of *A. esculenta* based on ITS data, however, COI-5P provides a conflicting ID, indicating the presence of *Alaria crispa*.

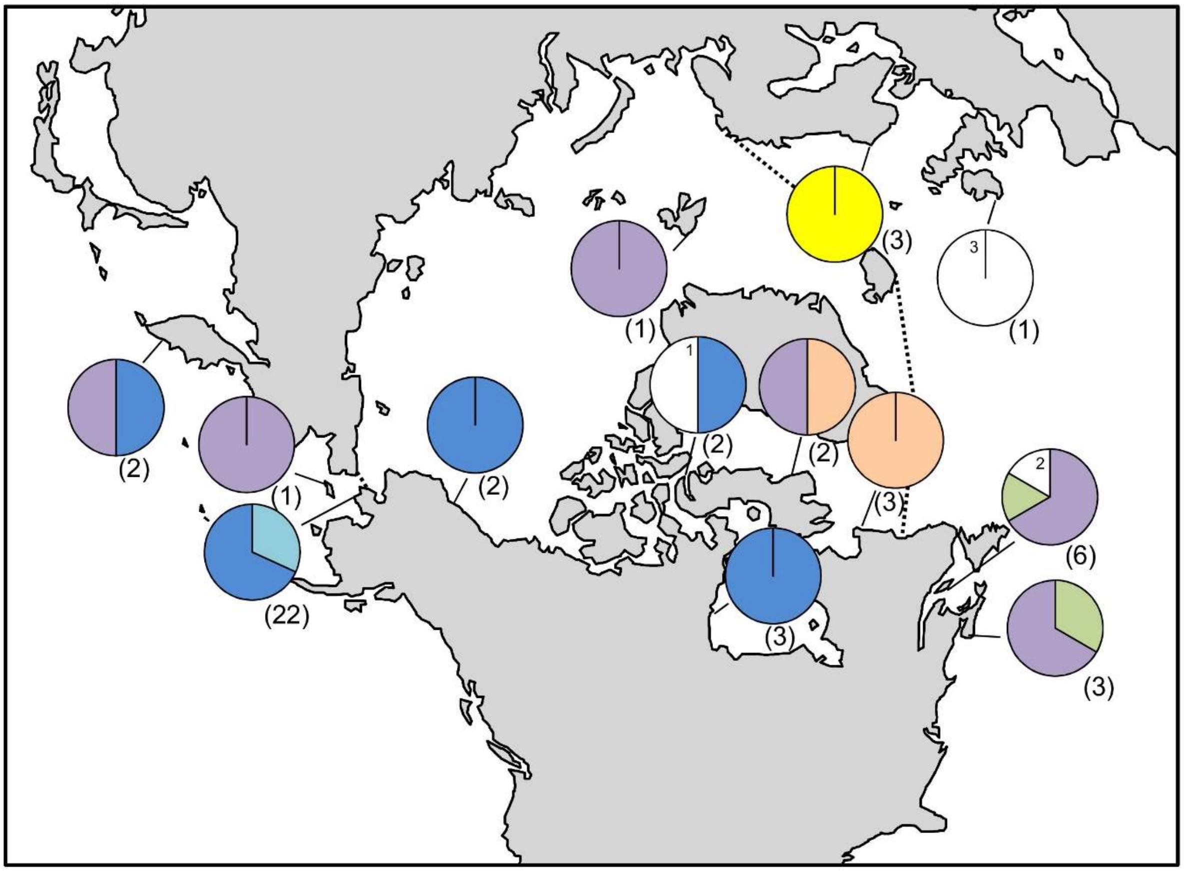

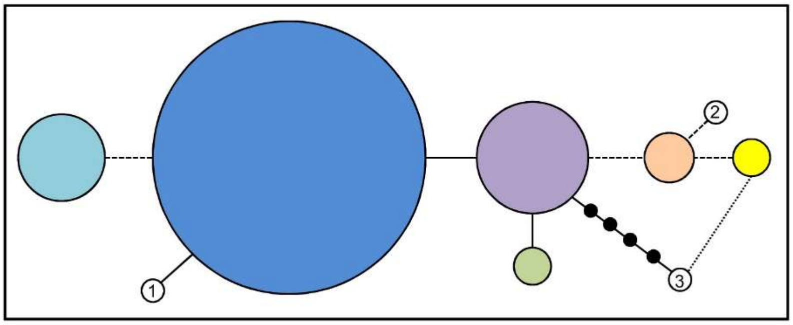

**Figure S45.** *Alaria esculenta* haplotype map and network based on ITS data. In the map, numbers in parentheses refer to sample sizes from given locales, whereas numbers adjacent to white portions of pie charts refer to a haplotype sampled only once in the accompanying network. The dashed line indicates delineation of the Arctic Ocean. In the haplotype network, numbered haplotypes in white reference back to the map. Black circles indicate hypothesized (e.g. unsampled) haplotypes between clades. Circle size is proportional to the sampling frequency of a given haplotype. Dashed lines indicated insertions/deletions, while the square dot line indicates a substitution shared at the same site between two haplotypes.

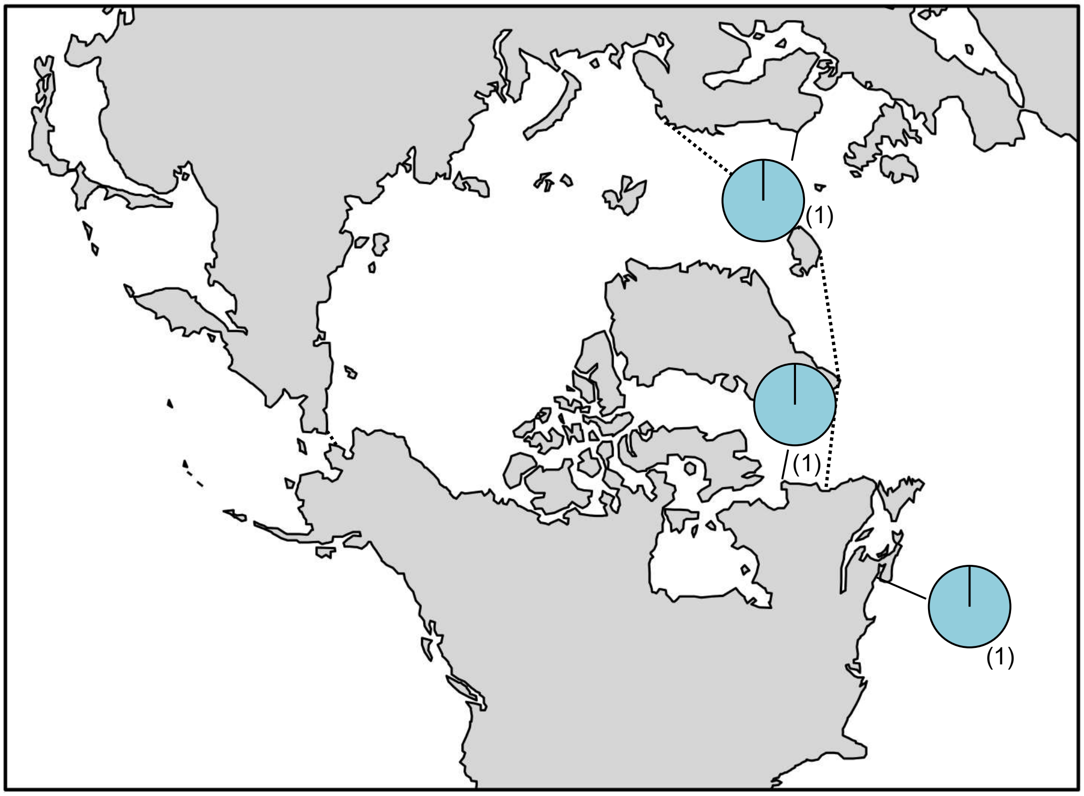

**Figure S46.** *Ascophyllum nodosum* haplotype map based on COI-5P data. In the map, numbers in parentheses refer to sample sizes from given locales. The dashed line indicates delineation of the Arctic Ocean.

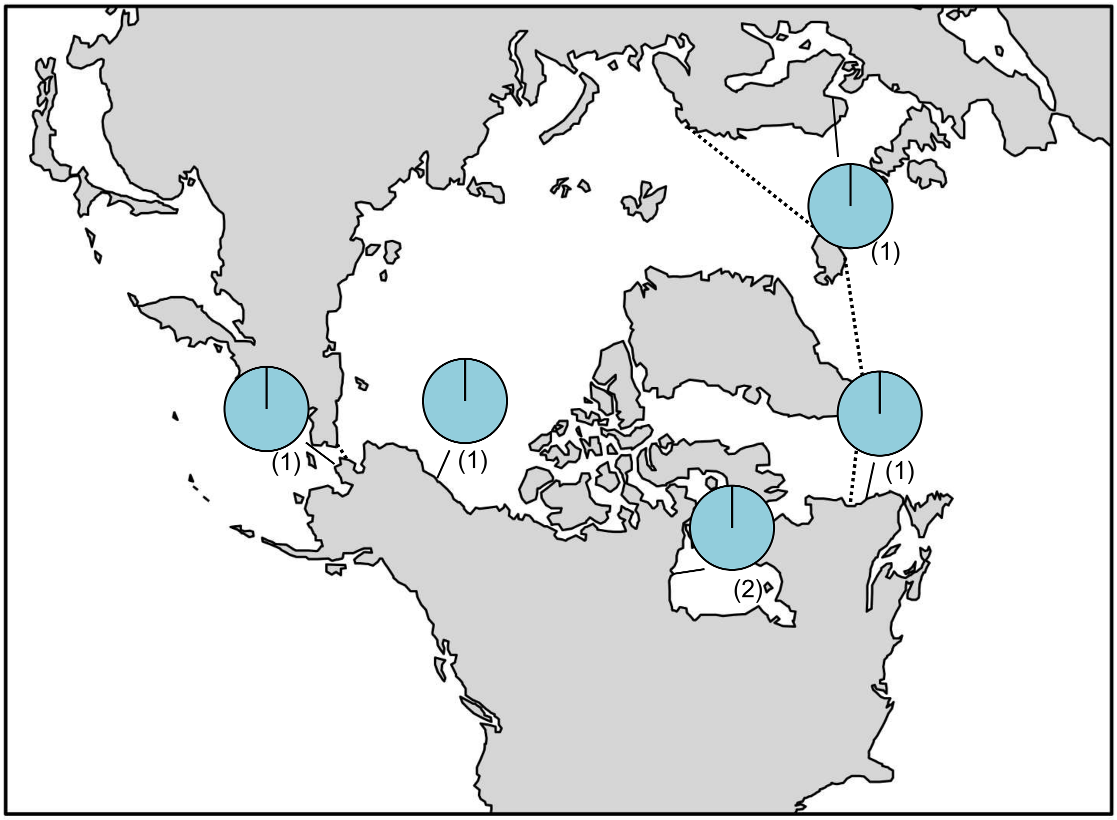

**Figure S47.** *Battersia arctica* haplotype map based on *rbc*L-3P data. In the map, numbers in parentheses refer to sample sizes from given locales. The dashed line indicates delineation of the Arctic Ocean.

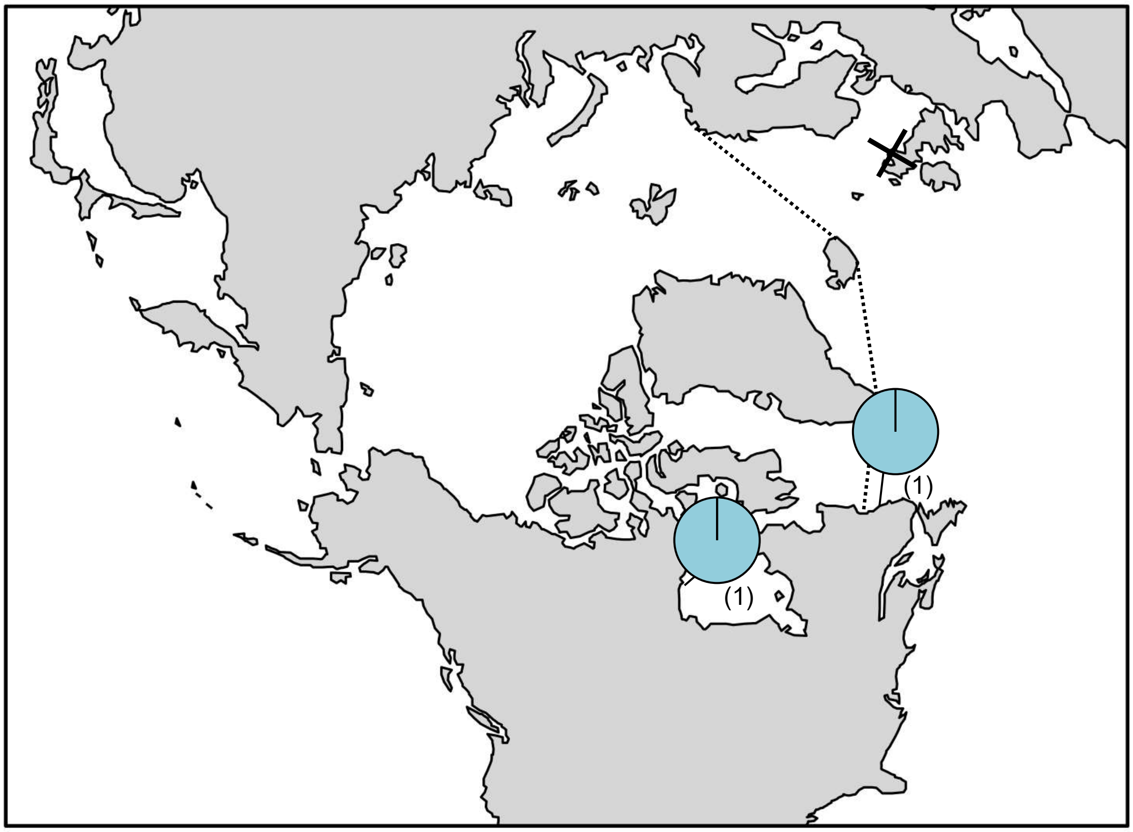

**Figure S48.** *Battersia racemosa* haplotype map based on COI-5P data. In the map, numbers in parentheses refer to sample sizes from given locales. The dashed line indicates delineation of the Arctic Ocean, while the black X indicates the location of genetically verified *Battersia racemosa* based on *rbc*L.

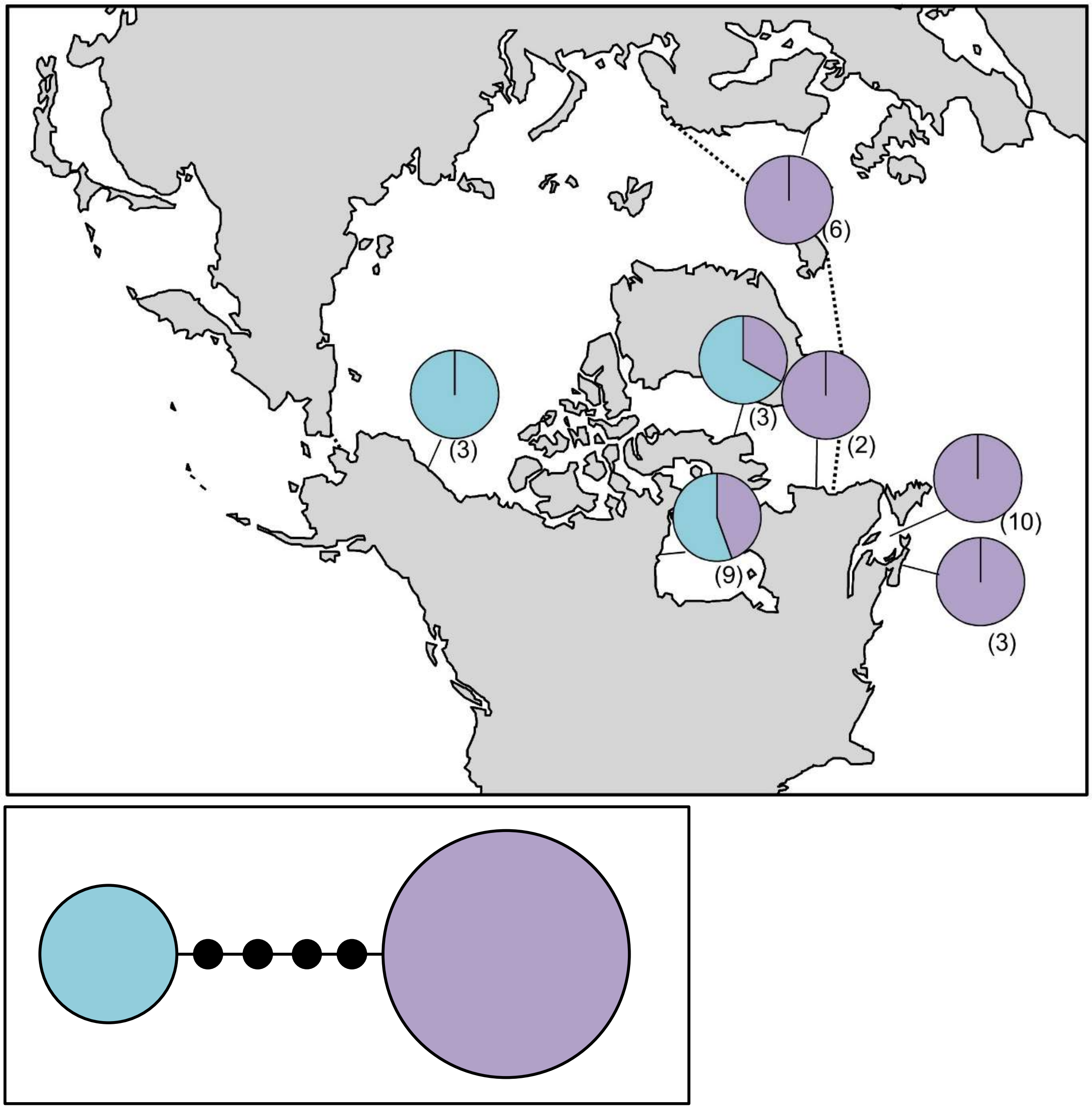

**Figure S49.** *Chaetopteris plumosa* haplotype map and network based on COI-5P data. In the map, numbers in parentheses refer to sample sizes from given locales. The dashed line indicates delineation of the Arctic Ocean. In the haplotype network, black circles indicate hypothesized (e.g. unsampled) haplotypes between clades. Circle size is proportional to the sampling frequency of a given haplotype.

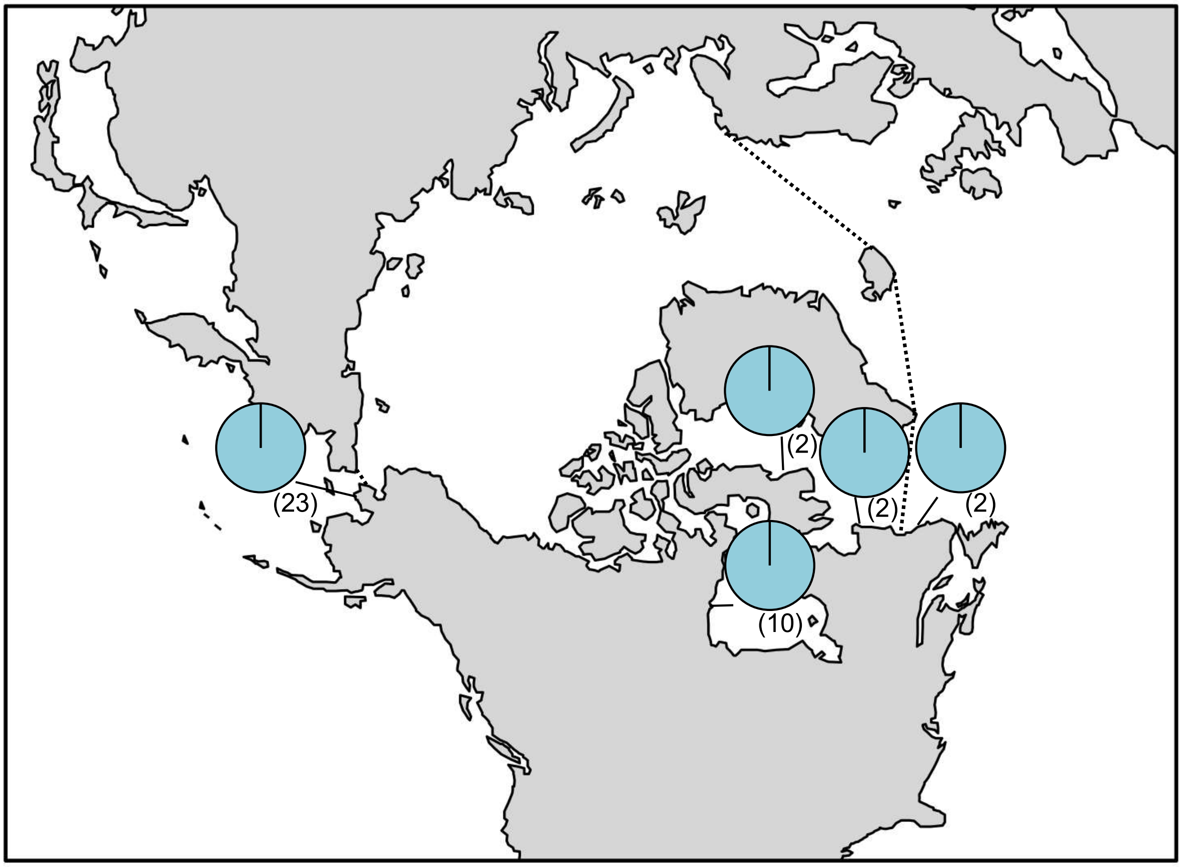

**Figure S50.** *Chorda borealis* haplotype map based on COI-5P data. In the map, numbers in parentheses refer to sample sizes from given locales. The dashed line indicates delineation of the Arctic Ocean.

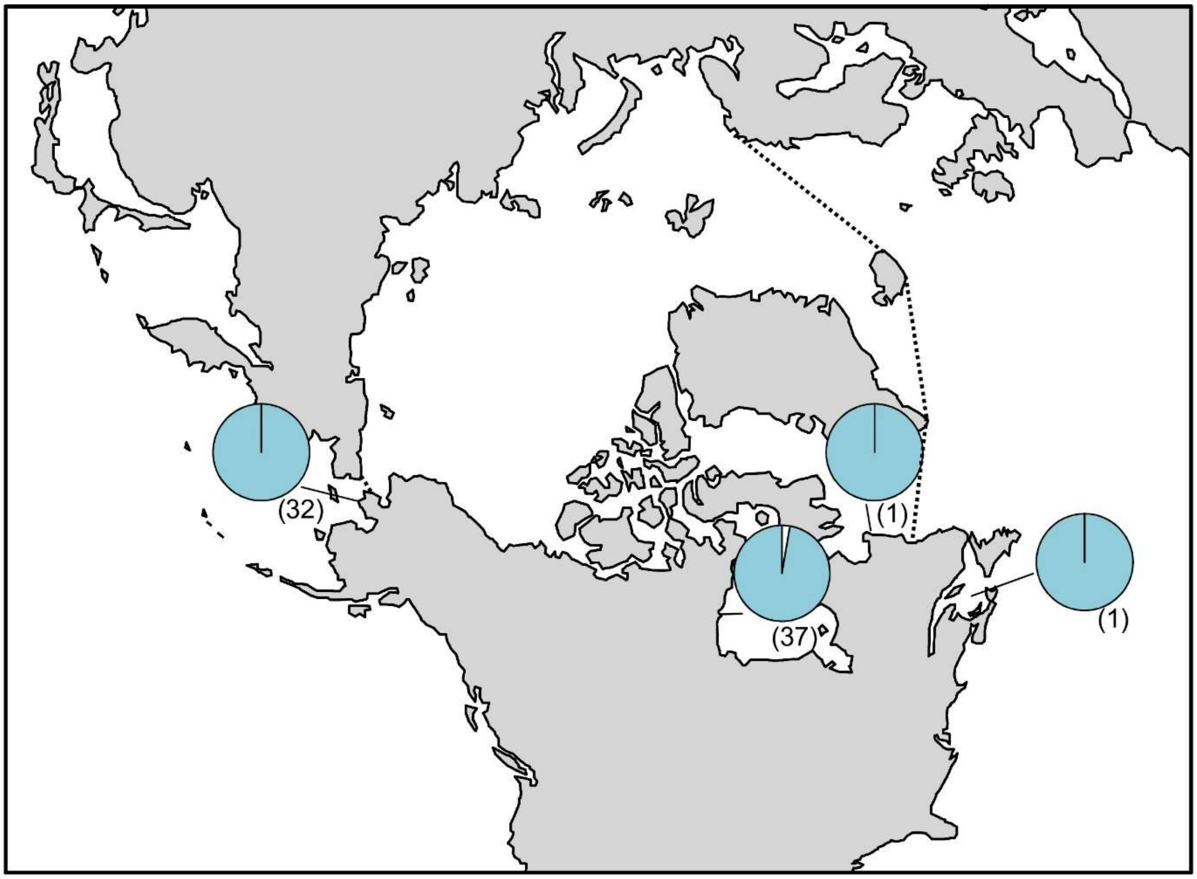

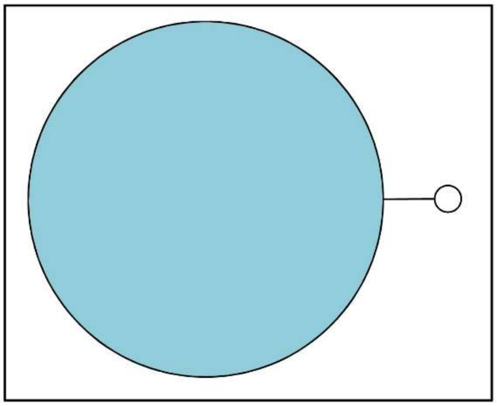

**Figure S51.** *Chordaria chordaeformis* haplotype map based on COI-5P data. In the map, numbers in parentheses refer to sample sizes from given locales. The dashed line indicates delineation of the Arctic Ocean.

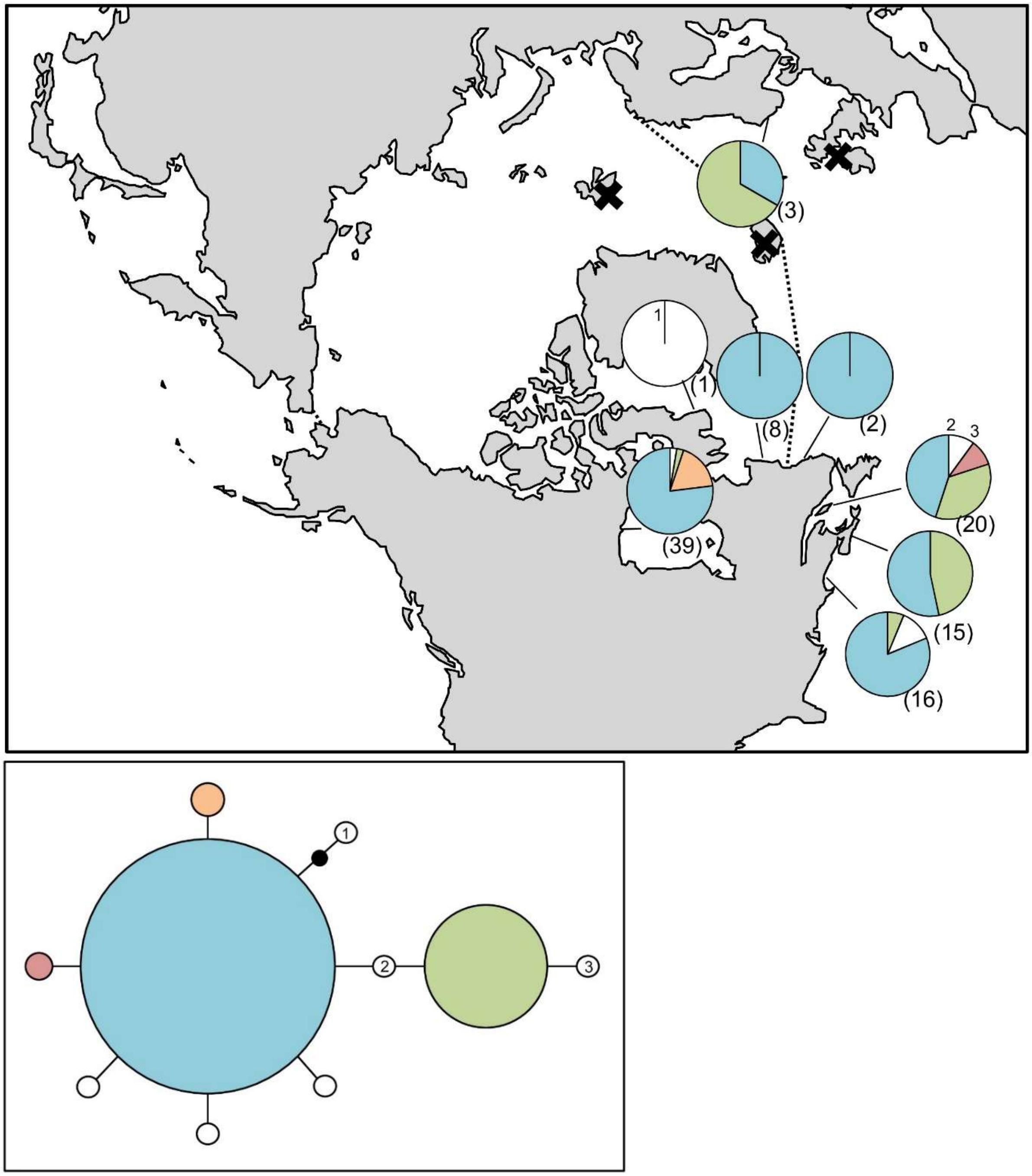

**Figure S52.** *Chordaria flagelliformis* haplotype map and network based on COI-5P data. In the map, numbers in parentheses refer to sample sizes from given locales, whereas numbers adjacent to white portions of pie charts refer to a haplotype sampled only once in the accompanying network. The dashed line indicates delineation of the Arctic Ocean, while black X's indicate the locations of genetically verified *Chordaria flagelliformis* based on *rbc*L. In the haplotype network, numbered haplotypes in white reference back to the map. Black circles indicate hypothesized (e.g. unsampled) haplotypes between clades. Circle size is proportional to the sampling frequency of a given haplotype.

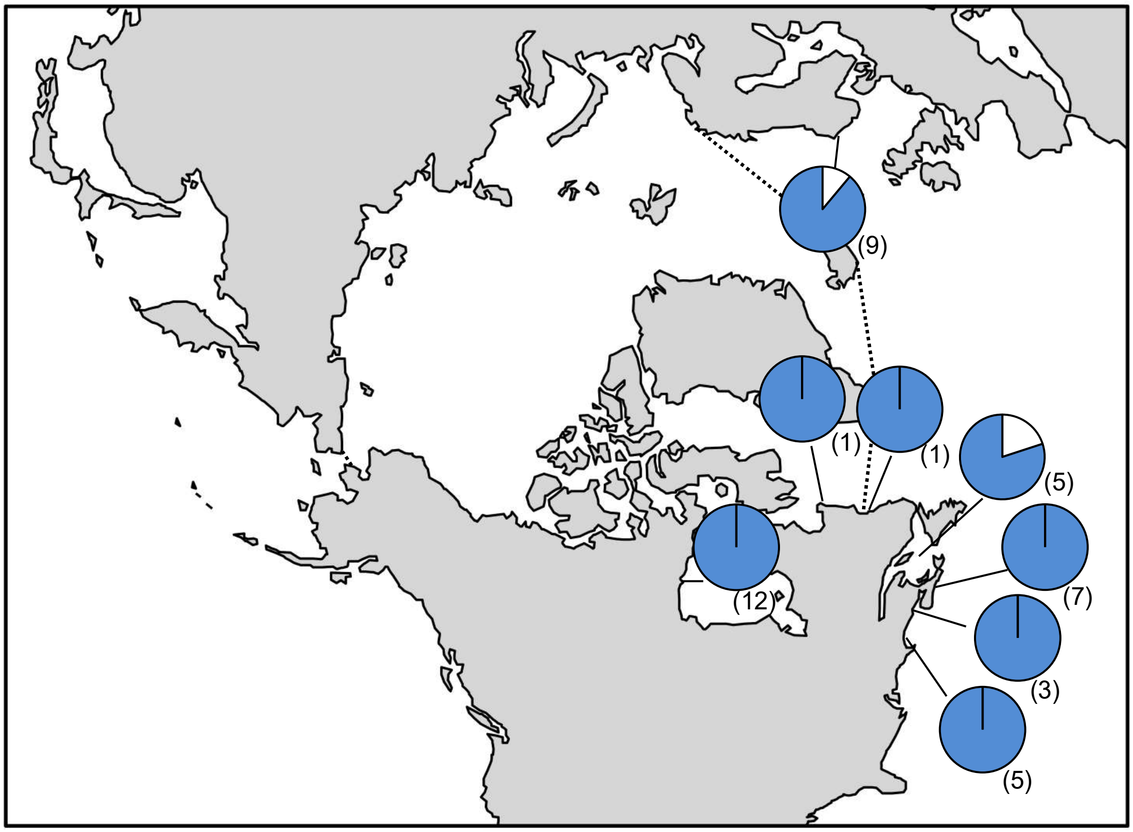

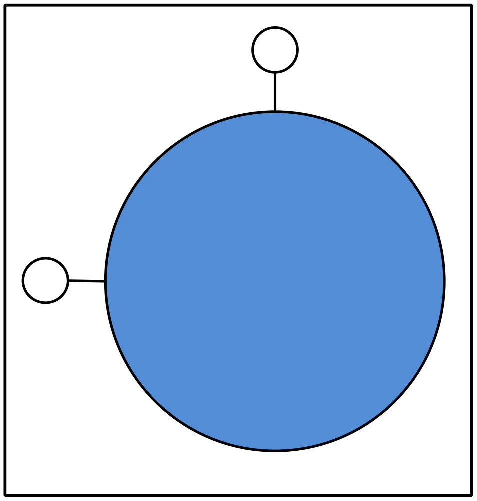

**Figure S53.** *Desmarestia* sp. 1aculeata haplotype map and network based on COI-5P data. In the map, numbers in parentheses refer to sample sizes from given locales. The dashed line indicates delineation of the Arctic Ocean. In the haplotype network, circle size is proportional to the sampling frequency of a given haplotype.

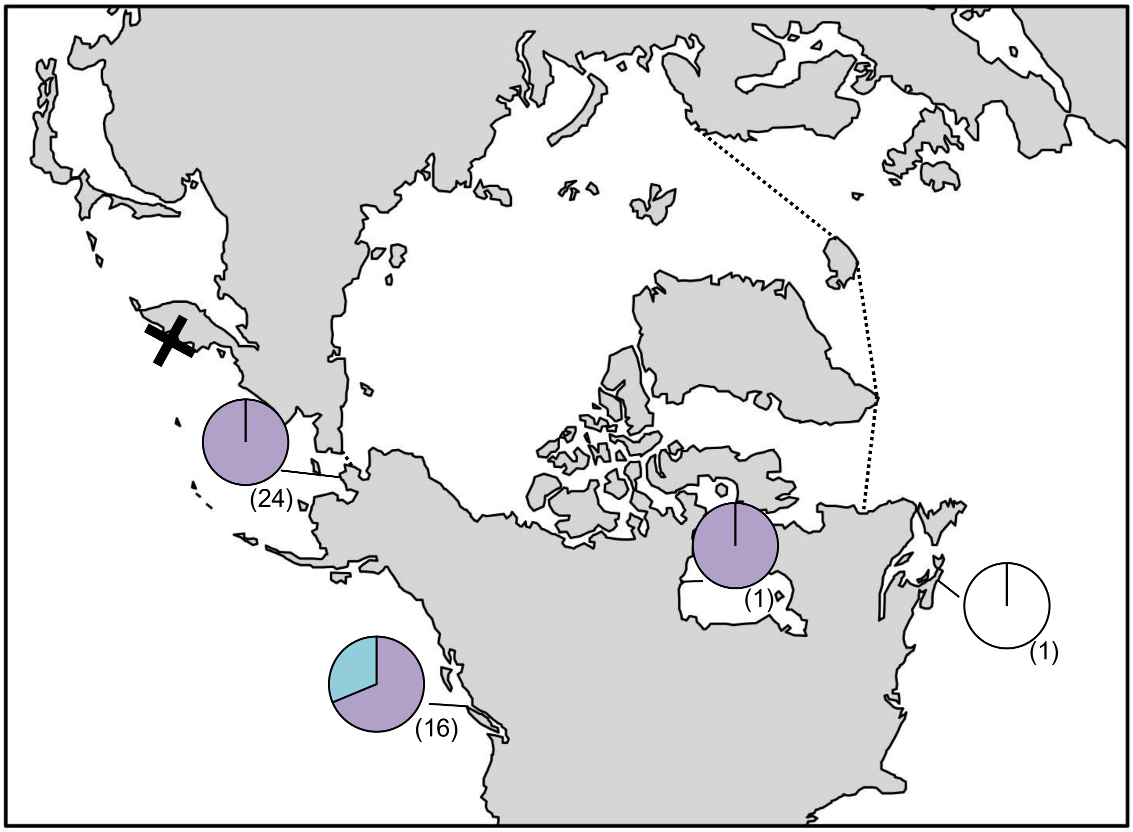

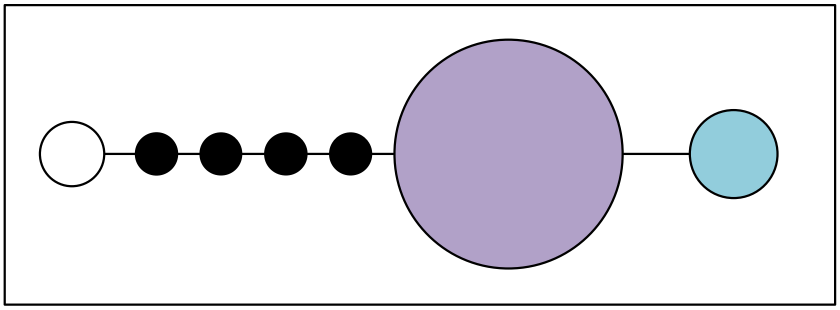

**Figure S54.** *Dictyosiphon* sp. 1GWS haplotype map and network based on COI-5P data. In the map, numbers in parentheses refer to sample sizes from given locales. The dashed line indicates delineation of the Arctic Ocean, while the black X indicates a location for genetically verified *Dictyosiphon* sp. 1GWS based on *rbc*L. In the haplotype network, black circles indicate hypothesized (e.g. unsampled) haplotypes between clades. Circle size is proportional to the sampling frequency of a given haplotype.

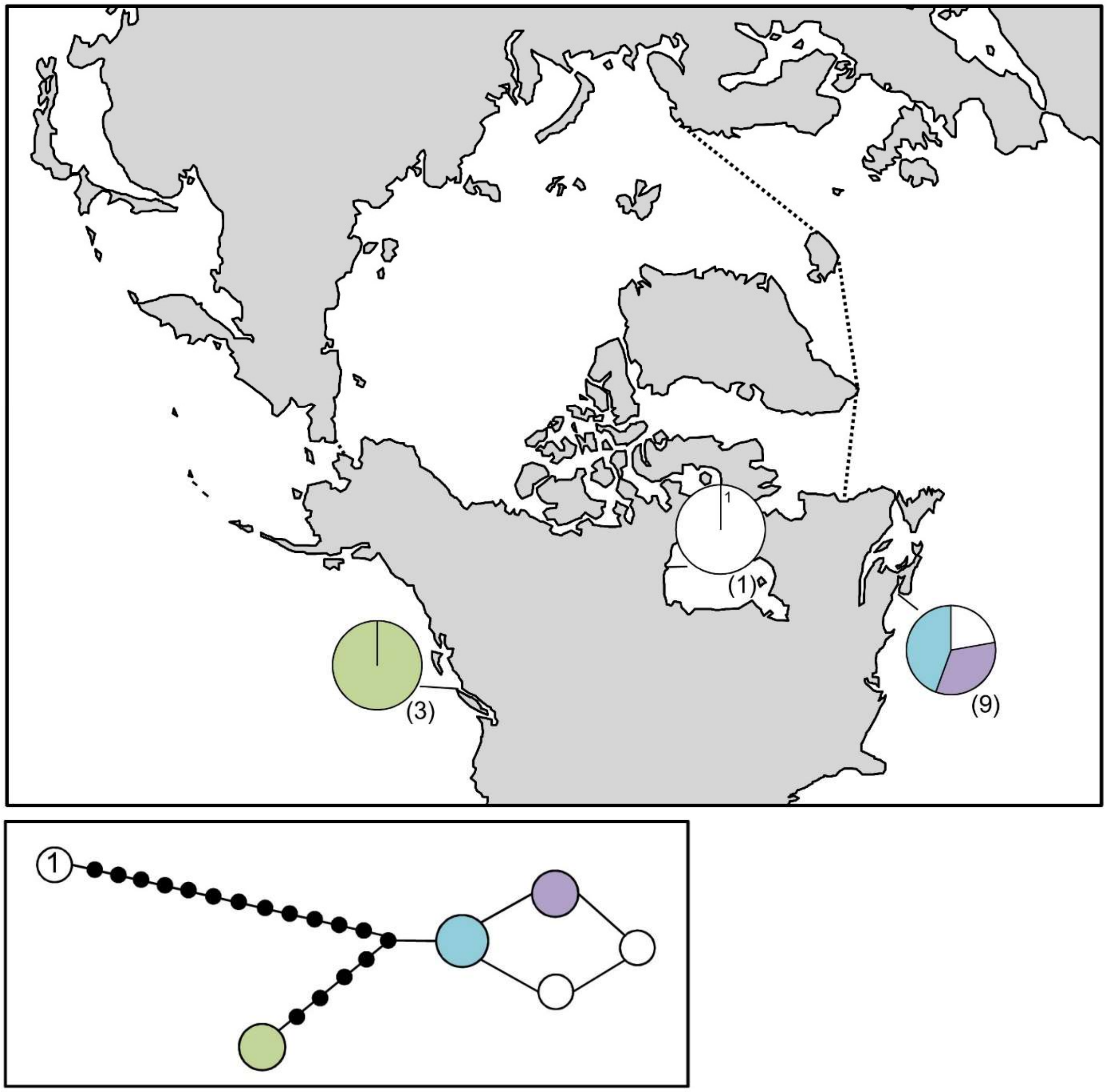

**Figure S55.** *Dictyosiphon* sp. 3GWS haplotype map and network based on COI-5P data. In the map, numbers in parentheses refer to sample sizes from given locales. The dashed line indicates delineation of the Arctic Ocean. In the haplotype network, black circles indicate hypothesized (e.g. unsampled) haplotypes between clades. Circle size is proportional to the sampling frequency of a given haplotype.

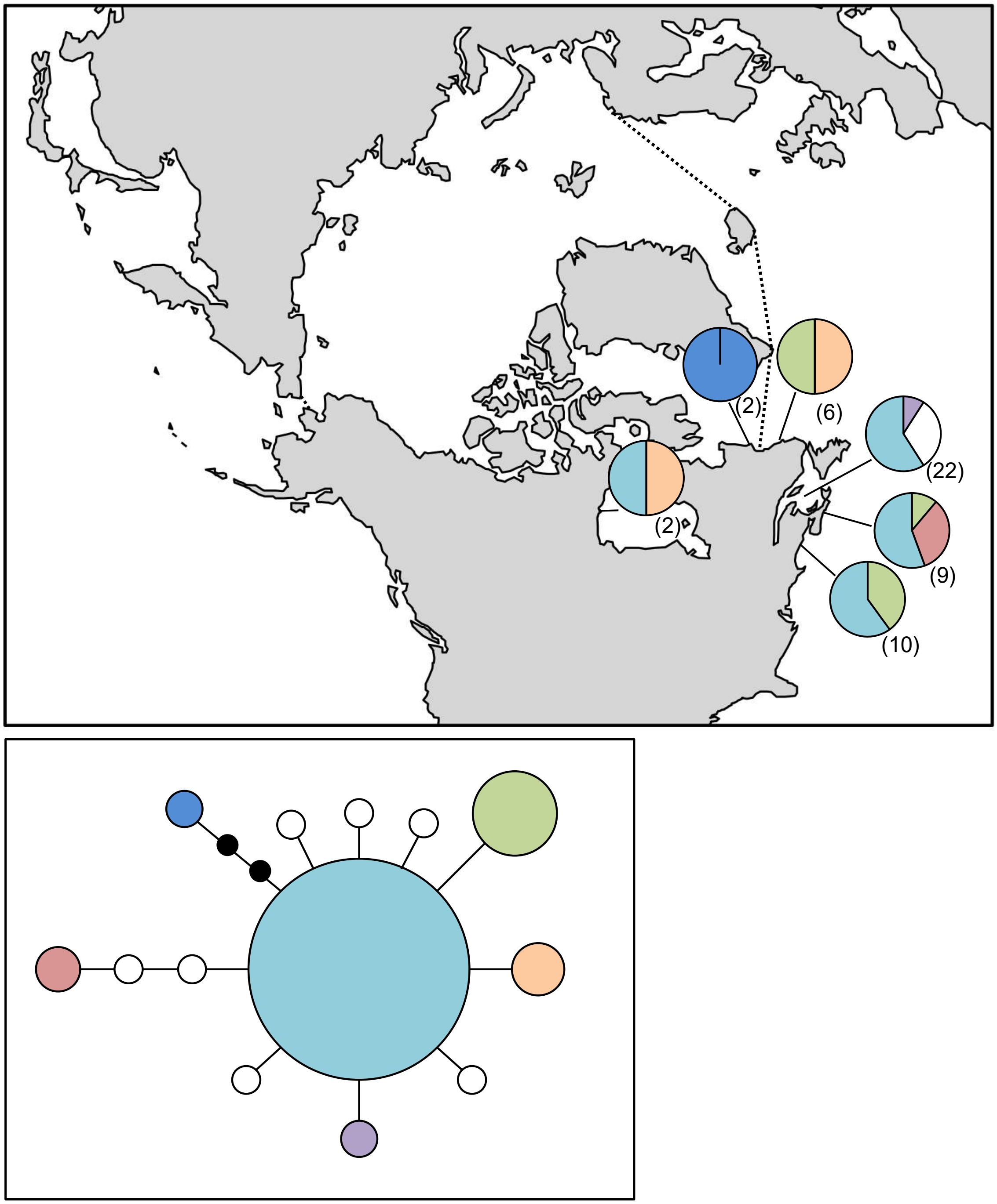

**Figure S56.** *Dictyosiphon foeniculaceus* haplotype map and network based on COI-5P data. In the map, numbers in parentheses refer to sample sizes from given locales. The dashed line indicates delineation of the Arctic Ocean. In the haplotype network, the black circles indicate hypothesized (e.g. unsampled) haplotypes between clades. Circle size is proportional to the sampling frequency of a given haplotype.

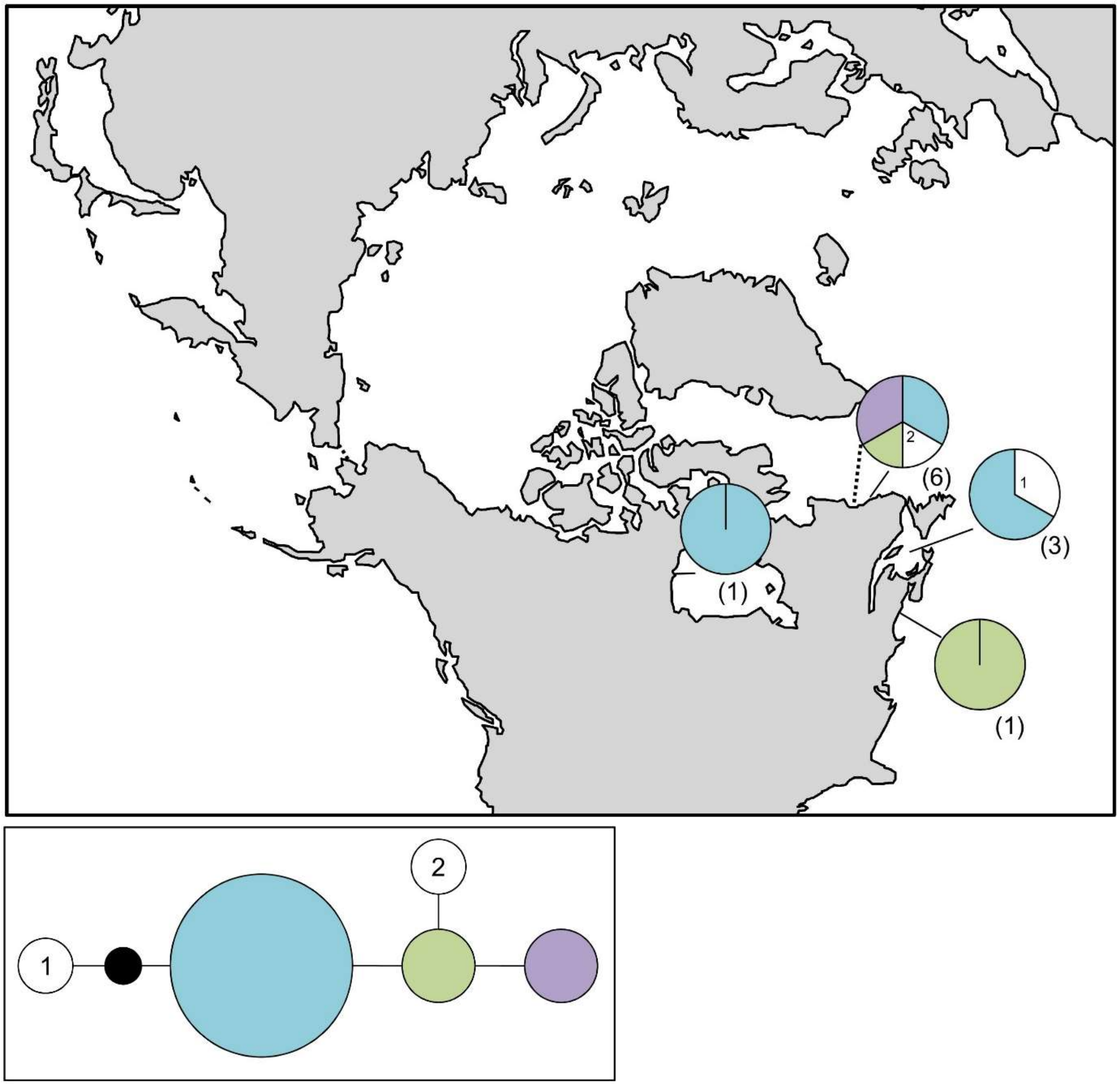

**Figure S57.** *Ectocarpus* sp. 1siliculosus haplotype map and network based on COI-5P data. In the map, numbers in parentheses refer to sample sizes from given locales. The dashed line indicates delineation of the Arctic Ocean. In the haplotype network, the black circles indicate hypothesized (e.g. unsampled) haplotypes between clades. Circle size is proportional to the sampling frequency of a given haplotype.

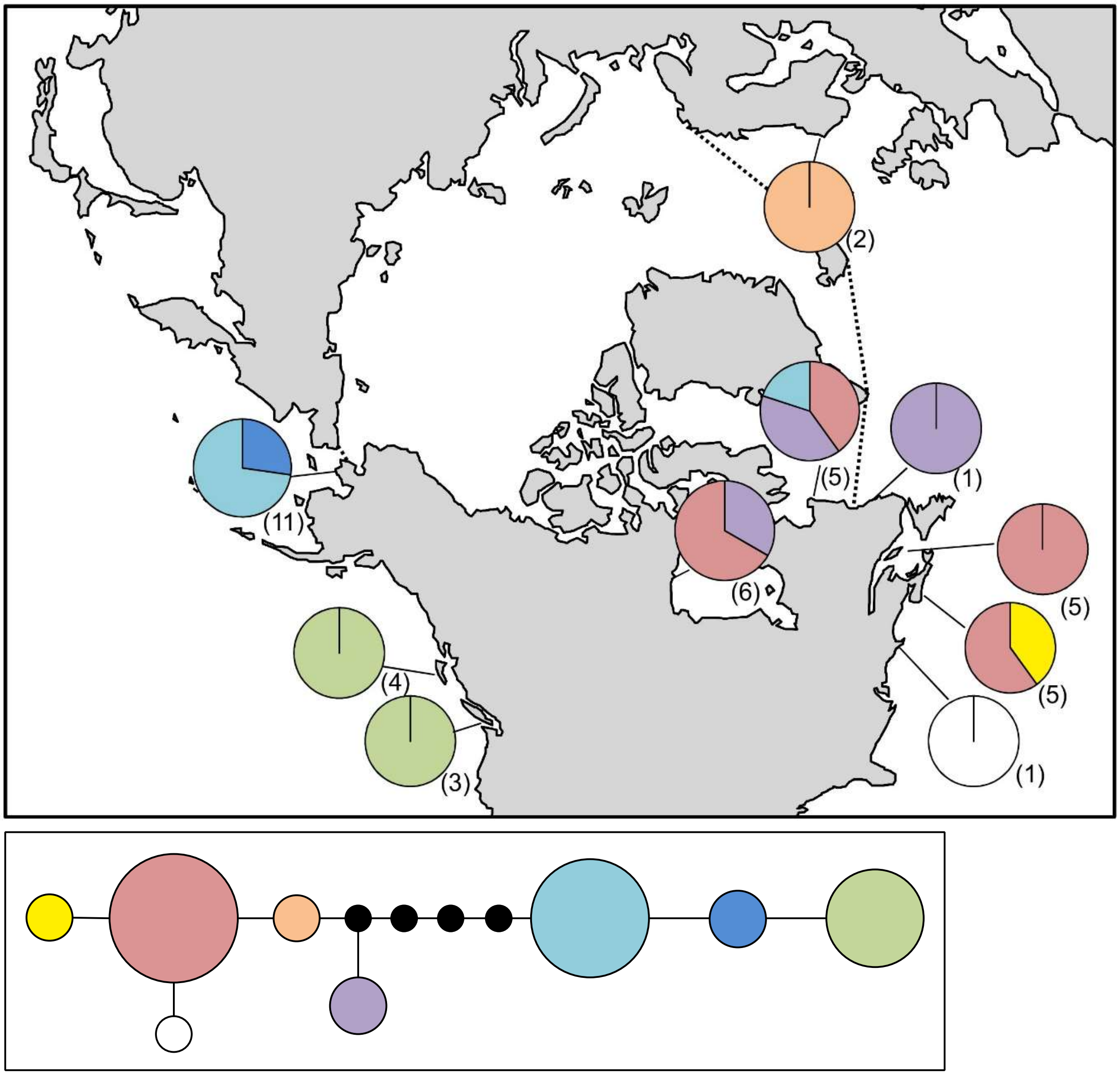

**Figure S58.** *Eudesme borealis* haplotype map and network based on COI-5P data. In the map, numbers in parentheses refer to sample sizes from given locales. The dashed line indicates delineation of the Arctic Ocean. In the haplotype network, the black circles indicate hypothesized (e.g. unsampled) haplotypes between clades. Circle size is proportional to the sampling frequency of a given haplotype.

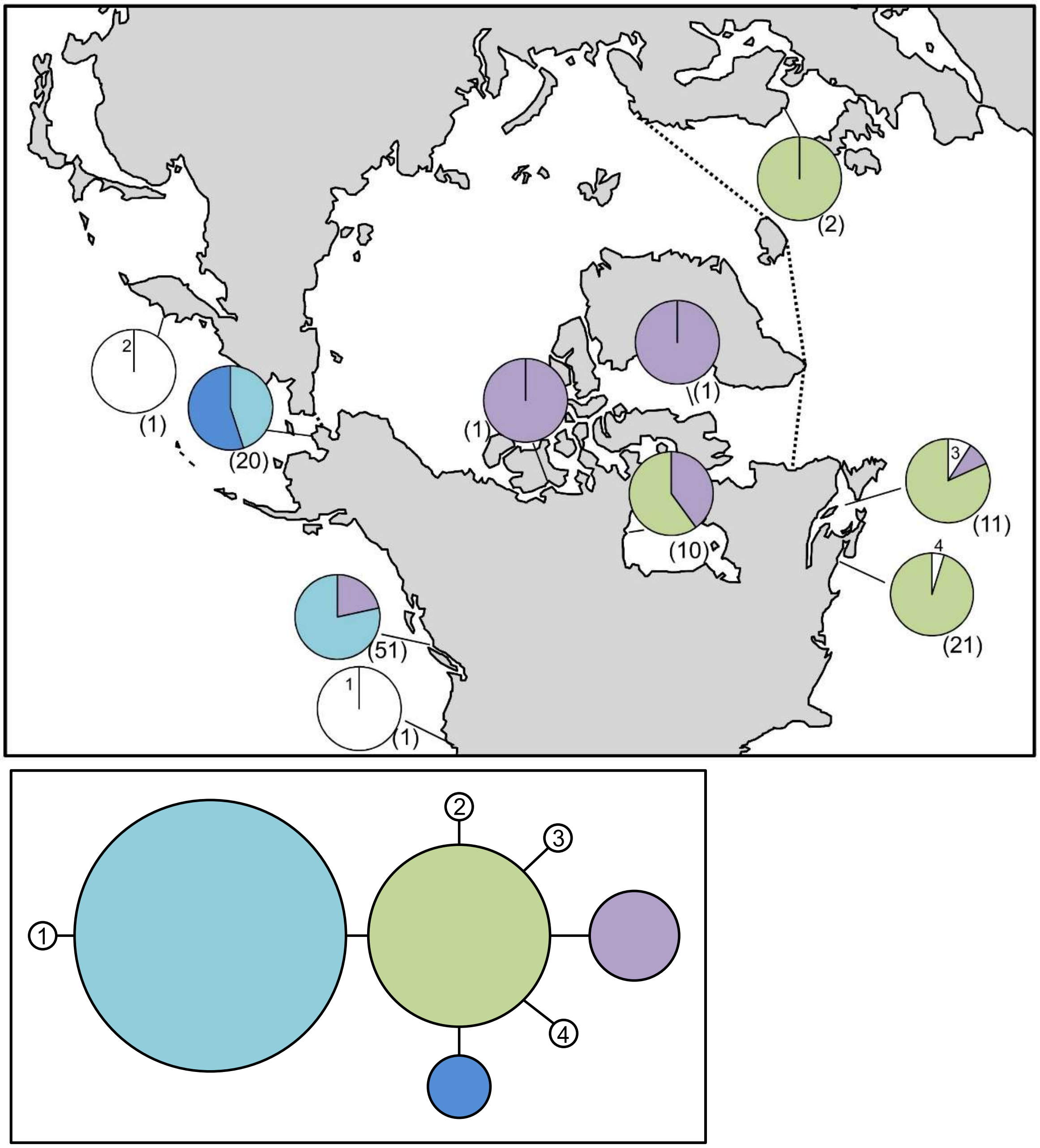

**Figure S59.** *Fucus distichus* haplotype map and network based on COI-5P data. In the map, numbers in parentheses refer to sample sizes from given locales, whereas numbers adjacent to white portions of pie charts refer to a haplotype sampled only once in the accompanying network. The dashed line indicates delineation of the Arctic Ocean. In the haplotype network, numbered haplotypes in white reference back to the map. Circle size is proportional to the sampling frequency of a given haplotype.

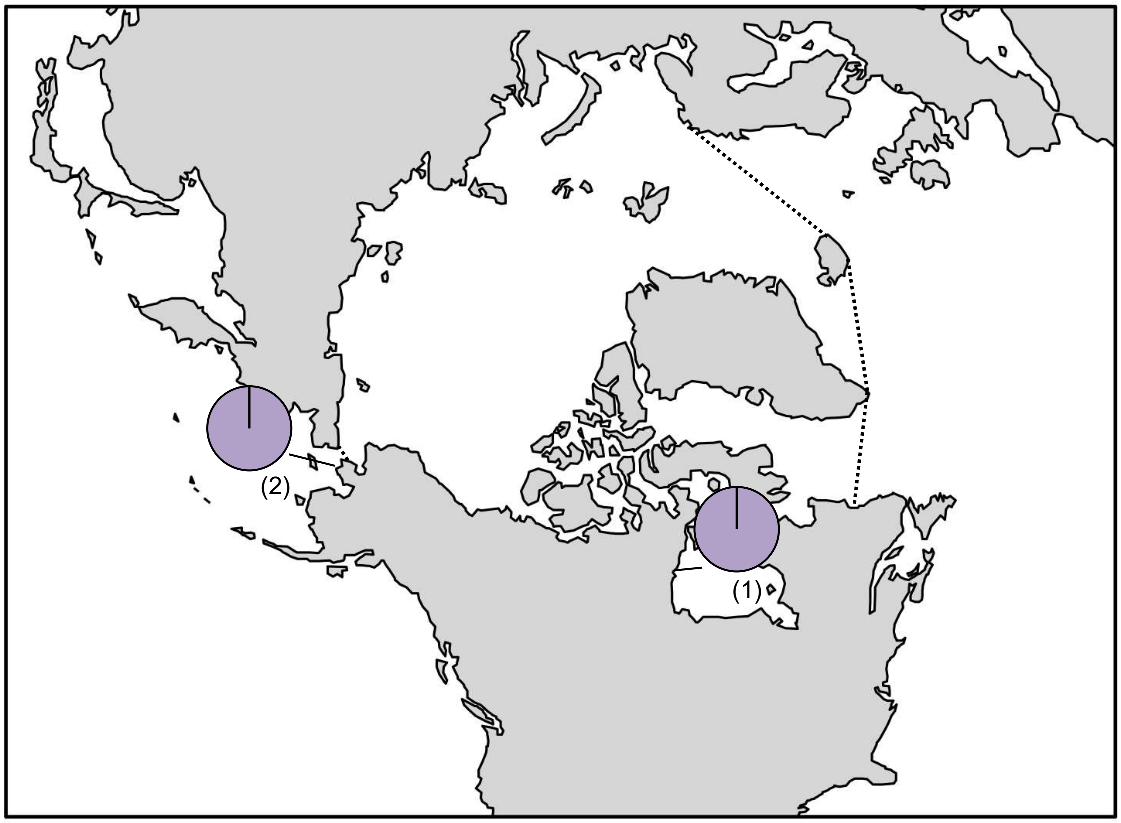

**Figure S60.** *Halosiphon sp. 2tomentosus* haplotype map based on COI-5P data. In the map, numbers in parentheses refer to sample sizes from given locales. The dashed line indicates delineation of the Arctic Ocean.

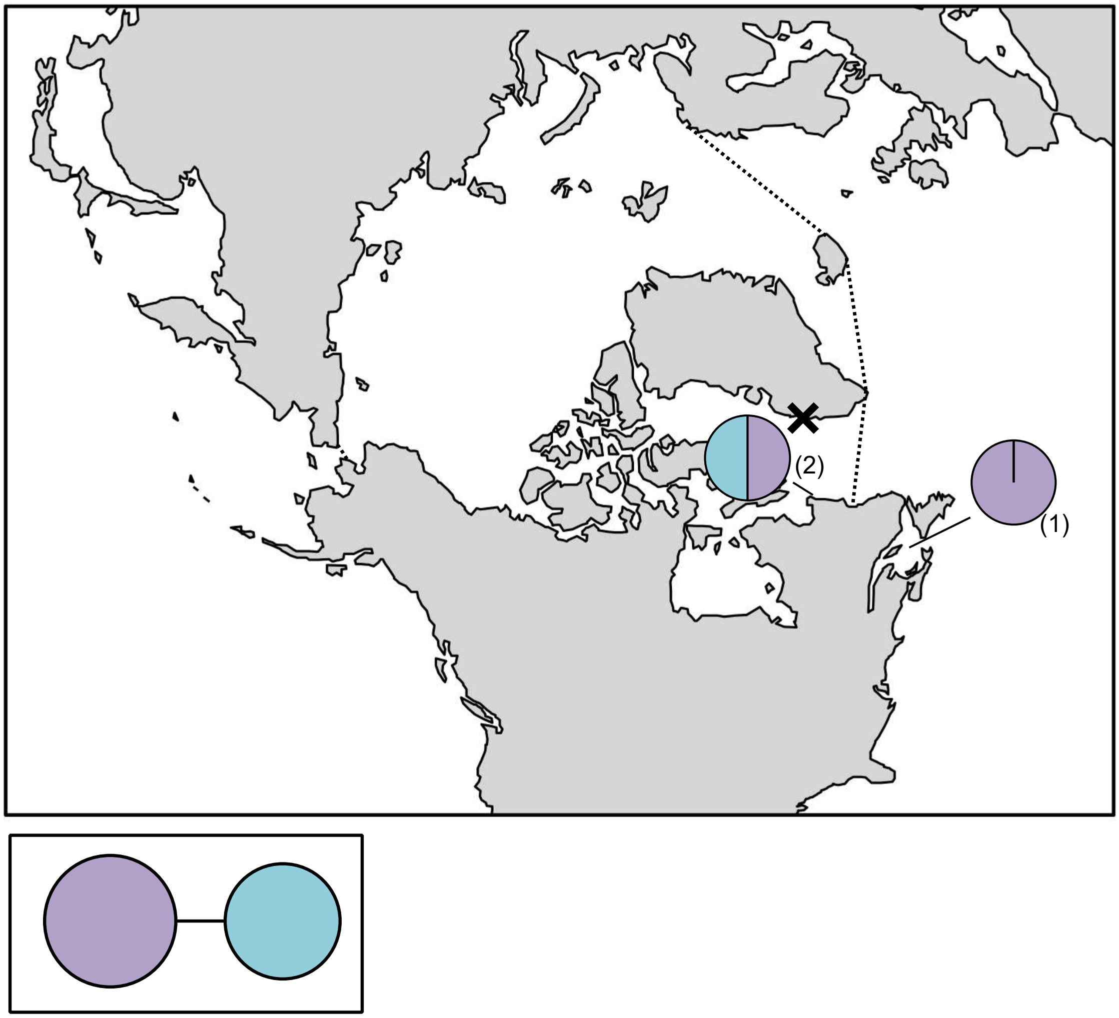

**Figure S61.** *Halothrix lumbricalis* haplotype map based on COI-5P data. In the map, numbers in parentheses refer to sample sizes from given locales. The dashed line indicates delineation of the Arctic Ocean, while the black X indicates a location for genetically verified *Halothrix lumbricalis* based on *rbc*L. In the haplotype network, circle size is proportional to the sampling frequency of a given haplotype.

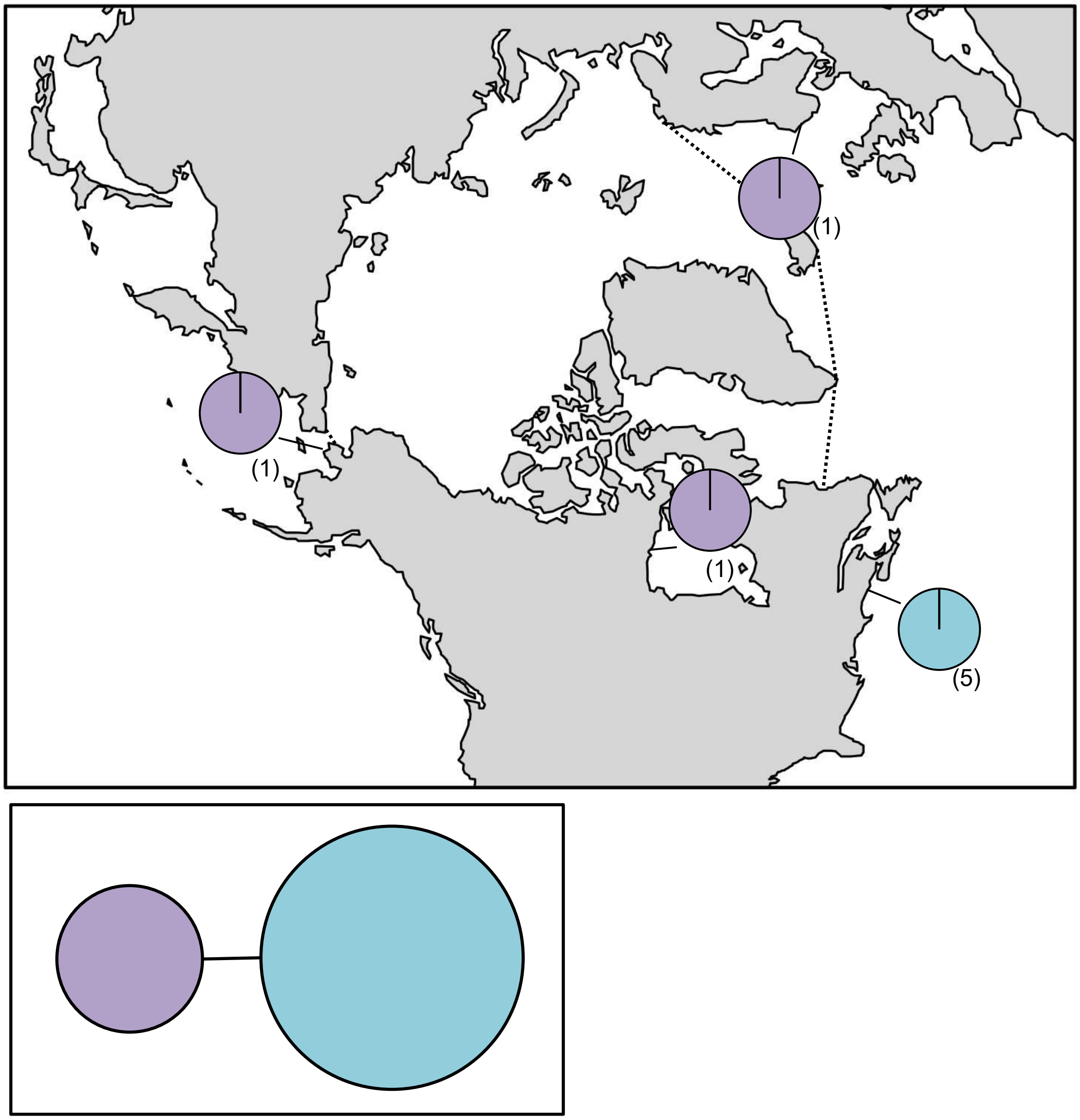

**Figure S62.** *Haplospora globosa* haplotype map and network based on COI-5P data. In the map, numbers in parentheses refer to sample sizes from given locales. The dashed line indicates delineation of the Arctic Ocean. In the haplotype network, circle size is proportional to the sampling frequency of a given haplotype.

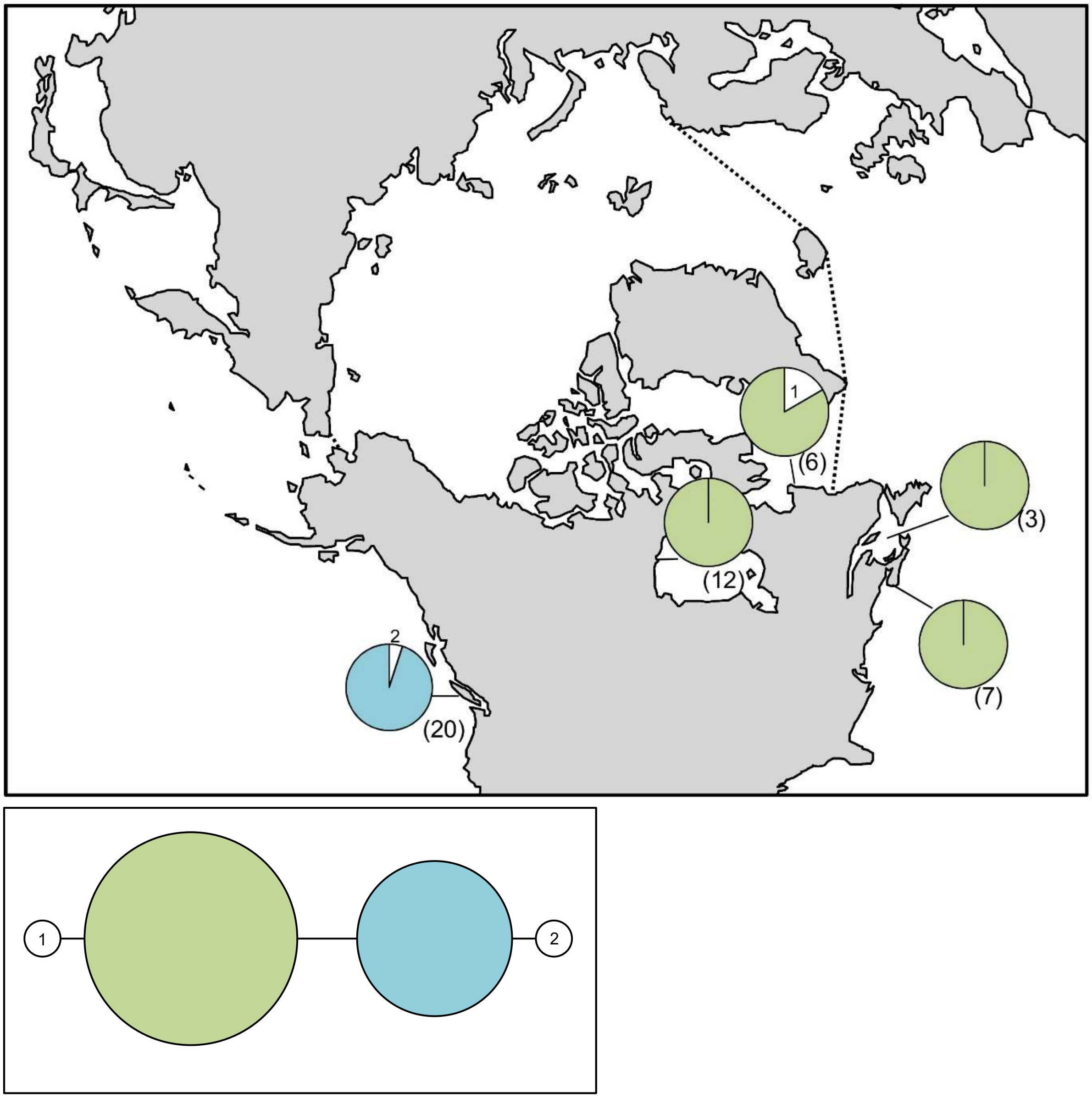

**Figure S63.** *Hedophyllum nigripes* haplotype map and network based on COI-5P data. In the map, numbers in parentheses refer to sample sizes from given locales, whereas numbers adjacent to white portions of pie charts refer to a haplotype sampled only once in the accompanying network. The dashed line indicates delineation of the Arctic Ocean. In the haplotype network, numbered haplotypes in white reference back to the map. Circle size is proportional to the sampling frequency of a given haplotype.

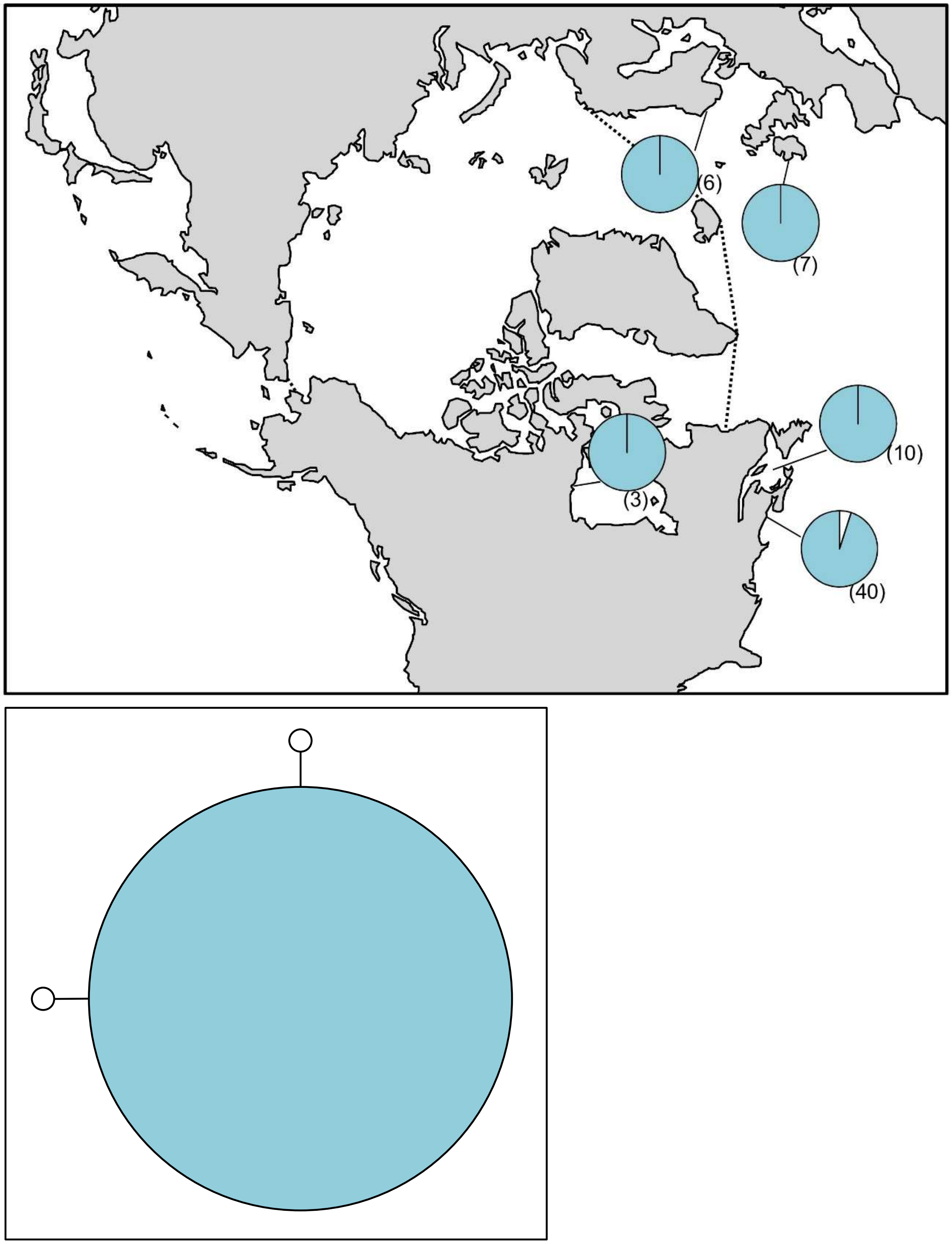

**Figure S64.** *Laminaria digitata* haplotype map and network based on COI-5P data. In the map, numbers in parentheses refer to sample sizes from given locales. The dashed line indicates delineation of the Arctic Ocean. In the haplotype network, circle size is proportional to the sampling frequency of a given haplotype.

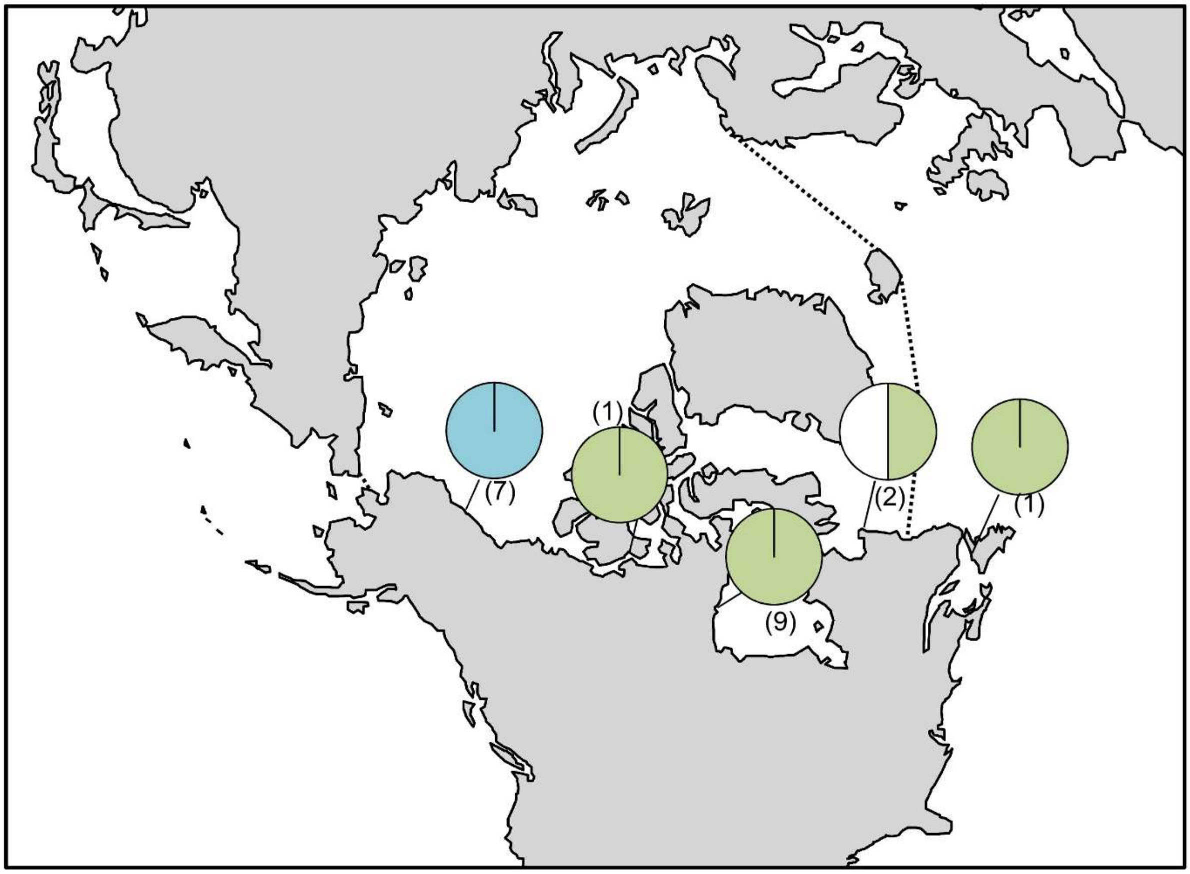

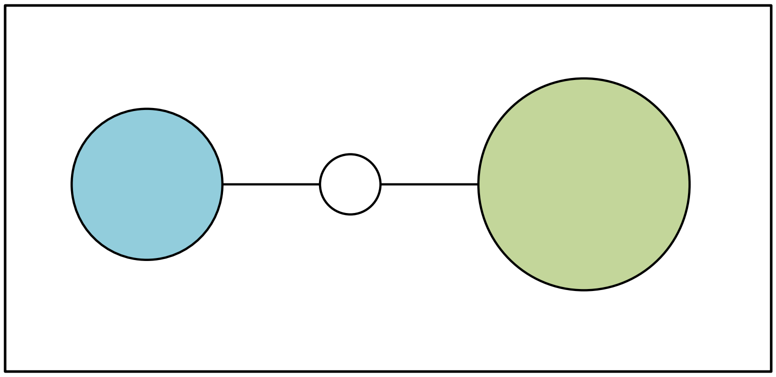

**Figure S65.** *Laminaria solidungula* haplotype map and network based on COI-5P data. In the map, numbers in parentheses refer to sample sizes from given locales. The dashed line indicates delineation of the Arctic Ocean. In the haplotype network, circle size is proportional to the sampling frequency of a given haplotype.

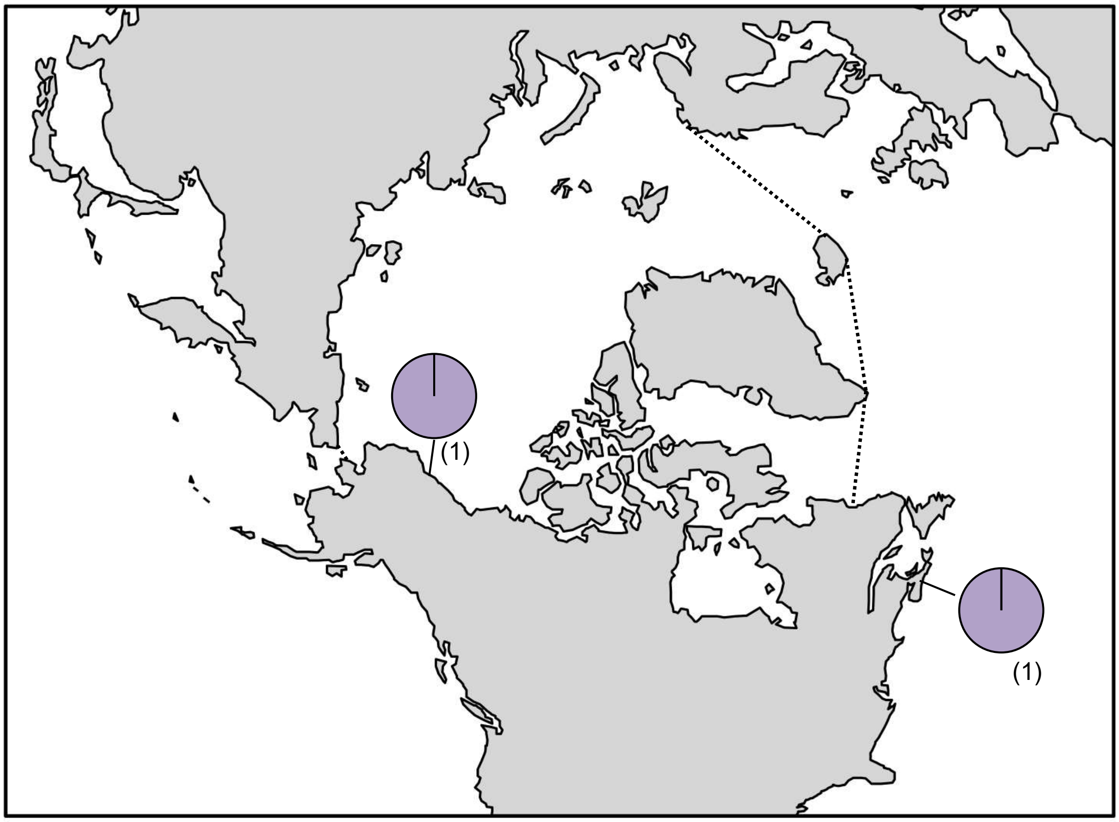

**Figure S66.** *Lithoderma* sp. 2GWS haplotype map based on *rbc*L-3P data. In the map, numbers in parentheses refer to sample sizes from given locales. The dashed line indicates delineation of the Arctic Ocean.

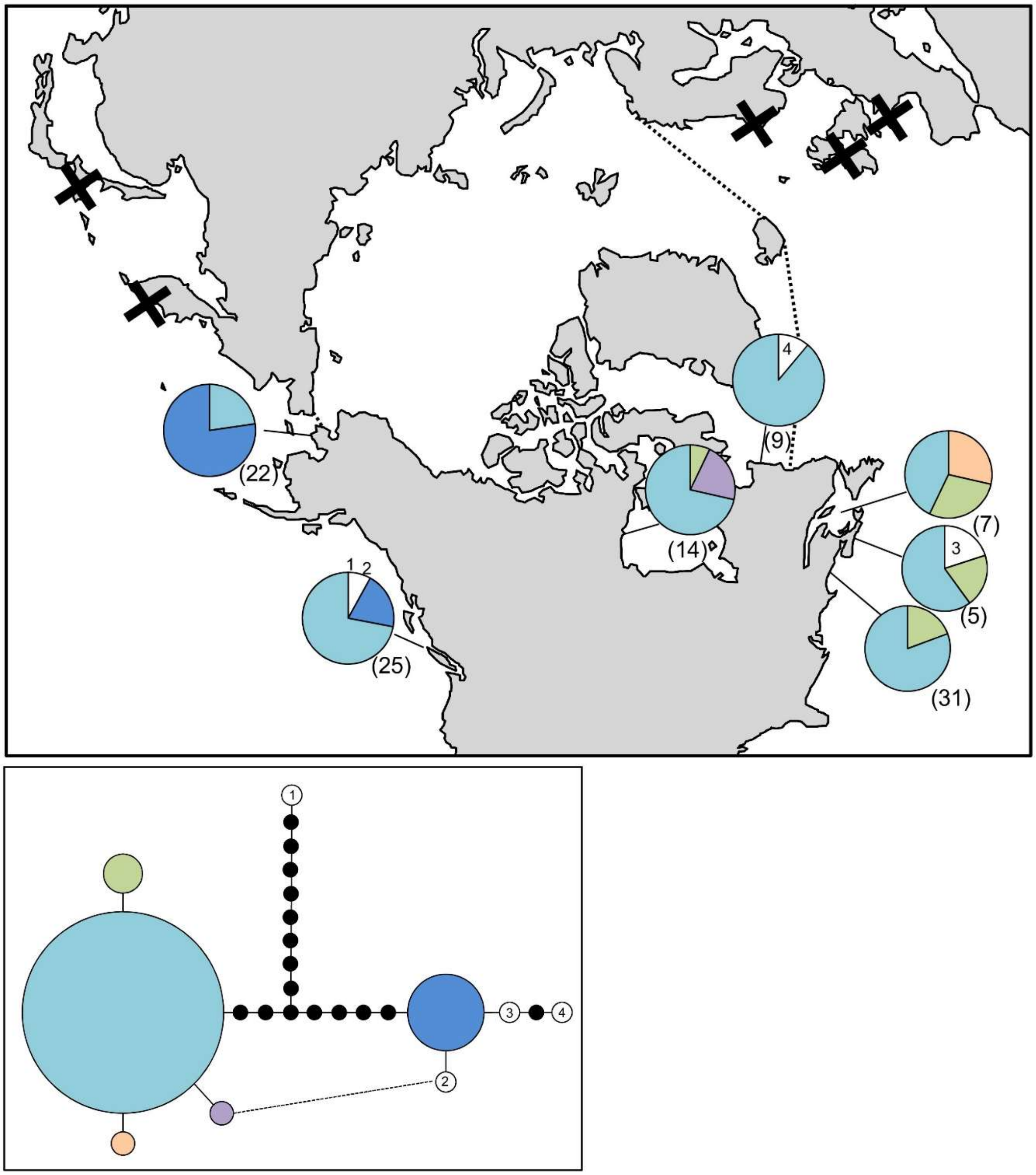

**Figure S67.** *Petalonia fascia* haplotype map and network based on COI-5P data. In the map, numbers in parentheses refer to sample sizes from given locales, whereas numbers adjacent to white portions of pie charts refer to a haplotype sampled only once in the accompanying network. The dashed line indicates delineation of the Arctic Ocean, while black X's indicate locations for genetically verified *Petalonia fascia* based on ITS, PSA, and *rbc*L. In the haplotype network, numbered haplotypes in white reference back to the map. Black circles indicate hypothesized (e.g. unsampled) haplotypes between clades. Circle size is proportional to the sampling frequency of a given haplotype. The dashed line indicates a substitution at the same site in two corresponding haplotypes.

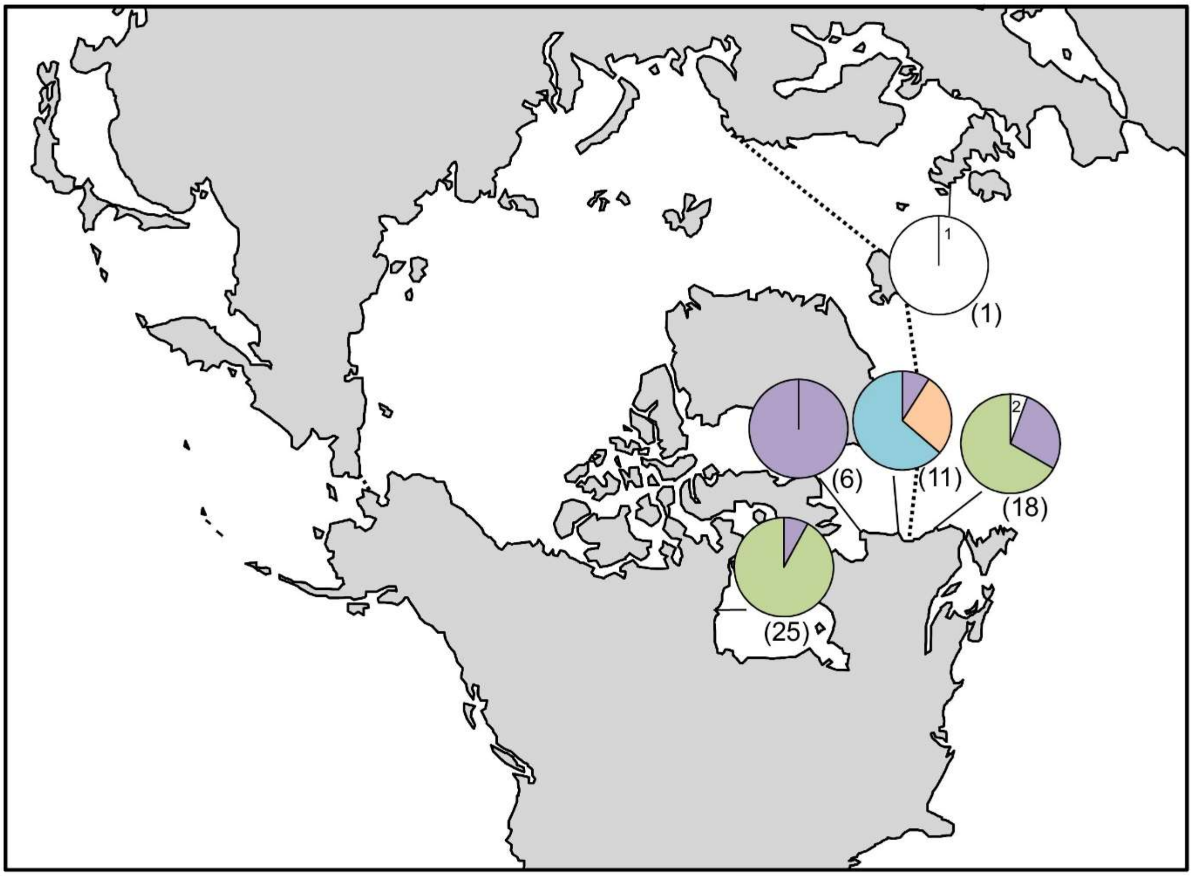

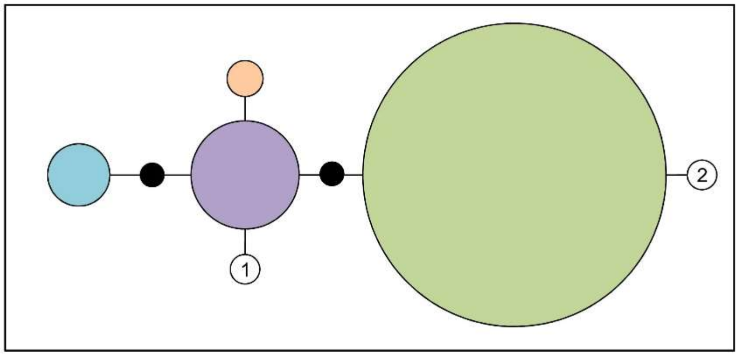

**Figure S68.** *Petalonia filiformis* haplotype map and network based on COI-5P data. In the map, numbers in parentheses refer to sample sizes from given locales. The dashed line indicates delineation of the Arctic Ocean. In the haplotype network, black circles indicate hypothesized (e.g. unsampled) haplotypes between clades. Circle size is proportional to the sampling frequency of a given haplotype.

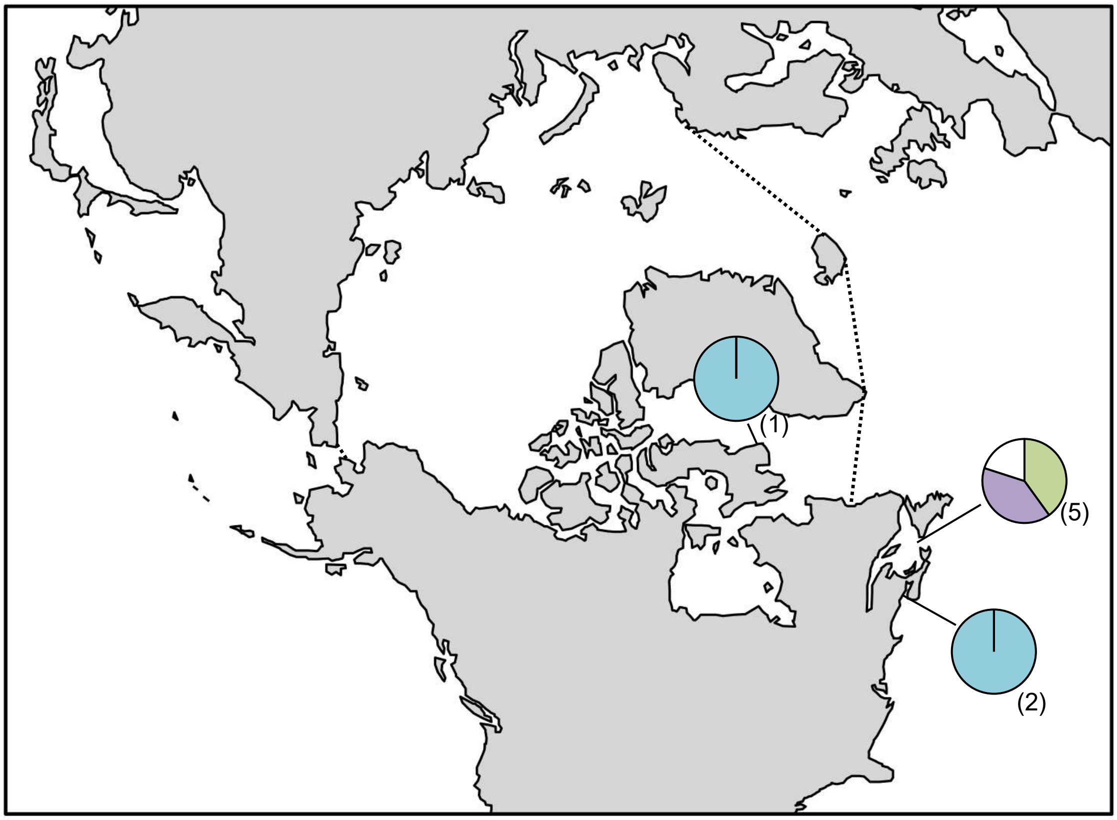

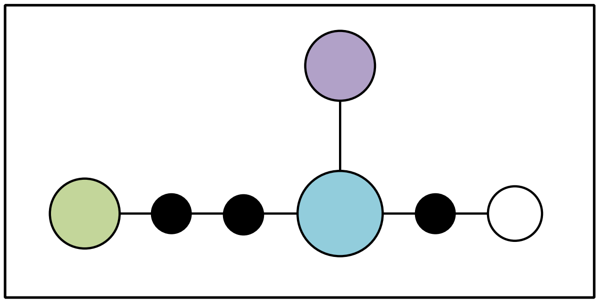

**Figure S69.** *Planosiphon complanatus* haplotype map and network based on COI-5P data. In the map, numbers in parentheses refer to sample sizes from given locales. The dashed line indicates delineation of the Arctic Ocean. In the haplotype network, black circles indicate hypothesized (e.g. unsampled) haplotypes between clades. Circle size is proportional to the sampling frequency of a given haplotype.

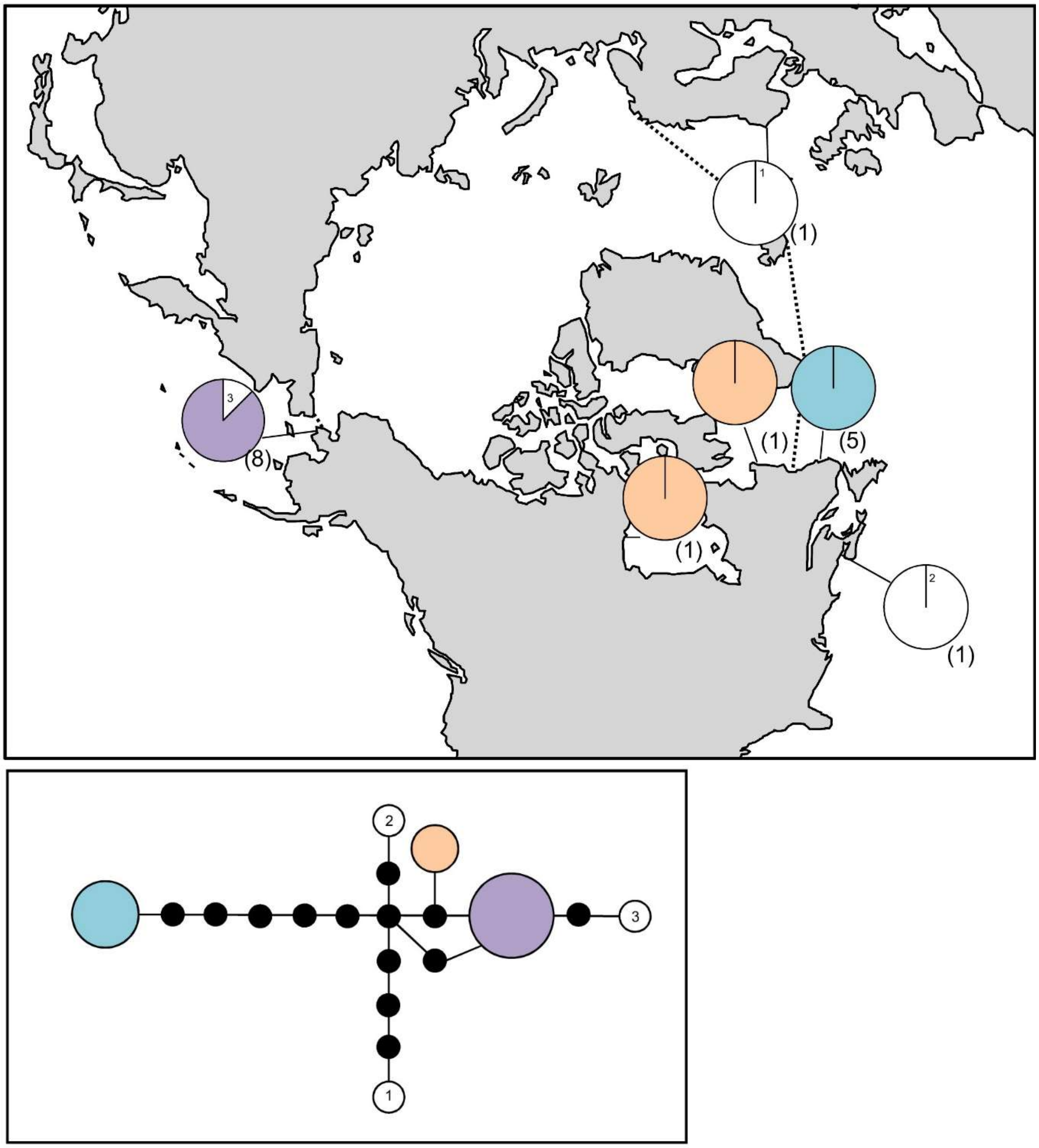

**Figure S70.** *Planosiphon zosterifolius* haplotype map and network based on COI-5P data. In the map, numbers in parentheses refer to sample sizes from given locales. The dashed line indicates delineation of the Arctic Ocean. In the haplotype network, black circles indicate hypothesized (e.g. unsampled) haplotypes between clades. Circle size is proportional to the sampling frequency of a given haplotype.

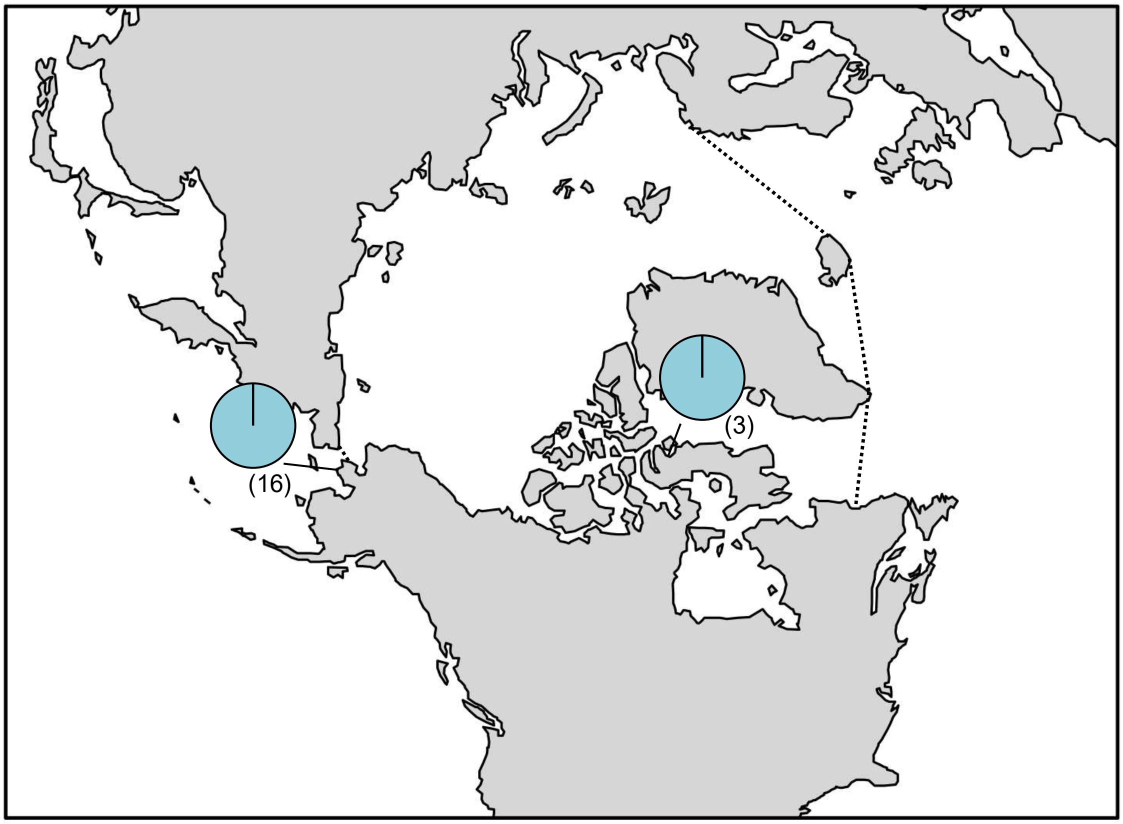

**Figure S71.** *Platysiphon glacialis* haplotype map based on COI-5P data. In the map, numbers in parentheses refer to sample sizes from given locales. The dashed line indicates delineation of the Arctic Ocean

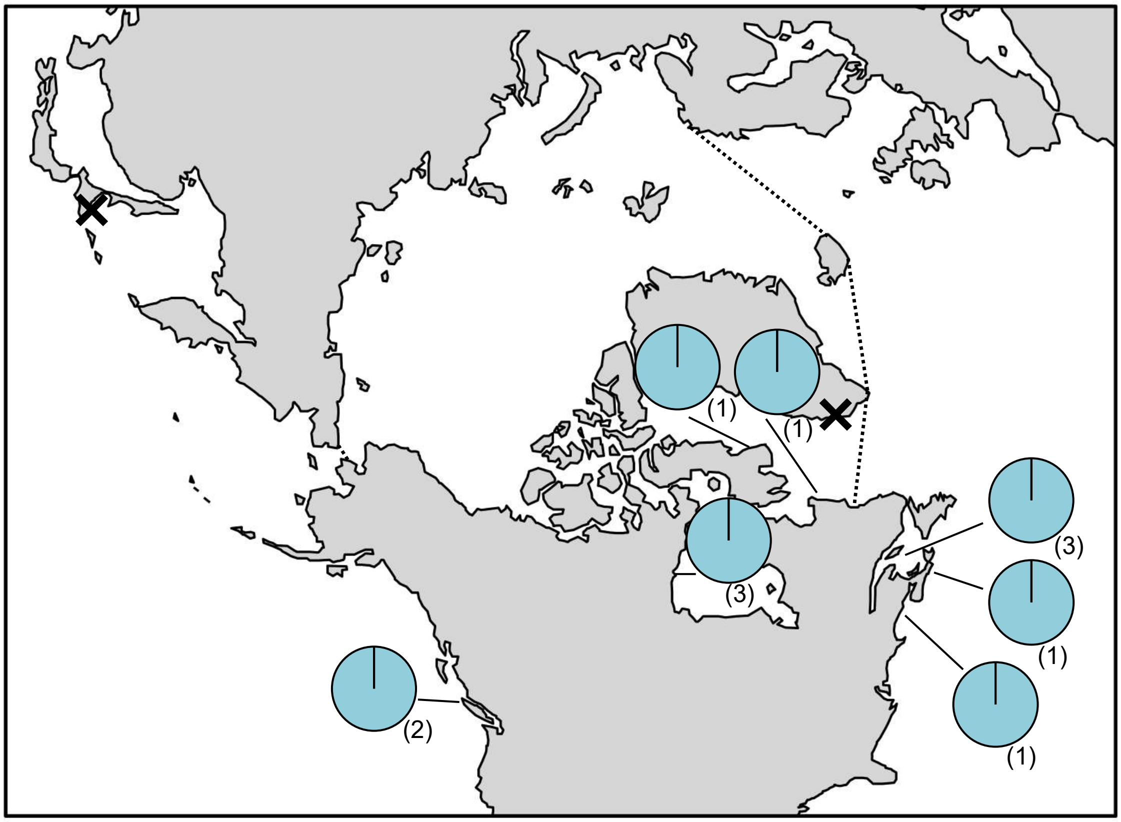

**Figure S72.** *Punctaria* sp. 2GWS haplotype map based on COI-5P data. In the map, numbers in parentheses refer to sample sizes from given locales. The dashed line indicates delineation of the Arctic Ocean. The black X's indicate locations for genetically verified *Punctaria* sp. 2GWS based on *rbc*L.

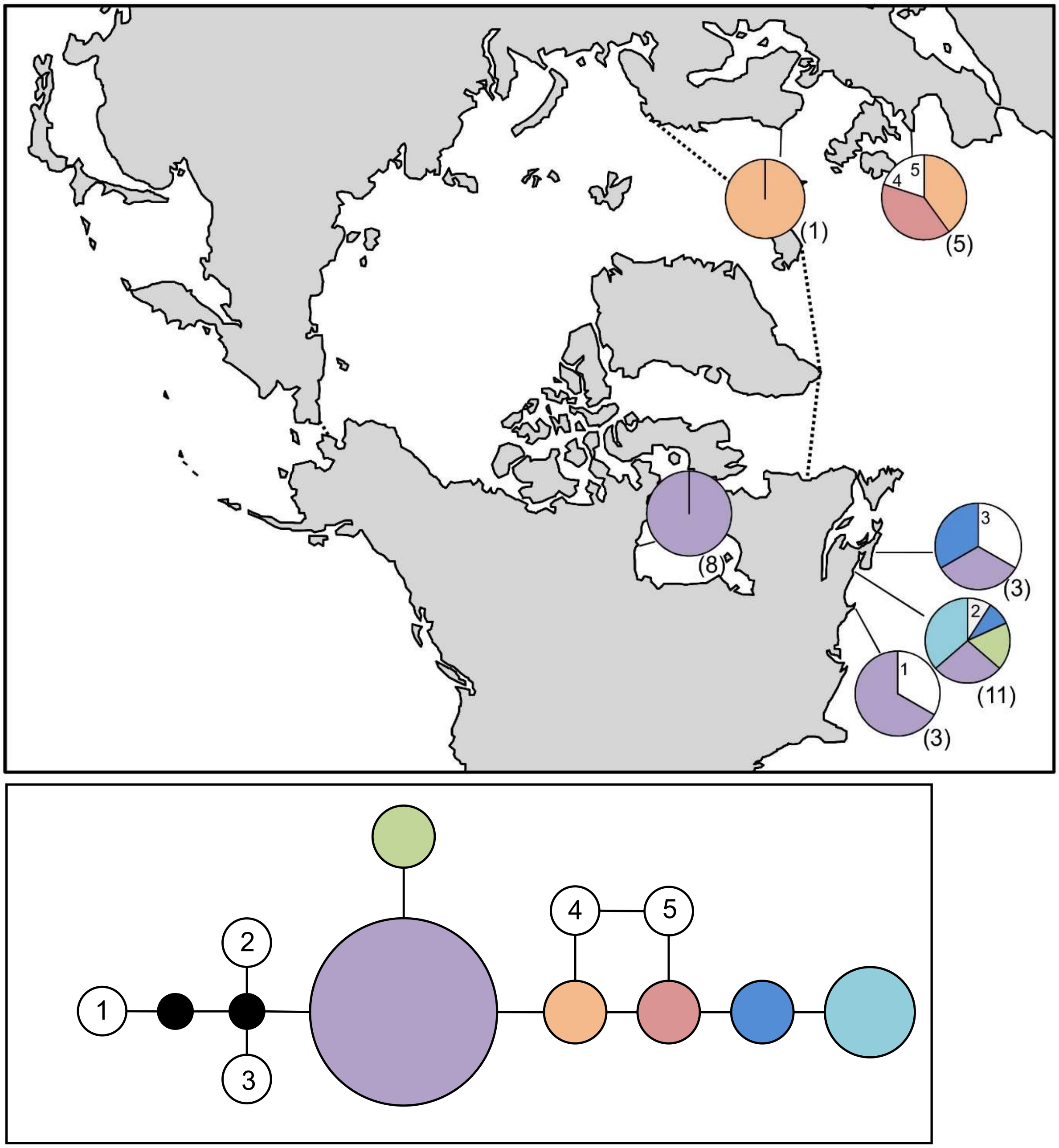

**Figure S73.** *Pylaiella littoralis* haplotype map and network based on COI-5P data. In the map, numbers in parentheses refer to sample sizes from given locales, whereas numbers adjacent to white portions of pie charts refer to a haplotype sampled only once in the accompanying network. The dashed line indicates delineation of the Arctic Ocean. In the haplotype network, numbered haplotypes in white reference back to the map. Black circles indicate hypothesized (e.g. unsampled) haplotypes between clades. Circle size is proportional to the sampling frequency of a given haplotype.

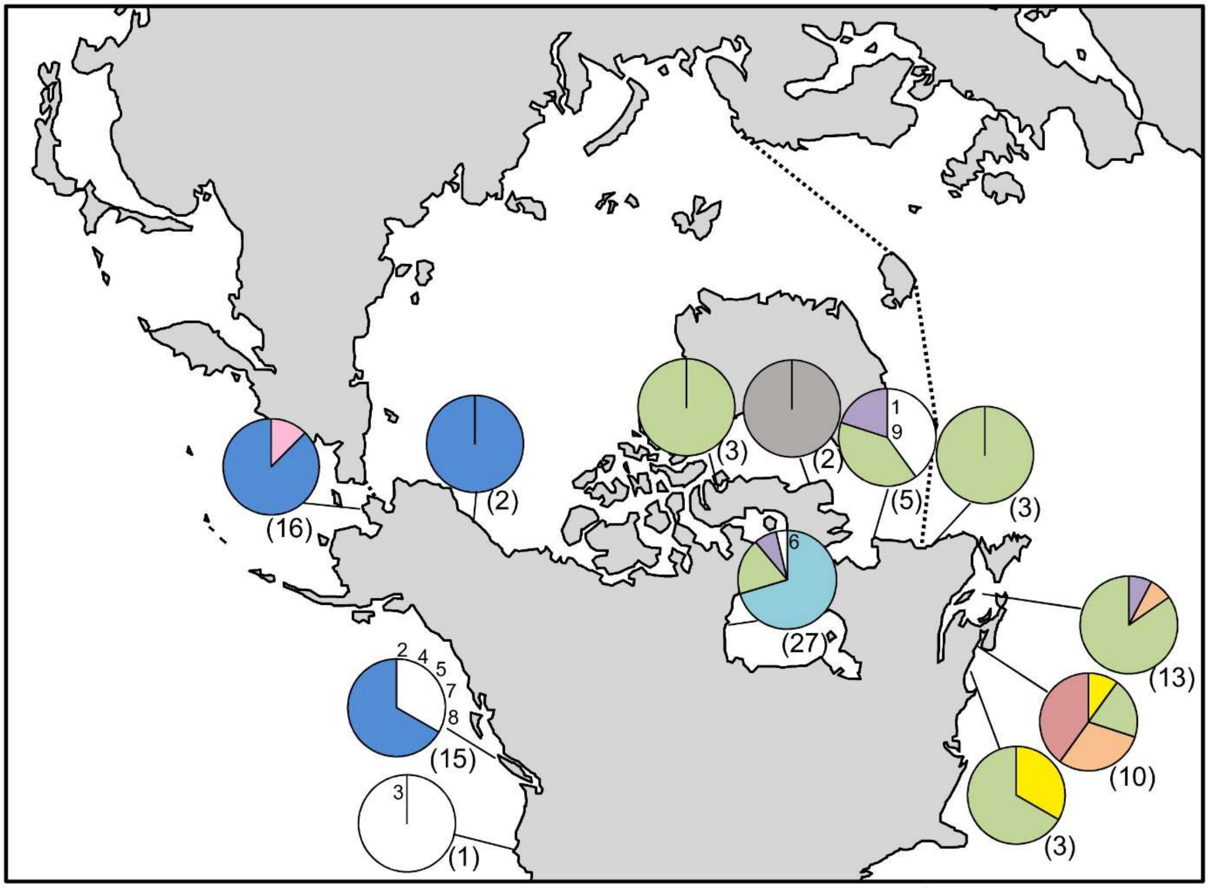

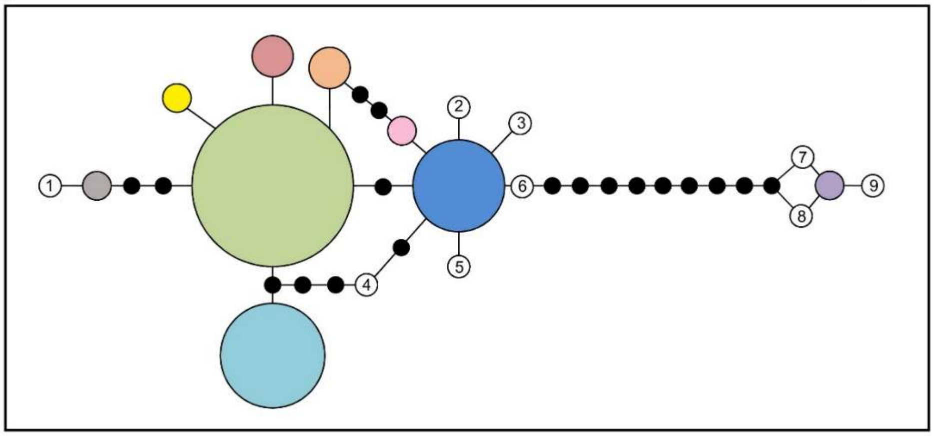

**Figure S74.** *Pylaiella washingtoniensis* haplotype map and network based on COI-5P data. In the map, numbers in parentheses refer to sample sizes from given locales, whereas numbers adjacent to white portions of pie charts refer to a haplotype sampled only once in the accompanying network. The dashed line indicates delineation of the Arctic Ocean. In the haplotype network, numbered haplotypes in white reference back to the map. Black circles indicate hypothesized (e.g. unsampled) haplotypes between clades. Circle size is proportional to the sampling frequency of a given haplotype.

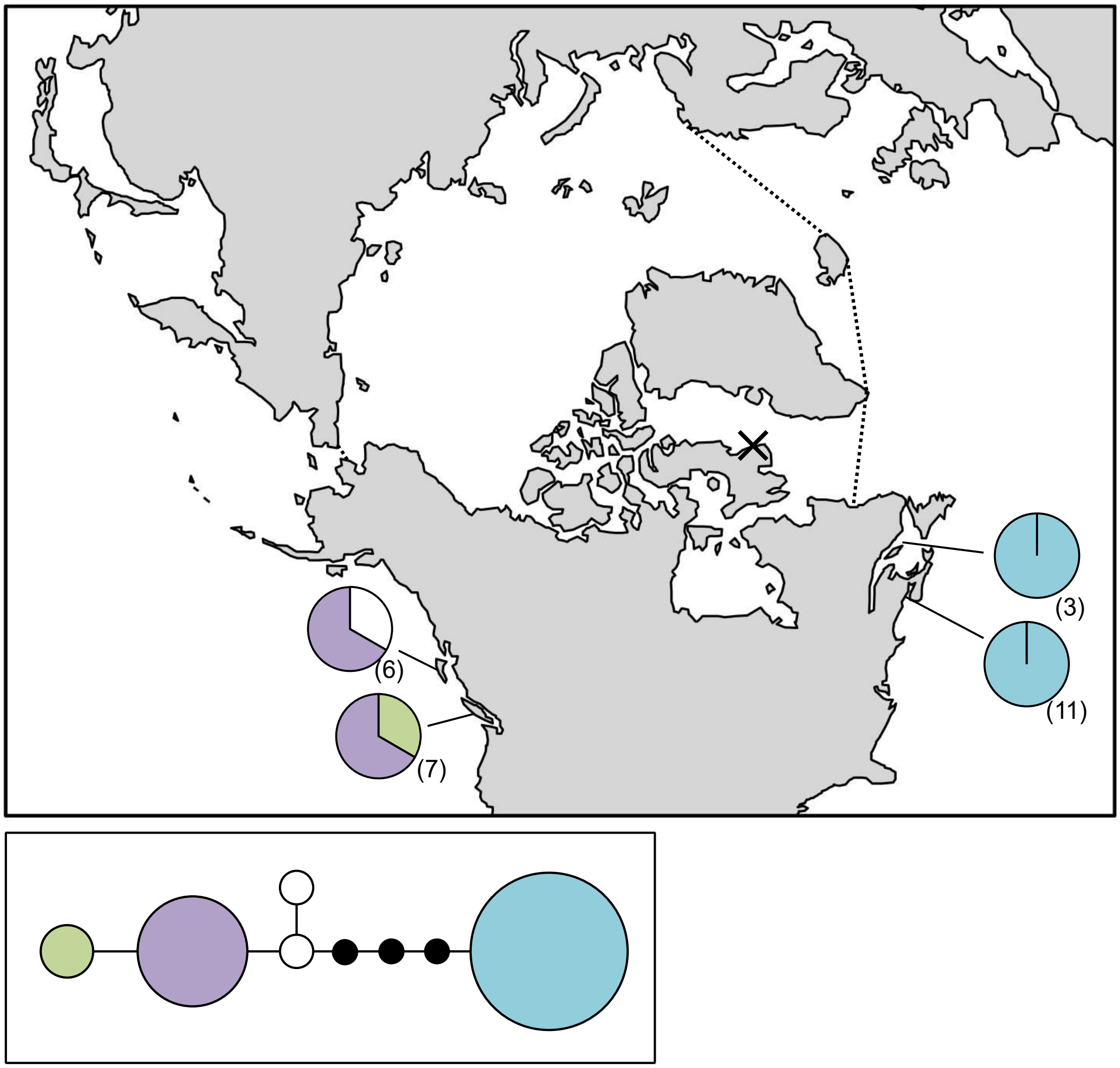

**Figure S75.** *Ralfsia fungiformis* haplotype map and network based on COI-5P data. In the map, numbers in parentheses refer to sample sizes from given locales. The dashed line indicates delineation of the Arctic Ocean. In the haplotype network, black circles indicate hypothesized (e.g. unsampled) haplotypes between clades. Circle size is proportional to the sampling frequency of a given haplotype.

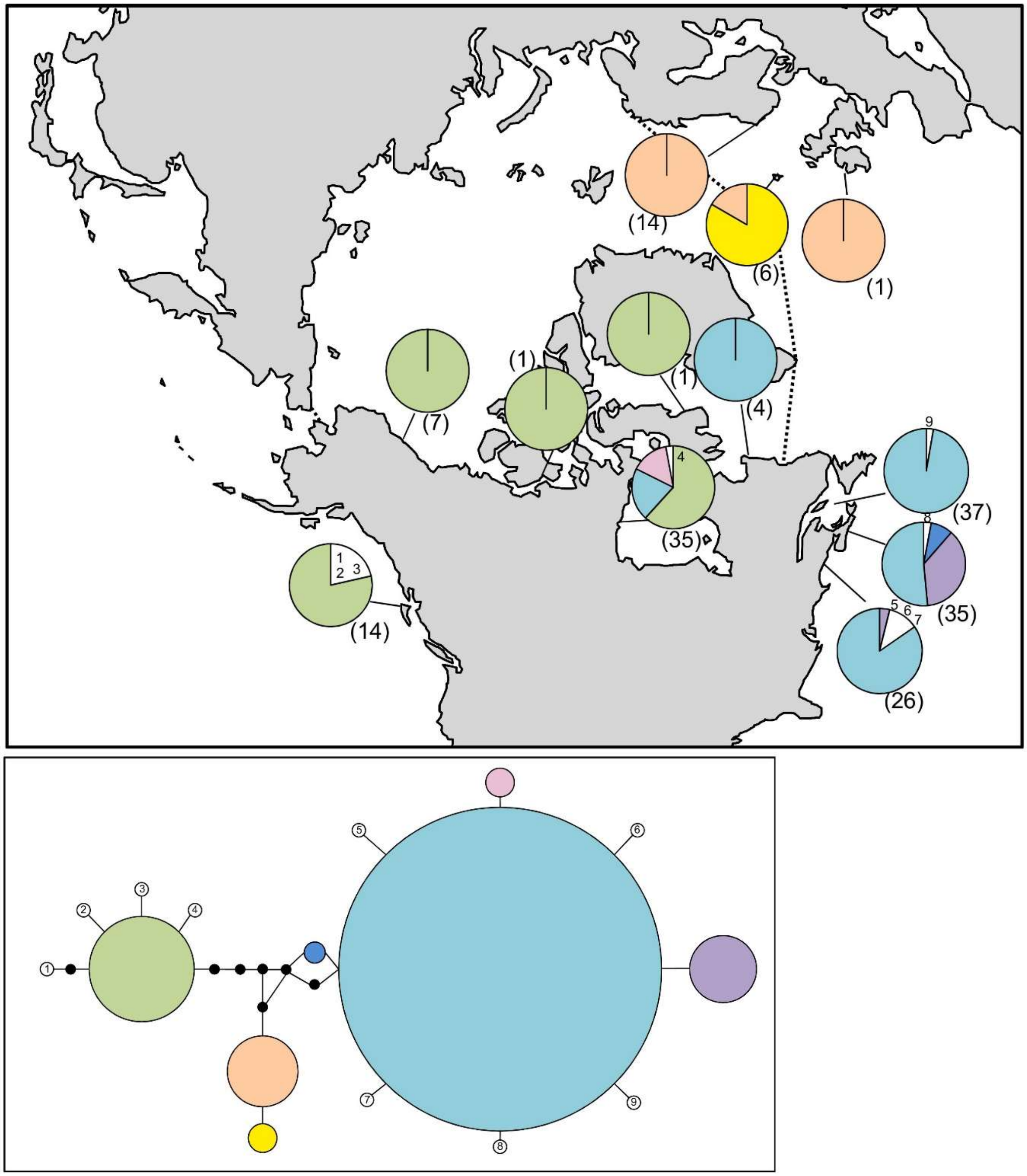

**Figure S76.** *Saccharina latissima* haplotype map and network based on COI-5P data. In the map, numbers in parentheses refer to sample sizes from given locales, whereas numbers adjacent to white portions of pie charts refer to a haplotype sampled only once in the accompanying network. The dashed line indicates delineation of the Arctic Ocean. In the haplotype network, numbered haplotypes in white reference back to the map. Black circles indicate hypothesized (e.g. unsampled) haplotypes between clades. Circle size is proportional to the sampling frequency of a given haplotype.

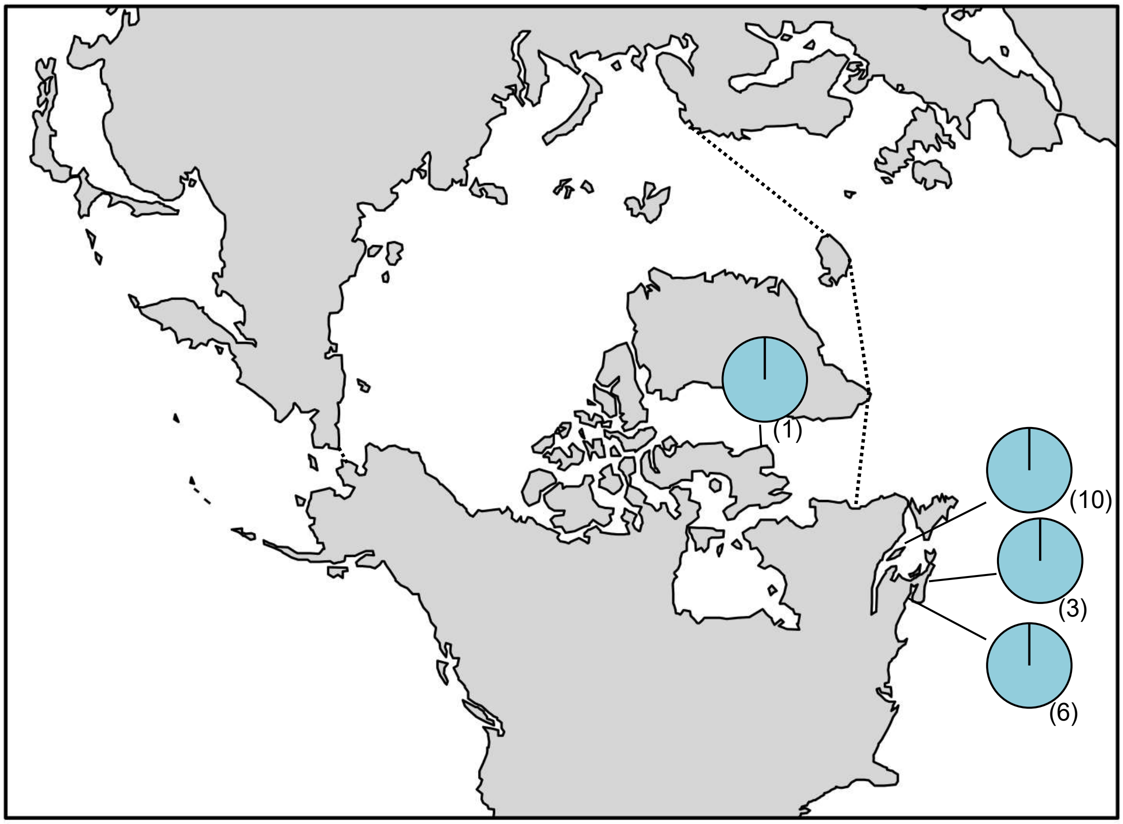

**Figure S77.** *Saccorhiza dermatodea* haplotype map and network based on COI-5P data. In the map, numbers in parentheses refer to sample sizes from given locales. The dashed line indicates delineation of the Arctic Ocean.

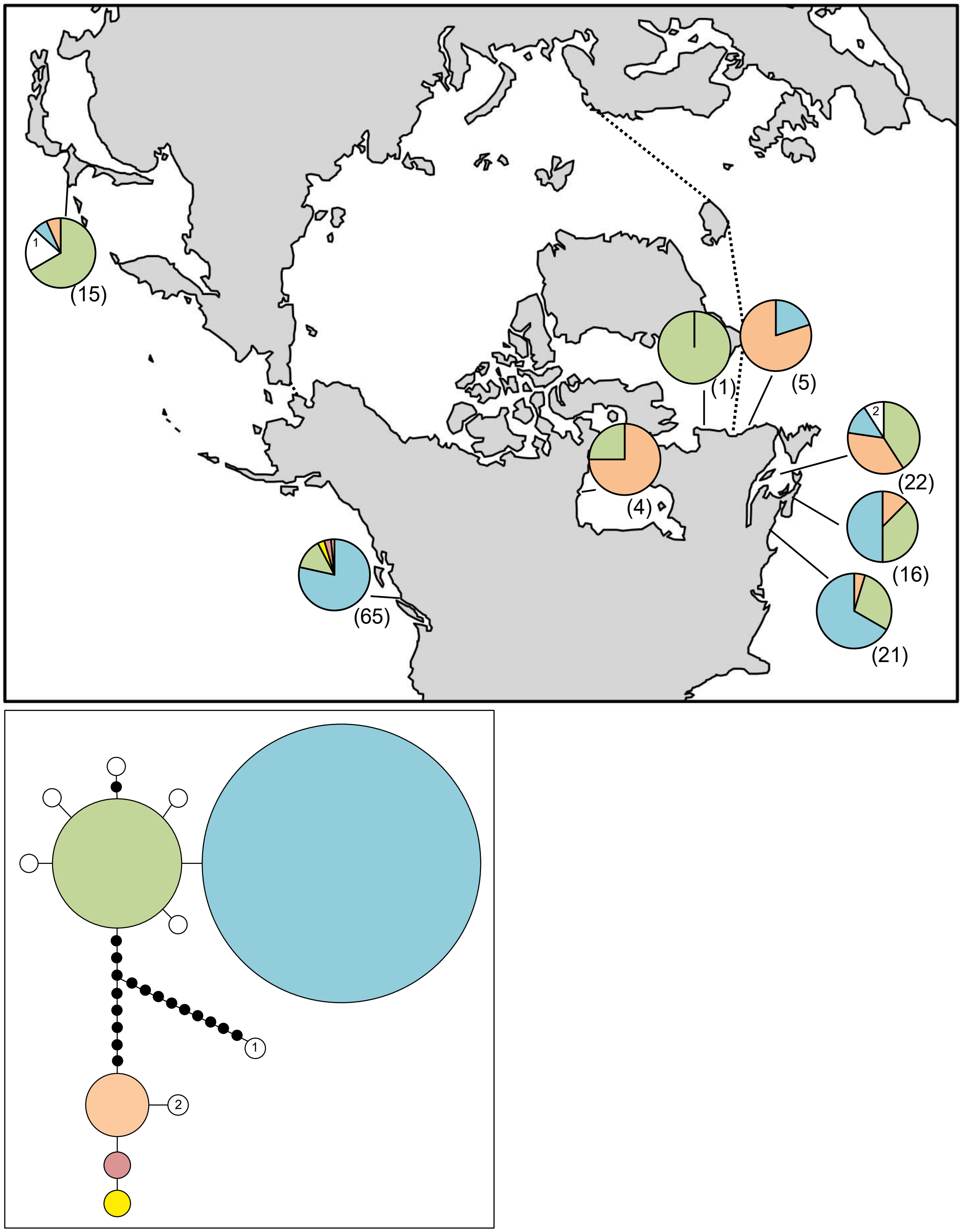

**Figure S78.** *Scytosiphon canaliculatus* haplotype map and network based on COI-5P data. In the map, numbers in parentheses refer to sample sizes from given locales. The dashed line indicates delineation of the Arctic Ocean. In the haplotype network, black circles indicate hypothesized (e.g. unsampled) haplotypes between clades. Circle size is proportional to the sampling frequency of a given haplotype.

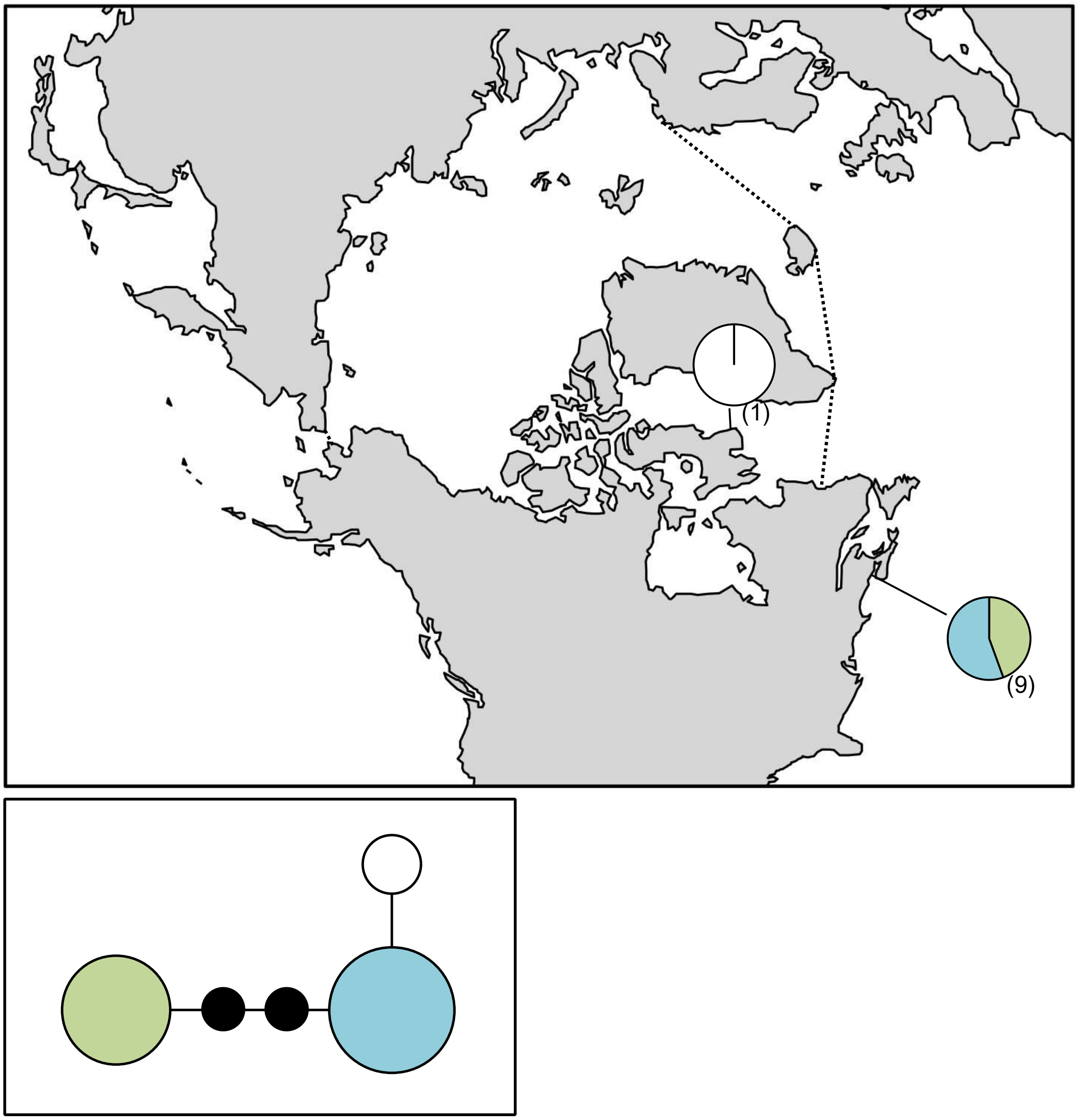

**Figure S79.** *Scytosiphon* sp. GroupJ haplotype map and network based on COI-5P data. In the map, numbers in parentheses refer to sample sizes from given locales. The dashed line indicates delineation of the Arctic Ocean. In the haplotype network, black circles indicate hypothesized (e.g. unsampled) haplotypes between clades. Circle size is proportional to the sampling frequency of a given haplotype.

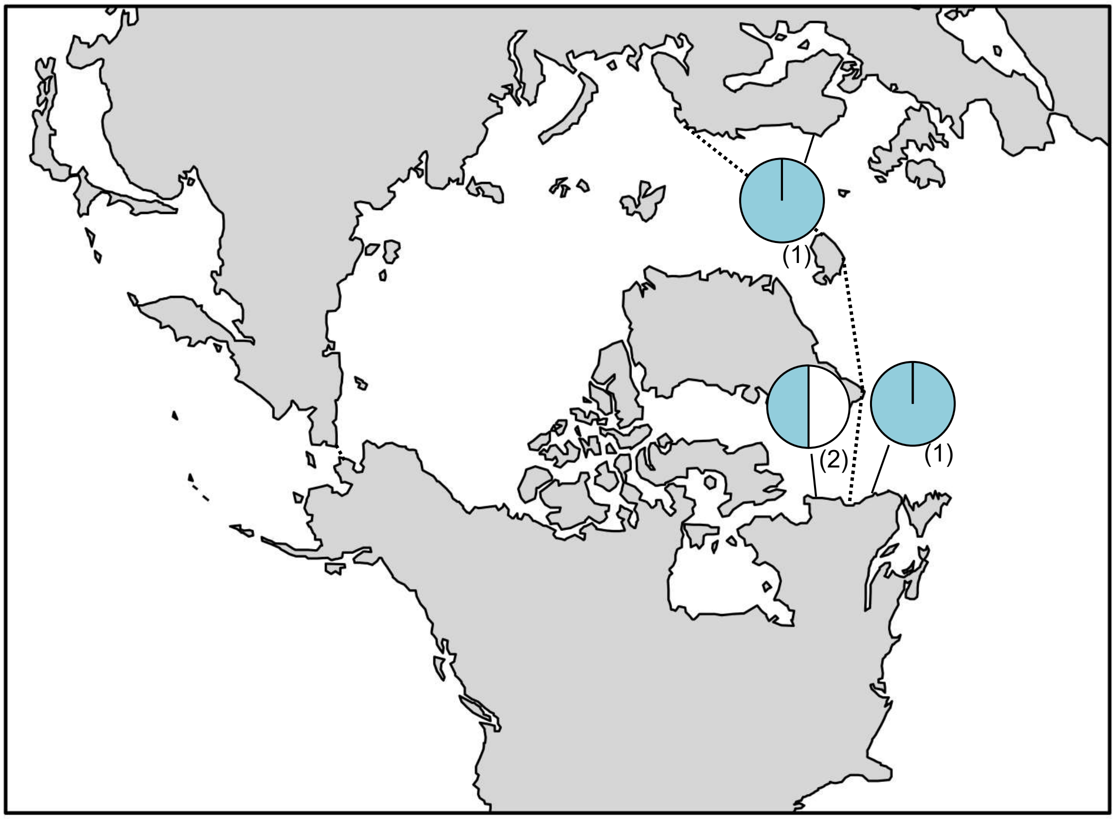

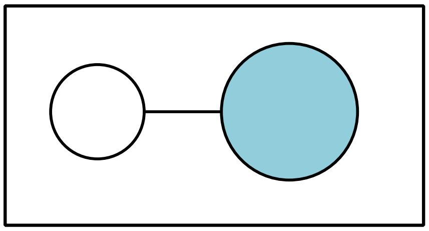

**Figure S80.** Tilopteridalean sp. 1GWS haplotype map and network based on COI-5P data. In the map, numbers in parentheses refer to sample sizes from given locales. The dashed line indicates delineation of the Arctic Ocean. In the haplotype network, circle size is proportional to the sampling frequency of a given haplotype.

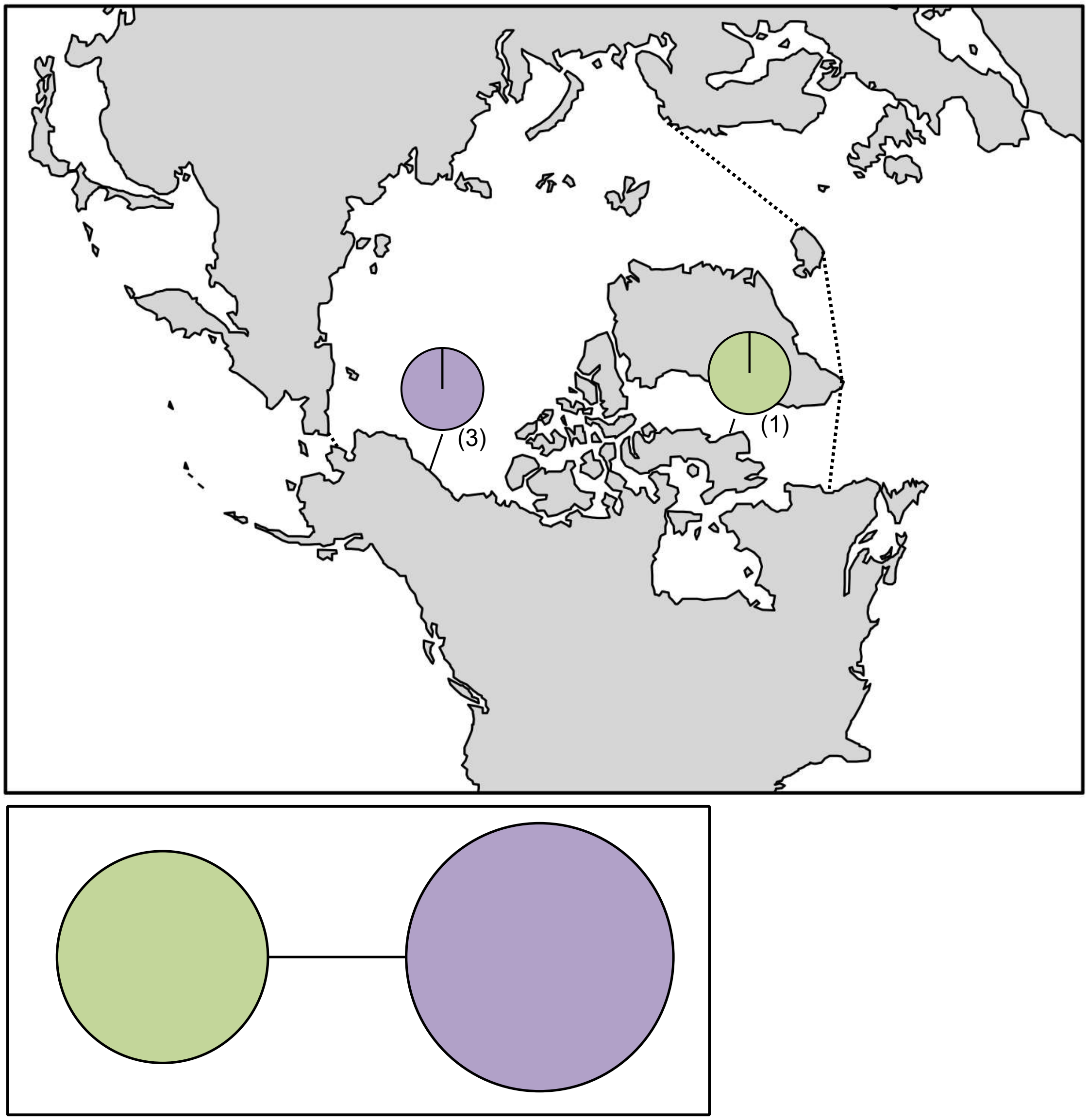

**Figure S81.** Tilopteridalean sp. 2GWS haplotype map and network based on COI-5P data. In the map, numbers in parentheses refer to sample sizes from given locales. The dashed line indicates delineation of the Arctic Ocean. In the haplotype network, circle size is proportional to the sampling frequency of a given haplotype.

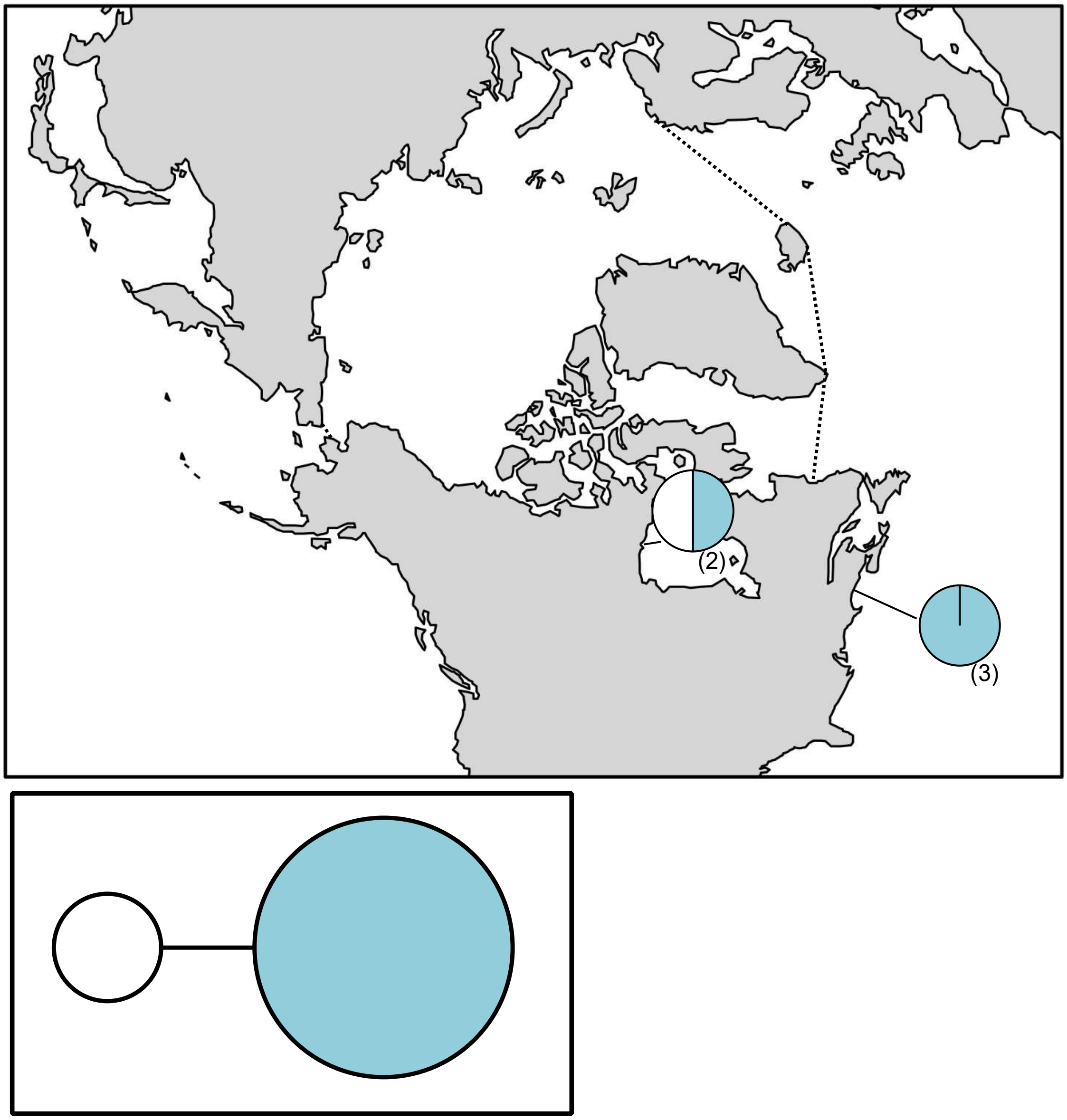

**Figure S82.** *Acrosiphonia* sp. 3GWS haplotype map and network based on *tufA* data. In the map, numbers in parentheses refer to sample sizes from given locales. The dashed line indicates delineation of the Arctic Ocean. In the haplotype network, circle size is proportional to the sampling frequency of a given haplotype.

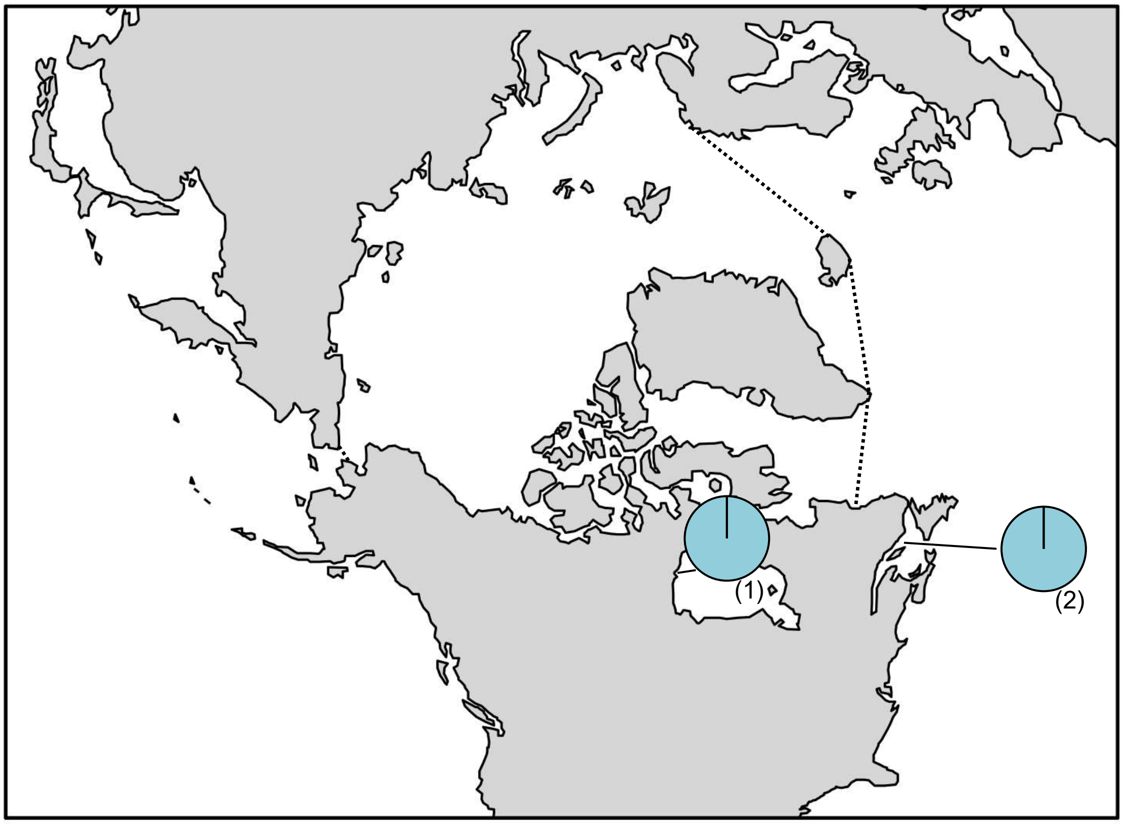

**Figure S83.** *Acrosiphonia* sp. 6GWS haplotype map based on *tufA* data. In the map, numbers in parentheses refer to sample sizes from given locales. The dashed line indicates delineation of the Arctic Ocean.

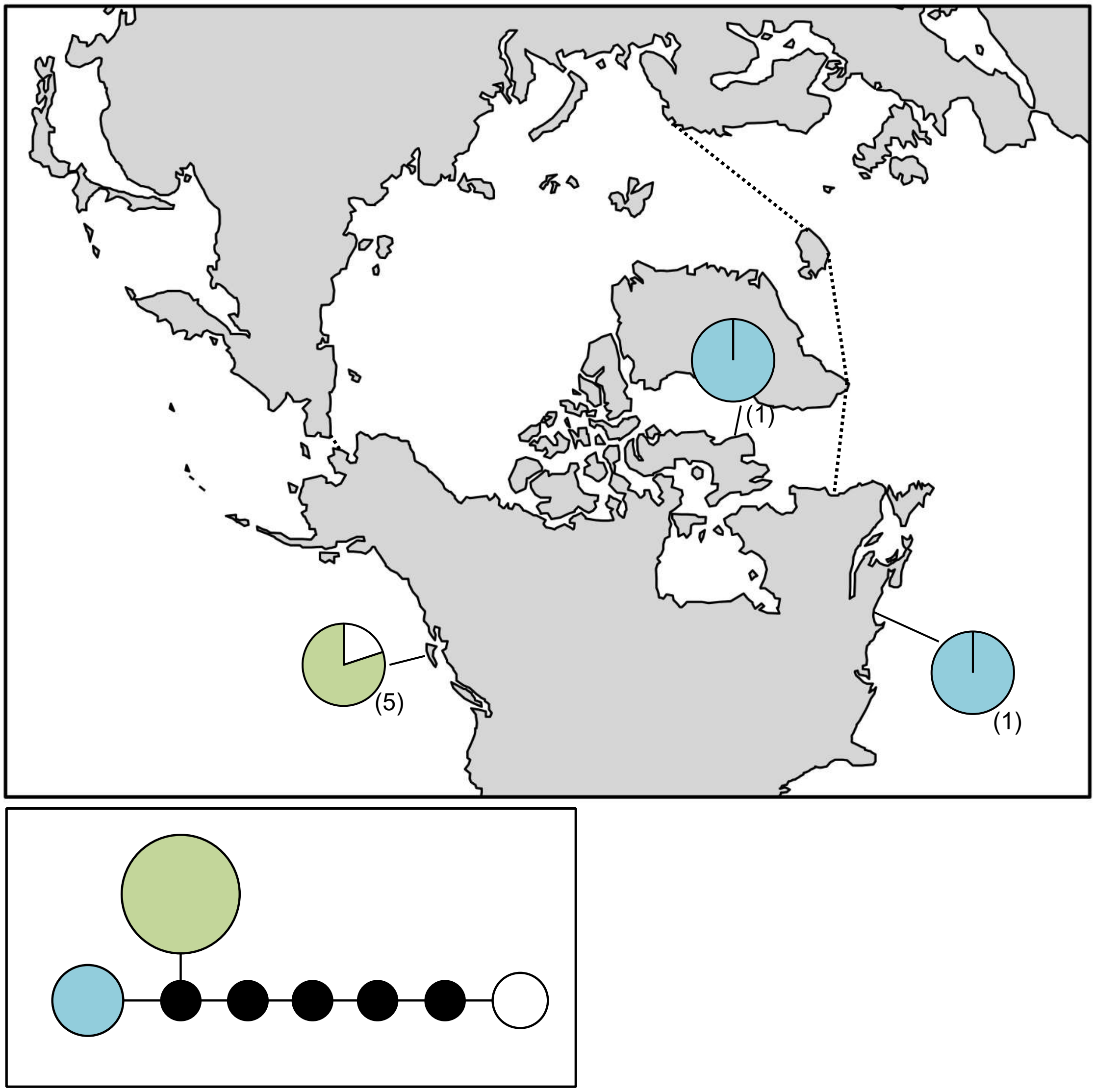

**Figure S84.** *Acrosiphonia* sp. 8GWS haplotype map and network based on *tufA* data. In the map, numbers in parentheses refer to sample sizes from given locales. The dashed line indicates delineation of the Arctic Ocean. In the haplotype network, the black circle indicates a hypothesized (e.g. unsampled) haplotypes between clades. Circle size is proportional to the sampling frequency of a given haplotype.

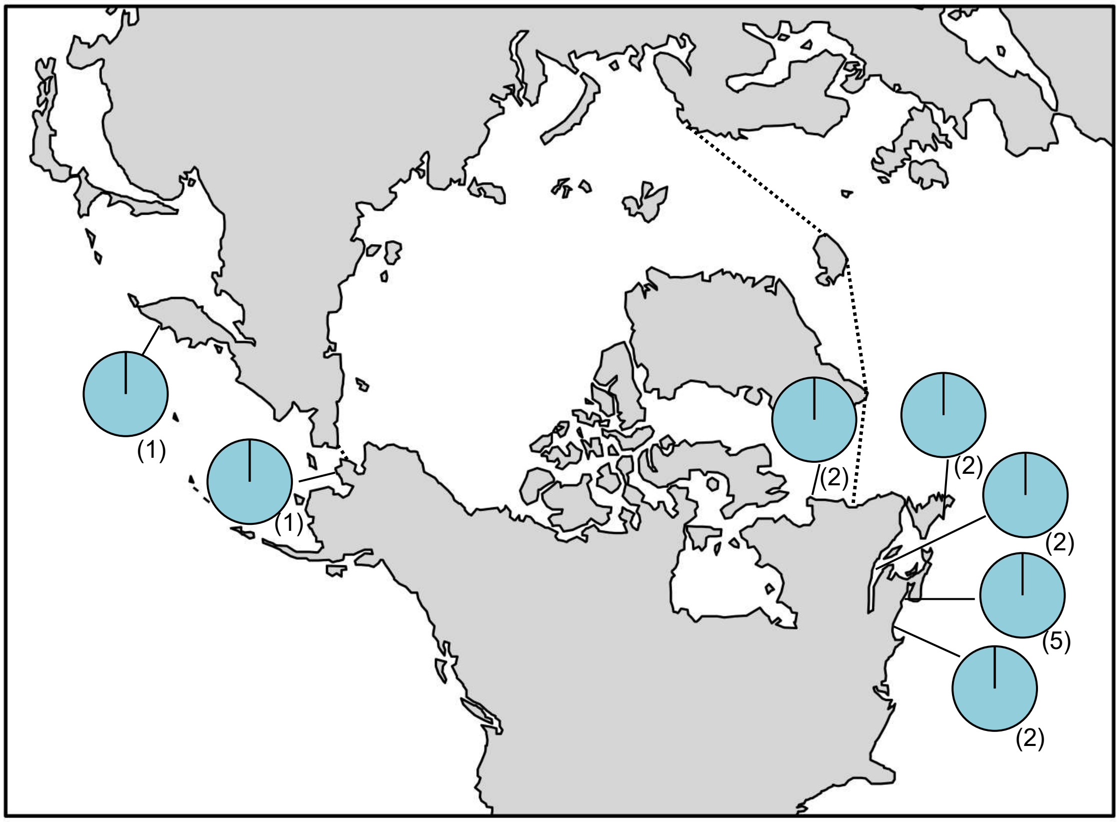

**Figure S85.** *Acrosiphonia sonderi* haplotype map based on *tufA* data. In the map, numbers in parentheses refer to sample sizes from given locales. The dashed line indicates delineation of the Arctic Ocean.

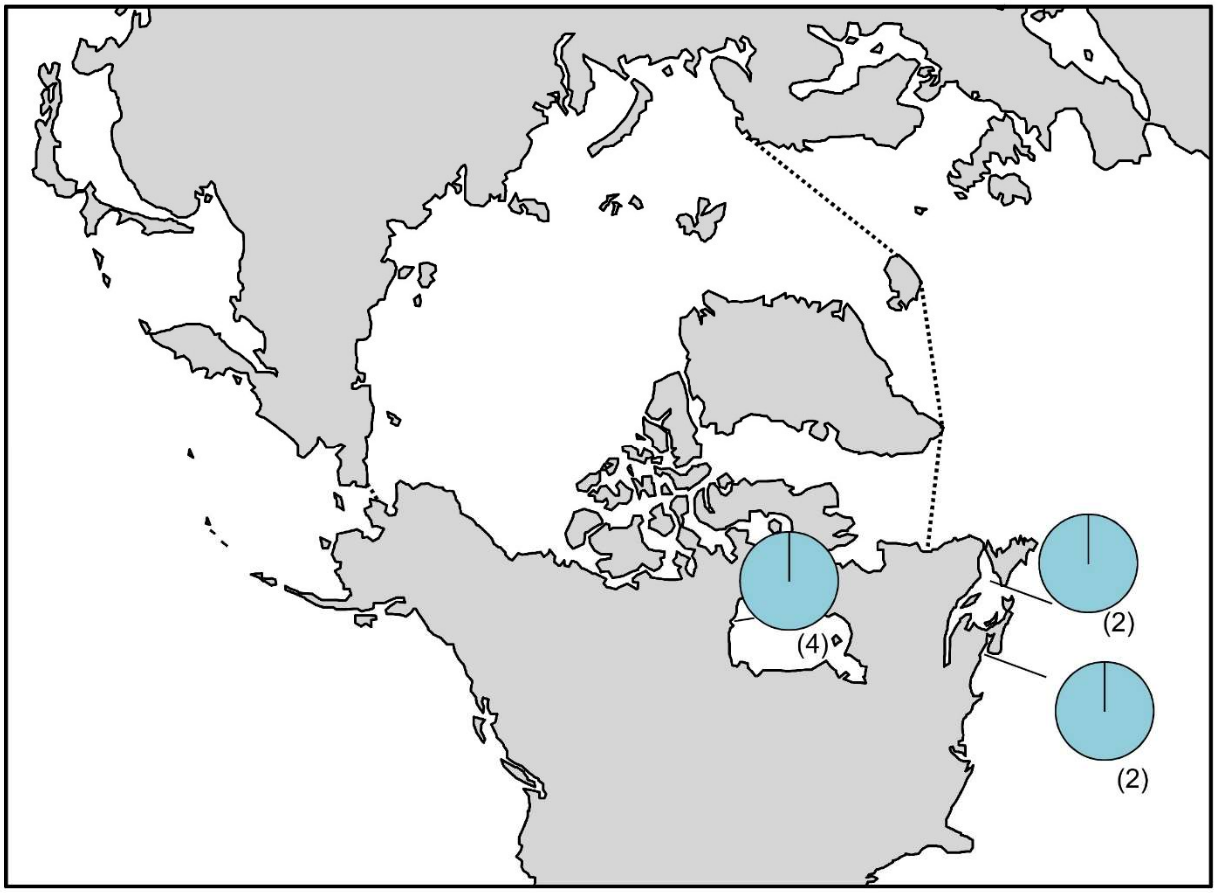

**Figure S86.** *Blidingia* sp. 3GWS haplotype map based on *tufA* data. In the map, numbers in parentheses refer to sample sizes from given locales. The dashed line indicates delineation of the Arctic Ocean.

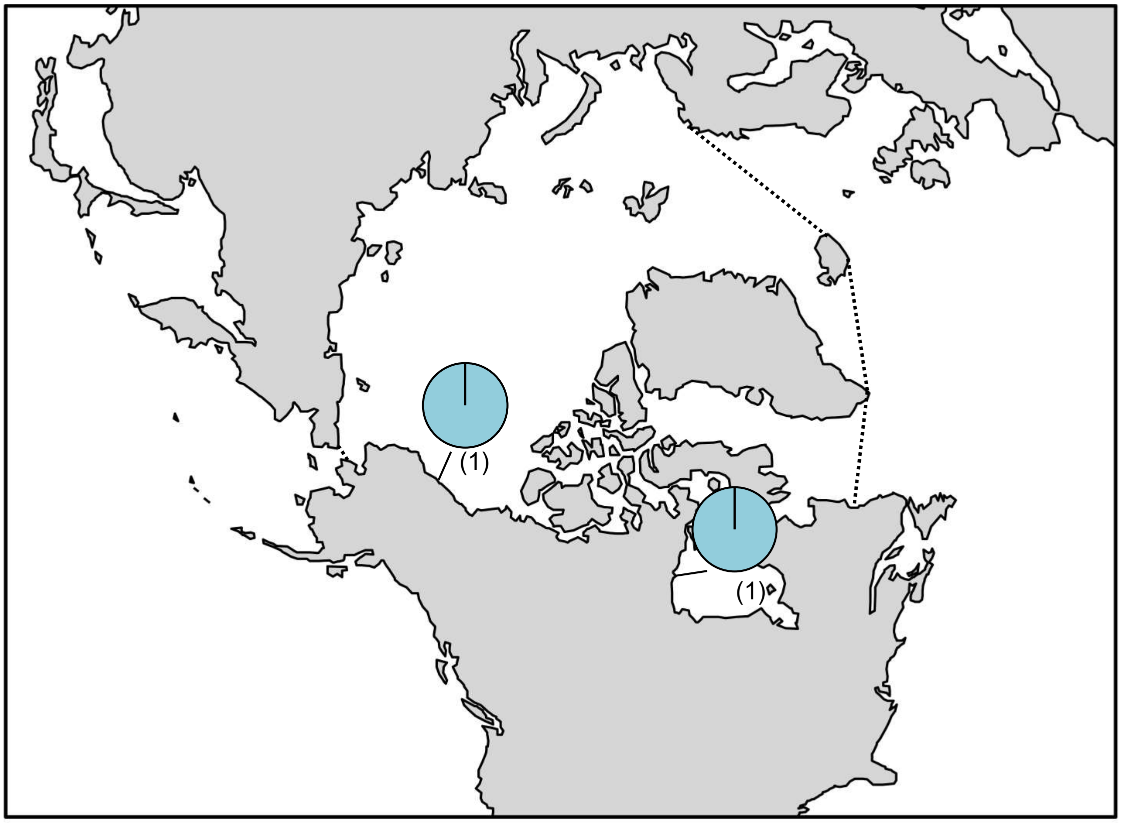

**Figure S87.** *Blidingia* sp. 5GWS haplotype map based on *tufA* data. In the map, numbers in parentheses refer to sample sizes from given locales. The dashed line indicates delineation of the Arctic Ocean.

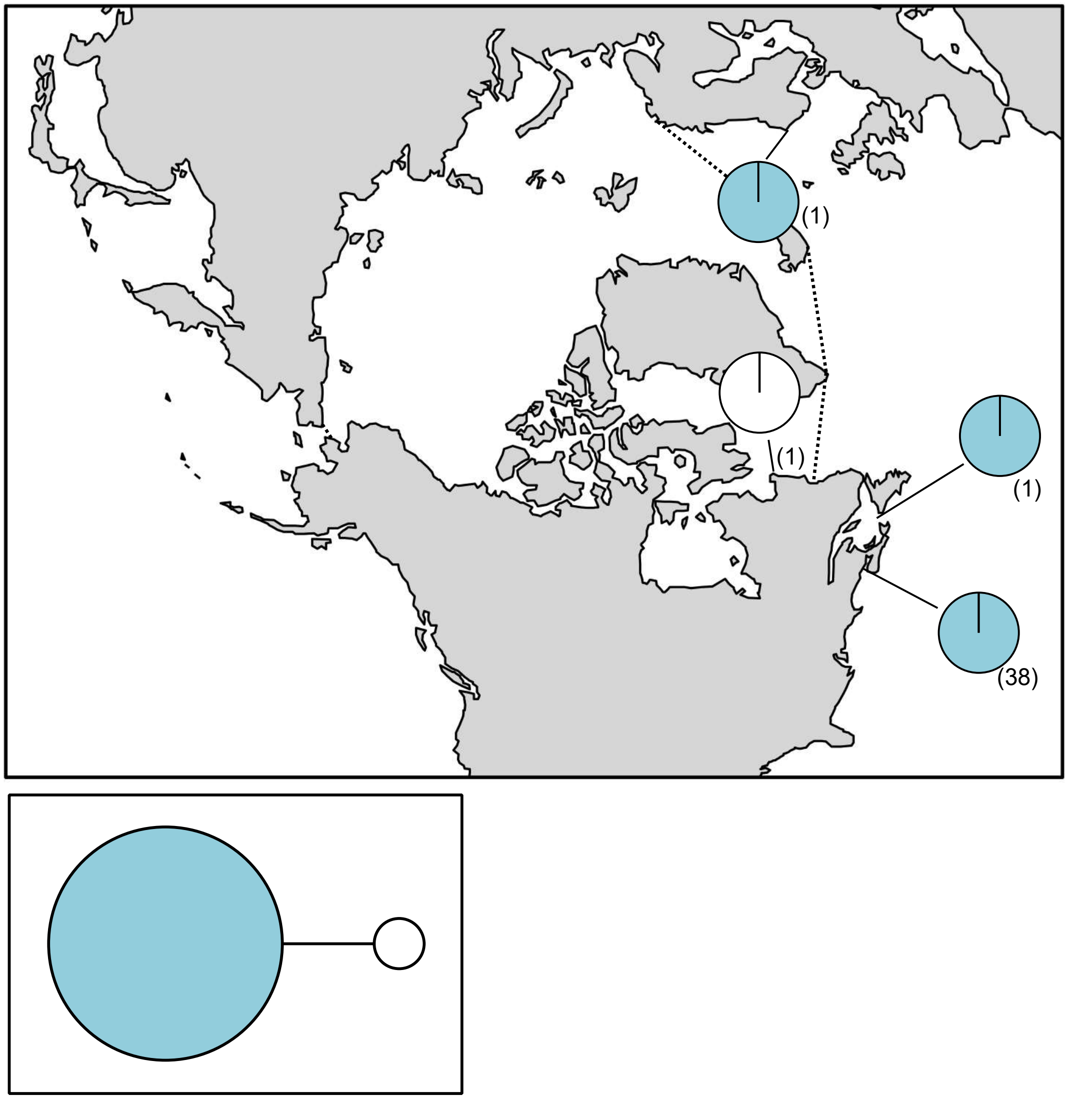

**Figure S88.** *Monostroma* sp. 2grevillei haplotype map based on *tufA* data. In the map, numbers in parentheses refer to sample sizes from given locales. The dashed line indicates delineation of the Arctic Ocean. In the haplotype network, circle size is proportional to the sampling frequency of a given haplotype.

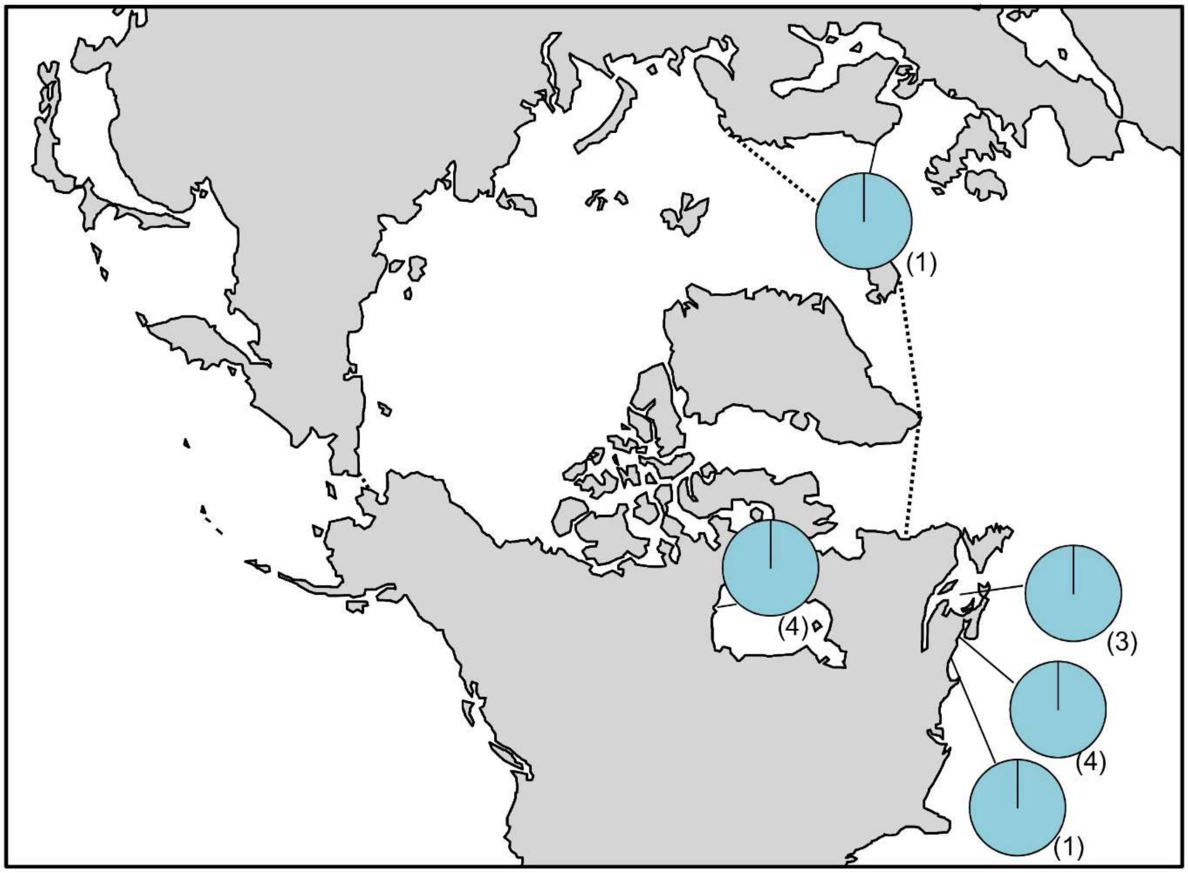

**Figure S89.** *Spongomorpha aeruginosa* haplotype map based on *tufA* data. In the map, numbers in parentheses refer to sample sizes from given locales. The dashed line indicates delineation of the Arctic Ocean.

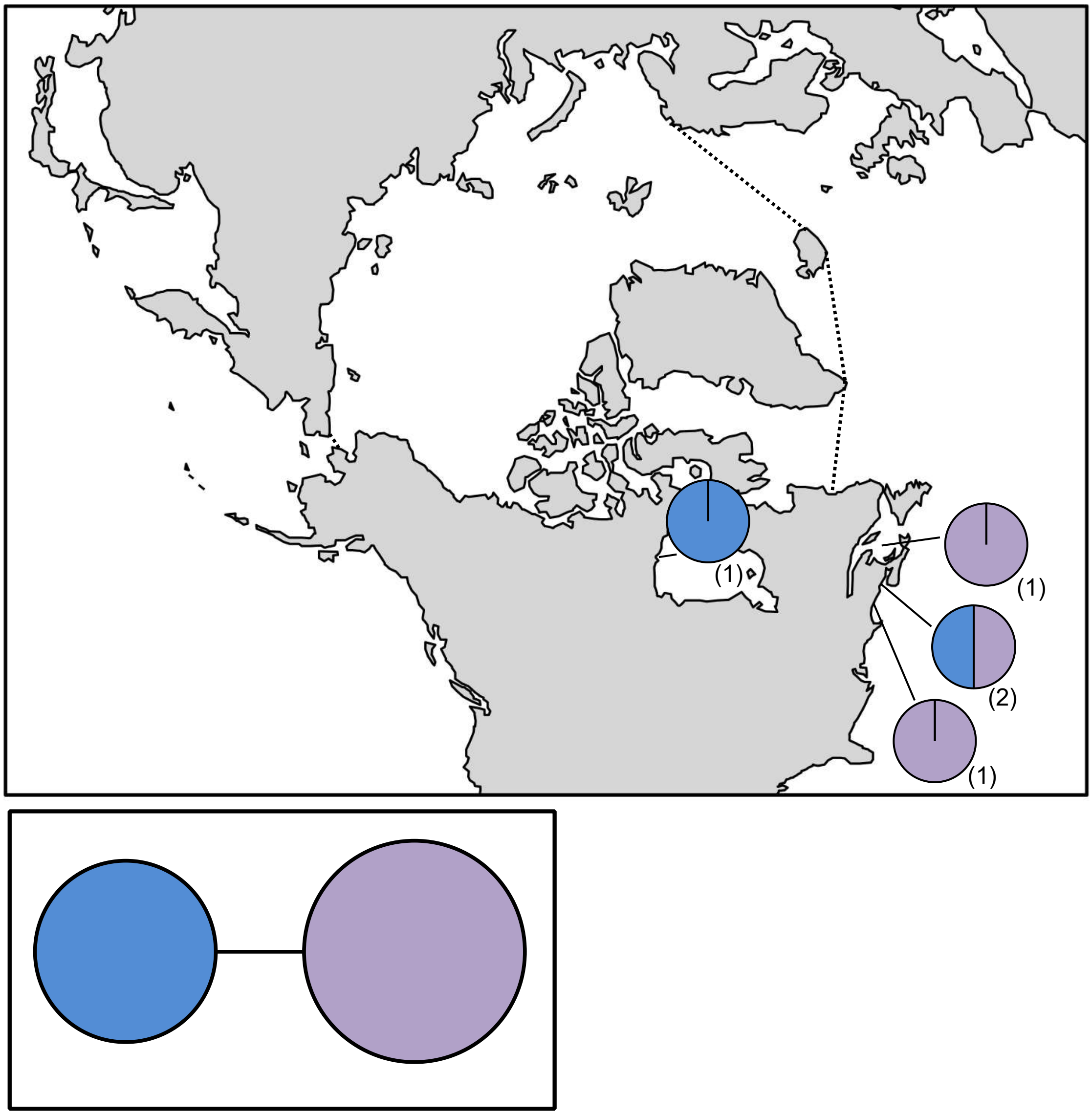

**Figure S90.** *Ulothrix flacca* haplotype map and network based on *rbc*L-3P data. In the map, numbers in parentheses refer to sample sizes from given locales. The dashed line indicates delineation of the Arctic Ocean. In the haplotype network, circle size is proportional to the sampling frequency of a given haplotype.

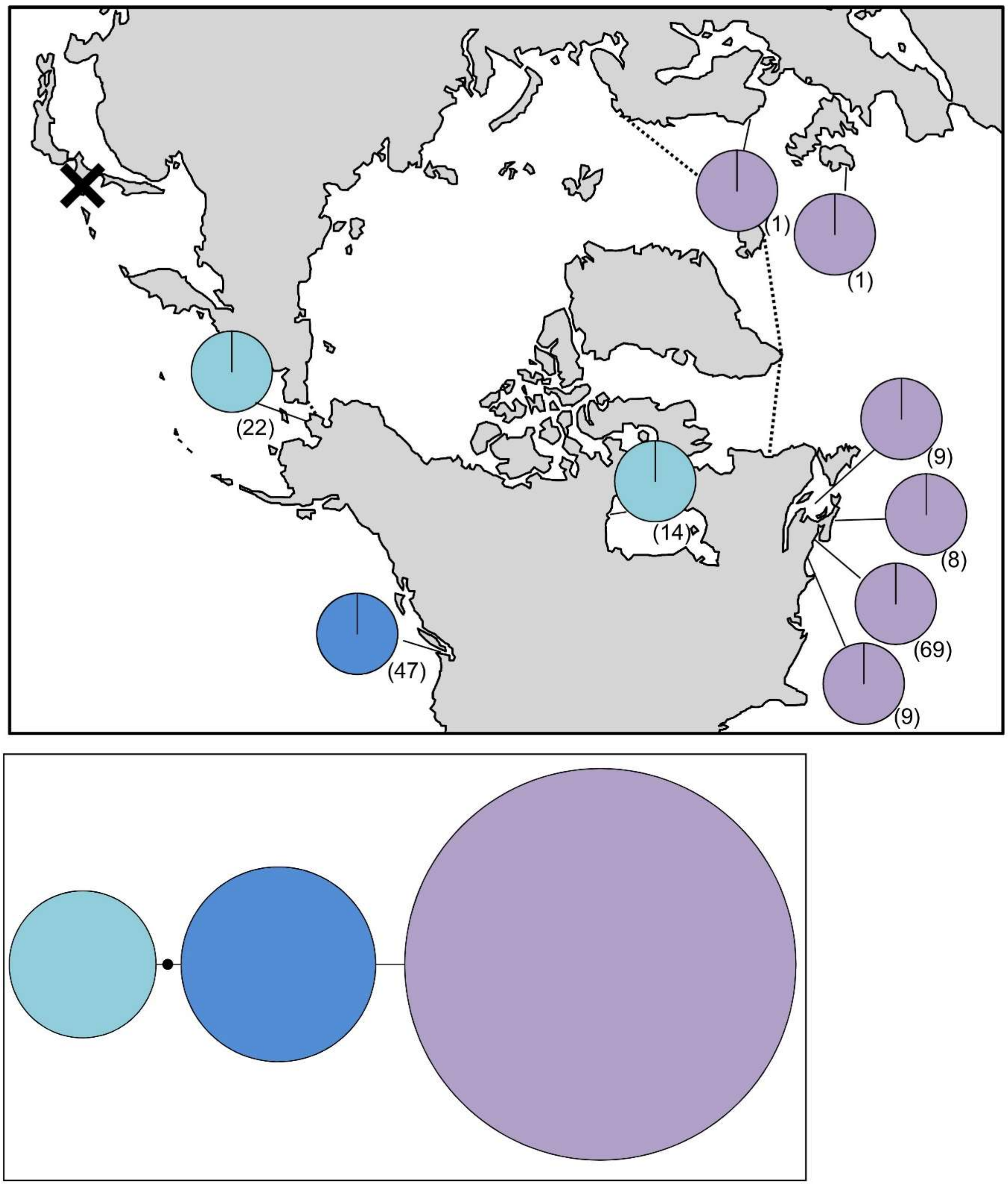

**Figure S91.** *Ulva fenestrata* haplotype map and network based on *tufA* data. In the map, numbers in parentheses refer to sample sizes from given locales. The dashed line indicates delineation of the Arctic Ocean, while the black X indicates a location with genetically verified *Ulva fenestrata* based on *rbc*L. In the haplotype network, the black circle indicates a hypothesized (e.g. unsampled) haplotype between clades. Circle size is proportional to the sampling frequency of a given haplotype.

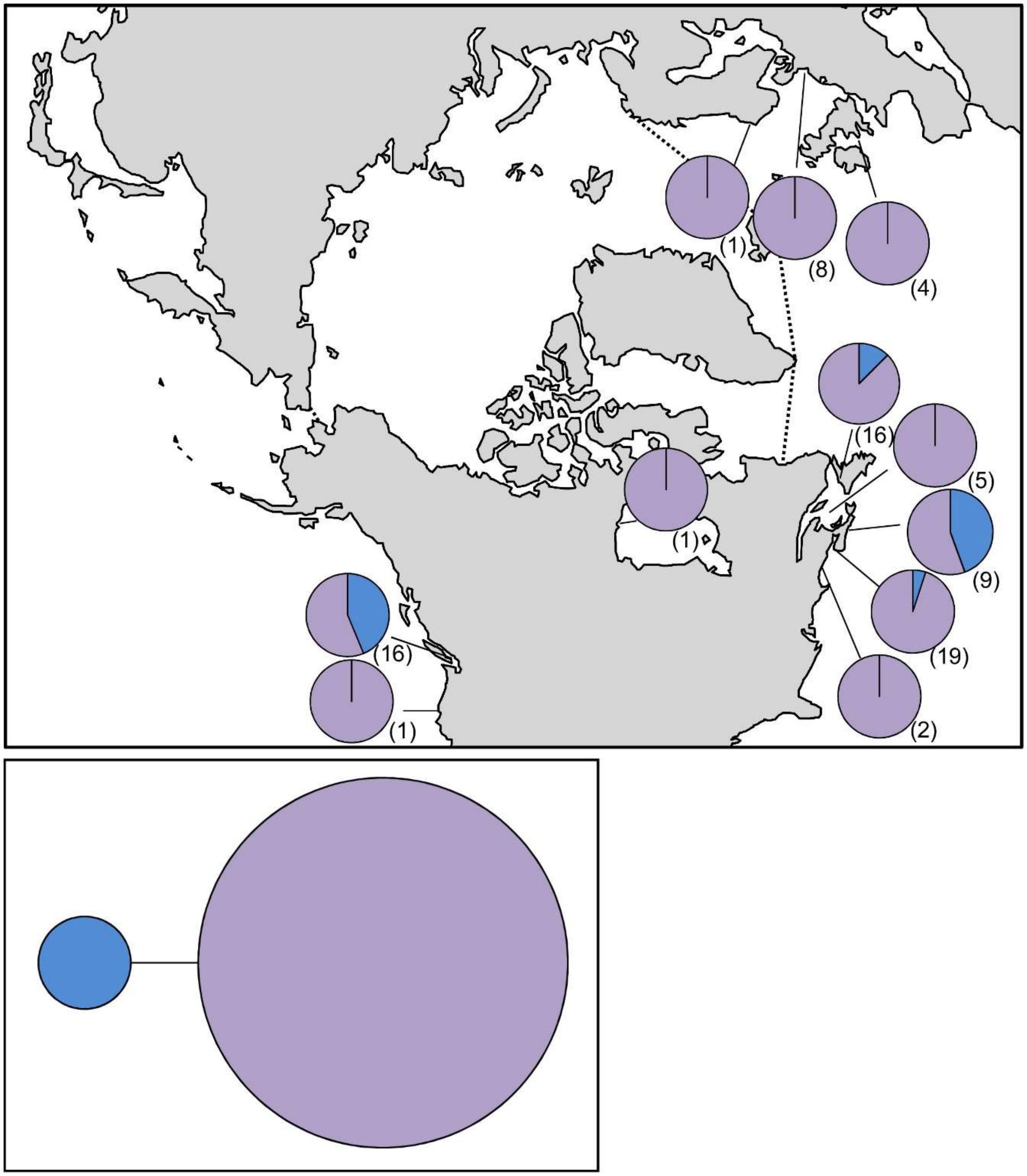

**Figure S92.** *Ulva intestinalis* haplotype map and network based on *tufA* data. In the map, numbers in parentheses refer to sample sizes from given locales. The dashed line indicates delineation of the Arctic Ocean. In the haplotype network, circle size is proportional to the sampling frequency of a given haplotype. The dashed line indicates haplotypes with a substitution at the same site.

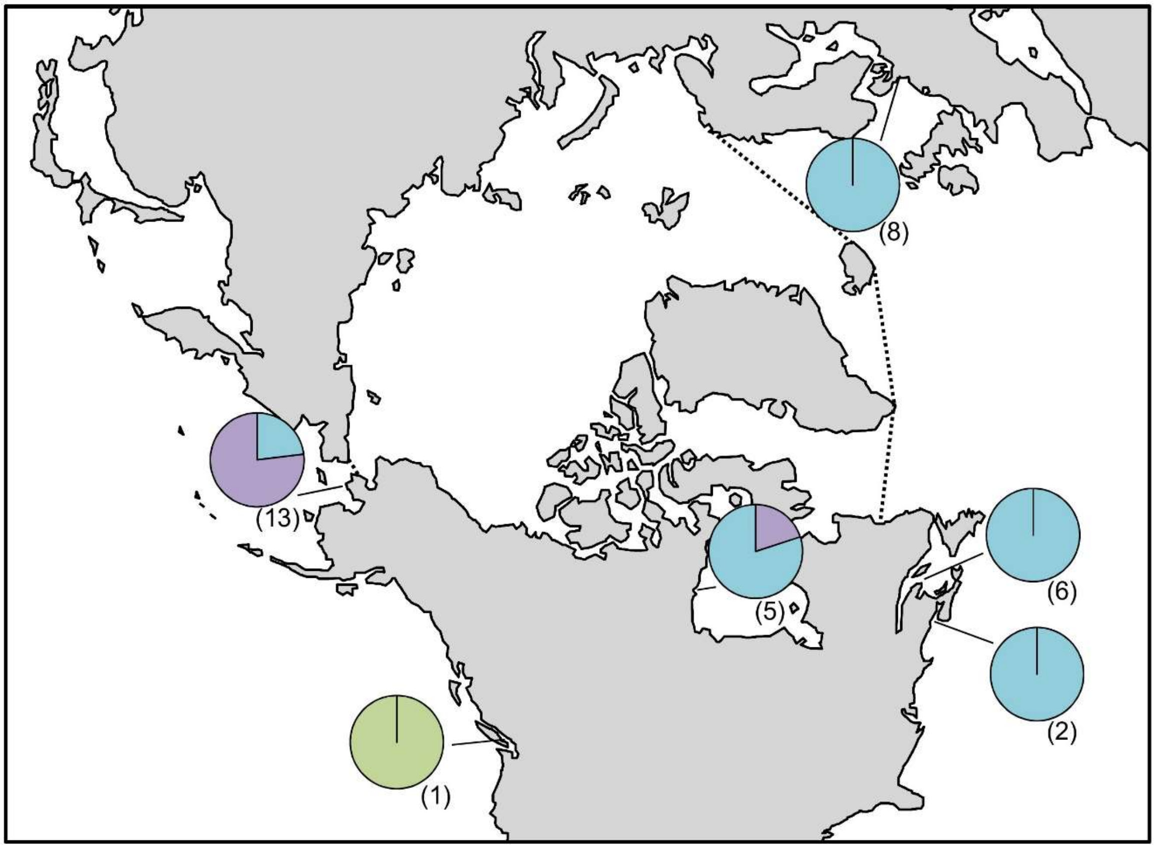

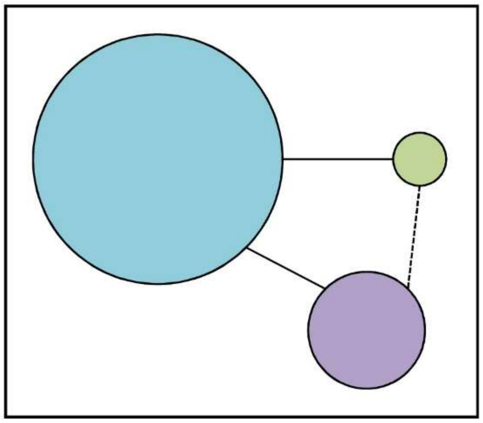

**Figure S93.** *Ulva* sp. 3linza haplotype map and network based on *tufA* data. In the map, numbers in parentheses refer to sample sizes from given locales. The dashed line indicates delineation of the Arctic Ocean. In the haplotype network, circle size is proportional to the sampling frequency of a given haplotype. The dashed line indicates haplotypes with a substitution at the same site.

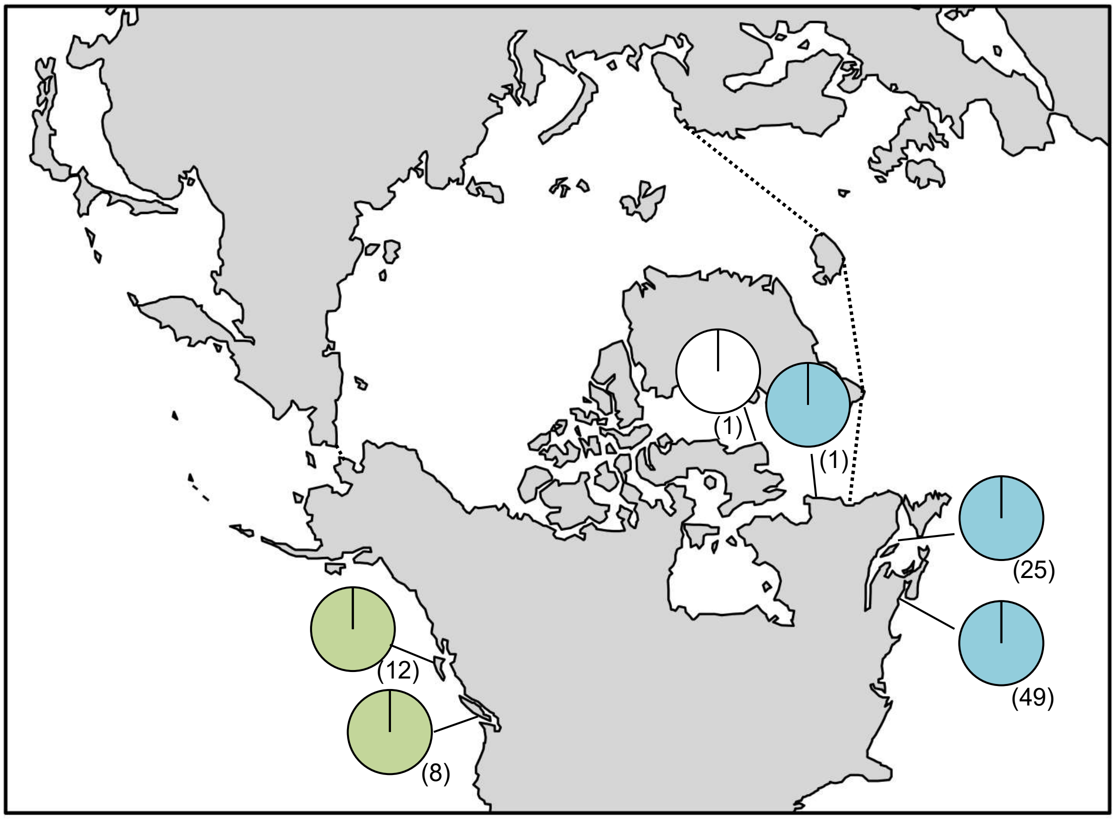

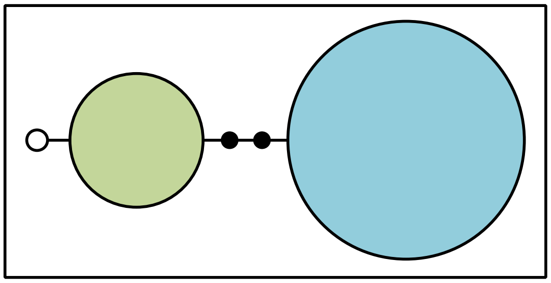

**Figure S94.** *Ulvaria obscura* haplotype map based on *tufA* data. In the map, numbers in parentheses refer to sample sizes from given locales. The dashed line indicates delineation of the Arctic Ocean. In the haplotype network, the black circle indicates a hypothesized (e.g. unsampled) haplotypes between clades. Circle size is proportional to the sampling frequency of a given haplotype.

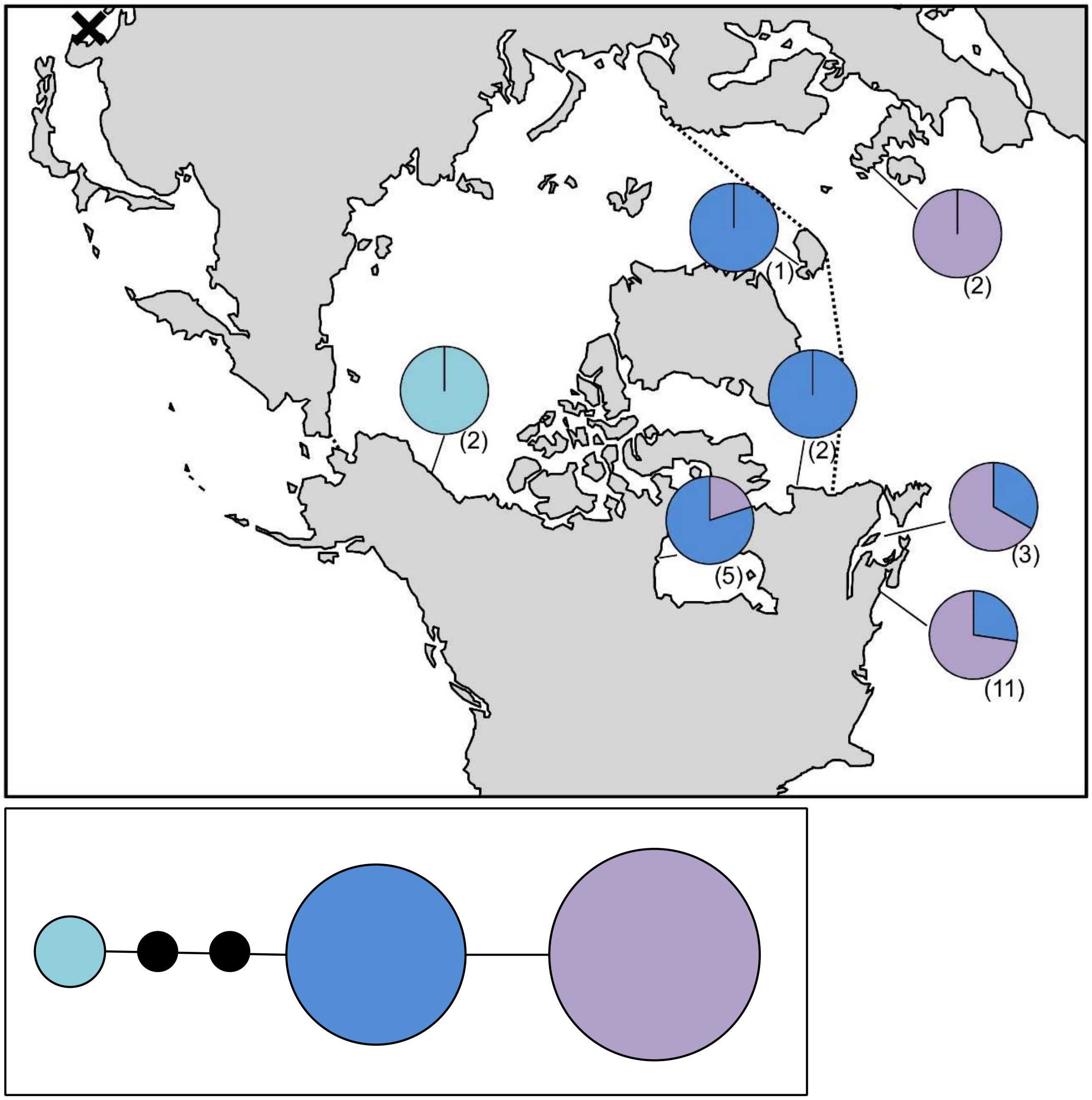

**Figure S95.** *Ulva prolifera* haplotype map and network based on *tufA* data. In the map, numbers in parentheses refer to sample sizes from given locales. The dashed line indicates delineation of the Arctic Ocean, while the black X indicates a location with genetically verified *Ulva prolifera* based on *rbc*L. In the haplotype network, the black circle indicates hypothesized (e.g. unsampled) haplotypes between clades. Circle size is proportional to the sampling frequency of a given haplotype.

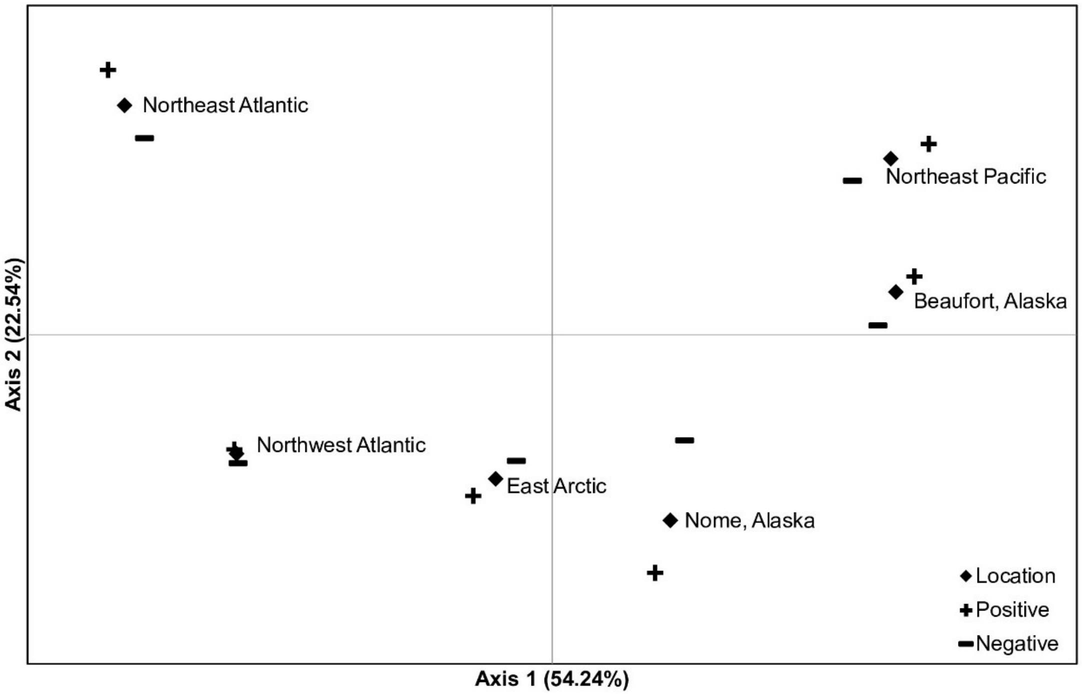

**Figure S96**. PCoA analysis without "normalizing" Beaufort, Northeast Pacific relationship.

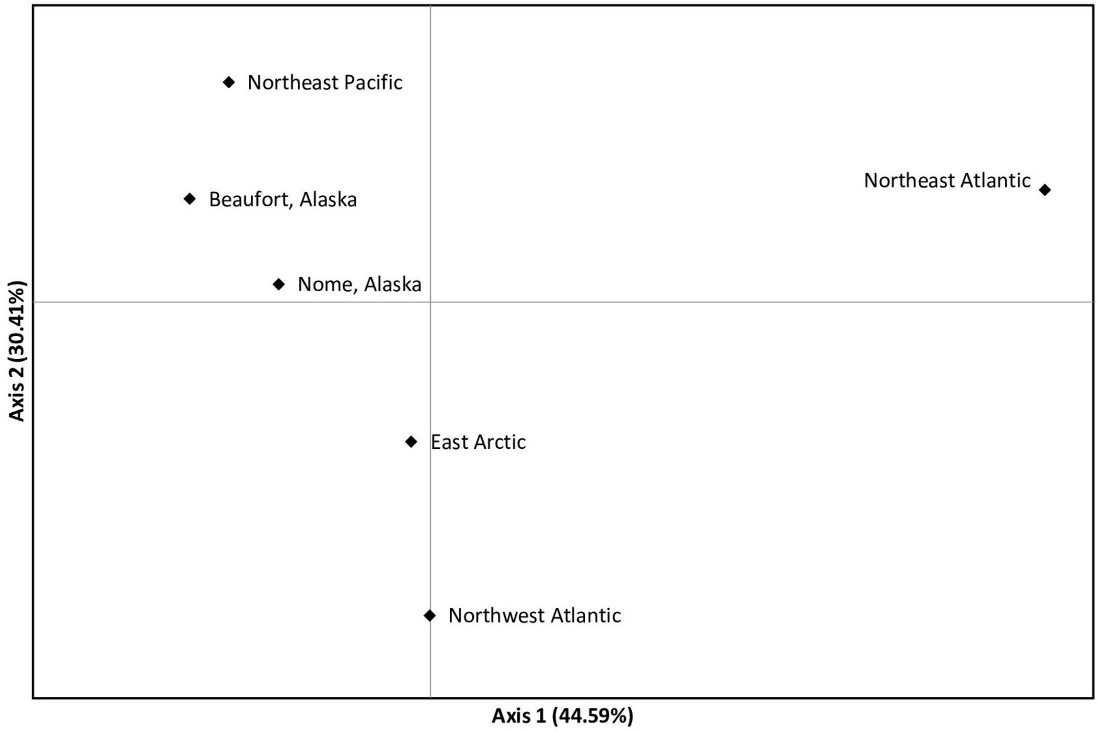

**Figure S97**. PCoA analysis with low sample populations (<10 individuals) removed from averages.

**Table S1**. General haplotype patterns and inferred origins in Arctic species of marine macroalgae. The Arctic basin is delineated according to the 10ºC isotherm for July, as per D'Odorico et al. (2013), and includes Northern Baffin Island through to Northern Labrador (Nain and Northwards), Svalbard, Northern Norway, the Siberian coastline, Northern Alaska (North of the Bering Strait), and the Northern Canadian coastline, including the Hudson Bay. Species with updated information regarding the origins of Arctic populations (relative to Saunders and McDevit [2013]) are indicated below the species name as updated, or are listed as new if they were not reported in that publication. Pa=Pacific, Ar=Arctic, At=Atlantic. Sample sizes refer to COI-5P data unless otherwise indicated. For the origin of Arctic specimens, ocean basins not in parentheses do not allow for an Arctic refugial populations (scenario 1), while those in parentheses indicate interpretation of haplotype data if Arctic refugial populations are considered as a possible source for contemporary Arctic populations (scenario 2). [1]Churchill, Manitoba, records cannot be accounted for by Pacific or Atlantic collections (e.g. unique Arctic species or haplotype[s] suggesting at Arctic periglacial refugial origins). [2]North Alaska records cannot be accounted for by Pacific or Atlantic collections. [3]Baffin Island and Northern Labrador records cannot be accounted for by Pacific or Atlantic collections.

| Species | Sample size (Pa/Ar/At) | Origin of Arctic specimens | Interpretation of haplotype patterns |
|---|---|---|---|
| Rhodophyta | | | |
| *Acrochaetium* sp.[1] | 0/1/0 | Uncertain (Arctic) | A single record exists for this genetic group, occurring in Churchill, leaving the origin uncertain. |
| *Ahnfeltia borealis* **(Updated)** (Figs. S3 & S4) | COI-5P: 25/44/2 *ycf35*: 24/35/2 | Pacific | Very little haplotype variation exists in this species. Divergent COI-5P haplotypes occur in the Northwest Atlantic, while Arctic haplotypes are monotypic, matching Pacific populations. *ycf35* haplotypes indicate two unique microsatellites occur in the Bering Sea, whereas only one of these microsatellites occurs in the Arctic. Available evidence therefore suggests a Pacific origin, possibly out of the Northwest Pacific given the low number of sampled haplotypes. |
| *Ahnfeltia plicata* (Fig. S5) | 0/2/140 | Atlantic | Haplotype patterns indicate this species has a long history in the Atlantic. Specimens of *Ahnfeltia* in the West Arctic and Bering Sea are assignable to *A. borealis* rather than *A. plicata*, suggesting that the latter species has a limited Artic distribution. Our verified collections in the North American Arctic are limited to two drift collections from Churchill, with |

| | | | |
|---|---|---|---|
| | | | Baffin Island and Labrador collections all assignable to *Ahnfeltia borealis* (n=10). |
| *Ceramium virgatum* **(New)** (Fig. S6) | 0/1/170 | Atlantic | A single Arctic collection along the coast of Labrador matches Northwest Atlantic populations, where this species appears to have survived multiple glaciations. |
| *Clathromorphum* sp. 9GWS **(Updated)** (Fig. S7) | 2/3/65 | Uncertain | Previously listed as *Phymatolithon lenormandii* in Saunders & McDevit (11), this genetic group has been updated to uncertain given the inclusion of North Pacific records and lack of haplotype variation throughout its genetically confirmed range. |
| *Clathromorphum circumscriptum* **(New)** (Fig. S8) | 0/1/6 | Atlantic | A single Arctic collection from Baffin Island matches a Northwest Atlantic haplotype. |
| *Clathromorphum compactum* **(New)** (Fig. S9) | 0/1/11 | Atlantic | A single Arctic collection along the coast of Labrador matches the Northwest Atlantic haplotype. |
| *Coccotylus brodiei* **(Updated)** (Fig. S10) | 0/1/101 | Atlantic | All verified Pacific and West Arctic records are attributable to *Coccotylus truncatus*; as such, the single Arctic collection, from Churchill appears to be of Atlantic origin. |
| *Coccotylus truncatus*[1, 2, 3] **(Updated)** (Figs. S11 & S12) | COI-5P: 12/65/55 ITS: 11/54/34 | Atlantic and Pacific (Arctic, Atlantic, and Pacific) | East and West Arctic populations appear to have been recolonized from the Atlantic and Pacific, respectively. ITS data, in particular, appear to have distinct East and West Arctic haplotypes, with admixing of populations in Churchill. Unique Arctic haplotypes (in Northern Alaska, Churchill, Northern Baffin Island, and Labrador) also suggest at possible Arctic contributions. |
| *Devaleraea ramentacea*[1, 2] **(Updated)** (Fig. S13 & S14) | COI-5P: 0/12/27 ITS 0/12/20 | Atlantic (Arctic and Atlantic) | All genetically verified specimens of *Devaleraea* from the North Pacific do not match this species (see Chapter 2); records of *Devaleraea ramentacea* in this flora thus remain uncertain. Haplotype patterns for this species are also consistent with a long history in the Northwest Atlantic (again, consistent with Chapter 2), with specimens sampled in |

| | | | |
|---|---|---|---|
| | | | Northern Labrador matching these populations. A unique Baffin Island and Arctic haplotype is relatively divergent from Atlantic collections, suggesting at possible Arctic contributions in this species. |
| *Dilsea socialis* **(Updated)** (Fig. S15) | 38/50/46 | Pacific | There was no haplotype variation in this species from the Western Arctic through to the Northwest Atlantic. A unique COI-5P haplotype was sampled on St. Lawrence Island in the North Pacific, but haplotype variation is notably absent from Nome, Alaska. It is likely this species has recently migrated out of the Northwest Pacific, which would be consistent with phylogeographic analyses (see Chapter 2); Pacific origins are tentatively inferred, pending further sampling in the Northwest Pacific. |
| *Euthora cristata*[3] **(New)** (Fig. S16) | 30/4/94 | Atlantic (Arctic and Atlantic) | This species is reported in the Northern Pacific and in Northern Alaska (22), though we were unable to genetically verify this species from these locations. Limited Arctic records along Northern Labrador indicate Northwest Atlantic origins in these populations. A unique and highly divergent haplotype from Baffin Island (most closely allied with Pacific populations) suggest at an additional refugial contribution to Arctic populations. |
| *Fimbrifolium dichotomum* **(New)** (Fig. S17) | 0/7/9 | Atlantic | This species is genetically verified from the Northwest Atlantic, matching specimens sampled along Northern Labrador. |
| *Haemescharia polygyna*[3] | *rbc*L: 0/5/0 | Uncertain (Arctic) | This genetic group appears to be restricted to the Eastern Canadian Arctic. |
| *Hildenbrandia* sp. 1Arct[3] **(New)** | 0/4/0 | Uncertain (Arctic) | This genetic group was recovered only in Northern Labrador. |
| *Leptophytum foecundum* **(New)** (Fig. S18) | 0/3/1 | Uncertain | This species has a unique haplotype in the West Arctic, and has been genetically verified in the Northwest Atlantic basin. Given the limited number of samples and difficulty associated with sampling red crusts, the origin of Arctic populations for this species remains unknown. |

| | | | |
|---|---|---|---|
| *Leptosiphonia flexicaulis* **(New)** (Fig. S19) | 0/1/38 | Atlantic | A single Arctic record from Northern Labrador matches Northwest Atlantic populations. |
| *Lithothamnion glaciale* (Fig. S20) | 4/4/44 | Atlantic | Limited North Pacific and Arctic collections suggest this species survived in the Atlantic and migrated Westward through the Arctic. Haplotype variation also suggests this species survived recent glaciation in the Northwest Atlantic. |
| *Lithothamnion lemoineae* **(New)** (Fig. S21) | 1/2/8 | Atlantic | Inferences are problematic in this group given there is a single genetic record from the North Pacific. Arctic records from Baffin Island and Northern Labrador match Northwest Atlantic haplotypes, and while this basin is tentatively listed as the origin for Arctic populations, more sampling is needed in the Western Arctic and North Pacific. |
| *Membranoptera carpophylla*[3] **(New)** | 0/1/0 | Uncertain (Arctic) | A single genetic record exists for this species from Northern Labrador. Phylogeographic analyses indicated this genetic group is nested in a clade of Atlantic species, suggesting it has origins in this basin (1). |
| *Membranoptera fabriciana* **(New)** (Fig. S22) | 0/5/23 | Atlantic | As with *Membranoptera carpophylla*, this species is nested in a clade of Atlantic species, further suggesting Arctic populations have origins in this basin. Indeed, the Arctic haplotype matches Northwest Atlantic populations. |
| *Odonthalia dentata*[2, 3] **(Updated)** (Fig. S23) | 21/48/67 | Atlantic and Pacific (Arctic, Atlantic, and Pacific) | Arctic populations share haplotypes with both Pacific and Northwest Atlantic basins. Haplotype variation also suggests this species has survived glaciation in the North Pacific and Northeast Atlantic, and the Last Glacial Maximum in the Northwest Atlantic (as evidenced by numerous rare private haplotypes). The haplotype patterns suggest at the establishment of the Northwest Atlantic flora from the North Pacific prior to the Last Glacial Maximum (Northeast Atlantic populations must have been established sometime during the Late Pleistocene; 1), followed by the establishment of contemporary Arctic populations from the Pacific and likely from the Atlantic (though more data are needed to confirm an |

| | | | |
|---|---|---|---|
| | | | Atlantic contribution to modern day Arctic flora). Unique rare haplotypes in Northern Alaska and Northern Labrador also suggest at an additional refugial population contributing to Arctic recolonization. |
| *Palmaria palmata* (Fig. S24) | 0/12/68 | Atlantic | Arctic collections are a clear north extension of Northwest Atlantic populations. Though this species is reported from Northern Alaska and the North Pacific, it has not been genetically verified from these regions (these reports likely represent other species of *Palmaria*; 22). |
| *Peyssonnelia rosenvingei* **(New)** (Fig. S25) | 1/7/34 | Uncertain | A single haplotype extends from the Northern Bering Sea to the Northwest Atlantic, suggesting at recent dispersal across the Arctic, through the direction of migration remains uncertain. |
| *Phycodrys fimbriata*[2, 3] **(Updated)** (Fig. S26) | 18/54/69 | Atlantic (Arctic and Atlantic) | A number of unique Arctic haplotypes occur in this species, particularly in Northern Alaska and along the coasts of Baffin Island and Northern Labrador. One Arctic haplotype can be confirmed as Atlantic in origin, while a widely distributed haplotype (occurring in the North Pacific through to Southern Labrador) suggests at recent trans-Arctic dispersal, though the direction of migration remains uncertain. Detailed population level analyses are needed in this species to determine what role, if any, Pacific populations played in Arctic recolonization since the Last Glacial Maximum. |
| *Phymatolithon tenue* **(New)** (Fig. S27) | 0/1/1 | Uncertain | As in *Leptophytum foecundum*, the West Arctic haplotype does not match the Atlantic mitotype, however, there are too few collections to determine the source population in this species. |
| *Polysiphonia* sp. 1stricta (Fig. S28) | 28/8/34 | Uncertain | This genetic group has no COI-5P haplotype variation despite a broad trans-Arctic distribution. This group is another contender for having origins in the Northwest Pacific. More sampling and/or a more variable marker is required; for now, origins of Arctic populations remain uncertain. |

| | | | |
|---|---|---|---|
| *Polysiphonia* sp. 3stricta[1] (Fig. S29) | 0/5/26 | Atlantic (Arctic) | This genetic group has only been confirmed in the Northwest Atlantic, with North Pacific collections attributable to *P*. sp. 1stricta. Phylogeographic analyses also suggest this species has origins in the North Atlantic (1). As such, Arctic populations tentatively have origins in the Northwest Atlantic, however, unique Churchill haplotypes suggest at Arctic refugial contributions in this species. |
| *Ptilota gunneri* (Fig. S30) | 0/1/41 | Atlantic | A rare member of the Northwest Atlantic flora, this species appears to have recent origins in the Northeast Atlantic, with recent migration into the Arctic. |
| *Ptilota serrata* **(New)** (Fig. S31) | 2/11/63 | Uncertain | Though this species is genetically verified from the Bering Sea and the Northeast Pacific (MG762003, MG762004), COI data is restricted to the Atlantic and Arctic basins. As such, the source for Arctic populations remains uncertain, pending COI data from the Pacific. |
| *Rhodochorton purpureum*[3] **(New)** (Fig. S32) | *rbc*L-3P: 0/3/1 | Atlantic (Arctic and Atlantic) | The limited number of genetically verified records suggests this species has recent origins in the Northwest Atlantic. |
| *Rhodomela lycopodioides* (Fig. S33) | 0/5/106 | Atlantic | A single collection from Churchill, matches the COI-5P haplotype of European populations, however, the ITS type for this specimen matches the Northwest Atlantic, indicating admixture between trans-Atlantic populations has occurred. |
| *Rhodomela sibirica*[2] **(Updated)** (Fig. S34 & S35) | COI-5P: 23/48/0 ITS: 22/42/0 | Pacific (Arctic and Pacific) | COI-5P haplotype variation in the North Pacific suggests this species has a long history in the area, with haplotype variation declining towards the Eastern Arctic. COI-5P haplotype variation is highest, however, in Northern Alaska. Similarly, ITS data suggested that specimens from the East Arctic were recolonized out of the North Pacific, while specimens from the West Arctic are distinct from these populations; refugial Arctic contributions are a possibility. |
| *Rhodomela* sp. 1virgata[1, 3] (Fig. S36) | 14/17/7 | Uncertain (Arctic) | Arctic collections do not match North Pacific or Northwest Atlantic haplotypes, with the latter two populations sharing |

| | | | |
|---|---|---|---|
| | | | haplotypes. The location of origin Arctic populations therefore remains uncertain. |
| *Rhodomela virgata* **(Updated)** (Fig. S37) | 37/37/2 | Pacific | Haplotype variation is monotypic through the Arctic and into the Atlantic, while several private haplotypes occur in the North Pacific, suggesting this species recently migrated Eastward through the Arctic. |
| *Rhodophysema kjellmanii*[1] | 0/2/0 | Uncertain (Arctic) | Two records exist for this genetic group from Churchill. As such, the origin of this group remains uncertain. |
| *Rhodophysemopsis hyperborea*[3] **(New)** | 0/1/0 | Uncertain (Arctic) | A single genetic record exists from Northern Labrador. The origin of Arctic populations therefore remains uncertain. |
| *Savoiea arctica*[1, 3] **(Updated)** (Fig. S38) | 0/11/5 | Atlantic (Arctic and Atlantic) | North Pacific collections of *Savoiea* are not attributable to this species (1), suggesting *Savoiea arctica* is limited to the Atlantic basin in its distribution. Given the lack of North Pacific records, Arctic populations in this species, tentatively, are inferred to have origins in the Northwest Atlantic. Two unique Arctic haplotypes are notable in this species, but may be an artifact of the low number of records from the Northwest Atlantic. |
| *Scagelia pylaisaei*[3] (Fig. S39) | 65/24/32 | Pacific (Arctic and Pacific) | A phylogenetic break occurs between Atlantic and Pacific populations, with Arctic collections matching Pacific populations. The Pacific lineage also appears to be admixing with the Atlantic populations along the coast of Labrador. |
| *Turnerella pennyi* **(New)** (Fig. S40) | *rbc*L-3P: 0/9/4 | Atlantic | This species is genetically verified in the Arctic and in the Northwest Atlantic, meaning Arctic populations likely have recent origins out of the Atlantic. |
| *Waernia mirabilis* **(New)** (Fig. S41) | 0/6/4 | Atlantic | Arctic specimens in Northern Labrador match a Northwest Atlantic haplotype. |
| *Wildemania miniata* **(New)** (Fig. S42) | 0/6/45 | Atlantic | As in *Waernia mirabilis*, Arctic specimens match Northwest Atlantic populations. |
| Phaeophyceae | | | |

| | | | |
|---|---|---|---|
| *Agarum clathratum* **(Updated)** (Fig. S43) | 101/9/50 | Pacific | Haplotype variation indicates this species has a relatively long history in the Pacific. Haplotype variation also declines from the Pacific through to the Northwest Atlantic, suggesting at recent Pacific origins. Barring the Northwest Pacific collections, the nearly monotypic haplotype variation extending through the Arctic into the Northwest Atlantic is reminiscent of several species listed here (e.g. *A. borealis*, *Chorda borealis*, *Dilsea socialis*, and *Polysiphonia* sp. 1stricta). |
| *Alaria esculenta*[1, 2, 3] **(Updated)** (Fig. S44 & S45) | COI-5P: 0/17/52 ITS: | Atlantic (Arctic and Atlantic) | A shallow genetic break occurs between Northwest and Northeast Atlantic populations, with the Northeast Atlantic haplotype matching specimens in Northern Labrador. COI-5P data for Bering Sea specimens, however, were attributable to *Alaria crispa.* Nonetheless, ITS data suggest these specimens are *Alaria esculenta* sensu lato, sharing genetic signatures with *A. crispa* and Arctic *A. esculenta*; introgression or incomplete lineage sorting between Nome and Arctic *Alaria* is a possibility. Given the limited sampling in the North Pacific, Arctic haplotypes may have originated from the Pacific. These patterns may also be indicative of an Arctic refugial population. |
| *Ascophyllum nodosum* **(New)** (Fig. S46) | 0/1/2 | Atlantic | Despite being an abundant member of intertidal flora in cold temperate Northwest Atlantic waters, genetic records are limited in this group. This is an Atlantic species, making this the origin for our lone Arctic specimen, collected as drift in Northern Labrador. |
| *Battersia arctica* **(New)** (Fig. S47) | *rbc*L-3P: 1/3/2 | Uncertain | This species has limited records in both the North Pacific and the Northwest Atlantic both matching the Arctic haplotype, rendering assignment of a source for Arctic populations problematic. |
| *Battersia racemosa* **(Updated)** (Fig. S48) | 0/1/1 | Atlantic | A single Arctic record exists from Churchill, matching a Northwest Atlantic record. In addition, this genetic group is verified in the Northeast Atlantic (based on *rbc*L; AJ287880). |

| | | | |
|---|---|---|---|
| | | | Origins for sampled Arctic populations are therefore tentatively inferred as being Atlantic. |
| *Chaetopteris plumosa* [1, 2, 3] **(New)** (Fig. S49) | 0/17/19 | Atlantic (Arctic and Atlantic) | A genetic break occurs between the two haplotypes sampled, both of which occur in the Arctic. One haplotype is attributable to the Atlantic, while the other may originate from the Pacific, though more sampling is needed to confirm this. Haplotype patterns may also indicate origins in Arctic refugia. Interpretation of these patterns is tentative, pending further study. |
| *Chorda borealis* **(Updated)** (Fig. S50) | 23/14/2 | Uncertain | Haplotype variation is monotypic in this genetic group, extending from the North Pacific to the Northwest Atlantic (Makkovik, Newfoundland); thus, the hypothesized Pacific origins inferred by Saunders & McDevit (11) are, at present, not supported by the additional collections available here. |
| *Chordaria chordaeformis*[1] **(Updated)** (Fig. S51) | 32/21/1 | Uncertain (Arctic) | Despite extensive sampling, COI-5P is nearly monotypic, except for a single private haplotype recovered in Churchill, lending uncertainty and possible Arctic periglacial origins to this species. |
| *Chordaria flagelliformis*[1, 3] (Fig. S52) | 0/48/56 | Atlantic (Arctic and Atlantic) | Haplotype variation in this species suggests at a long history in the Northwest Atlantic. Despite unique haplotypes in the Arctic collections, the lack of North Pacific records matching this species (which were attributable to a closely related genetic group) suggests Arctic populations originated from the Atlantic basin. This species is also genetically verified in Northern Europe, including Svalbard, based on *rbc*L data (AB066076, AB066073, AB066075, JN599169). |
| Chordariacean sp. 4nov | 0/1/1 | Atlantic | With only two records, this genetic group is tentatively hypothesized to have originated from the Atlantic. |
| *Cladosiphon* sp. 1NFLD[3] **(New)** | 0/1/0 | Uncertain (Arctic) | A single collection for this genetic group occurs in Northern Labrador. |
| *Desmarestia* sp. 1aculeata (Fig. S53) | 0/13/30 | Atlantic | Specimens of *Desmarestia* in the Northern Bering Sea were attributable to *Desmarestia* sp. 2aculeata and *Desmarestia viridis*. This genetic group, on the other hand, has only been |

| | | | |
|---|---|---|---|
| | | | recovered in the Atlantic, making this the source for Arctic populations. |
| *Dictyosiphon* sp. 1GWS (Fig. S54) | 40/1/1 | Pacific | The single Arctic record from Churchill, matches a Pacific haplotype, while a single Atlantic collection appears to be divergent from Pacific collections. This genetic group is also verified from Russia based on *rbc*L data (AY372973). |
| *Dictyosiphon* sp. 3GWS[1] (Fig. S55) | 3/1/9 | Uncertain (Arctic) | The single Arctic record from Churchill is quite divergent from Pacific and Atlantic populations, possibly representing its own species. |
| *Dictyosiphon foeniculaceus*[3] (Fig. S56) | 0/4/47 | Atlantic (Arctic and Atlantic) | Haplotype variation in the Northwest Atlantic suggests this species has survived multiple glaciations in the area, with records of *Dictyosiphon* from the North Pacific attributable to *Dictyosiphon* sp. 1GWS. Arctic haplotypes match Atlantic populations. |
| Ectocarpoid sp. **(New)** | 0/2/0 | Uncertain (Arctic) | This genetic group has been sampled from Northern Labrador (our study) and from Baffin Island (LT546267; 15). |
| *Ectocarpus* sp. 1siliculosus **(New)** (Fig. S57) | 0/1/10 | Atlantic | A single Arctic collection from Churchill matches a Northwest Atlantic haplotype, the only area this genetic group has been recovered from. |
| *Ectocarpus* sp. **(New)** | 0/2/0 | Uncertain (Arctic) | As above, this genetic group was sampled in our study, in Northern Labrador, and by Küpper *et al.* (15) on Baffin Island (LT546288). |
| *Eudesme borealis* **(New)** (Fig. S58) | 18/11/14 | Atlantic and Pacific | Haplotypes sampled in Northern Labrador match Atlantic and Pacific records, indicating admixture between these populations in the Arctic. |
| *Eudesme* sp.[1] **(New)** | 0/9/0 | Uncertain (Arctic) | A new genetic group was recently sampled in Churchill (MB). |
| *Fucus distichus*[1] (Fig. S59) **(Updated)** | 73/12/34 | Atlantic (Arctic) | One Arctic haplotype recovered in this matches Atlantic populations, while another is widespread between the Atlantic and Pacific basins; Atlantic origins are therefore tentatively inferred in this species with further population level work needed. Recent work also suggests this species survived in |

| | | | |
|---|---|---|---|
| | | | Arctic refugial populations, seeding the Atlantic and Pacific out of the Arctic (14). |
| *Halosiphon* sp. 2tomentosus (Fig. S60) | 2/1/0 | Pacific | PCR success was low for Arctic *Halosiphon* suggesting primer issues. Given the lack of Atlantic records, Arctic populations in this species are, tentatively, of Pacific origin, though an effort should be made to generate sequence data in previously failed PCRs for *Halosiphon*, particularly from the Northwest Atlantic. This genetic group, however, is previously reported from the North Pacific, with *rbc*L data for Arctic collections matching Pacific rather than Atlantic collections for this species (11). |
| *Halothrix lumbricalis*[3] **(New)** (Fig. S61) | 0/2/1 | Atlantic (Arctic and Atlantic) | Two Arctic specimens were sampled from Northern Labrador, one of which matched a Northwest Atlantic haplotype, while the other was unique. This group is genetically verified from Greenland (published as *Elachista fucicola*; AF055398). |
| *Haplospora globosa* **(Updated)** (Fig. S62) | 1/1/6 | Uncertain | A single Arctic collection matches a haplotype sampled in both the Northeast Atlantic and the North Pacific, meaning a source for Arctic populations cannot be inferred. |
| *Hedophyllum nigripes*[3] (Fig. S63) | 20/18/10 | Atlantic (Arctic and Atlantic) | Arctic records match an Atlantic haplotype. |
| *Heterosaundersella* sp. 1NFLD[3] | 0/1/0 | Uncertain (Arctic) | A single record for this genetic group was recovered from Northern Labrador. |
| *Laminaria digitata* (Fig. S64) | 0/3/63 | Atlantic | The source for Arctic populations is hypothesized to be the Atlantic, where this species is believed to have evolved (2). |
| *Laminaria solidungula*[2, 3] **(Updated)** (Fig. S65) | 0/19/1 | Atlantic (Arctic and Atlantic) | East and West Arctic specimens represent different mitotypes, with the East Arctic haplotype matching a Northwest Atlantic record. The inference of Pacific contributions to Arctic recolonization awaits genetic confirmation of this species in the Pacific. |
| *Leptonematella fasciculata*[1] | 0/3/0 | Uncertain (Arctic) | The few records that exist for this genetic group occur in Churchill, meaning a source population cannot be inferred. |

| | | | |
|---|---|---|---|
| *Lithoderma* sp. 2GWS (Fig. S66) | *rbc*L-3P: 0/1/1 | Uncertain | Records of this species are from Northern Alaska and the Northwest Atlantic, however, given the disjunct sampling distribution and limited number of records, the source basin for this species remains uncertain. |
| *Petalonia fascia*[1,3] (Fig. S67) | 47/23/43 | Uncertain (Arctic) | One Arctic haplotype matches Atlantic collections, but a complex haplotype network in this species extends throughout the Atlantic and Pacific. A unique haplotype occurs in Churchill, sharing a substitution site with a Pacific haplotype. In addition, this species is genetically verified in the Northwest Pacific (based on ITS and *rbc*L data; AY154725, AB578997), and from the Northeast Atlantic (based on PSA and *rbc*L data; AY372953, AB860190, AB860189), which are not included in our haplotype map/network. In sum, it remains unclear where Arctic populations originated from. |
| *Petalonia filiformis*[3] **(Updated)** (Fig. S68) | 0/25/19 | Atlantic (Arctic and Atlantic) | Genetic variation along the coast of Labrador and the absence of specimens attributable to *Petalonia filiformis* from the North Pacific, suggest this species survived the Last Glacial Maximum in the Northwest Atlantic and has subsequently moved into the Canadian Arctic. |
| *Planosiphon complanatus* **(New)** (Fig. S69) | 0/1/7 | Atlantic | A single Arctic specimen from Baffin Island matches Northwest Atlantic collections. |
| *Planosiphon zosterifolius*[3] **(New)** (Fig. S70) | 8/2/7 | Uncertain (Arctic) | Two Arctic collections (Churchill and Northern Labrador) do not match limited collections from the Northern Bering Sea and the North Atlantic. The origin of Arctic populations therefore remains uncertain at this time. |
| *Platysiphon glacialis* **(New)** (Fig. S71) | 16/3/0 | Pacific | This species previously had genetically verified records only from Northern Baffin Island (22). The presence of this species in the North Pacific suggests this is the source for Arctic populations. |
| *Punctaria* sp. 2GWS (Fig. S72) | 2/5/5 | Uncertain | Limited collections and a lack of haplotypes in this group impedes further consideration of the source for Arctic populations. In addition, this group is genetically verified in |

| | | | |
|---|---|---|---|
| | | | Greenland (AF055410) and Japan (AB302316) based on *rbc*L data. |
| *Pylaiella littoralis* (Fig. S73) | 0/8/23 | Atlantic | Haplotype variation suggests this species has survived multiple glaciations on both sides of the North Atlantic, with a single Arctic haplotype matching Northwest Atlantic variation. Specimens in the North Pacific are attributable to other genetics groups within *Pylaiella* (11). |
| *Pylaiella washingtoniensis*[1, 3] **(Updated)** (Fig. S74) | 32/39/29 | Atlantic and Pacific (Arctic, Atlantic, and Pacific) | A haplotype recovered in the West Arctic matches Pacific populations, while the haplotypes in Churchill are either unique or of Atlantic origin. Baffin Island and Northern Labrador similarly display unique haplotypes despite reasonably extensive sampling in the Atlantic and Pacific basins. |
| *Ralfsia fungiformis* **(New)** (Fig. S75) | 13/1(*rbc*L)/14 | Uncertain | This species is genetically verified from Baffin Island based on *rbc*L data, however, we were unable to generate COI-5P data to include in the haplotype map/network. Given Atlantic and Pacific populations appear to be divergent, COI-5P data from Arctic collections will likely be informative as to recent origins in these populations. |
| *Saccharina latissima*[1] (Fig. S76) | 14/50/119 | Atlantic and Pacific (Arctic, Atlantic, and Pacific) | Atlantic and Pacific lineages are in secondary contact in the Arctic, a finding consistent with ITS data (3). |
| *Saccorhiza dermatodea* **(New)** (Fig. S77) | 0/1/19 | Atlantic | A single Arctic record from Baffin Island matches a Northwest Atlantic haplotype. |
| *Scytosiphon* sp. 1crust[1] | 0/1/0 | Uncertain (Arctic) | A single record exists for this genetic group from Churchill. |
| *Scytosiphon canaliculatus* (Fig. S78) | 80/5/64 | Uncertain | The two Arctic haplotypes sampled in this species are trans-Arctic, extending from the Northwest Pacific to the Northwest Atlantic, meaning the origin of Arctic populations remains uncertain at this time. |

| | | | |
|---|---|---|---|
| *Scytosiphon* sp. GroupJ[3] **(New)** (Fig. S79) | 0/1/9 | Uncertain (Arctic) | This genetic group appears to have a disjunct distribution, with collections limited to Baffin Island and the New England States, with the single Arctic collection representing a unique haplotype. The origin of Arctic populations therefore remains uncertain at this time. |
| *Stictyosiphon soriferus*[1] | 0/1/0 | Uncertain (Arctic) | A single genetic record exists for this genetic group, occurring in Churchill. |
| *Stictysiphon tortilis*[1] | 0/11/0 | Uncertain (Arctic) | Genetic records for this species are limited to Cambridge Bay and Churchill. |
| Tilopteridalean sp. 1GWS[3] **(New)** (Fig. S80) | 0/2/2 | Atlantic (Arctic and Atlantic) | Two Arctic collections are from Northern Labrador. One matches North Atlantic collections, while the other represents a unique haplotype; the origin of Arctic populations is therefore tentatively Atlantic, though more collections are needed in this genetic group. |
| Tilopteridalean sp. 2GWS[2, 3] (Fig. S81) | 0/4/0 | Uncertain (Arctic) | This genetic group has only been collected in the Arctic, meaning recent origins remain uncertain. |
| Tilopteridalean sp. 3GWS[2] | 0/2/0 | Uncertain (Arctic) | Only two records exist for this genetic group, both from Northern Alaska. |
| *Ulvophyceae* | | | |
| *Acrosiphoni*a sp. 3GWS[1] (Fig. S82) | *tufA*: 0/2/3 | Atlantic (Arctic, Atlantic) | Few genetic records exist for this genetic group, however, the origin of Arctic populations is tentatively assigned to the Atlantic. |
| *Acrosiphonia* sp. 6GWS **(Updated)** (Fig. S83) | *tufA*: 0/1/2 | Atlantic | The same scenario occurs in this species as in *Acrosiphonia* sp. 3GWS. |
| *Acrosiphonia* sp. 8GWS **(New)** (Fig. S84) | *tufA*: 5/1/1 | Atlantic | This genetic group occurs in all three oceans, however, the single Arctic collection from Baffin Island matches a single Northwest Atlantic record. |
| *Acrosiphonia sonderi* **(New)** (Fig. S85) | *tufA*: 2/2/11 | Uncertain | A single haplotype extends from the Northwest Pacific to the Northwest Atlantic, meaning the origin of Arctic populations remains uncertain at this time. |
| *Blidingia* sp. 3GWS **(New)** | *tufA*: 0/4/4 | Atlantic | This genetic group was only recovered from the Arctic and Atlantic, making the latter the putative source population. |

| | | | |
|---|---|---|---|
| (Fig. S86) | | | |
| *Blidingia* sp. 5GWS[1, 2] (Fig. S87) | *tufA*: 0/2/0 | Uncertain (Arctic) | Two genetic records exist for this genetic group, both occurring in the Arctic. |
| *Monostroma* sp. 2grevillei[3] **(New)** (Fig. S88) | *tufA*: 0/1/40 | Uncertain (Arctic) | A single Arctic record for this genetic group does not match the single North Atlantic haplotype, making the origin of Arctic populations uncertain at this time. |
| *Spongomorpha aeruginosa* (Fig. S89) | *tufA*: 0/1/9 | Atlantic | This species lacks North Pacific records and is not reported from the Pacific. As such, the source for Arctic populations is inferred to be the Atlantic. |
| *Rosenvingiella* sp.[3] **(New)** | *tufA*: 0/1/0 | Uncertain (Arctic) | A single Arctic collection for this genetic group exists from Northern Labrador. |
| *Ulothrix flacca* (Fig. S90) | *rbc*L-3P: 0/1/4 | Atlantic | We did not recover this species while sampling the North Pacific, however, it is reported from the area (22). The origin of Arctic populations is therefore tentatively Atlantic. |
| Ulotrichales spp. | *tufA*: 0/3/0 | Uncertain (Arctic) | Two new genetic groups (possibly three) with uncertain assignments occur in Churchill. |
| *Ulva fenestrata* (Fig. S91) | *tufA*: 69/14/97 | Pacific | North Pacific populations match the Arctic haplotype recovered in Churchill, corroborating the inference of Pacific origins made by Saunders and McDevit (11; previously reported as *Ulva lactuca*). This species is also confirmed in Japan (based on *rbc*L; AB097622). |
| *Ulva intestinalis* **(New)** (Fig. S92) | *tufA*: 17/1/65 | Uncertain | A single Churchill collection matches a widespread haplotype, leaving the origin of the Arctic specimen uncertain. |
| *Ulva* sp. 3linza **(New)** (Fig. S93) | *tufA*: 14/5/16 | Pacific | One Arctic haplotype is assignable to Pacific populations, while another widespread haplotype (also in the Arctic) has uncertain origins. Declining haplotype diversity eastwards into the North Atlantic suggest at an entirely Pacific origin for Arctic and North Atlantic populations. |
| *Ulva obscura*[3] **(New)** (Fig. S94) | *tufA*: 20/2/74 | Atlantic (Arctic and Atlantic) | Pacific and Atlantic haplotypes are distinct in this species, with one Arctic specimen matching the latter. The second Arctic specimen represents a unique haplotype but allies most closely with Pacific collections. |

| | | | |
|---|---|---|---|
| *Ulva prolifera*[2] (Fig. S95) | *tufA*: 0/10/16 | Atlantic (Arctic and Atlantic) | East Arctic haplotypes match Atlantic populations, while the West Arctic haplotype differs from Atlantic populations by three substitutions, sharing signatures with a closely related genetic group (*Ulva* sp. 2prolifera) sampled in Northern British Columbia (though this specimen differs by another five substitutions from the West Arctic haplotype). This species is also confirmed in the Northwest Pacific (based on *rbc*L; KP233770). Given species delineations are not clear in this group, Atlantic origins for Arctic populations are tentatively inferred, but further sampling in the Pacific is likely to revise this scenario. |
| *Ulva* sp. | *tufA*: 0/11/0 | Uncertain (Arctic) | A new genetic group most closely matching *Ulva intestinalis* (97%) occurs in Churchill. |

**Table S2**. Kruskal-Wallis test results with Dunn's post hoc tests with Bonferroni corrections for groups wherein the null hypothesis was rejected. NE=Northeast, NW=Northwest.

| Variable | Total N | Test Statistic | Degrees of freedom | Asymptotic Sig. (2-sided test) | |
|---|---|---|---|---|---|
| Tajima's D | 73 | 14.554 | 5 | .012 | |
| Sample 1-Sample 2 | Test Statistic | Std. Error | Std. Test Statistic | Sig. | Adj. Sig. |
| NE Pacific-NW Atlantic | -3.807 | 9.936 | -.383 | .702 | 1.000 |
| NE Pacific-NE Atlantic | 12.133 | 12.848 | .944 | .345 | 1.000 |
| NE Pacific-East Arctic | 21.920 | 9.726 | 2.254 | .024 | .363 |
| NE Pacific-Nome | -23.433 | 10.249 | -2.286 | .022 | .333 |
| NE Pacific-Beaufort | 26.133 | 12.848 | 2.034 | .042 | .629 |
| NW Atlantic-NE Atlantic | 8.326 | 10.664 | .781 | .435 | 1.000 |
| NW Atlantic-East Arctic | 18.113 | 6.578 | 2.754 | .006 | .088 |
| NW Atlantic-Nome | -19.626 | 7.328 | -2.678 | .007 | .111 |
| NW Atlantic-Beaufort | 22.326 | 10.664 | 2.094 | .036 | .544 |
| NE Atlantic-East Arctic | 9.787 | 10.469 | .935 | .350 | 1.000 |
| NE Atlantic-Nome | -11.300 | 10.956 | -1.031 | .302 | 1.000 |
| NE Atlantic-Beaufort | 14.000 | 13.419 | 1.043 | .297 | 1.000 |
| East Arctic-Nome | -1.513 | 7.042 | -.215 | .830 | 1.000 |
| East Arctic-Beaufort | 4.213 | 10.469 | .402 | .687 | 1.000 |
| Nome-Beaufort | 2.700 | 10.956 | .246 | .805 | 1.000 |
| Variable | Total N | Test Statistic | Degrees of freedom | Asymptotic Sig. (2-sided test) | |
| Number of private haplotypes (NPH) | 119 | 11.989 | 5 | .035 | |
| Sample 1-Sample 2 | Test Statistic | Std. Error | Std. Test Statistic | Sig. | Adj. Sig. |
| Beaufort-Nome | -11.012 | 11.147 | -.988 | .323 | 1.000 |
| Beaufort-East Arctic | -13.321 | 10.231 | -1.302 | .193 | 1.000 |
| Beaufort-NE Atlantic | -15.216 | 12.410 | -1.226 | .220 | 1.000 |
| Beaufort-NW Atlantic | -30.011 | 10.350 | -2.900 | .004 | .056 |
| Beaufort-NE Pacific | -32.912 | 13.398 | -2.457 | .014 | .210 |
| Nome-East Arctic | 2.308 | 9.532 | .242 | .809 | 1.000 |
| Nome-NE Atlantic | 4.204 | 11.840 | .355 | .723 | 1.000 |
| Nome-NW Atlantic | 18.998 | 9.660 | 1.967 | .049 | .738 |
| Nome-NE Pacific | 21.900 | 12.872 | 1.701 | .089 | 1.000 |

| | | | | | |
|---|---|---|---|---|---|
| East Arctic-NE Atlantic | -1.896 | 10.982 | -.173 | .863 | 1.000 |
| East Arctic-NW Atlantic | -16.690 | 8.586 | -1.944 | .052 | .779 |
| East Arctic-NE Pacific | -19.592 | 12.087 | -1.621 | .105 | 1.000 |
| NE Atlantic-NW Atlantic | -14.794 | 11.093 | -1.334 | .182 | 1.000 |
| NE Atlantic-NE Pacific | -17.696 | 13.980 | -1.266 | .206 | 1.000 |
| NW Atlantic-NE Pacific | 2.902 | 12.188 | .238 | .812 | 1.000 |

| Variable | Total N | Test Statistic | Degrees of freedom | Asymptotic Sig. (2-sided test) |
|---|---|---|---|---|
| Gene diversity (h) | 112 | 8.242 | 5 | .143 |

| Variable | Total N | Test Statistic | Degrees of freedom | Asymptotic Sig. (2-sided test) |
|---|---|---|---|---|
| Number of effective alleles (Ne) | 112 | 7.986 | 5 | .157 |

| Variable | Total N | Test Statistic | Degrees of freedom | Asymptotic Sig. (2-sided test) |
|---|---|---|---|---|
| Pi | 112 | 9.439 | 5 | .093 |

| Variable | Total N | Test Statistic | Degrees of freedom | Asymptotic Sig. (2-sided test) |
|---|---|---|---|---|
| Number of haplotypes (Na) | 112 | 11.591 | 5 | .041 |

| Sample 1-Sample 2 | Test Statistic | Std. Error | Std. Test Statistic | Sig. | Adj. Sig. |
|---|---|---|---|---|---|
| Beaufort-NE Atlantic | -2.197 | 12.557 | -.175 | .861 | 1.000 |
| Beaufort-Nome | -11.808 | 10.805 | -1.093 | .274 | 1.000 |
| Beaufort-East Arctic | -19.011 | 9.950 | -1.911 | .056 | .841 |
| Beaufort-NE Pacific | -26.958 | 13.849 | -1.947 | .052 | .774 |
| Beaufort-NW Atlantic | -28.296 | 10.187 | -2.778 | .005 | .082 |
| NE Atlantic-Nome | -9.611 | 11.875 | -.809 | .418 | 1.000 |
| NE Atlantic-East Arctic | 16.814 | 11.102 | 1.514 | .130 | 1.000 |
| NE Atlantic-NE Pacific | -24.761 | 14.699 | -1.685 | .092 | 1.000 |
| NE Atlantic-NW Atlantic | -26.099 | 11.315 | -2.307 | .021 | .316 |
| Nome-East Arctic | 7.202 | 9.073 | .794 | .427 | 1.000 |
| Nome-NE Pacific | 15.150 | 13.233 | 1.145 | .252 | 1.000 |
| Nome-NW Atlantic | 16.488 | 9.333 | 1.767 | .077 | 1.000 |
| East Arctic-NE Pacific | -7.948 | 12.545 | -.634 | .526 | 1.000 |
| East Arctic-NW Atlantic | -9.286 | 8.327 | -1.115 | .265 | 1.000 |

| | | | | | |
|---|---|---|---|---|---|
| NE Pacific-NW Atlantic | -1.338 | 12.734 | -.105 | .916 | 1.000 |
| Variable | Total N | Test Statistic | Degrees of freedom | Asymptotic Sig. (2-sided test) | |
| Number of polymorphic sites (Npoly) | 112 | 13.244 | 5 | .021 | |
| Sample 1-Sample 2 | Test Statistic | Std. Error | Std. Test Statistic | Sig. | Adj. Sig. |
| Beaufort-NE Atlantic | -9.648 | 12.601 | -.766 | .444 | 1.000 |
| Beaufort-Nome | -12.267 | 10.842 | -1.131 | .258 | 1.000 |
| Beaufort-East Arctic | -27.322 | 9.984 | -2.737 | .006 | .093 |
| Beaufort-NW Atlantic | -29.022 | 10.222 | -2.839 | .005 | .068 |
| Beaufort-NE Pacific | -31.779 | 13.897 | -2.287 | .022 | .333 |
| NE Atlantic-Nome | -2.618 | 11.916 | -.220 | .826 | 1.000 |
| NE Atlantic-East Arctic | 17.673 | 11.140 | 1.586 | .113 | 1.000 |
| NE Atlantic-NW Atlantic | -19.374 | 11.354 | -1.706 | .088 | 1.000 |
| NE Atlantic-NE Pacific | -22.131 | 14.750 | -1.500 | .134 | 1.000 |
| Nome-East Arctic | 15.055 | 9.104 | 1.654 | .098 | 1.000 |
| Nome-NW Atlantic | 16.756 | 9.365 | 1.789 | .074 | 1.000 |
| Nome-NE Pacific | 19.513 | 13.279 | 1.469 | .142 | 1.000 |
| East Arctic-NW Atlantic | -1.701 | 8.356 | -.204 | .839 | 1.000 |
| East Arctic-NE Pacific | -4.458 | 12.588 | -.354 | .723 | 1.000 |
| NW Atlantic-NE Pacific | 2.757 | 12.778 | .216 | .829 | 1.000 |

**Table S3:** Summary statistics for populations of Arctic marine macroalgae with Arctic populations. *n* = sample size, bp = number of basepairs, $N_{poly}$ = number of polymorphic nucleotide sites, Na = number of haplotypes, Ne = number of effective alleles, $N_{PH}$ = number of private haplotypes, *h* = haplotype diversity, $\theta_\pi$ = nucleotide diversity (Standard Deviation), *D* = Tajima's test for neutralilty, *p*= Tajima's D test probability.

| Location | *n* | bp | $N_{poly}$ | Na | Ne | $N_{PH}$ | *h* | $\theta_\pi$ | *D* | *p* |
|---|---|---|---|---|---|---|---|---|---|---|
| ***Ahnfeltia borealis* COI-5P** | | | | | | | | | | |
| **NE Pacific** | 1 | 638 | - | - | - | 0 | - | - | - | - |
| **Nome** | 24 | 638 | 0 | 1 | 1 | 0 | 0 | 0.0000 | - | - |
| **Beaufort** | 23 | 638 | 0 | 1 | 1 | 0 | 0 | 0.0000 | - | - |
| **East Arctic** | 21 | 638 | 0 | 1 | 1 | 0 | 0 | 0.0000 | - | - |
| **NW Atlantic** | 2 | 638 | 10 | 2 | 2 | 2 | 0.5 | 0.01567 | - | - |
| **Overall** | 71 | 638 | 10 | 3 | 1.058 | - | 0.055 | 0.00044 | **-2.32662** | **<0.01** |
| ***Ahnfeltia borealis ycf35*** | | | | | | | | | | |
| **NE Pacific** | 1 | 915 | - | - | - | 1 | - | - | - | - |
| **Nome** | 23 | 915 | 0 | 2 | 2 | 1 | 0.423 | 0.0000 | - | - |
| **Beaufort** | 20 | 915 | 0 | 2 | 1 | 0 | 0.000 | 0.0000 | - | - |
| **East Arctic** | 15 | 915 | 1 | 2 | 2 | 1 | 0.124 | 0.00015 | -1.15945 | >0.10 |
| **NW Atlantic** | 2 | 915 | 0 | 1 | 1 | 1 | 0.000 | 0.0000 | - | - |
| **Overall** | 61 | 915 | 4 | 5 | 1.456 | - | 0.313 | 0.00031 | -1.4440 | >0.10 |
| ***Coccotylus truncatus* COI-5P** | | | | | | | | | | |
| **Nome** | 12 | 660 | 3 | 3 | 1.674 | 2 | 0.403 | 0.00117 | -0.72873 | >0.10 |
| **Beaufort** | 30 | 660 | 3 | 3 | 2.261 | 1 | 0.558 | 0.00175 | 1.23301 | >0.10 |
| **East Arctic** | 34 | 660 | 8 | 4 | 2.050 | 3 | 0.512 | 0.00134 | -1.62142 | >0.05 |
| **NW Atlantic** | 54 | 660 | 7 | 7 | 2.292 | 5 | 0.564 | 0.00173 | -0.66903 | >0.10 |
| **NE Atlantic** | 1 | 660 | - | - | - | 1 | - | - | - | - |
| **Overall** | 131 | 660 | 18 | 14 | 3.629 | - | 0.724 | 0.00240 | -1.43473 | >0.10 |
| ***Coccotylus truncatus* ITS** | | | | | | | | | | |
| **Nome** | 11 | 672 | 1 | 2 | 1.198 | 0 | 0.165 | 0.00026 | -0.64112 | >0.10 |
| **Beaufort** | 30 | 672 | 3 | 4 | 1.525 | 2 | 0.344 | 0.00056 | -0.8272 | >0.10 |
| **East Arctic** | 25 | 672 | 3 | 4 | 2.510 | 0 | 0.604 | 0.00127 | 0.57304 | >0.10 |
| **NW Atlantic** | 35 | 672 | 4 | 5 | 3.551 | 2 | 0.718 | 0.00155 | 0.54407 | >0.10 |
| **Overall** | 101 | 672 | 7 | 8 | 4.879 | - | 0.795 | 0.00209 | 0.38203 | >0.10 |
| ***Devaleraea ramentacea* COI-5P** | | | | | | | | | | |
| **East Arctic** | 12 | 603 | 4 | 3 | 2.182 | 2 | 0.542 | 0.00291 | 1.14977 | >0.10 |
| **NW Atlantic** | 27 | 603 | 7 | 5 | 1.609 | 4 | 0.379 | 0.00142 | -1.60503 | >0.05 |
| **Overall** | 39 | 603 | 10 | 7 | 2.214 | - | 0.548 | 0.00249 | -1.09565 | >0.10 |
| ***Devaleraea ramentacea* ITS** | | | | | | | | | | |
| **Location** | *n* | bp | $N_{poly}$ | Na | Ne | $N_{PH}$ | *h* | $\theta_\pi$ | *D* | *p* |

| | | | | | | | | | | |
|---|---|---|---|---|---|---|---|---|---|---|
| **East Arctic** | 12 | 940 | 15 | 7 | 4.235 | 5 | 0.764 | 0.00754 | **2.63065** | **<0.01** |
| **NW Atlantic** | 20 | 940 | 3 | 8 | 5.128 | 6 | 0.847 | 0.00091 | 0.43931 | >0.10 |
| **Overall** | 32 | 940 | 16 | 13 | 7.111 | - | 0.859 | 0.00443 | 0.43483 | >0.10 |
| ***Dilsea socialis* COI-5P** | | | | | | | | | | |
| **Nome** | 38 | 645 | 1 | 2 | 1.054 | 1 | 0.051 | 0.00008 | -1.12863 | >0.10 |
| **Beaufort** | 28 | 645 | 0 | 1 | 1.000 | 0 | 0.000 | 0.000 | - | - |
| **East Arctic** | 21 | 645 | 0 | 1 | 1.000 | 0 | 0.000 | 0.000 | - | - |
| **NW Atlantic** | 47 | 645 | 0 | 1 | 1.000 | 0 | 0.000 | 0.000 | - | - |
| **Overall** | 134 | 645 | 1 | 2 | 1.015 | - | 0.015 | 0.00002 | -0.99538 | >0.10 |
| ***Odonthalia dentata* COI-5P** | | | | | | | | | | |
| **Nome** | 21 | 632 | 3 | 4 | 2.609 | 2 | 0.617 | 0.00179 | 0.94342 | >0.10 |
| **Beaufort** | 24 | 632 | 3 | 4 | 2.215 | 1 | 0.549 | 0.00123 | -0.08767 | >0.10 |
| **East Arctic** | 24 | 632 | 3 | 4 | 1.823 | 2 | 0.451 | 0.00081 | -0.91578 | >0.10 |
| **NW Atlantic** | 41 | 632 | 3 | 4 | 1.223 | 2 | 0.182 | 0.00030 | -1.57109 | >0.10 |
| **NE Atlantic** | 26 | 632 | 0 | 1 | 1.000 | 1 | 0.000 | 0.0000 | - | - |
| **Overall** | 136 | 632 | 14 | 11 | 4.346 | - | 0.770 | 0.00425 | -0.04925 | >0.10 |
| ***Palmaria palmata* COI-5P** | | | | | | | | | | |
| **East Arctic** | 12 | 591 | 0 | 1 | 1.000 | 0 | 0.000 | 0.000 | - | - |
| **NW Atlantic** | 48 | 591 | 9 | 9 | 1.580 | 8 | 0.367 | 0.00077 | **-2.18796** | **<0.01** |
| **NE Atlantic** | 19 | 591 | 8 | 9 | 3.800 | 9 | 0.737 | 0.00190 | -1.74253 | >0.05 |
| **Overall** | 79 | 591 | 23 | 18 | 2.392 | - | 0.582 | 0.00525 | -1.01224 | >0.10 |
| ***Phycodrys fimbriata* COI-5P** | | | | | | | | | | |
| **Nome** | 18 | 634 | 1 | 2 | 1.800 | 1 | 0.444 | 0.00074 | 1.16615 | >0.10 |
| **Beaufort** | 32 | 634 | 2 | 3 | 1.290 | 2 | 0.225 | 0.00038 | -1.04684 | >0.10 |
| **East Arctic** | 22 | 634 | 8 | 5 | 1.635 | 3 | 0.388 | 0.00238 | -1.02928 | >0.10 |
| **NW Atlantic** | 69 | 634 | 10 | 10 | 3.843 | 8 | 0.740 | 0.00218 | -0.90722 | >0.10 |
| **Overall** | 141 | 634 | 15 | 16 | 4.174 | - | 0.760 | 0.00406 | -0.14046 | >0.10 |
| ***Rhodomela sibirica* COI-5P** | | | | | | | | | | |
| **Nome** | 23 | 646 | 2 | 3 | 2.159 | 1 | 0.537 | 0.00094 | 0.27506 | >0.10 |
| **Beaufort** | 29 | 646 | 4 | 4 | 2.396 | 2 | 0.538 | 0.00206 | 0.79250 | >0.10 |
| **East Arctic** | 19 | 646 | 0 | 1 | 1.000 | 0 | 0.000 | 0.000 | - | - |
| **Overall** | 71 | 646 | 4 | 5 | 2.882 | - | 0.653 | 0.00158 | 0.49806 | >0.10 |
| ***Rhodomela sibirica* ITS** | | | | | | | | | | |
| **Nome** | 22 | 594 | 1 | 4 | 2.444 | 2 | 0.591 | 0.00069 | 1.06453 | >0.10 |
| **Beaufort** | 27 | 594 | 0 | 1 | 1.000 | 1 | 0.000 | 0.00000 | - | - |
| **East Arctic** | 15 | 594 | 1 | 2 | 1.471 | 0 | 0.320 | 0.00031 | -0.40885 | >0.10 |
| **Overall** | 64 | 594 | 2 | 5 | 3.501 | - | 0.714 | 0.00155 | **2.19262** | **<0.05** |
| ***Rhodomela virgata* COI-5P** | | | | | | | | | | |
| **Location** | *n* | bp | $N_{poly}$ | Na | Ne | $N_{PH}$ | *h* | $\theta_{\pi}$ | *D* | *p* |

| | | | | | | | | | | |
|---|---|---|---|---|---|---|---|---|---|---|
| **Nome** | 37 | 560 | 4 | 5 | 1.253 | 4 | 0.492 | 0.00039 | **-1.88391** | **<0.05** |
| **Beaufort** | 1 | 560 | - | - | - | 0 | - | - | - | - |
| **East Arctic** | 36 | 560 | 0 | 1 | 1.000 | 0 | 0.000 | 0.00000 | - | - |
| **NW Atlantic** | 2 | 560 | 0 | 1 | 1.000 | 0 | 0.000 | 0.00000 | - | - |
| **Overall** | 76 | 560 | 4 | 5 | 1.113 | - | 0.102 | 0.00019 | **-1.81504** | **<0.05** |
| ***Rhodomela* sp. 1virgata COI-5P** | | | | | | | | | | |
| **Nome** | 14 | 638 | 1 | 2 | 1.153 | 1 | 0.133 | 0.00022 | -1.15524 | >0.10 |
| **East Arctic** | 17 | 638 | 5 | 6 | 3.400 | 5 | 0.706 | 0.00205 | -0.37121 | >0.10 |
| **NW Atlantic** | 7 | 638 | 1 | 2 | 1.324 | 1 | 0.245 | 0.00045 | -1.00623 | >0.10 |
| **Overall** | 38 | 638 | 6 | 8 | 2.971 | - | 0.663 | 0.00159 | -0.78481 | >0.10 |
| ***Scagelia pylaisaei* COI-5P** | | | | | | | | | | |
| **NE Pacific** | 29 | 617 | 10 | 10 | 3.805 | 8 | 0.737 | 0.00182 | -1.78359 | >0.05 |
| **Nome** | 36 | 617 | 1 | 2 | 1.117 | 1 | 0.105 | 0.00017 | -0.81338 | >0.10 |
| **East Arctic** | 24 | 617 | 13 | 3 | 1.646 | 1 | 0.392 | 0.00683 | 0.72999 | >0.10 |
| **NW Atlantic** | 32 | 617 | 10 | 10 | 2.246 | 10 | 0.555 | 0.00147 | **-1.97921** | **<0.05** |
| **Overall** | 121 | 617 | 27 | 22 | 3.073 | - | 0.675 | 0.00793 | -0.07822 | >0.10 |
| ***Alaria esculenta* COI-5P** | | | | | | | | | | |
| **Beaufort** | 3 | 473 | 0 | 1 | 1.000 | 0 | 0.000 | 0.00000 | - | - |
| **East Arctic** | 14 | 473 | 5 | 2 | 1.508 | 0 | 0.337 | 0.00383 | 0.53054 | >0.10 |
| **NW Atlantic** | 31 | 473 | 1 | 2 | 1.067 | 2 | 0.062 | 0.00014 | -1.14473 | >0.10 |
| **NE Atlantic** | 22 | 473 | 2 | 3 | 1.204 | 2 | 0.169 | 0.00038 | -1.51481 | >0.10 |
| **Overall** | 70 | 473 | 8 | 6 | 3.010 | - | 0.668 | 0.00421 | 0.51221 | >0.10 |
| ***Alaria esculenta* ITS** | | | | | | | | | | |
| **NW Pacific** | 2 | 518 | 1 | 2 | 2.000 | 0 | 0.500 | 0.00129 | 1.63299 | >0.10 |
| **Nome** | 23 | 518 | 1 | 3 | 1.924 | 1 | 0.480 | 0.00017 | -0.86025 | >0.10 |
| **Beaufort** | 2 | 518 | 0 | 1 | 1.000 | 0 | 0.000 | 0.00000 | - | - |
| **East Arctic** | 11 | 518 | 2 | 4 | 2.814 | 2 | 0.645 | 0.00134 | 0.59464 | >0.10 |
| **NW Atlantic** | 9 | 518 | 1 | 3 | 1.976 | 2 | 0.494 | 0.00071 | 0.48809 | >0.10 |
| **NE Atlantic** | 4 | 518 | 4 | 2 | 1.600 | 2 | 0.375 | 0.00332 | 0.48523 | >0.10 |
| **Overall** | 51 | 518 | 7 | 9 | 3.764 | - | 0.734 | 0.00146 | -1.03265 | >0.10 |
| ***Chaetopteris plumosa* COI-5P** | | | | | | | | | | |
| **Beaufort** | 3 | 544 | 0 | 1 | 1.000 | 0 | 0.000 | 0.00000 | - | - |
| **East Arctic** | 14 | 544 | 4 | 2 | 2.000 | 0 | 0.500 | 0.00396 | **2.34668** | **<0.05** |
| **NW Atlantic** | 12 | 544 | 0 | 1 | 1.000 | 0 | 0.000 | 0.00000 | - | - |
| **NE Atlantic** | 6 | 544 | 0 | 1 | 1.000 | 0 | 0.000 | 0.00000 | - | - |
| **Overall** | 35 | 544 | 4 | 2 | 1.690 | - | 0.408 | 0.00309 | 1.80157 | >0.05 |
| ***Chorda borealis* COI-5P** | | | | | | | | | | |
| **Nome** | 23 | 664 | 0 | 1 | 1.000 | 0 | 0.000 | 0.000 | - | - |
| **Location** | *n* | bp | $N_{poly}$ | Na | Ne | $N_{PH}$ | *h* | $\theta_{\pi}$ | *D* | *p* |

| | | | | | | | | | | |
|---|---|---|---|---|---|---|---|---|---|---|
| **East Arctic** | 14 | 664 | 0 | 1 | 1.000 | 0 | 0.000 | 0.000 | - | - |
| **NW Atlantic** | 2 | 664 | 0 | 1 | 1.000 | 0 | 0.000 | 0.000 | - | - |
| **Overall** | 39 | 664 | 0 | 1 | 1.000 | - | 0.000 | 0.000 | - | - |
| ***Chordaria chordaeformis* COI-5P** | | | | | | | | | | |
| **Nome** | 32 | 619 | 0 | 1 | 1.000 | 0 | 0.000 | 0.00000 | - | - |
| **East Arctic** | 38 | 619 | 1 | 2 | 1.054 | 1 | 0.051 | 0.00009 | -1.12863 | >0.10 |
| **NW Atlantic** | 1 | 619 | - | - | - | 0 | - | - | - | - |
| **Overall** | 71 | 619 | 1 | 2 | 1.029 | - | 0.028 | 0.00005 | -1.06579 | >0.10 |
| ***Chordaria flagelliformis* COI-5P** | | | | | | | | | | |
| **East Arctic** | 48 | 623 | 6 | 5 | 1.540 | 3 | 0.351 | 0.00074 | -1.68843 | >0.05 |
| **NW Atlantic** | 53 | 623 | 6 | 7 | 2.235 | 5 | 0.553 | 0.00170 | -0.49712 | >0.10 |
| **NE Atlantic** | 3 | 623 | 2 | 2 | 1.800 | 0 | 0.444 | 0.00214 | - | - |
| **Overall** | 104 | 623 | 10 | 10 | 1.994 | - | 0.499 | 0.00144 | -1.35439 | >0.10 |
| ***Desmarestia* sp. 1aculeata COI-5P** | | | | | | | | | | |
| **East Arctic** | 12 | 564 | 0 | 1 | 1.000 | 0 | 0.000 | 0.00000 | - | - |
| **NW Atlantic** | 21 | 564 | 1 | 2 | 1.100 | 1 | 0.091 | 0.00017 | -1.16356 | >0.10 |
| **NE Atlantic** | 9 | 564 | 1 | 2 | 1.246 | 1 | 0.198 | 0.00039 | -1.08823 | >0.10 |
| **Overall** | 42 | 564 | 2 | 3 | 1.101 | - | 0.092 | 0.00017 | -1.48214 | >0.10 |
| ***Eudesme borealis* COI-5P** | | | | | | | | | | |
| **NE Pacific** | 7 | 523 | 0 | 1 | 1.000 | 1 | 0.000 | 0.00000 | - | - |
| **Nome** | 11 | 523 | 1 | 2 | 1.658 | 1 | 0.397 | 0.00083 | 0.67135 | >0.10 |
| **East Arctic** | 11 | 523 | 7 | 3 | 2.283 | 0 | 0.562 | 0.00445 | -0.10637 | >0.10 |
| **NW Atlantic** | 12 | 523 | 4 | 4 | 2.057 | 2 | 0.514 | 0.00180 | -1.59840 | >0.05 |
| **NE Atlantic** | 2 | 523 | 0 | 1 | 1.000 | 1 | 0.000 | 0.00000 | - | - |
| **Overall** | 43 | 523 | 10 | 8 | 5.011 | - | 0.800 | 0.00721 | 1.44750 | >0.10 |
| ***Fucus distichus* COI-5P** | | | | | | | | | | |
| **NW Pacific** | 1 | 569 | - | - | - | 1 | - | - | - | - |
| **NE Pacific** | 52 | 569 | 3 | 3 | 1.570 | 1 | 0.363 | 0.00126 | 0.16907 | >0.10 |
| **Nome** | 20 | 569 | 2 | 2 | 1.980 | 1 | 0.495 | 0.00183 | 1.98958 | >0.05 |
| **East Arctic** | 12 | 569 | 1 | 2 | 2.000 | 0 | 0.500 | 0.00096 | 1.48617 | >0.10 |
| **NW Atlantic** | 32 | 569 | 3 | 4 | 1.213 | 2 | 0.176 | 0.00033 | -1.72954 | >0.05 |
| **NE Atlantic** | 2 | 569 | 0 | 1 | 1.000 | 0 | 0.000 | 0.00000 | - | - |
| **Overall** | 119 | 569 | 7 | 8 | 3.356 | - | 0.702 | 0.00173 | -0.56053 | >0.10 |
| ***Hedophyllum nigripes* COI-5P** | | | | | | | | | | |
| **NE Pacific** | 20 | 578 | 1 | 2 | 1.105 | 2 | 0.095 | 0.00017 | -1.16439 | >0.10 |
| **East Arctic** | 18 | 578 | 1 | 2 | 1.117 | 1 | 0.105 | 0.00019 | -1.16467 | >0.10 |
| **NW Atlantic** | 10 | 578 | 0 | 1 | 1.000 | 0 | 0.000 | 0.00000 | - | - |
| **Overall** | 48 | 578 | 3 | 4 | 2.110 | - | 0.526 | 0.00100 | -0.29683 | >0.10 |
| **Location** | *n* | bp | $N_{poly}$ | Na | Ne | $N_{PH}$ | *h* | $\theta_{\pi}$ | *D* | *p* |

| | | | | | | | | | | |
|---|---|---|---|---|---|---|---|---|---|---|
| ***Laminaria solidungula* COI-5P** | | | | | | | | | | |
| **Beaufort** | 7 | 603 | 0 | 1 | 1.000 | 1 | 0.000 | 0.00000 | - | - |
| | 12 | 603 | 1 | 2 | 1.180 | 1 | 0.153 | 0.00028 | -1.14053 | >0.10 |
| **NW Atlantic** | 1 | 603 | - | - | - | 0 | - | - | - | - |
| **Overall** | 20 | 603 | 2 | 3 | 2.062 | - | 0.515 | 0.001630 | 1.74879 | >0.05 |
| ***Petalonia fascia* COI-5P** | | | | | | | | | | |
| **NE Pacific** | 25 | 632 | 18 | 4 | 1.781 | 2 | 0.438 | 0.00684 | -0.50872 | >0.10 |
| **Nome** | 22 | 632 | 9 | 2 | 1.541 | 0 | 0.351 | 0.00524 | 1.14418 | >0.10 |
| **East Arctic** | 23 | 632 | 14 | 4 | 1.579 | 2 | 0.367 | 0.00216 | **-2.26245** | **<0.01** |
| **NW Atlantic** | 43 | 632 | 13 | 5 | 1.873 | 3 | 0.466 | 0.00149 | **-2.11431** | **<0.01** |
| **Overall** | 113 | 632 | 24 | 10 | 2.263 | - | 0.558 | 0.00580 | -0.65300 | >0.10 |
| ***Petalonia filliformis* COI-5P** | | | | | | | | | | |
| **East Arctic** | 42 | 658 | 5 | 4 | 2.641 | 2 | 0.621 | 0.00261 | 1.20875 | >0.10 |
| **NW Atlantic** | 18 | 658 | 3 | 3 | 1.906 | 1 | 0.475 | 0.00146 | 0.28158 | >0.10 |
| **NE Atlantic** | 1 | 658 | - | - | - | 1 | - | - | - | - |
| **Overall** | 61 | 658 | 7 | 6 | 2.512 | - | 0.602 | 0.00237 | 0.10495 | >0.10 |
| ***Pylaiella washingtoniensis* COI-5P** | | | | | | | | | | |
| **NE Pacific** | 16 | 656 | 7 | 7 | 2.415 | 6 | 0.586 | 0.00133 | **-2.06208** | **<0.05** |
| **Nome** | 16 | 656 | 1 | 2 | 1.280 | 1 | 0.219 | 0.00036 | -0.44832 | >0.10 |
| **Beaufort** | 2 | 656 | 0 | 1 | 1 | 0 | 0.000 | 0.00000 | - | - |
| **East Arctic** | 37 | 656 | 20 | 7 | 1.350 | 5 | 0.652 | 0.00667 | -0.29077 | >0.10 |
| **NW Atlantic** | 29 | 656 | 17 | 5 | 2.330 | 3 | 0.517 | 0.00243 | **-2.17855** | **<0.01** |
| **Overall** | 100 | 656 | 27 | 18 | 5.291 | - | 0.811 | 0.00506 | -1.09162 | >0.10 |
| ***Saccharina latissima* COI-5P** | | | | | | | | | | |
| **NE Pacific** | 14 | 614 | 4 | 4 | 1.581 | 3 | 0.367 | 0.00093 | **-1.79759** | **<0.05** |
| **Beaufort** | 7 | 614 | 0 | 1 | 1.000 | 0 | 0.000 | 0.00000 | - | - |
| **East Arctic** | 41 | 614 | 8 | 4 | 2.325 | 2 | 0.570 | 0.00520 | 2.01140 | >0.05 |
| **NW Atlantic** | 98 | 614 | 7 | 8 | 1.604 | 7 | 0.377 | 0.00067 | -1.64224 | >0.05 |
| **NE Atlantic** | 21 | 614 | 2 | 2 | 1.569 | 2 | 0.363 | 0.00124 | 0.85355 | >0.10 |
| **Overall** | 181 | 614 | 19 | 16 | 3.325 | - | 0.699 | 0.00541 | -0.10982 | >0.10 |
| ***Ulva fenestrata tufa*** | | | | | | | | | | |
| **NE Pacific** | 47 | 746 | 0 | 1 | 1.000 | 1 | 0.000 | 0.000 | - | - |
| **Nome** | 22 | 746 | 0 | 1 | 1.000 | 0 | 0.000 | 0.000 | - | - |
| **East Arctic** | 14 | 746 | 0 | 1 | 1.000 | 0 | 0.000 | 0.000 | - | - |
| **NW Atlantic** | 95 | 746 | 0 | 1 | 1.000 | 0 | 0.000 | 0.000 | - | - |
| **NE Atlantic** | 2 | 746 | 0 | 1 | 1.000 | 0 | 0.000 | 0.000 | - | - |
| **Overall** | 180 | 746 | 3 | 3 | 2.509 | - | 0.601 | 0.00153 | 1.95022 | >0.05 |

**Table S4.** Primers used for amplification of various genes in marine macroalgae. Primer sequence portions underlined and in bold type for M13LF3 and M13Rx forward and reverse primers, respectively, refer to sequencing primers (M13F and M13R).

| Locus | Taxa | Primers | Primer sequence (5'-3') | Reference/thermocycling regime |
|---|---|---|---|---|
| COI-5P | Red algae | M13LF3 | **TGTAAAACGACGGCCAGT**ACHAA YCAYAARGATATHGG | 4 |
| | | M13Rx | **CAGGAAACAGCTATGAC**ACTTCT GGRTGICCRAARAAYCA | |
| COI-5P | Brown algae | GazF2 | CCAACCAYAAAGATATWGGTAC3 | 5 |
| | | GazR2 | GGATGACCAAARAACCAAAA | |
| COI-5P | *Coccotylus* | GWSFn | TCAACAAAYCAYAAAGATATYGG3 | 4 |
| | | GWSRx | ACTTCTGGRTGICCRAARAAYCA | |
| *tufA* | Green algae | TufGF4 | GGNGCNGCNCAAATGGAYGG | 6 |
| | | TufAR | CCTTCNCGAATMGCRAAWCGC | |
| *rbc*L-3P | *Rhodochorton* | F57 | GTAATTCCATATGCTAAAATGGG | 4 |
| | | rbcLrevNEW | ACATTTGCTGTTGGAGTYTC | |
| *rbc*L-3p | *Battersia*, *Lithoderma* | L2 | AAAAGTGACCGTTATGAATC | 4 |
| | | L8 | CCAATAGTACCACCACCAAAT | |
| *rbc*L-3P | *Ulothrix* | GrbcLFi | TCTCARCCWTTYATGCGTTGG | 6 |
| | | 1385R | AATTCAAATTTAATTTCTTTCC | |
| ITS | *Coccotylus truncatus* | P1 | GGAAGGAGAAGTCGTAACAAGG | 4 |
| | | G4 | CTTTTCCTCCGCTTATTGATATG | |
| ITS | *Rhodomela sibirica* | RLycF1 | TAGGGGTACAGTGGTCTCAC | Newly developed primers. Thermocycling regime followed Saunders and Moore (4). |
| | | RLycR1 | GAATCATTCGTCCTAAACGTC | |
| *ycf35* | *Ahnfeltia borealis* | ycf35F1 | CTTGCGCTTTCGCGTCTTTCT | Newly developed. Thermocycling regime: 95ºC for 2 mins; 35 cycles of 93ºC for 1 min, 55ºC for 1 min, and 72ºC for 2 mins; 72ºC for 2 mins. |
| | | ycf35R1 | CGCTAGATTTAGGTTCTAGTG | |

**Table S5.** Pairwise $\Phi_{ST}$ values for species of macroalgae with Arctic populations. For $\Phi_{ST}$, values above the diagonal represent *p*-values, with significant results bolded. Populations with low sample sizes (n<10) are flagged with an asterisk. Populations with a single collection were removed from analyses.

| **Species\|Marker (# of haplotypes\|# of polymorphic sites)** | | | | | | |
|---|---|---|---|---|---|---|
| *Ahnfeltia borealis*\|COI-5P (3\|10) | | | | | | |
| Population pairwise $\Phi_{ST}$ | *n* | Northeast Pacific | Nome, Alaska | Beaufort, Alaska | East Arctic | Northwest Atlantic |
| Northeast Pacific* | 1 | --- | --- | --- | --- | --- |
| Nome, Alaska | 24 | --- | --- | 1.0 | 1.0 | **0.001*** |
| Beaufort, Alaska | 23 | --- | 0.000 | --- | 1.0 | **0.001*** |
| East Arctic | 21 | --- | 0.000 | 0.000 | --- | **<0.001*** |
| Northwest Atlantic | 2 | --- | 0.851* | 0.846* | 0.833* | --- |
| *Ahnfeltia borealis*\|*ycf-35* (6\|7) | | | | | | |
| Population pairwise $\Phi_{ST}$ | *n* | Northeast Pacific | Nome, Alaska | Beaufort, Alaska | East Arctic | Northwest Atlantic |
| Northeast Pacific* | 1 | --- | --- | --- | --- | --- |
| Nome, Alaska | 23 | --- | --- | **0.010** | **0.019** | **0.001*** |
| Beaufort, Alaska | 20 | --- | 0.256 | --- | 0.420 | **<0.001*** |
| East Arctic | 15 | --- | 0.209 | 0.020 | --- | **0.010*** |
| Northwest Atlantic | 2 | --- | 0.815* | 1.000* | 0.976* | --- |
| *Coccotylus truncatus*\|COI-5P (14\|18) | | | | | | |
| Population pairwise $\Phi_{ST}$ | *n* | Nome, Alaska | Beaufort, Alaska | East Arctic | Northwest Atlantic | Northeast Atlantic |
| Nome, Alaska | 12 | --- | **0.003** | **0.001** | **0.001** | --- |
| Beaufort, Alaska | 30 | 0.270 | --- | **0.001** | **0.001** | --- |
| East Arctic | 34 | 0.672 | 0.573 | --- | **0.001** | --- |
| Northwest Atlantic | 54 | 0.444 | 0.378 | 0.182 | --- | --- |

| | | | | | | |
|---|---|---|---|---|---|---|
| Northeast Atlantic* | 1 | --- | --- | --- | --- | --- |
| *Coccotylus truncatus*\|ITS (8\|8) | | | | | | |
| Population pairwise $\Phi_{ST}$ | *n* | Nome, Alaska | Beaufort, Alaska | East Arctic | Northwest Atlantic | |
| Nome, Alaska | 11 | --- | **0.001** | **0.001** | **0.002** | |
| Beaufort, Alaska | 30 | 0.697 | --- | **0.001** | **0.001** | |
| East Arctic | 25 | 0.571 | 0.739 | --- | **0.002** | |
| Northwest Atlantic | 35 | 0.293 | 0.640 | 0.173 | --- | |
| *Devaleraea ramentacea*\|COI-5P (7\|10) | | | | | | |
| Population pairwise $\Phi_{ST}$ | *n* | East Arctic | Northwest Atlantic | | | |
| East Arctic | 12 | --- | **0.001** | | | |
| Northwest Atlantic | 27 | 0.435 | --- | | | |
| *Devaleraea ramentacea*\|ITS (13\|28) | | | | | | |
| Population pairwise $\Phi_{ST}$ | *n* | East Arctic | Northwest Atlantic | | | |
| East Arctic | 12 | --- | **0.001** | | | |
| Northwest Atlantic | 20 | 0.346 | --- | | | |
| *Dilsea socialis*\|COI-5P (2\|1) | | | | | | |
| Population pairwise $\Phi_{ST}$ | *n* | Nome, Alaska | Beaufort, Alaska | East Arctic | Northwest Atlantic | |
| Nome, Alaska | 38 | --- | 0.428 | 0.332 | 0.448 | |
| Beaufort, Alaska | 28 | 0.000 | --- | 1.0 | 1.0 | |
| East Arctic | 21 | 0.000 | 0.000 | --- | 1.0 | |
| Northwest Atlantic | 47 | 0.006 | 0.000 | 0.000 | --- | |
| *Odonthalia dentata*\|COI-5P (11\|14) | | | | | | |

<table>
<tr><td>Population pairwise $\Phi_{ST}$</td><td>n</td><td>Nome, Alaska</td><td>Beaufort, Alaska</td><td>East Arctic</td><td>Northwest Atlantic</td><td>Northeast Atlantic</td></tr>
<tr><td>Nome, Alaska</td><td>21</td><td>---</td><td>0.010</td><td>0.001</td><td>0.001</td><td>0.001</td></tr>
<tr><td>Beaufort, Alaska</td><td>24</td><td>0.173</td><td>---</td><td>0.001</td><td>0.001</td><td>0.001</td></tr>
<tr><td>East Arctic</td><td>24</td><td>0.557</td><td>0.574</td><td>---</td><td>0.001</td><td>0.001</td></tr>
<tr><td>Northwest Atlantic</td><td>41</td><td>0.440</td><td>0.520</td><td>0.639</td><td>---</td><td>0.001</td></tr>
<tr><td>Northeast Atlantic</td><td>26</td><td>0.926</td><td>0.946</td><td>0.964</td><td>0.981</td><td>---</td></tr>
<tr><td colspan="7">Palmaria palmata|COI-5P (18|23)</td></tr>
<tr><td>Population pairwise $\Phi_{ST}$</td><td>n</td><td>East Arctic</td><td>Northwest Atlantic</td><td>Northeast Atlantic</td><td></td><td></td></tr>
<tr><td>East Arctic</td><td>12</td><td>---</td><td>0.418</td><td>0.001</td><td></td><td></td></tr>
<tr><td>Northwest Atlantic</td><td>48</td><td>0.000</td><td>---</td><td>0.001</td><td></td><td></td></tr>
<tr><td>Northeast Atlantic</td><td>19</td><td>0.908</td><td>0.915</td><td>---</td><td></td><td></td></tr>
<tr><td colspan="7">Phycodrys fimbriata|COI-5P (16|15)</td></tr>
<tr><td>Population pairwise $\Phi_{ST}$</td><td>n</td><td>Nome, Alaska</td><td>Beaufort, Alaska</td><td>East Arctic</td><td>Northwest Atlantic</td><td></td></tr>
<tr><td>Nome, Alaska</td><td>18</td><td>---</td><td>0.001</td><td>0.121</td><td>0.001</td><td></td></tr>
<tr><td>Beaufort, Alaska</td><td>32</td><td>0.250</td><td>---</td><td>0.007</td><td>0.001</td><td></td></tr>
<tr><td>East Arctic</td><td>22</td><td>0.060</td><td>0.097</td><td>---</td><td>0.001</td><td></td></tr>
<tr><td>Northwest Atlantic</td><td>69</td><td>0.709</td><td>0.773</td><td>0.635</td><td>---</td><td></td></tr>
<tr><td colspan="7">Rhodomela sibirica|COI-5P (4|5)</td></tr>
<tr><td>Population pairwise $\Phi_{ST}$</td><td>n</td><td>Nome, Alaska</td><td>Beaufort, Alaska</td><td>East Arctic</td><td></td><td></td></tr>
<tr><td>Nome, Alaska</td><td>23</td><td>---</td><td>0.002</td><td>0.001</td><td></td><td></td></tr>
<tr><td>Beaufort, Alaska</td><td>29</td><td>0.238</td><td>---</td><td>0.001</td><td></td><td></td></tr>
<tr><td>East Arctic</td><td>19</td><td>0.435</td><td>0.452</td><td>---</td><td></td><td></td></tr>
<tr><td colspan="7">Rhodomela sibirica|ITS (5|6)</td></tr>
</table>

| Population pairwise $\Phi_{ST}$ | *n* | Nome, Alaska | Beaufort, Alaska | East Arctic | |
|---|---|---|---|---|---|
| Nome, Alaska | 22 | --- | **0.001** | **0.001** | |
| Beaufort, Alaska | 27 | 0.877 | --- | **0.001** | |
| East Arctic | 15 | 0.414 | 0.959 | --- | |
| *Rhodomela virgata*\|COI-5P (5\|4) | | | | | |
| Population pairwise $\Phi_{ST}$ | *n* | Nome, Alaska | Beaufort, Alaska | East Arctic | Northwest Atlantic |
| Nome, Alaska | 37 | --- | --- | 0.695 | 0.204 |
| Beaufort, Alaska | 1 | --- | --- | --- | --- |
| East Arctic | 36 | 0.000 | --- | --- | 1.0 |
| Northwest Atlantic | 2 | 0.000 | --- | 0.000 | --- |
| *Rhodomela* sp. 1virgata\|COI-5P (8\|6) | | | | | |
| Population pairwise $\Phi_{ST}$ | *n* | Nome, Alaska | East Arctic | Northwest Atlantic | |
| Nome, Alaska | 14 | --- | **0.001** | 0.555* | |
| East Arctic | 17 | 0.446 | --- | **0.001*** | |
| Northwest Atlantic | 7 | 0.028* | 0.359* | --- | |
| *Scagelia pylaisaei*\|COI-5P (22\|27) | | | | | |
| Population pairwise $\Phi_{ST}$ | n | Northeast Pacific | Nome, Alaska | East Arctic | Northwest Atlantic |
| Northeast Pacific | 29 | --- | **0.001** | **0.014** | **0.001** |
| Nome, Alaska | 36 | 0.103 | --- | **0.004** | **0.001** |
| East Arctic | 24 | 0.133 | 0.211 | --- | **0.001** |
| Northwest Atlantic | 32 | 0.903 | 0.953 | 0.726 | --- |
| *Alaria esculenta*\|COI-5P (6\|8) | | | | | |

| Population pairwise $\Phi_{ST}$ | *n* | Beaufort, Alaska | East Arctic | Northwest Atlantic | Northeast Atlantic | | |
|---|---|---|---|---|---|---|---|
| Beaufort, Alaska | 3 | --- | 0.326* | **0.003*** | **0.004*** | | |
| East Arctic | 14 | 0.000* | --- | **0.001** | **0.001** | | |
| Northwest Atlantic | 30 | 0.980* | 0.770 | --- | **0.001** | | |
| Northeast Atlantic | 22 | 0.966* | 0.799 | 0.947 | --- | | |
| *Alaria esculenta*\|ITS (9\|11) | | | | | | | |
| Population pairwise $\Phi_{ST}$ | *n* | Northwest Pacific | Nome, Alaska | Beaufort, Alaska | East Arctic | Northwest Atlantic | Northeast Atlantic |
| Northwest Pacific | 2 | --- | 0.162* | 1.000* | 0.493* | 0.373* | 0.079* |
| Nome, Alaska | 23 | 0.117* | --- | 0.156* | **0.023** | **0.001*** | **0.001*** |
| Beaufort, Alaska | 2 | 0.000* | 0.000* | --- | 0.505* | **0.034*** | 0.068* |
| East Arctic | 11 | 0.000* | 0.161 | 0.013* | --- | **0.018*** | **0.002*** |
| Northwest Atlantic | 9 | 0.070* | 0.525* | 0.534* | 0.235* | --- | **0.001*** |
| Northeast Atlantic | 4 | 0.373* | 0.664* | 0.529* | 0.486* | 0.536* | --- |
| *Chaetopteris plumosa*\|COI-5P (2\|4) | | | | | | | |
| Population pairwise $\Phi_{ST}$ | *n* | Beaufort, Alaska | East Arctic | Northwest Atlantic | Northeast Atlantic | | |
| Beaufort, Alaska | 3 | --- | 0.210* | **0.001*** | **0.001*** | | |
| East Arctic | 14 | 0.250* | --- | **0.005** | **0.047*** | | |
| Northwest Atlantic | 12 | 1.000* | 0.438 | --- | 1.000* | | |
| Northeast Atlantic | 6 | 1.000* | 0.344* | 0.000* | --- | | |
| *Chorda borealis*\|COI-5P (1\|0) | | | | | | | |
| Population pairwise $\Phi_{ST}$ | *n* | Nome, Alaska | East Arctic | Northwest Atlantic | | | |
| Nome, Alaska | 23 | --- | 1.0 | 1.0* | | | |

| | | | | | | |
|---|---|---|---|---|---|---|
| East Arctic | 14 | 0.000 | --- | 1.0* | | |
| Northwest Atlantic | 2 | 0.000* | 0.000* | --- | | |
| *Chordaria chordaeformis*\|COI-5P (2\|1) | | | | | | |
| Population pairwise $\Phi_{ST}$ | *n* | Nome, Alaska | East Arctic | Northwest Atlantic | | |
| Nome, Alaska | 32 | --- | 1.0 | --- | | |
| East Arctic | 38 | 0.000 | --- | --- | | |
| Northwest Atlantic | 1 | --- | --- | --- | | |
| *Chordaria flagelliformis*\|COI-5P (10\|10) | | | | | | |
| Population pairwise $\Phi_{ST}$ | *n* | East Arctic | Northwest Atlantic | Northeast Atlantic | | |
| East Arctic | 48 | --- | **0.001** | 0.007* | | |
| Northwest Atlantic | 53 | 0.187 | --- | 0.349* | | |
| Northeast Atlantic | 3 | 0.607* | 0.059* | --- | | |
| *Desmarestia* sp. 1aculeata\|COI-5P (3\|2) | | | | | | |
| Population pairwise $\Phi_{ST}$ | *n* | East Arctic | Northwest Atlantic | Northeast Atlantic | | |
| East Arctic | 12 | --- | 0.369 | 0.439* | | |
| Northwest Atlantic | 21 | 0.000 | --- | 0.525* | | |
| Northeast Atlantic | 9 | 0.034* | 0.031* | --- | | |
| *Eudesme borealis*\|COI-5P (8\|10) | | | | | | |
| Population pairwise $\Phi_{ST}$ | *n* | Northeast Pacific | Nome, Alaska | East Arctic | Northwest Atlantic | Northeast Atlantic |
| Northeast Pacific | 7 | --- | **<0.001*** | **<0.001*** | **<0.001*** | **<0.001*** |
| Nome, Alaska | 11 | 0.793* | --- | **<0.001** | **<0.001** | **0.013*** |
| East Arctic | 11 | 0.770* | 0.742 | --- | **0.041** | 0.438* |
| Northwest Atlantic | 12 | 0.916* | 0.891 | 0.156 | --- | 0.045* |

<table>
<tr><td>Northeast Atlantic</td><td>2</td><td>1.000*</td><td>0.926*</td><td>0.021*</td><td>0.431*</td><td colspan="2">---</td></tr>
<tr><td colspan="8">Fucus distichus|COI-5P (8|7)</td></tr>
<tr><td>Population pairwise $\Phi_{ST}$</td><td>n</td><td>Northwest Pacific</td><td>Northeast Pacific</td><td>Nome, Alaska</td><td>East Arctic</td><td>Northwest Atlantic</td><td>Northeast Atlantic</td></tr>
<tr><td>Northwest Pacific</td><td>1</td><td>---</td><td>---</td><td>---</td><td>---</td><td>---</td><td>---</td></tr>
<tr><td>Northeast Pacific</td><td>52</td><td>---</td><td>---</td><td><0.001</td><td><0.001</td><td><0.001</td><td>0.054*</td></tr>
<tr><td>Nome, Alaska</td><td>20</td><td>---</td><td>0.350</td><td>---</td><td><0.001</td><td><0.001</td><td>0.396*</td></tr>
<tr><td>East Arctic</td><td>12</td><td>---</td><td>0.493</td><td>0.448</td><td>---</td><td>0.001</td><td>0.477*</td></tr>
<tr><td>Northwest Atlantic</td><td>32</td><td>---</td><td>0.554</td><td>0.487</td><td>0.422</td><td>---</td><td>0.169*</td></tr>
<tr><td>Northeast Atlantic</td><td>2</td><td>---</td><td>0.407*</td><td>0.190*</td><td>0.172*</td><td>0.000*</td><td>---</td></tr>
<tr><td colspan="8">Hedophyllum nigripes|COI-5P (4|3)</td></tr>
<tr><td>Population pairwise $\Phi_{ST}$</td><td>n</td><td>Northeast Pacific</td><td>East Arctic</td><td>Northwest Atlantic</td><td></td><td colspan="2"></td></tr>
<tr><td>Northeast Pacific</td><td>20</td><td>---</td><td><0.001</td><td><0.001*</td><td></td><td colspan="2"></td></tr>
<tr><td>East Arctic</td><td>18</td><td>0.905</td><td>---</td><td>0.359*</td><td></td><td colspan="2"></td></tr>
<tr><td>Northwest Atlantic</td><td>10</td><td>0.936*</td><td>0.000*</td><td>---</td><td></td><td colspan="2"></td></tr>
<tr><td colspan="8">Laminaria solidungula|COI-5P (3|2)</td></tr>
<tr><td>Population pairwise $\Phi_{ST}$</td><td>n</td><td>Beaufort, Alaska</td><td>East Arctic</td><td>Northwest Atlantic</td><td></td><td colspan="2"></td></tr>
<tr><td>Beaufort, Alaska</td><td>7</td><td>---</td><td><0.001*</td><td>---</td><td></td><td colspan="2"></td></tr>
<tr><td>East Arctic</td><td>12</td><td>0.944*</td><td>---</td><td>---</td><td></td><td colspan="2"></td></tr>
<tr><td>Northwest Atlantic</td><td>1</td><td>---</td><td>---</td><td>---</td><td></td><td colspan="2"></td></tr>
<tr><td colspan="8">Petalonia fascia|COI-5P (10|24)</td></tr>
<tr><td>Population pairwise $\Phi_{ST}$</td><td>n</td><td>Northeast Pacific</td><td>Nome, Alaska</td><td>East Arctic</td><td>Northwest Atlantic</td><td colspan="2"></td></tr>
<tr><td>Northeast Pacific</td><td>25</td><td>---</td><td><0.001</td><td>0.027</td><td>0.001</td><td colspan="2"></td></tr>
<tr><td>Nome, Alaska</td><td>22</td><td>0.370</td><td>---</td><td><0.001</td><td><0.001</td><td colspan="2"></td></tr>
</table>

| | | | | | | |
|---|---|---|---|---|---|---|
| East Arctic | 23 | 0.098 | 0.670 | --- | 0.176 | |
| Northwest Atlantic | 43 | 0.182 | 0.744 | 0.017 | --- | |
| *Petalonia filliformis*\|COI-5P (5\|6) | | | | | | |
| Population pairwise $\Phi_{ST}$ | *n* | East Arctic | Northwest Atlantic | Northeast Atlantic | | |
| East Arctic | 42 | --- | 0.125 | --- | | |
| Northwest Atlantic | 18 | 0.042 | --- | --- | | |
| Northeast Atlantic | 1 | --- | --- | --- | | |
| *Pylaiella washingtoniensis*\|COI-5P (18\|27) | | | | | | |
| Population pairwise $\Phi_{ST}$ | *n* | Northeast Pacific | Nome, Alaska | Beaufort, Alaska | East Arctic | Northwest Atlantic |
| Northeast Pacific | 16 | --- | 0.489 | 0.582* | **0.001** | **0.001** |
| Nome, Alaska | 16 | 0.015 | --- | 0.231* | **0.001** | **0.001** |
| Beaufort, Alaska | 2 | 0.000* | 0.000* | --- | 0.198* | 0.069* |
| East Arctic | 37 | 0.385 | 0.408 | 0.217* | --- | **0.001** |
| Northwest Atlantic | 29 | 0.583 | 0.623 | 0.496* | 0.157 | --- |
| *Saccharina latissima*\|COI-5P (16\|19) | | | | | | |
| Population pairwise $\Phi_{ST}$ | *n* | Northeast Pacific | Beaufort, Alaska | East Arctic | Northwest Atlantic | Northeast Atlantic |
| Northeast Pacific | 14 | --- | 0.406* | **0.005** | **<0.001** | **<0.001** |
| Beaufort, Alaska | 7 | 0.000* | --- | 0.082* | **<0.001*** | **<0.001*** |
| East Arctic | 41 | 0.241 | 0.201* | --- | **<0.001** | **<0.001** |
| Northwest Atlantic | 98 | 0.933 | 0.939* | 0.644 | --- | **<0.001** |
| Northeast Atlantic | 21 | 0.881 | 0.896* | 0.590 | 0.908 | --- |
| *Ulva fenestrata*\|*tufa* (3\|3) | | | | | | |
| Population pairwise $\Phi_{ST}$ | *n* | Northeast Pacific | Nome, Alaska | East Arctic | Northwest Atlantic | Northeast Atlantic |

| | | | | | | |
|---|---|---|---|---|---|---|
| Northeast Pacific | 47 | --- | **<0.001** | **<0.001** | **<0.001** | **<0.001*** |
| Nome, Alaska | 22 | 1.000 | --- | 1.000 | **<0.001** | **<0.001*** |
| East Arctic | 14 | 1.000 | 0.000 | --- | **<0.001** | **<0.001*** |
| Northwest Atlantic | 95 | 1.000 | 1.000 | 1.000 | --- | 1.000* |
| Northeast Atlantic | 2 | 1.000* | 1.000* | 1.000* | 0.000* | --- |
